# Macroevolutionary diversity of traits and genomes in the model yeast genus *Saccharomyces*

**DOI:** 10.1101/2022.03.30.486421

**Authors:** David Peris, Emily J. Ubbelohde, Meihua Christina Kuang, Jacek Kominek, Quinn K. Langdon, Marie Adams, Justin A. Koshalek, Amanda Beth Hulfachor, Dana A. Opulente, David J. Hall, Katie Hyma, Justin C. Fay, Jean-Baptiste Leducq, Guillaume Charron, Christian R. Landry, Diego Libkind, Carla Gonçalves, Paula Gonçalves, José Paulo Sampaio, Qi-Ming Wang, Feng-Yan Bai, Russel L. Wrobel, Chris Todd Hittinger

## Abstract

Species is the fundamental unit to quantify biodiversity. In recent years, the model yeast *Saccharomyces cerevisiae* has seen an increased number of studies related to its geographical distribution, population structure, and phenotypic diversity. However, seven additional species from the same genus have been less thoroughly studied, which has limited our understanding of the macroevolutionary leading to the diversification of this genus over the last 20 million years. Here, we report the geographies, hosts, substrates, and phylogenetic relationships for approximately 1,800 *Saccharomyces* strains, covering the complete genus with unprecedented breadth and depth. We generated and analyzed complete genome sequences of 163 strains and phenotyped 128 phylogenetically diverse strains. This dataset provides insights about genetic and phenotypic diversity within and between species and populations, quantifies reticulation and incomplete lineage sorting, and demonstrates how gene flow and selection have affected traits, such as galactose metabolism. These findings elevate the genus *Saccharomyces* as a model to understand biodiversity and evolution in microbial eukaryotes.

## Introduction

Global climate change is expected to significantly impact biodiversity and human health ^1^. Thus, it is increasingly important to catalog and understand the origins of biological diversity. While the species is the fundamental unit to quantify biodiversity from a biological perspective ^2^, the study of only one or a few representatives of each species biases our understanding of the true diversity of a species ^3^. This limitation is especially problematic when current species delineations are not in full agreement with the boundaries of gene flow or when traits vary widely within a species ^4^. Phenotypes can vary within a species or genus due to gene flow, selection, or other evolutionary processes ^5^. Thus, it is vital that the scientific community quantifies biodiversity and strives to understand both its ecological and evolutionary contexts.

Quantifying and understanding the origins of biodiversity will advance fundamental science while also identifying and prioritizing bioresources that contribute to food, medicine, fuels, and other value-added compounds^2^. Whole genome sequencing has empowered researcher’s in this endeavor, and ongoing initiatives, such as the Earth BioGenome Project and the European Reference Genome Atlas (ERGA), envision cataloging most of the individual species on Earth ^6, 7^. Unfortunately, these studies are particularly biased toward multicellular organisms, such as insects, vertebrates, and plants, for which multiple species have been identified, geographic patterns have been described, and phenotypic traits are often visible ^6^. In other species, such as microbial eukaryotes, macroevolutionary processes have been less thoroughly studied and received less attention for species- or genus-wide genome sequencing efforts. Nonetheless, microbial eukaryotes, such as yeasts, are great model organisms due to their small genomes, ease of genetic manipulation, and large number of genes that are orthologous with multicellular eukaryotes ^8^.

A major factor in the lack of quantification of eukaryotic microbes has been the influence of the hypothesis proposed by Baas Becking in 1934 and promulgated by Beijerinck that “everything is everywhere, but, the environment selects” ^9^. Nevertheless, expanded strain isolation from the wild and genome sequencing have shown that eukaryotic microbes, like multicellular organisms, also have geographical structure ^10, 11^. While large-scale whole genome sequencing studies have investigated the evolutionary history of the model yeast *Saccharomyces cerevisiae* and its closest relative, *Saccharomyces paradoxus* ^12–14^, the six other non-hybrid species of the genus *Saccharomyces* have been less thoroughly studied ^15–18^. In particular, several new and diverse lineages of *Saccharomyces* have recently been delineated ^13, 14, 19–28^, but the genetic and phenotypic diversities of each species have not been studied in a comparative context ^29^, which has limited our understanding of the macroevolutionary processes driving diversification in this important genus.

In this study, we cover the genetic and phenotypic diversity of the model eukaryotic genus *Saccharomyces* with unprecedented breadth and depth—reporting geographies, hosts, substrates, and phylogenetic relationships for approximately 1,800 *Saccharomyces* strains. We generate and analyze high-quality genome sequences for representative strains of all available phylogenetic lineages, and we sequence and phenotype more than a hundred *Saccharomyces* strains to quantify the genetic and phenotypic variation across this macroevolutionary timescale (13.3-19.3 million years ^30^). With this global dataset, we quantify diversity and divergence within and between species and populations, several types of natural reticulation events, and the influence of ecology and incomplete lineage sorting. This work elevates the genus *Saccharomyces* as a model for understanding biodiversity, population structure, and macroevolutionary processes in microbial eukaryotes. This fundamental understanding also provides a much needed framework for identifying and prioritizing key bioresources.

## Results

### *The Palearctic and Fagales preponderance of* Saccharomyces

To place newly isolated *Saccharomyces* strains in the context of existing datasets ^12, 13, 18, 23–25, 31–33^, we partially sequenced an additional 275 *COX2* and 129 *COX3* mitochondrial genes from key strains. In total, we analyzed the mitochondrial sequences of ∼1,800 *Saccharomyces* strains isolated mostly from bark substrates (52 % of wild isolates) from multiple continents (Figure 1A,C Figure S1, S2 and Table S1). Across the genus, 85 % of wild isolates were associated with the order Fagales, which includes oak and beech trees. In contrast, 89 % of *S. cerevisiae* strains analyzed here were isolated from anthropic environments (Figure 1C, Figure S2A).

**Figure 1.**
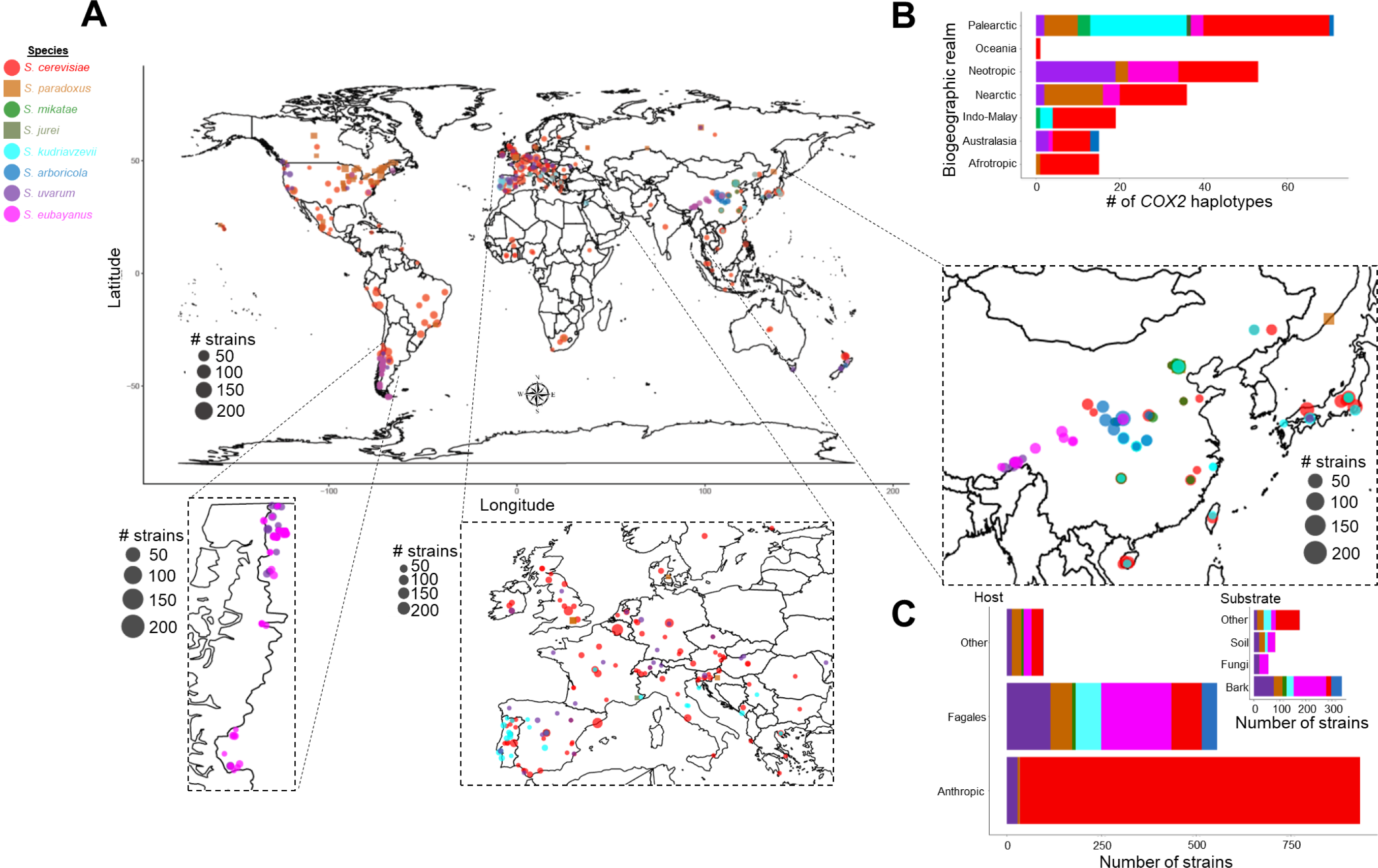
Geographic distribution of *Saccharomyces* strains. A) Map showing the locations where *Saccharomyces* strains have been isolated, scaled by size to the number of strains studied here. Symbols and colors designate the species. Ecological and geographic information about the strains can be found in Table S1. B) Stacked bar plot showing the number of *COX2* haplotypes isolated in each biogeographic realm (Figure 2A). The data shows many *COX2* haplotypes from the Palearctic region, pointing to Asia as a hotspot of diversity. Bars are colored by species. The map was generated using the `map_data` function implemented in `R` package `ggplot2`^99^. C) Bar plots represent the total number of strains from each *Saccharomyces* species grouped by host (external plot) or substrates (inner plot) (full details in Table S1 and Figure S2). Human-related environments, such as vineyards, were grouped in the “Anthropic” hosts category and removed from the substrate plot. Bar plots are colored according to species.

*Saccharomyces* mitochondrial genomes were highly polymorphic, with a large number of haplotypes inferred for *COX2* (Figure 1B, 2A) and *COX3* (Figure S3, Table S1). Our results indicate that the Palearctic biogeographic realm, which includes China and Europe, contained haplotypes from all species and more haplotypes than any other biogeographic realm (Figure 1B). The centrality of Palearctic *COX2* haplotypes in the phylogenetic network (Figure 2A) corroborates the hypothesis that many *Saccharomyces* lineages originated in this region, particularly East Asia ^25, 28, 34, 35^.

**Figure 2.**
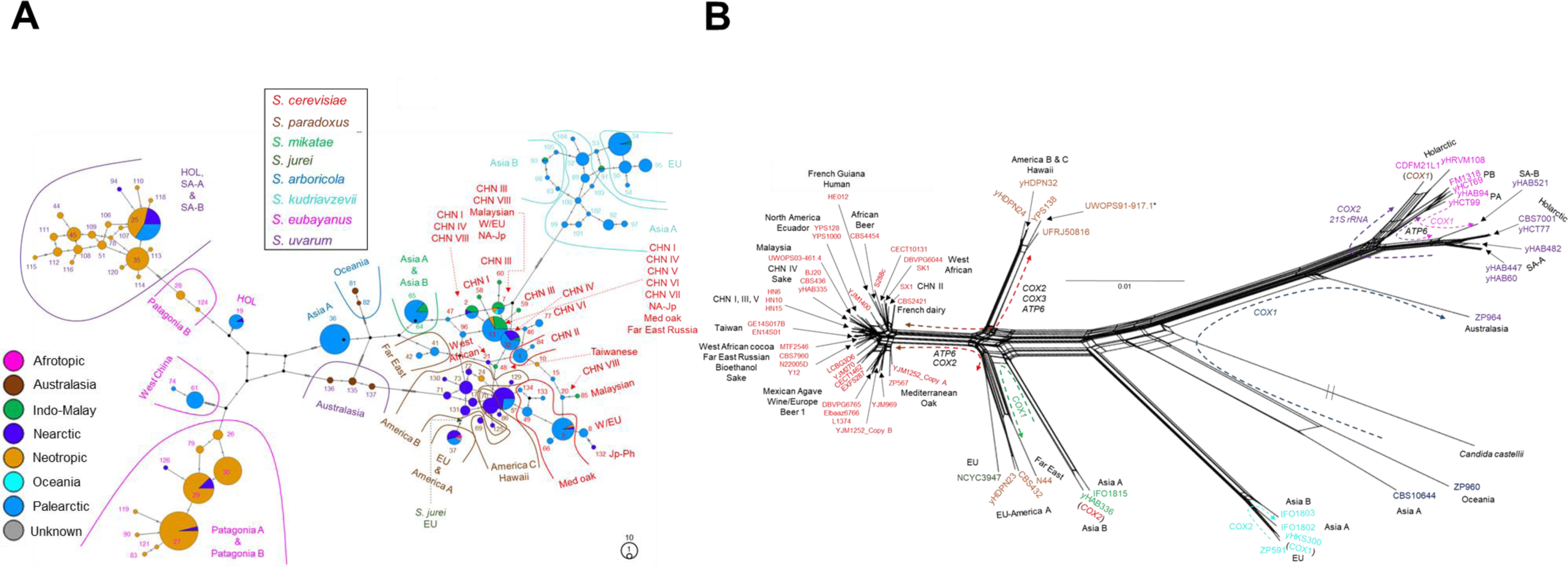
Extensive mitochondrial gene flow and introgression between *Saccharomyces* >lineages. A) Templeton, Crandall, and Sing (TCS) phylogenetic network of 739 partial *COX2* sequences from wild *Saccharomyces* strains. *COX2* haplotype classification, for the wild and anthropic *Saccharomyces* strains, is shown in Table S1. Haplotypes are represented by circles. Circle size is scaled according to the haplotype frequency. Pie charts show the frequency of haplotypes based on biogeographic realm. The number of mutations separating each haplotype are indicated by lines on the edges connecting different haplotype circles. Haplotype numbers and populations are highlighted in the panel and colored according to species designations. CHN: China; EU: Europe; HOL: Holarctic; Jp-Ph: Japan-Philippines (=Sake-Philippines); Med oak: Mediterranean oak; NA-Jp: North America-Japan (=North America); SA-A: South America A; SA-B: South America B; W/EU: Wine/European. B) Neighbor-Net phylogenetic network reconstructed using a concatenated alignment of the coding sequences of 10 mitochondrial genes (*ATP6*, *ATP8*, *ATP9*, *COB*, *COX1*, *COX2*, *COX3*, *VAR1*, and the genes encoding 15S rRNA and 21S rRNA) for 64 sequenced *Saccharomyces* strains representing all known *Saccharomyces* lineages that were available (Table S2). Strain names are colored according to species designations. Population names are highlighted in black. The scale is given in nucleotide substitution per site. Arrows highlight mitochondrial gene flow (intraspecies) and introgressions (interspecies) detected from individual gene trees (Figure S4); affected genes are shown close to the arrows with the color indicated by the species donor. Gene flow and introgressions unique to a *Saccharomyces* strain are indicated between parentheses. A similar phylogenetic network for the *COX3* mitochondrial gene is shown in Figure S3, which is more congruent with the concatenated data shown in panel B than the data for *COX2* shown in panel A. The asterisk indicates that UWOPS91-917.1 did not contain the introgression of *COX3* from *S. cerevisiae* found in other *Saccharomyces paradoxus* America B and C strains. Most of the *Saccharomyces jurei* (NCYC3947) protein-coding sequences were more closely related to the *S. paradoxus* Far East-EU clade, rather than to *Saccharomyces mikatae* (Figure S4).

### Genomic structural variation is common between Saccharomyces lineages

From our global *Saccharomyces* collection, we sequenced and assembled 22 high-quality genomes, including representatives for each major phylogenetic lineage (Table S2); these assemblies had nearly complete chromosomes with additional unplaced scaffolds ranging from 0 to 39 (Table S2). We also included 16 previously published assemblies, one of which we substantially improved, bringing the total here to 38 high-quality genome assemblies (Table S2). In addition, we generated sixteen complete mitochondrial genome assemblies, corrected the size of the previously published *Saccharomyces jurei* mitochondrial genome ^18^, and assembled two new 2-µm plasmids (Table S2). Structurally, species varied by GC contents, chromosome lengths, mitochondrial genome sizes, and the synteny of nuclear and mitochondrial genomes, usually due to a modest number of translocations (Figure S5-S8, Supplementary Note 1).

### *Analyses revealed new* Saccharomyces *lineages*

To better illuminate population-level diversity, especially for previously under-sampled species, 163 sequenced *Saccharomyces* strains were analyzed using several population and phylogenomic approaches (Table S2, see Online Material & Methods). Our analyses revealed new lineages of *S. kudriavzevii* and of *S. mikatae* (Figure S9C,D); we consider yeast lineages to be clades of strains with shared ancestries that have frequently interbred, even though they are not strictly panmictic populations. Two *S. kudriavzevii* strains, originally isolated in China, belonged to a newly identified lineage (Figure S9D), but they had fewer fixed differences compared to European (EU) strains (5.5 thousand SNPs) than to strains from the Asia A lineage (10.2 thousand SNPs). In haplotype and phylogenetic networks, mitochondrial gene sequences for these two strains were located between Asia A and EU haplotypes or unexpectedly close to Asia A (Figure 2A, S3, S4B,E). Interestingly, despite the geographic proximity of this lineage to Asia A, only ∼12 % of the nuclear genome of these strains was more divergent from EU than from the Asia A *S. kudriavzevii* population (Table S3, Figure S9D, S10Hi-ii), suggesting that these strains are descendants of an ancestral admixture event. Specifically, large portions of their genome are most closely related to EU (∼87 %), and small portions most closely related to Asia A (∼12 %). Two distinct populations were revealed for *S. mikatae*, one of which (Asia A) may have up to three cryptic lineages and a large number of segregating polymorphisms (Figure S9C), possibly from lineages yet to be discovered.

### *Differentiation and divergence of* Saccharomyces *lineages and species*

Studying all *Saccharomyces* species together, we inferred two or more populations, with an average of about 3 populations per species (Figure 3, Figure S9), except for *S. cerevisiae*, due partly to its multiple domestication events. *S. cerevisiae*, with 16 or more populations and extensive admixture ^13, 19, 25, 36, 37^, had relatively low genetic diversity compared to other species, with an average genetic distance only slightly higher than *S. mikatae* (Figure 3C, S11I). Despite the low sequence diversity, phenotypic and ecological factors better differentiated *S. cerevisiae* into distinct lineages or populations than in the other *Saccharomyces* species (Figure S9A). In contrast, *Saccharomyces paradoxus* was the most diverse species (1.95 % average pairwise divergence), followed by *S. kudriavzevii* and *S. uvarum* (Figure 3C, S11I). *Saccharomyces eubayanus* likely has diversity levels similar to *S. uvarum* ^24^, but the Sichuan and West Asia lineages ^22^ were not available for genome sequencing. Each species was separated from its closest relative by a genetic divergence of ∼10 % (Figure S11A-D,G-H), except for *S. arboricola* and *S. kudriavzevii* (Figure S11E,F). The differentiation among *S. kudriavzevii*, *S. arboricola,* and *S. paradoxus*, as measured by FST, was considerably lower than among the other *Saccharomyces* species (Figure S12), an indication that these three species harbor more variation that is not fixed between other members of the genus.

**Figure 3.**
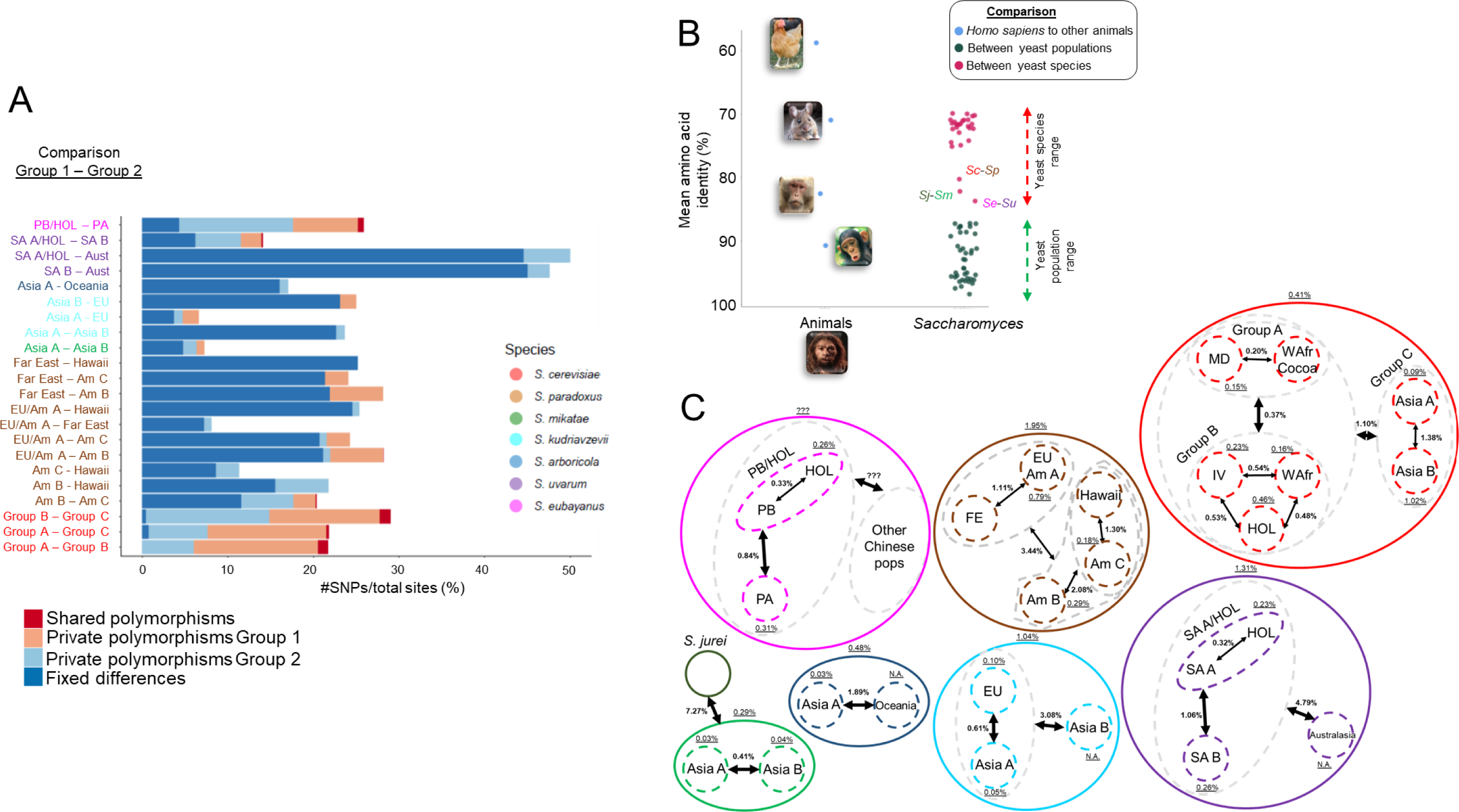
Species and population-level diversity in *Saccharomyces*. A) Percentage of private segregating polymorphisms, fixed differences, and shared polymorphisms among SNPs found in pairwise comparisons between supported populations, except for *S. cerevisiae* where populations were grouped according to PCA and co-ancestry for better resolution (Figure S9A iv and v). B) Right: dot plot of mean amino acid identities (AAI) calculated from pairwise comparisons between populations and between species. Left: dot plot for comparisons of *Homo sapiens* with *Pan troglodytes*, *Macaca mulatta*, *Mus musculus*, and *Gallus gallus*. C) Global picture of the percentage of the Tamura-Nei-corrected pairwise genetic distance between populations and within *Saccharomyces* species. ???: values cannot be inferred because West China and Sichuan strains were unavailable for whole genome sequencing. N. A.: not applicable because only one strain was available from this population. Am: America; EU: European; FE: Far East; HOL: Holarctic; MD: Mediterranean Domesticated group; SA-A: South America A; SA-B: South America B; *Sc*: *S. cerevisiae*; *Se*: *S. eubayanus*; *Sj*: *S. jurei*; *Sm*: *S. mikatae*; *Sp*: *S. paradoxus*; WAfr: West African; IV: China IV.

In *Saccharomyces*, levels of <85 % of amino acid identity (AAI) in a set of core single-copy eukaryotic genes differentiated species, while population-level AAI values were higher (Figure 3B). The lowest AAI value within a species was the comparison between the Asia B and EU populations of *S. kudriavzevii*, whose value was between the AAI values of the *Homo sapiens/Pan troglodytes* and *Homo sapiens*/*Macaca mulatta* comparisons. *Saccharomyces paradoxus* America A versus EU produced the highest AAI value (Figure 3B), which is consistent with the hypothesis that these populations were very recently derived due to migration from Europe to North America ^38^. The minimum AAI between *Saccharomyces* species was comparable to the comparison between *Homo sapiens* and *Mus musculus* (<70 % AAI).

### The non-nuclear genome is more permeable to introgression and gene flow than the nuclear genome

To explore the stability of the relationships among *Saccharomyces* populations and species, we analyzed 38 high-quality nuclear genomes of representative strains using a phylogenomic framework to investigate 3850 conserved genes. The `ASTRAL` coalescent species tree and `BUCKy` concordance primary tree agreed with previous studies (Figure 4A, Figure S13) ^15, 18, 28, 39^. Species-level branches were highly supported, while some branches close to the tips were not. Internal branch support values decreased outside of the *S. cerevisiae*-*S. paradoxus* clade and the *S. uvarum*-*S. eubayanus* clade, a phenomenon previously observed ^30, 40^ and proposed to be due to hybridization involving ancestors of *S. kudriavzevii* ^41^. Alternatively, the short coalescent units near the divergence of *S. arboricola* and *S. kudriavzevii* (Figure 4A) and the low relative differentiation of *S. arboricola* and *S. kudriavzevii* with the rest of species (Figure S12E,F) suggest a more nuanced model. Specifically, we propose that the conflicting data between genes are the result of diversification over a relatively narrow window of time, which allowed for the retention of considerable ancestral polymorphisms through incomplete lineage sorting (ILS); ancient gene flow between lineages in the early stages of speciation; or both. These patterns have been seen frequently across the tree of life ^42^.

**Figure 4.**
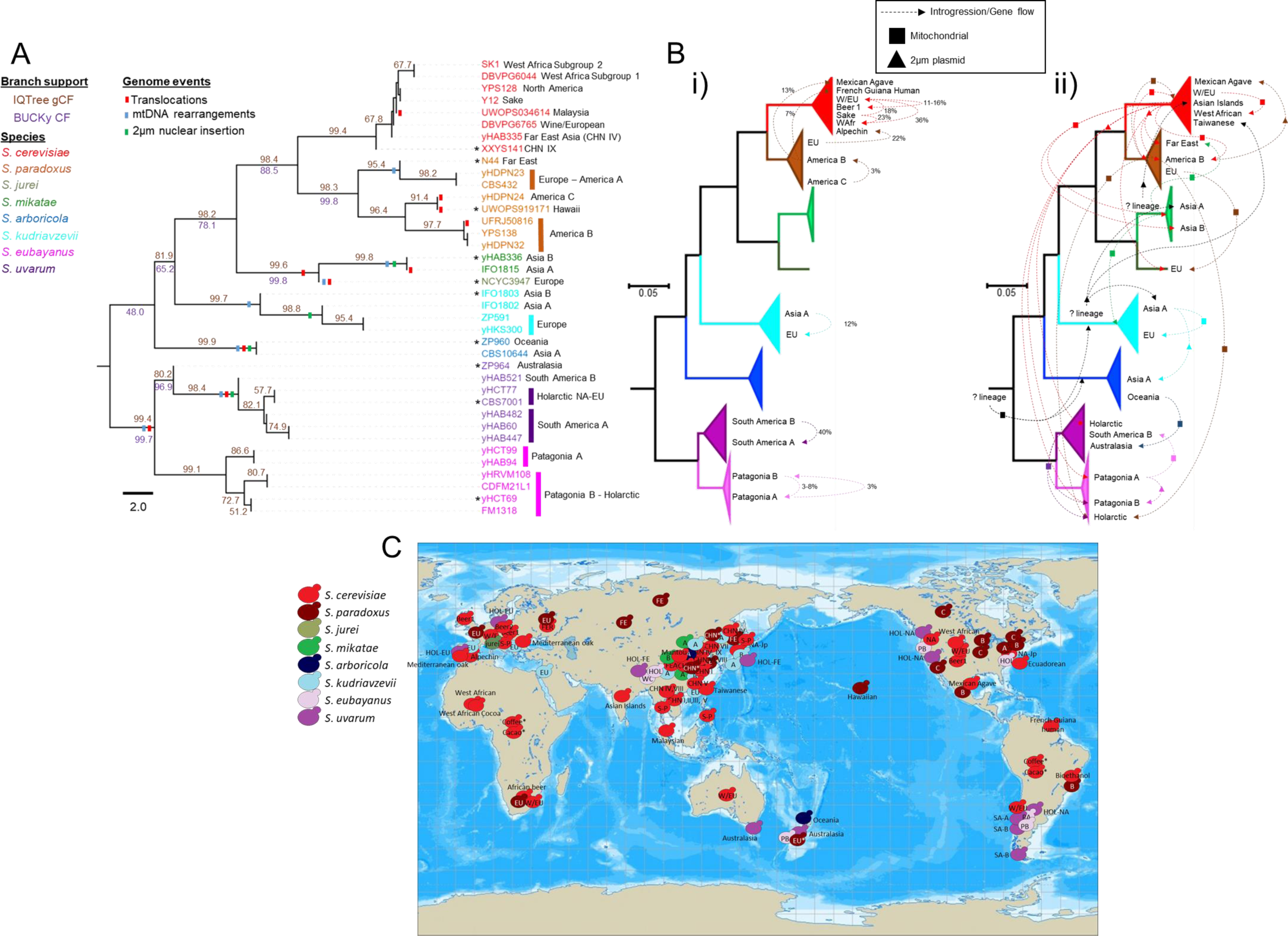
Vertical inheritance and incomplete lineage sorting dominated in the nuclear genome, while introgression and gene flow were widespread among cytoplasmically inherited genetic elements. A) Coalescent tree (species tree) for *Saccharomyces* lineages. Two values of concordance factors (CFs) are shown. Brown CFs were generated by `IQTree` using a collection of Maximum Likelihood phylogenetic trees (3850 genes) and the `ASTRAL` species tree. The normalized score was 0.97, which indicates that 97 % of input gene quartet trees are satisfied by the `ASTRAL` species tree. Purple CFs were generated by `BUCKy` using a collection of sample trees during Bayesian reconstruction in `MrBayes` and representative strains, mostly selected from Asia (asterisks). Other gene tree topologies are shown in Figure S13. Chromosomal translocations (Figure S5) and mitochondrial rearrangements (Figure S7,S8) are reported by red and blue bars on branches, respectively. The insertion of a 2-µm plasmid gene into the nuclear genome (Table S4) is represented by green bars on branches. The scale is coalescent units. B) Maximum-likelihood phylogenetic tree of all studied *Saccharomyces* strains reconstructed using the common `BUSCO` genes and collapsed to the species level (full tree in Figure 5B). Scale bars show the number of substitutions per site. Population names are only shown for those involved in gene flow or introgression based on the genome-wide analysis. B i) Summary of detected nuclear gene flow (between populations) and introgression (between species). The quantified percent of genome contribution by the donor is indicated near to the dashed arrow. *Saccharomyces cerevisiae* introgressions were congruent with previous reports ^13, 19, 64^. B ii) summary of detected gene flow and introgression for the mitochondrial genome (squared symbol) and 2-µm plasmid (triangle symbol). The direction of the arrow indicates the donor lineage. Unknown donor lineages are colored in black. Strain names, branches, and arrows are colored according to the species designations or their donors. C) Geographic locations of the different *Saccharomyces* populations. We omitted the global distribution of Wine/European *S. cerevisiae* population for clarity. The location of populations, for which strains were not studied here, are indicated with an asterisk symbol. Species-specific populations are colored according to the left legend.

To further explore the phylogenetic stability of species boundaries, we applied reciprocal monophyly tests for each species using 3850 ML gene trees (Table S5). *Saccharomyces cerevisiae* and *S. paradoxus* only failed to be monophyletic in 17 and 57 genes, respectively. Gene flow from *S. cerevisiae* to *S. paradoxus* EU and America A were detected, as previously documented ^43^, but the most frequent source of conflict was the location of the *S. cerevisiae* CHNIX lineage. This lineage sometimes grouped as an early-diverging member of the *S. paradoxus* clade or as an outgroup to both *S. cerevisiae* and *S. paradoxus*, topologies and branch lengths that are consistent with ILS. The *S. uvarum* Australasian lineage produced an even more striking pattern, again consistent with ILS, where more than 700 genes placed it as an early-diverging lineage of the *S. eubayanus* clade. At the species level, the Bayesian pipeline revealed many genes that supported alternative topologies, especially where the phylogenetic locations of *S. arboricola, S. kudriavzevii*, and the *S. mikatae/S. jurei* clade varied, and the consensus species tree was only supported by ∼1824 genes (48 % of a total of 3801 genes for this pipeline) (Figure S13). The presence of *Kluyveromyces lactis* in the dataset for the Bayesian pipeline, which was necessary to root the tree during phylogenetic reconstruction, might have decreased the support for internal branches compared with the ML pipeline (Figure 4A).

This conflict can be recapitulated using phylogenetic networks reconstructed using genes in 38 high-quality genomes (Table S2, Figure S14A) annotated with the Yeast Genome Annotation Pipeline (`YGAP`) and using 14 `BUSCO` genes common to all (160 strains) phenotyped and previously sequenced strains (Table S2, Figure S14B). Collectively, these results support a model of rapid radiation of some lineages with the retention of ancestral polymorphisms.

Within species, we observed much lower concordance factors at nodes (Figure 4A), which highlights ongoing gene flow within and between lineages. We next examined our sequenced and phenotyped strains (Table S2) for genome-wide signals of gene flow between recognized lineages (Figure S10). These analyses suggested that nuclear gene flow was infrequent. Only 9.25 % of the *Saccharomyces* strains, from five of the eight species, showed strong evidence of admixture (Figure S10, Table S3). Admixture was mostly observed in domesticated *S. cerevisiae* strains and was accompanied by higher levels of heterozygosity, which was generally low across the genus (Figure S15). The genomic contributions of the minor parental donor averaged 14.29 % (Figure 4Bi, Table S3). The smallest values belonged to two strains of *S. paradoxus* America C with contributions from America B, which were previously named the America C* lineage ^14^, as well as two *S. eubayanus* strains. In the latter cases, one strain was from each Patagonian population, but it had genomic contributions from the other Patagonian population. The highest value of genomic contribution by a minor donor in our dataset was found in a South America B strain, which had 39.53 % of its genome from South America A origin (Figure 4Bi, Figure S10I). This strain also showed one of the two highest levels of heterozygosity for wild strains (Figure S15), further suggesting a recent admixture event. The low levels of heterozygosity for the rest of admixed strains might point to the rapid fixation of lineage-specific alleles following haploselfing, intratetrad mating, or a return-to-growth event. Although we found some evidence of gene flow between populations, rarer introgressions between species (Figure S16, Table S3, ^13^), and considerable evidence of incomplete lineage sorting, we conclude that the phylogenies of nuclear genes were generally consistent with the accepted species relationships.

We next tested how the species tree compared with phylogenies generated using the mitochondrial genome. A preliminary view of mitochondrial synteny among *Saccharomyces* immediately suggested the possibility of considerable incongruence. For example, mitochondrial genome synteny is conserved in *S. cerevisiae* and *S. paradoxus*, except in the EU–America A and Far East populations of *S. paradoxus* (Figure S7, S8A, ^44^). The *S. jurei* nuclear genome was mostly syntenic with *S. mikatae* strains (Figure S5), but its mitochondrial genome was syntenic with the *S. paradoxus* EU and America A populations (Figure S8B) and differed from the *S. mikatae* Asia A population (Figure S7).

The *S. uvarum* Australasian population and *S. eubayanus* were syntenic in both their nuclear and mitochondrial genomes (Figure S5, S8E), while the other *S. uvarum* populations inherited derived mitochondrial and nuclear rearrangements (Figure S5, S8D). At the nucleotide level, both *COX2* and *COX3* phylogenetic networks disagreed with the nuclear genome in some cases. In both mitochondrial phylogenetic networks, population haplotypes from some species were more closely related to other species haplotypes than to their same-species haplotypes (Figure 2A, S3) due to lineage-specific introgressions. For example, *S. paradoxus* America B and C strains were connected to *S. cerevisiae* haplotypes. Similarly, *S. eubayanus* West China and *S. uvarum* Australasian strains likely experienced introgressions. A phylogenetic network for mitochondrial genes of the 64 high-quality mitochondrial genomes (Table S2, Figure 2B), supported the broader *COX2* and *COX3* results (Figure 2B, Figure S4). In addition to previously detected mitochondrial introgressions between species and gene flow between populations ^44–47^, we also detected new cases of mitochondrial introgressions and gene flow for *S. kudriavzevii*, *S. jurei,* and *S. mikatae* (Figure 4Bii, Figure S4). The *S. arboricola* and *S. kudriavzevii* mitochondrial genomes also had some affinity for the *Candida (Nakaseomyces) castellii* outgroup, as suggested by their exacerbated subtended edges in the network (Figure S4), so ancestral polymorphisms or introgression from unknown *Saccharomyces* lineages might have contributed to the mitochondrial genomes of these species. We conclude that events of introgressions and gene flow between mitochondrial genomes have been much more frequent than in the nuclear genome (Figure 4Bi, Bii).

Similarly, 22 interspecies transfers were detected for the 2-µm plasmid (Figure 4Bii, Figure S17), which is also cytoplasmically inherited. The *S. cerevisiae* 2-µm plasmid seems to be highly mobile, and we detected it in four other species. Sixteen strains had both cytoplasmic 2-µm plasmid genes and plasmid genes that had been transferred to the nuclear genome, a phenomenon previously noted for a handful of strains ^48^ (Table S4). We also detected a transfer from a hypothesized unknown source into the *S. cerevisiae* Taiwanese lineage ^13^, as well as to a *S. mikatae* Asia A strain and a *S. kudriavzevii* Asia A strain (Figure S17A). Given its sister relationship with the previously detected *S. kudriavzevii* 2-µm plasmid, this unknown lineage may also be a close relative of *S. kudriavzevii* (Figure S17A). Taken together, our results suggest that introgressions and gene flow involving the nuclear genome are limited in wild environments, while introgression and gene flow involving the cytoplasmically inherited mitochondrial genome and the 2-µm plasmid are much more frequent (Figure 4), likely because they can occur without involving karyogamy ^49^, or be aided by the activity of free-standing homing endonucleases ^47, 50^.

### Complex ancestries promote phenotypic diversity

To explore phenotypic variation across the genus *Saccharomyces*, we phenotyped 128 of the sequenced *Saccharomyces* strains, focusing on phylogenetically distinct lineages from different species (Table S2, S6, Figure S9). We tested the ability of these strains to grow in different carbon sources, temperatures, and stresses (Supplementary Note 2). Growth characteristics varied among *Saccharomyces* species depending on the conditions tested (Figure S18-S22). Interestingly, *S. mikatae* had some of the lowest genetic diversity values but had some of the highest phenotypic diversity (Figure 3C, 5A, Figure S23). In contrast, *S. eubayanus* and *S. uvarum* strains were mostly overlapping in a principal component analysis (PCA) and were less phenotypically diverse than the other species (Figure 5A, S23), indicating strains from these sister species have similar traits in the conditions tested (Figure 5A, Figure S24A,C). These results highlight how phenotypically diverse the *Saccharomyces* genus is and offer new bioresources for industrial applications.

**Figure 5.**
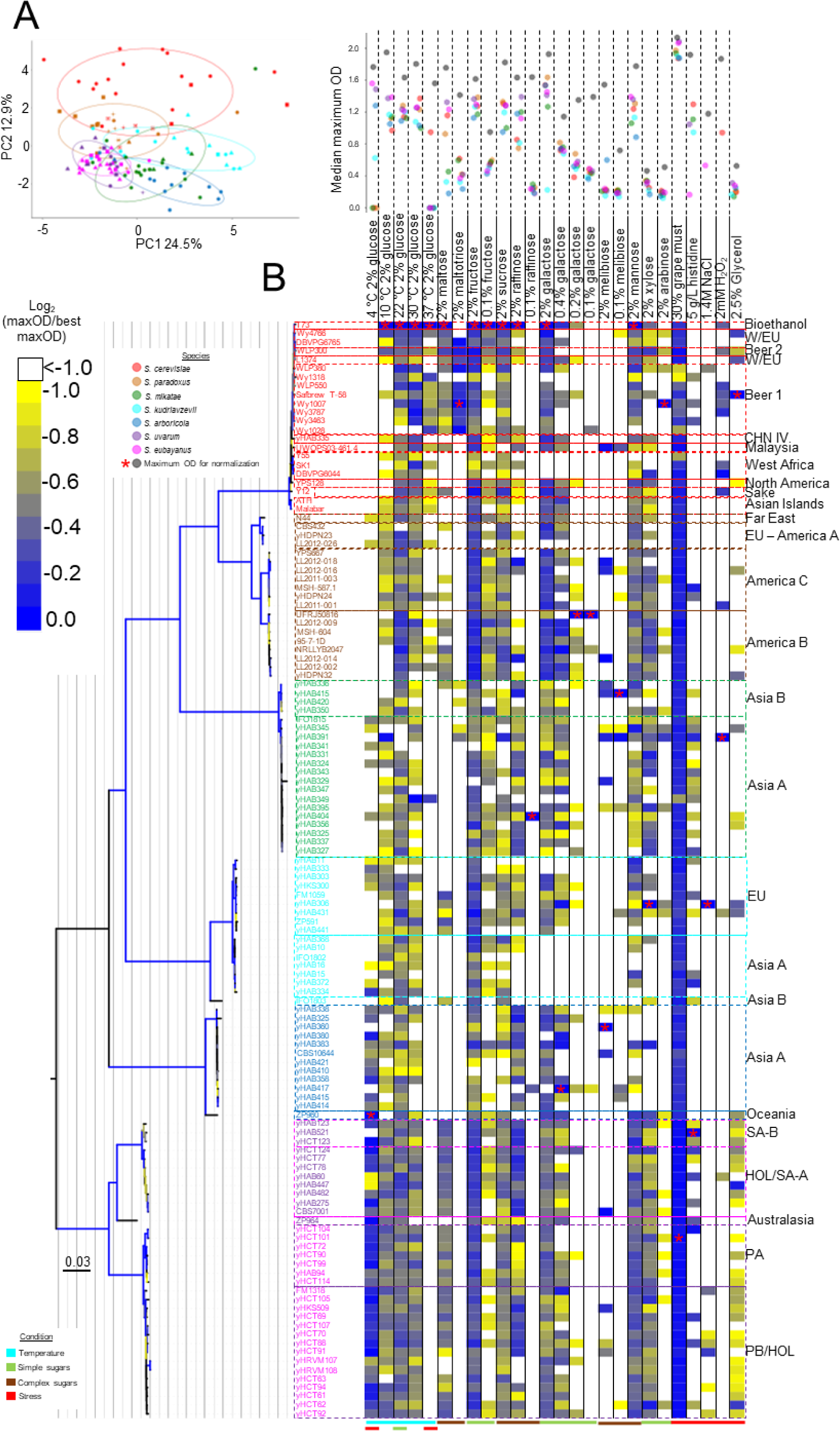
The genus *Saccharomyces* is phenotypically diverse. A) Principal component analysis (PCA) of PC1 and PC2 of the maximum OD600 calculated from growth curves (n = 3) calculated from an array of twenty-six media conditions (Table S6). PC1 and PC2 accounted for 37.4 % of the total variation. A higher image resolution PCA with growth condition weights can be found in Figure S24A. The variation explained by each component is shown in Figure S24B, and a plot of PC1 and PC3 is shown in Figure S24C. Strains are colored according to their species designations, and different shapes represent their population or lineage designation (see below). B) Heatmap showing the maximum OD600, normalized by the highest value for each growth condition as indicated by a red asterisk. Heat colors from yellow (low growth) to blue (high growth) are scaled according to the bar in the left. White colors indicate log2 values lower than -1 or no detected growth. Growth conditions are columns, and strains are rows. The dot plot above the growth conditions shows the maximum OD600 value used for normalizing the data for each growth condition (grey dot), and the colored dots are median maximum OD600 value for each *Saccharomyces* species. A maximum-likelihood (ML) phylogenetic tree of 14 orthologs (∼8.7 Kbp) for the phenotyped strains is shown to the left of the heatmap. Branches are colored according to their bootstrap support (minimum, yellow – maximum, dark blue). Strain names are colored according to species designations. Population designations are written to the right of the heatmap. The bottom colored bars highlight the conditions tested: temperature, simple or complex sugars, and stress. CHN: China; EU: Europe; HOL/SA-A: Holarctic/South America A; PA: Patagonia A; PB/HOL: Patagonia B/Holarctic; SA-B: South America B. iTOL tree at http://bit.ly/2VthpGT.

Temperature tolerance was an important condition (Figure S25 S26) for species differentiation (Figure 5A). *Saccharomyces eubayanus* and *S. uvarum* grew the best at lower temperatures (Figure 5B, S18, S26C-E), while *S. cerevisiae* and *S. paradoxus* grew the worst at lower temperatures and instead grew best at higher temperatures (Figure 5B, S26C-E). *Saccharomyces mikatae*, *S. arboricola,* and *S. kudriavzevii* also grew well at lower temperatures, which supports the hypothesis that lower temperature growth is an ancestral trait of the genus *Saccharomyces* ^51, 52^ and might influence in the ecological and geographic distribution of *Saccharomyces* lineages.

The utilization pathway for the sugars *GAL*actose and *MEL*ibiose is well studied and highly variable in the genus *Saccharomyces* (Figure S27A) ^53–56^. Making use of our diverse genomic and phenotypic dataset, we explored the ancestry of the individual genes involved in the *GAL*/*MEL* pathway (Figure S28) to determine potential genetic bases of variabilities in growth on galactose and melibiose (Figure 6A,B, S27B,C). Previous studies have observed loss-of-function mutations in some genes of the pathway in *S. cerevisiae* ^56, 57^, ancient pseudogenization of the entire *GAL* pathway in the *S. kudriavzevii* Asia A and B populations and retention of a functional pathway in the EU population ^58, 59^, and ancient alleles in some *S. cerevisiae* strains whose origin predates the diversification of the genus ^60–63^. Our new analyses here found additional variation that suggests that some of the variation in galactose or melibiose growth was the consequence of gene flow between populations of the same species or introgression between species (Figure 6A, S27B, S28). For example, two strains of *S. paradoxus* from America C with evidence of gene flow from America B population (Figure S9G) were capable of growing on melibiose, likely because they acquired an active *MEL1* gene from the America B population (Figure S27C, S28H). Introgressions for genes conferring melibiose utilization were also detected between *S. cerevisiae* and *S. paradoxus* ^55, 57^.

**Figure 6.**
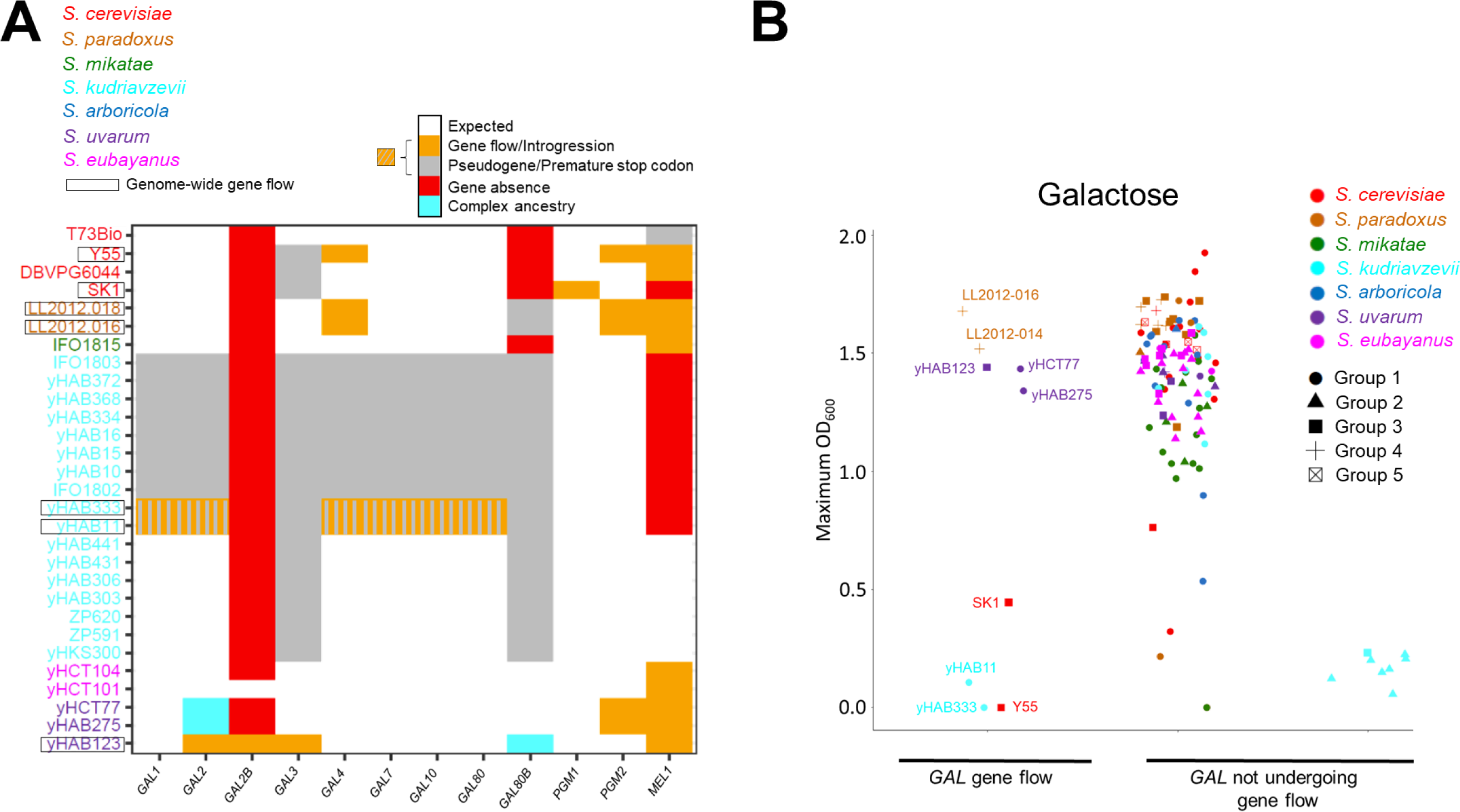
Phenotypic diversity and complex ancestries. A) *Saccharomyces* strains affected by gene-flow for the *GAL* regulon genes. Names of strains with genome-wide admixture (Table S3) are boxed. Strain names are colored according to species designations. Complete genes with a phylogenetic position (Figure S28) as expected based on population genomic analysis (Figure S9) are labeled as white. Genes acquired from another lineage by gene flow are labelled orange. Genes with premature stop codons or in a more advanced state of pseudogenization are labelled gray. Genes with a complex ancestry, such as unexpectedly ancient alleles, are labelled cyan. Genes not detected by any of the methods employed in this study (see Online Material and Methods) were considered absent and are labelled red. B) Maximum biomass production (OD600) on 2 % galactose. Each point is a strain colored by species designation. Data was split based on whether (left) or not (right) gene flow had occurred. Asia A and B *S. kudriavzevii* (on the right) were separated from the rest of *Saccharomyces* data points for clarity. The groups are defined as follows: i) *S. cerevisiae*: Group1 (Domesticated strains: Bioethanol, Beer 1 & 2, Wine/European, Sake), Group 3 (West African), Group 4 (CHN IV), Group 5 (Asian Islands, Malaysian, North American). ii) *S. paradoxus*: Group 1 (European), Group 2 (Far East), Group 3 (America B), Group 4 (America C). iii) *S*. *mikatae*: Group 1 (Asia A), Group 2 (Asia B). iv) *S*. *kudriavzevii*: Group 1 (EU), Group 2 (Asia A), Group 3 (Asia B). v) *S*. *arboricola*: Group 1 (Asia A), Group 2 (Oceania). vi) *S*. *uvarum*: Group 1 (Holarctic), Group 2 (South America A), Group 3 (South America B), Group 4 (Australasia). vii) *S*. *eubayanus*: Group 1 (Holarctic), Group 2 (Patagonia B), Group 3 (Patagonia A).

The two new admixed strains of *S. kudriavzevii* provided an even more striking example of gene flow and selection. We previously inferred long-term balancing selection based on local selection regimes for the functional genes and inactivated pseudogenes of *S. kudriavzevii* ^58^, but the populations with inactive (Asia A and B) or active (EU) *GAL* networks were strongly differentiated by geography and population structure. Here we discovered two strains isolated from Southern China (Figure S1D, Table S2) that shared more than 87 % genome ancestry with EU strains (Figure S10H) and yet were unable to grow on galactose (Figure 6B). Phylogenetic analyses demonstrated that the loss of this trait was due to the acquisition of six *GAL* pseudogenes (at four loci: *GAL1/GAL10/GAL7*, *GAL4*, *GAL2*, and *GAL80*) from the *S. kudriavzevii* Asia A population after the diversification of EU and Asia A populations (Figure S28K). Since these two strains shared less than 12 % genome ancestry with the Asia A lineage, in the absence of selection against hybrid networks or against *GAL* activity in Asia, the odds are quite low (*p* = 0.12^4^ = 0.0002) that these closely related strains would have acquired pseudogenes by chance at all 4 *GAL* loci that are functional in the EU population. Notably, the only two *GAL* loci not transferred from the Asia A lineage by gene flow into the ancestors of these two strains were *GAL3* and *GAL80B* (Figure S28K, Figure S10H), two pseudogenes that were inactivated in the ancestor of all known strains of *S. kudriavzevii* ^58^.

The data also suggested that intricate selection dynamics may be occurring at the *GAL2* locus that are not simply qualitative. Most *S. eubayanus* and *S. uvarum* strains have a tandem duplication at the *GAL2* locus whose function is unknown ^17, 58–60^. Some *S. cerevisiae* strains from the CHNIII lineage that were isolated from milk fermentations also possess additional copies of *GAL2* whose origin predates the diversification of the genus; these strains lack functional copies of *HXT6* and *HXT7*, which encode hexose transporters, and seem to use *GAL2* to encode the transport of both galactose and glucose in dairy environments that are rich in lactose ^63^. Some *S. eubayanus* and *S. uvarum* strains have lost the *GAL2B* gene. Despite testing several growth conditions, including various galactose concentrations, the strains lacking *GAL2B* only displayed maximum growth rate differences at 30 °C on 2 % glucose, which was lower (Wilcoxon rank-sum test, *p*-value = 5.97 x 10^-4^, Figure S26F). This result suggests a similar model for the evolution of the *S. uvarum*/*S. eubayanus GAL2B* gene and the additional copies of *GAL2* in *S. cerevisiae*, wherein these additional copies of *GAL2* evolved to support glucose transport in specific ecological conditions. Notably, the single copies of *GAL2* from *S. eubayanus* Holarctic strains are an outgroup to the entire *S. uvarum/S. eubayanus* clade, including all known *GAL2* and *GAL2B* alleles (Figure S28B), suggesting that multiple ancient alleles are segregating at this locus due to balancing selection ^60^. Collectively, these results highlight how local selection regimes can maintain ancient polymorphisms, even in multi-locus gene networks.

## Discussion

### *Saccharomyces* diversification within and outside of Asia in association with plants

Several authors have postulated Asia as the geographical origin of *S. cerevisiae* and other species of *Saccharomyces* ^13, 22, 25, 28, 37, 64, 65^. Our present results provide evidence to support several rounds of speciation in Asia, as well as potentially the origin of the genus itself: i) the high genomic diversity in the Palearctic biogeographic realm, which includes Asia; ii) the centrality of Palearctic mitochondrial haplotypes to the mitochondrial network; iii) and ancestral polymorphisms in Asian strains that generate phylogenetic conflict and, in some cases, such as the *GAL* loci, phenotypic differences that are likely under strong selection. The presence of ancestral polymorphisms in several populations and species suggests that *Saccharomyces* diversification was rapid ^66^, that considerable gene flow continued prior to the generation of strong species barriers ^67–71^, or both. The presence of all species in association with trees of the order Fagales points to the adaptation of the last common ancestor of *Saccharomyces* to these hosts. However, there is still much to learn about the ecological distribution of yeasts in general, and *Saccharomyces* in particular ^72^, where sampling has often been biased toward bark and soil samples from Fagales. Even though most new lineages and species likely originated in Asia, our comprehensive global sampling and analyses strongly support the hypothesis that several lineages originated in South America, North America, Europe, and Oceania, including lineages of *S. eubayanus*, *S. paradoxus*, *S. uvarum*, *S. jurei*, and *S. arboricola* ^14, 21, 24, 26, 27, 31, 73–75^ (Figure 4D). These diversifications could be accompanied by the adaptation to new hosts. For example, *S. uvarum* and *S. eubayanus* lineages are frequently isolated from fungi associated with trees of the genus *Nothafagus* in South America. This influence of related *Nothafagus* hosts during diversification might help explain the similar phenotypic traits observed among *S. uvarum* and *S. eubayanus* strains.

The ecological and genetic factors driving this diversification of the genus could also be linked to temperature fluctuations during the Miocene epoch, which is coincident with *Saccharomyces* divergence times ^30^. Temperature fluctuations have played an important role in the diversification of plants ^76^ and animals ^77^, and temperature tolerance differentiate several *Saccharomyces* species and clades. In particular, the high temperature tolerance of *S. cerevisiae* and *S. paradoxus* ^51, 52, 78^ seems to be a derived trait. The influence of temperature during the diversification might be one of the reasons why we observe frequent introgressions in the mitochondrial genome ^44–47^, where species-specific mitotypes have been shown to strongly affect temperature tolerance ^50, 79^. Clear patterns of differentiation by geographic distribution and climatic conditions have also been detected for *Saccharomyces* mitotypes ^26, 33, 65, 80, 81^.

The role of introgressions during lineage diversification is still under debate, but nuclear introgressions between species have been mainly observed in human-associated environments, including the horizontal gene transfer of few genes ^82–84^, frequent admixture of domesticated *S. cerevisiae* strains ^13, 36, 62^, and interspecies hybridization of strains used to produce fermented beverages ^85–87^. In contrast, cytoplasmic genetic elements have undergone extensive introgression and gene flow even in wild strains of *Saccharomyces*, as previously seen in animals ^88–90^.

### *Saccharomyces* populations are often more genetically differentiated than multicellular eukaryotic species

Multicellular eukaryotes might be more permeable to interspecies introgression ^91, 92^ because animal and plant species are more closely related than species are in the genus *Saccharomyces*. The distinction is not entirely due to differences in taxonomic practice because, even when we considered phylogenetically distinct *Saccharomyces* lineages, only 9.25 % of *Saccharomyces* nuclear genomes were admixed. Spore viabilities lower than 1 % ^69, 93^ in crosses between strains have been considered sufficient to define yeast species using the biological species concept alone. When combined with phylogenetic and ecological species concepts, taxonomic authorities have accepted spore viabilities lower than 10 %, as seen for *S. eubayanus* and *S. uvarum*, which have the highest AAI values among currently recognized species ^94, 95^.

Our comparison of AAI values with multicellular eukaryotes suggests that species designations based on spore viability and other currently used criteria do not differentiate *Saccharomyces* species as finely as the criteria deployed by plant and animal taxonomists. If they did, what we currently consider *Saccharomyces* populations or lineages might be more analogous to the species designations of multicellular eukaryotes. Even so, current yeast taxonomic practice has the advantage of recognizing the ease with which genes of phenotypic importance flow between populations of the same species.

### Phenotypic diversity through complex ancestries

Phenotypic traits are gained and lost frequently in animals, plants, and fungi ^30, 96–98^. Alternatively, traits can be retained in a species by balancing selection when different lineages or populations maintain genes or even multi-locus gene networks encoding traits due to local adaptation or fluctuating conditions. For example, here we showed that some admixed *S. paradoxus* America C strains regained the ability to grow in melibiose by acquiring a functional *MEL1* gene from the *S. paradoxus* America B population. Even more strikingly, two admixed *S. kudriavzevii* strains, which were isolated in Asia but were more closely related to the EU population, lost the ability to grow in the presence of galactose by acquiring *GAL* pseudogenes from the Asia A population, directly demonstrating gene flow between Gal^+^ and Gal^-^ populations of *S. kudriavzevii* for the first time ^58^. Recent studies concluded that *S. cerevisiae* maintained alternative higher-activity versions of the *GAL* network due to segregating variation at multiple loci ^60^. Our new results here definitively show that qualitative variation can also segregate within a species for a multi-locus gene network, and indeed, suggest that pseudogenized genes may be preferred in some environments. We conclude that the maintenance of compatible alternative versions of gene networks, even at unlinked loci, may be more frequent than previously thought.

## Conclusion

The model genus *Saccharomyces* and the current dataset provide an important quantitative benchmark of the boundaries of lineages, populations, and species in terms of genetic variation, phenotypic variation, and the relationship between genotype and phenotype. Setting these boundaries helps characterize eukaryotic microbial biodiversity, understand ecological dynamics, and offers bioresources of industrial interest.

## Supporting information

SuppFiguresTextANDOnlineMethods

## Acknowledgments

We thank the University of Wisconsin Biotechnology Center DNA Sequencing Facility for providing Illumina and Sanger sequencing facilities and services; Maria Sardi, Audrey Gasch, and Ursula Bond for providing strains; Sean McIlwain for providing guidance for genome ultra-scaffolding; Yury V. Bukhman for discussing applications of the Growth Curve Analysis Tool (GCAT); Mick McGee for HPLC analysis; Raúl Ortíz-Merino for assistance during `YGAP` annotations; Jessica Leigh for assistance with `PopART`; Cecile Ané for suggestions about `BUCKy` utilization and phylogenetic network analyses; Samina Naseeb and Daniela Delneri for sharing preliminary multi-locus *Saccharomyces jurei* data; and Branden Timm, Brian Kyle, and Dan Metzger for computational assistance. Some computations were performed on Tirant III of the Spanish Supercomputing Network (‘‘Servei d’Informàtica de la Universitat de València”) under the project BCV-2021-1-0001 granted to DP, while others were performed at the Wisconsin Energy Institute and the Center for High-Throughput Computing of the University of Wisconsin-Madison. During a portion of this project, DP was a researcher funded by the European Union’s Horizon 2020 research and innovation programme Marie Sklodowska-Curie, grant agreement No. 747775, the Research Council of Norway (RCN) grant Nos. RCN 324253 and 274337, and the Generalitat Valenciana plan GenT grant No. CIDEGENT/2021/039. DP is a recipient of an Illumina Grant for Illumina Sequencing *Saccharomyces* strains in this study. QKL was supported by the National Science Foundation under Grant No. DGE-1256259 (Graduate Research Fellowship) and the Predoctoral Training Program in Genetics, funded by the National Institutes of Health (5T32GM007133). This material is based upon work supported in part by the Great Lakes Bioenergy Research Center, Office of Science, Office of Biological and Environmental Research under Award Numbers DE-SC0018409 and DE-FC02-07ER64494; the National Science Foundation under Grant Nos. DEB-1253634 and DEB-2110403; and the USDA National Institute of Food and Agriculture Hatch Project Number 1020204. C.T.H. is a H. I. Romnes Faculty Fellow, supported by the Office of the Vice Chancellor for Research and Graduate Education with funding from Wisconsin Alumni Research Foundation. QMW was supported by the National Natural Science Foundation of China (NSFC) under Grant Nos. 31770018 and 31961133020. CRL holds the Canada Research Chair in Cellular Systems and Synthetic Biology, and his research on wild yeast is supported by a NSERC Discovery Grant.

## Author contributions

DP performed most analyses (phenotyping, computational analyses, and figure plots) and data management; DP, CG, and QMW provided *COX2* and *COX3* sequences by PCR and Sanger sequencing; DP and JK designed the alignment pipeline; JK uploaded genomes to the gxseq.glbrc.org genome browser server; EJU and RLW confirmed *GAL* genes by PCR and Sanger sequencing; MCK performed growth rate correlation analyses and plots for different sugar concentrations; QKL, ABH, and DAO prepared paired-end Illumina libraries; DP, MA, and JAK prepared mate-pair Illumina libraries; DP and QKL designed the population genomic pipeline; QMW, FYB, JBL, CRL, JPS, PG, DL, DH, KH, and JCF contributed key strains to study design; DP and CTH conceived of and designed the study; DP and CTH wrote the manuscript with editorial input from JK, MCK, QKL, JCF, CRL, JBL, FYB, KH, PG, and JPS; and all co-authors approved the final version of the manuscript.

## Author information

### Data deposition statement

Code availability: https://perisd.github.io/Sac2.0/ website provides access to custom scripts and information regarding raw data. Raw data is deposited in FigShare (https://figshare.com/s/93614f0e128d86f2ed8e). Data availability: Strains physically used in this study (i.e. with codes FM[Number] (e.g. FM1198) or yHXX[Number] (e.g. yHAB33) are available from and have been submitted to the Portuguese Yeast Culture Collection (PYCC) (Table S1). *COX2* and *COX3* sequences were deposited in GenBank under accession nos. MH813536-MH813939. *GAL* genes that were Sanger-sequenced were deposited in GenBank under accession nos. OL660614-OL660618. Illumina sequencing data have been deposited in NCBI’s SRA database, Bioproject PRJNA475869. Genome assemblies and annotations are available at gxseq.glbrc.org and on European Nucleotide Archive (ENA) project accession number PRJEB48264.

## Online Material & Methods

### Extended tables

**Table S1.** List of strains used in this study.

**Table S2.** Sequencing and genome assembly statistics.

**Table S3.** Genome contributions in admixed and introgressed strains.

**Table S4.** *Saccharomyces* 2-µm plasmid information.

**Table S5.** Reciprocal monophyly tests.

**Table S6.** Kinetic growth parameter information for *Saccharomyces* strains.

**Table S7.** PCR primers and conditions.

### Supplementary Notes & Figures

**Supplementary Note 1** – Nuclear and mitochondrial genome diversity.

**Supplementary Note 2** – *Saccharomyces* phenotypic diversity.

**Figure S1.** Geographic locations of *Saccharomyces* populations.

**Figure S2.** Association biases for hosts and substrates of *Saccharomyces* strains.

**Figure S3.** COX3 phylogenetic network of *Saccharomyces* strains.

**Figure S4.** Phylogenetic networks of mitochondrial genes.

**Figure S5.** Genome dot plots of *Saccharomyces* strains compared to the *S. cerevisiae* S288C laboratory strain.

**Figure S6.** Highly diverse genomic architectures among *Saccharomyces* species.

**Figure S7.** Mitochondrial genome dot plots of *Saccharomyces* strains compared to the *S. cerevisiae* S288C laboratory strain.

**Figure S8.** Mitochondrial genome dot plots of *Saccharomyces* populations compared to other populations.

**Figure S9.** Population genomics of seven *Saccharomyces* species.

**Figure S10.** Genome-wide pairwise nucleotide sequence divergence plots for admixture *Saccharomyces* strains.

**Figure S11.** Genetic distance distributions.

**Figure S12.** Fst distributions.

**Figure S13.** `BUCKy` concordance primary tree and alternative topologies.

**Figure S14.** Phylogenomic network of *Saccharomyces* single-copy orthologous genes.

**Figure S15.** Levels of heterozygosity among *Saccharomyces* strains.

**Figure S16.** Introgressions between *S. cerevisiae* and *S. paradoxus*.

**Figure S17.** *Saccharomyces* 2 µm plasmid inheritance.

**Figure S18.** Percentage of *Saccharomyces* that grew above OD600=0.5 in various growth conditions.

**Figure S19.** Growth conditions promoting flocculation among *Saccharomyces* strains.

**Figure S20.** Growth variation in simple sugars across concentrations and the impact of Gal4-binding sites.

**Figure S21.** Lag time and maximum growth rate correlations between low and high sugar concentrations.

**Figure S22.** Lag time correlations between monosaccharides and their disaccharides or trisaccharides.

**Figure S23.** Phenotypic variance across *Saccharomyces* species.

**Figure S24.** Principal component analysis of maximum OD600.

**Figure S25.** Variance contributed to each component by growth condition.

**Figure S26.** Kinetic parameters of *Saccharomyces* strains in different growth conditions.

**Figure S27.** Melibiose phenotypic diversity generated through complex genomic ancestries.

**Figure S28.** Individual phylogenetics trees of the *GAL*/*MEL* pathway.

**Figure S29.** Maximum OD600 violin boxplots of *Saccharomyces* populations/groups.

**Figure S30.** Summary statistics of *Saccharomyces* genome assemblies.

