## Supplementary material for "Macroevolutionary diversity of traits and genomes in the model yeast genus *Saccharomyces*": SuppFiguresTextANDOnlineMethods: FigureS10.pdf

A GS5.6

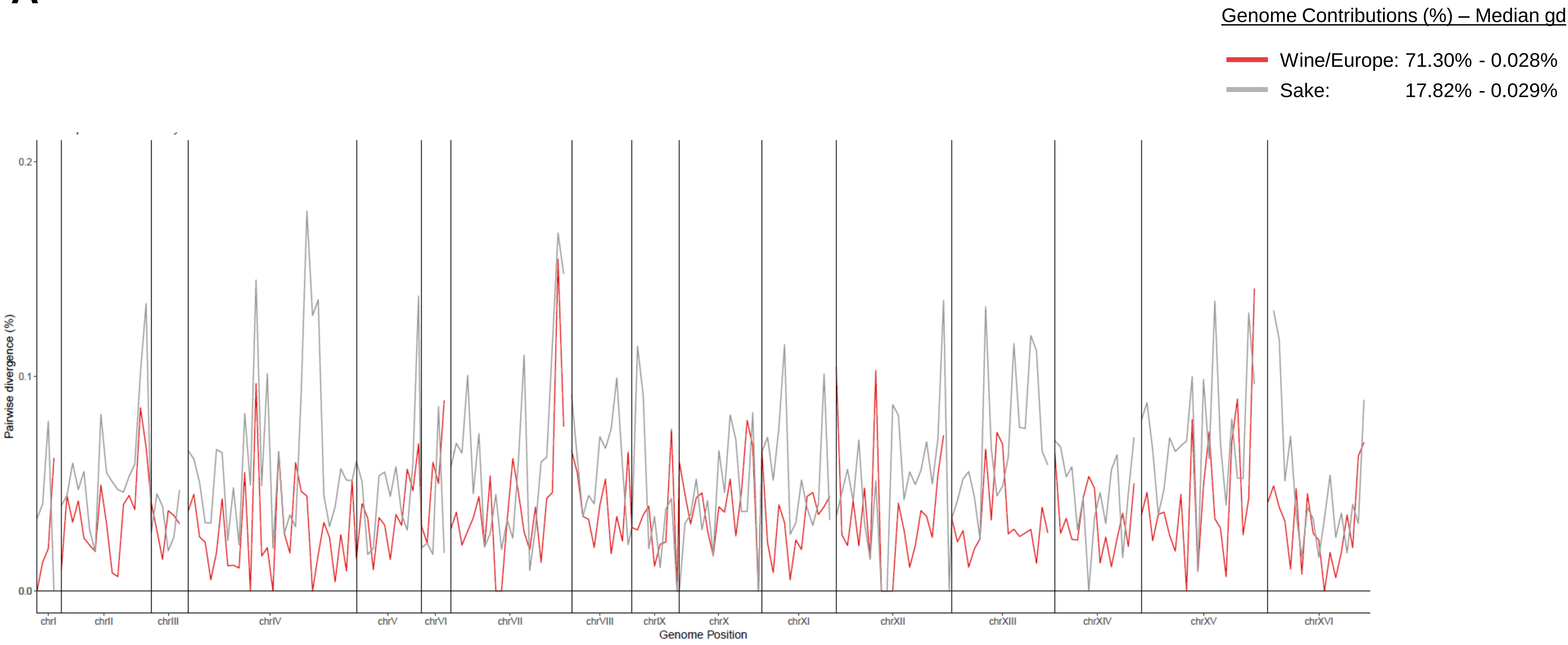

Genome Contributions (%) – Median gd
Wine/Europe: 71.30% - 0.028%
Sake: 17.82% - 0.029%
Pairwise divergence (%)
0.2
0.1
0.0
chrI
chrII
chrIII
chrIV
chrV
chrVI
chrVII
chrVIII
chrIX
chrX
chrXI
chrXII
chrXIII
chrXIV
chrXV
chrXVI
Genome Position

**B** SK1

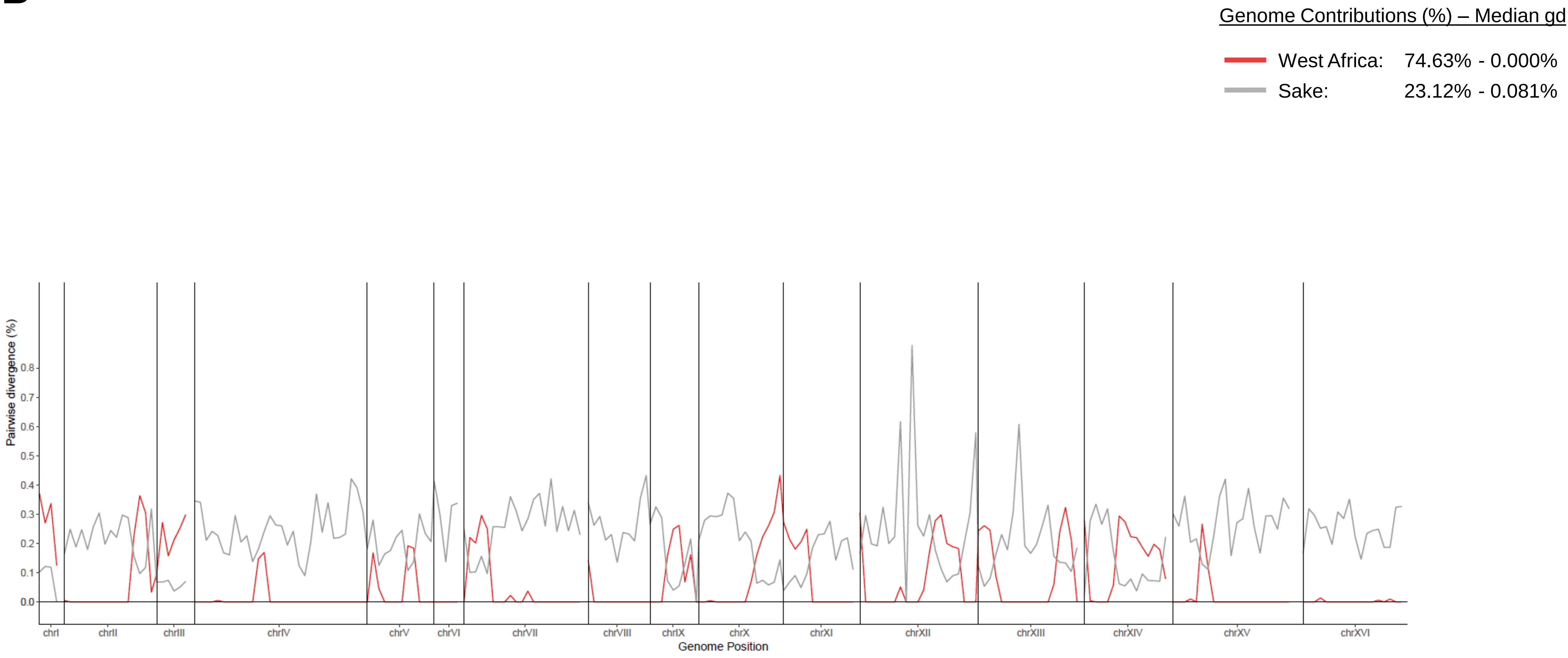
Genome Contributions (%) – Median gd
West Africa: 74.63% - 0.000%
Sake: 23.12% - 0.081%
Pairwise divergence (%)
0.0
0.1
0.2
0.3
0.4
0.5
0.6
0.7
0.8
chrI
chrII
chrIII
chrIV
chrV
chrVI
chrVII
chrVIII
chrIX
chrX
chrXI
chrXII
chrXIII
chrXIV
chrXV
chrXVI
Genome Position

**C** Y55

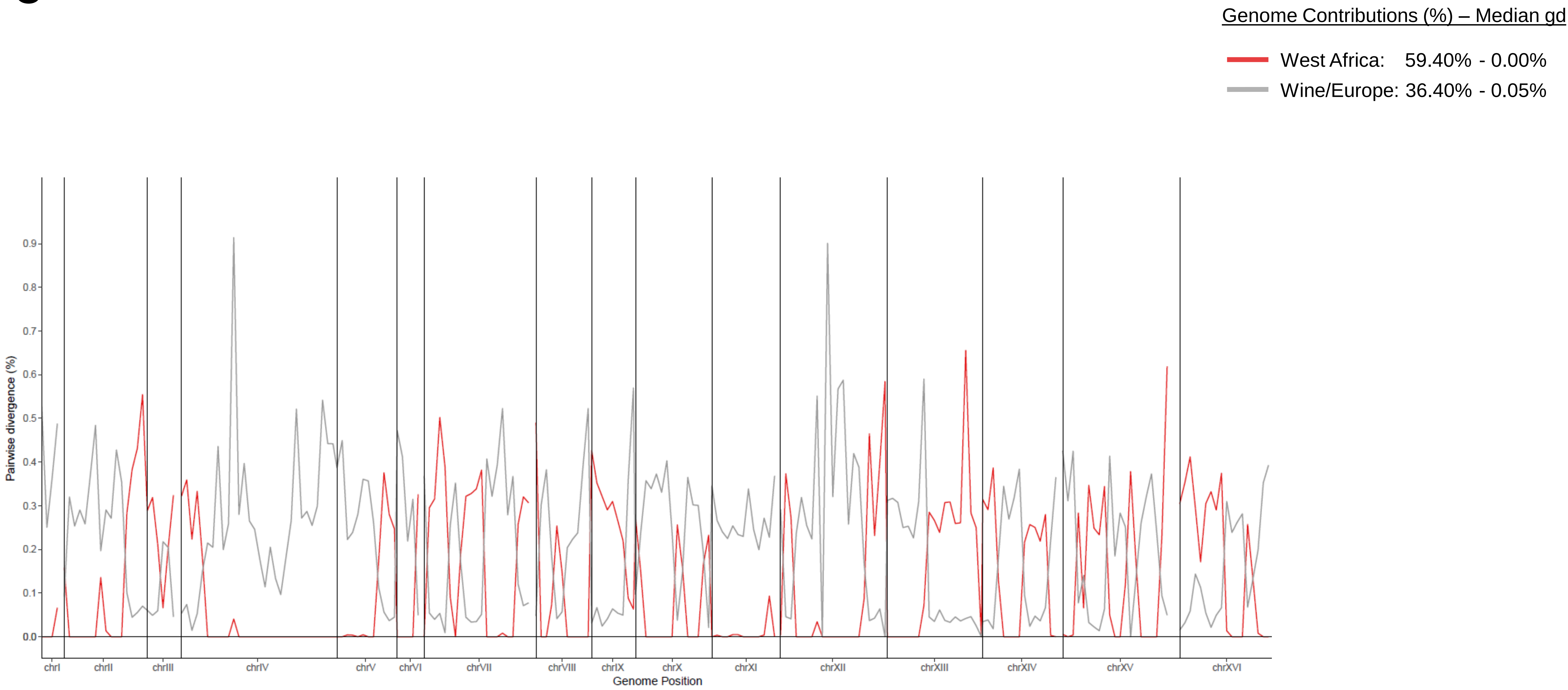
Genome Contributions (%) – Median gd
West Africa: 59.40% - 0.00%
Wine/Europe: 36.40% - 0.05%
Pairwise divergence (%)
0.0
0.1
0.2
0.3
0.4
0.5
0.6
0.7
0.8
0.9
chrI
chrII
chrIII
chrIV
chrV
chrVI
chrVII
chrVIII
chrIX
chrX
chrXI
chrXII
chrXIII
chrXIV
chrXV
chrXVI
Genome Position

D

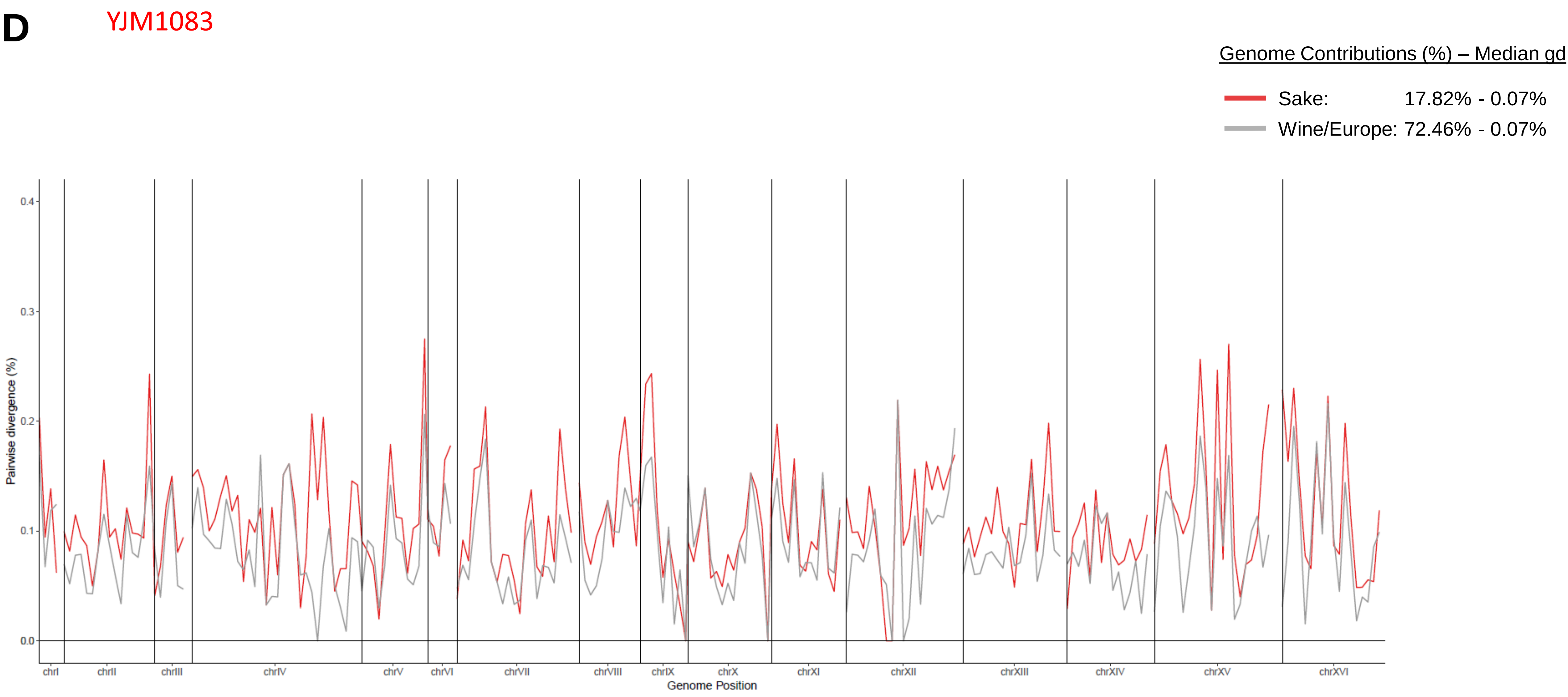
YJM1083
Genome Contributions (%) – Median gd
Sake: 17.82% - 0.07%
Wine/Europe: 72.46% - 0.07%
Pairwise divergence (%)
0.0
0.1
0.2
0.3
0.4
chrI
chrII
chrIII
chrIV
chrV
chrVI
chrVII
chrVIII
chrIX
chrX
chrXI
chrXII
chrXIII
chrXIV
chrXV
chrXVI
Genome Position

E

WLP380

Genome Contributions (%) – Median gd

Wine/Europe: 11.43% - 0.02%

Beer 1: 71.86% - 0.01%

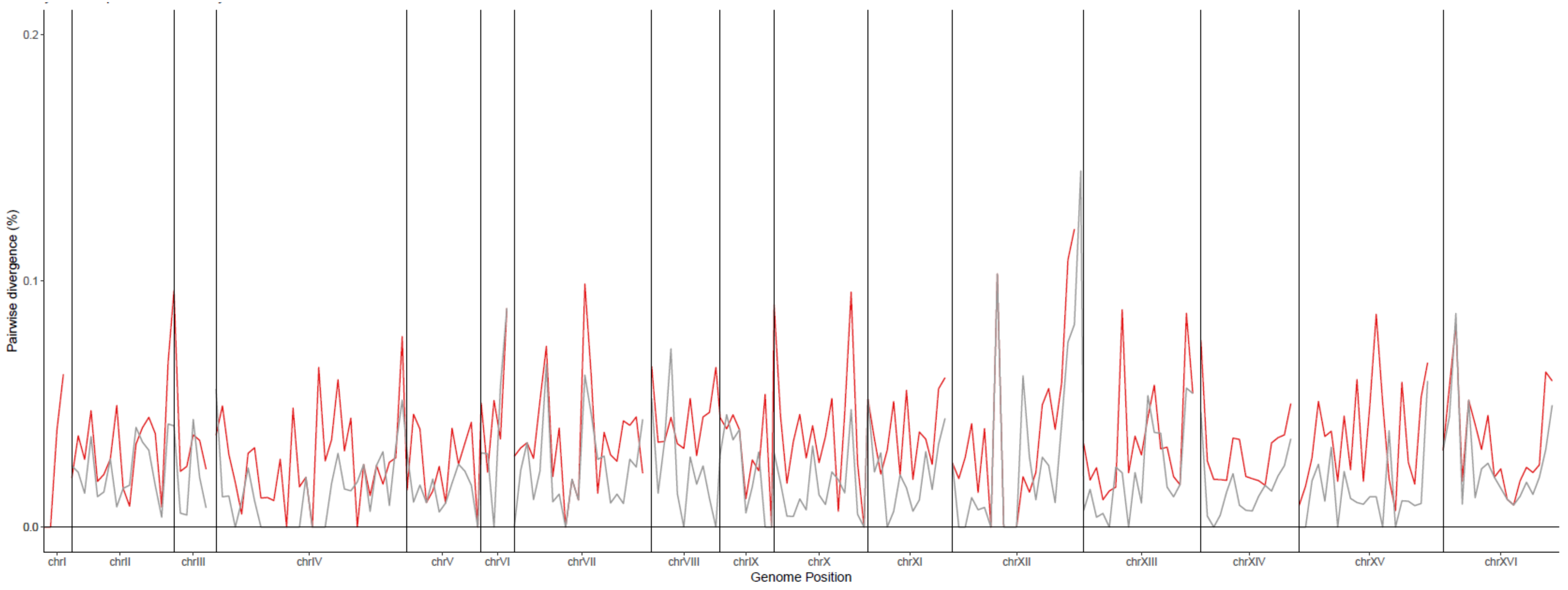

F

Wy1318

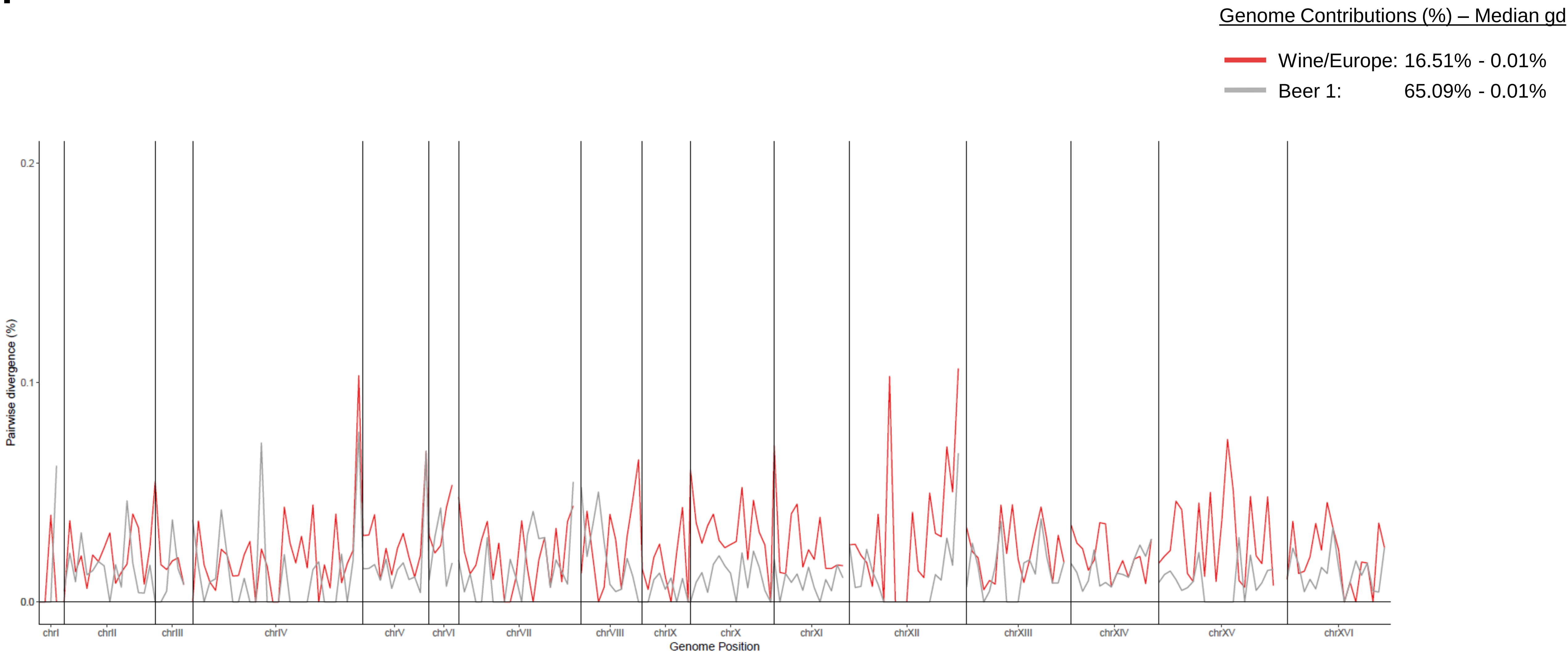
Genome Contributions (%) – Median gd
Wine/Europe: 16.51% - 0.01%
Beer 1: 65.09% - 0.01%
Pairwise divergence (%)
0.2
0.1
0.0
chrI
chrII
chrIII
chrIV
chrV
chrVI
chrVII
chrVIII
chrIX
chrX
chrXI
chrXII
chrXIII
chrXIV
chrXV
chrXVI
Genome Position

**G** LL2012-016/LL2012-018

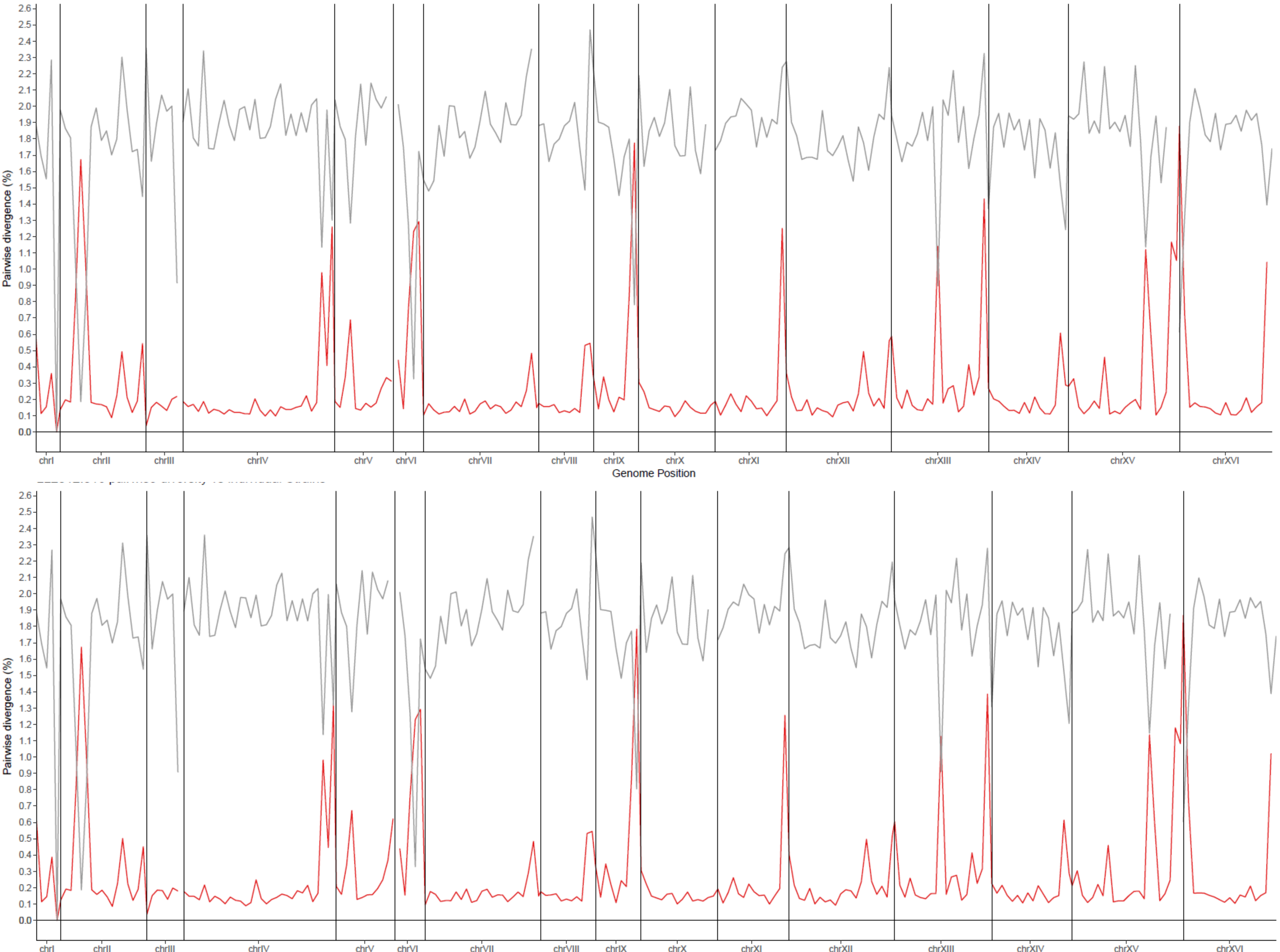


Genome Contributions (%) – Median gd

America C: 97.17% - 0.16%

America B: 2.46% - 0.78%

### H i) yHAB11/yHAB333

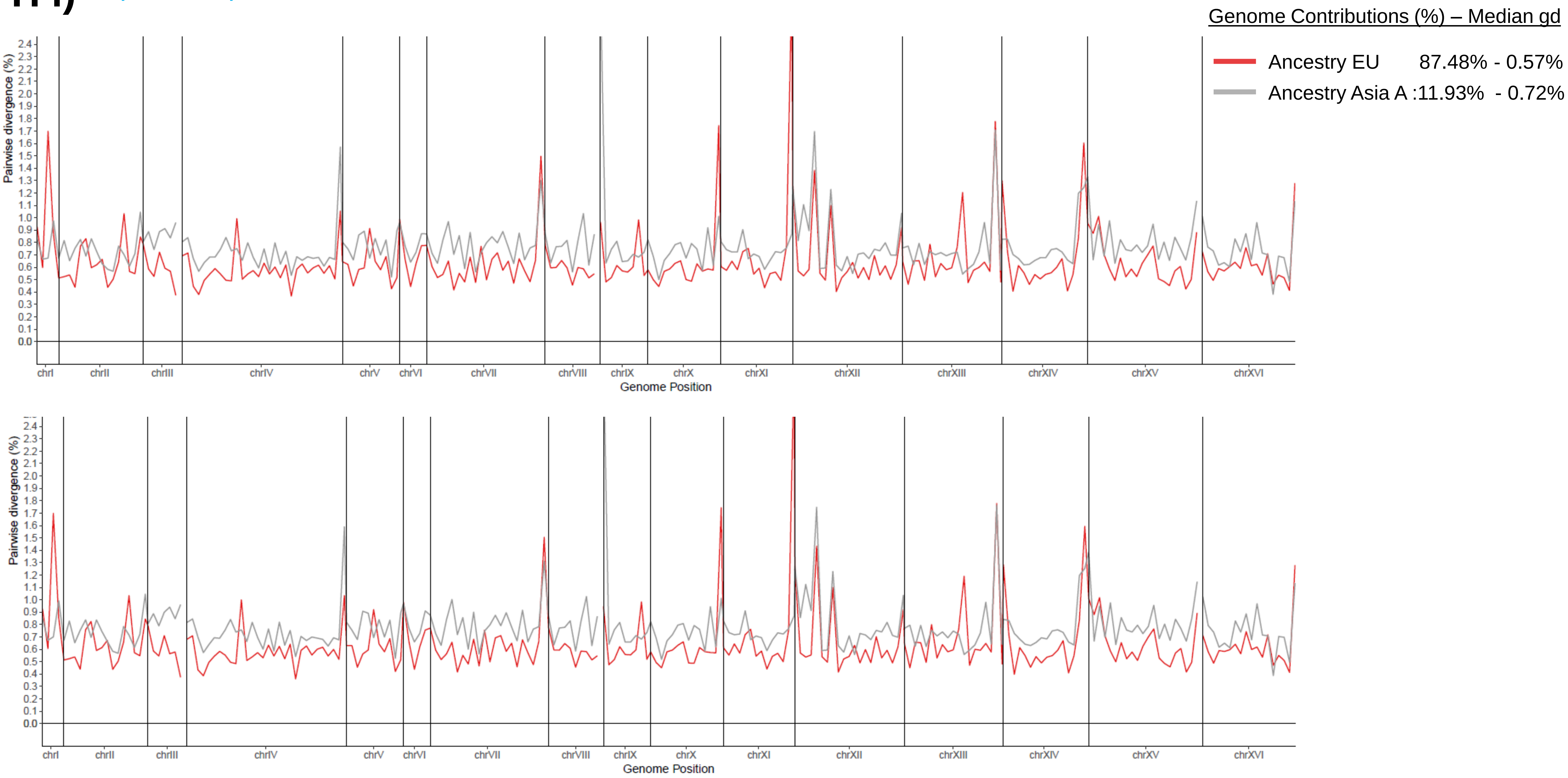

H ii) yHAB11/yHAB333

5 Kbp windows

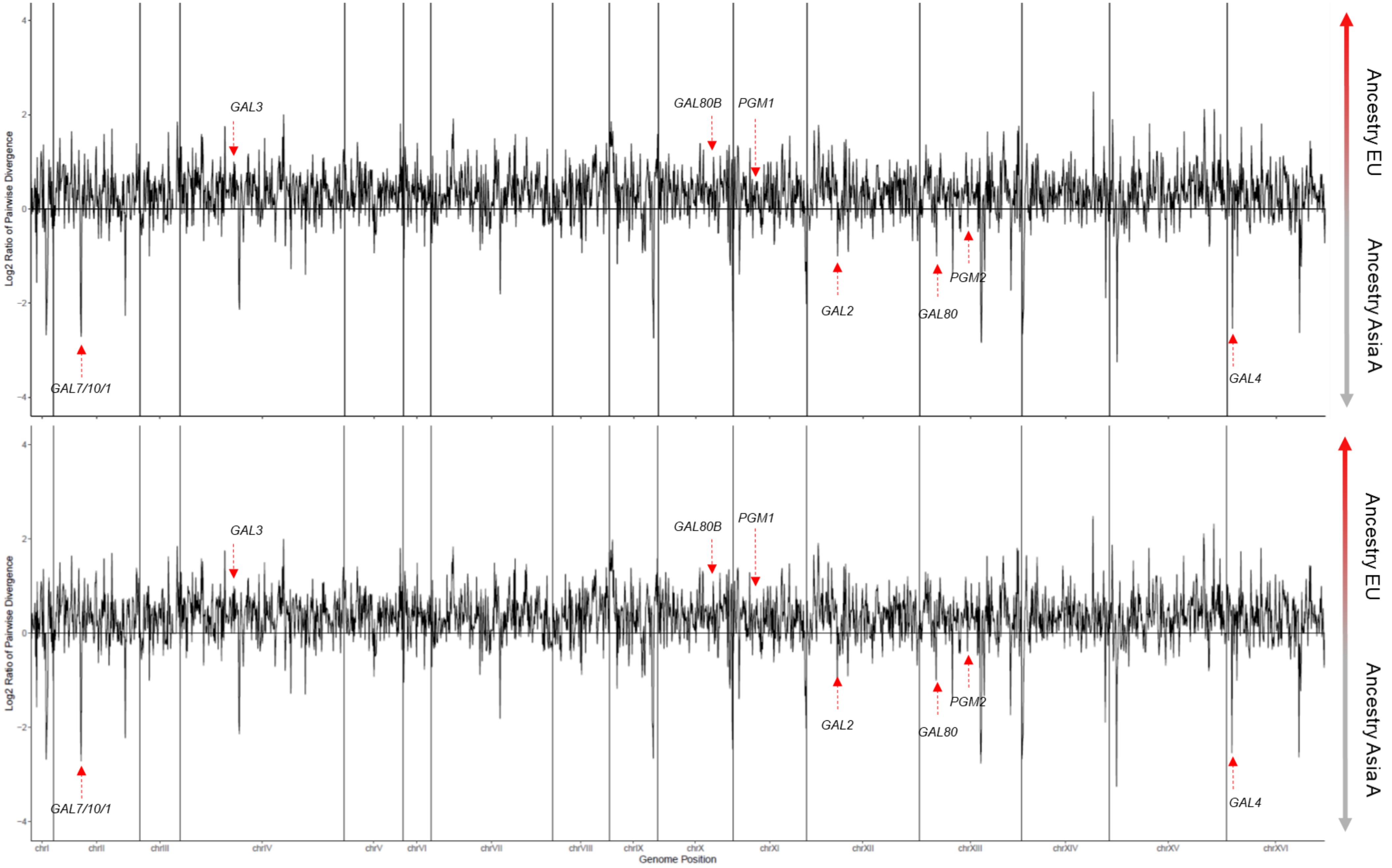

I

yHAB123

Genome Contributions (%) – Median gd

South America A: 39.53% - 0.09%

South America B: 58.74% - 0.25%

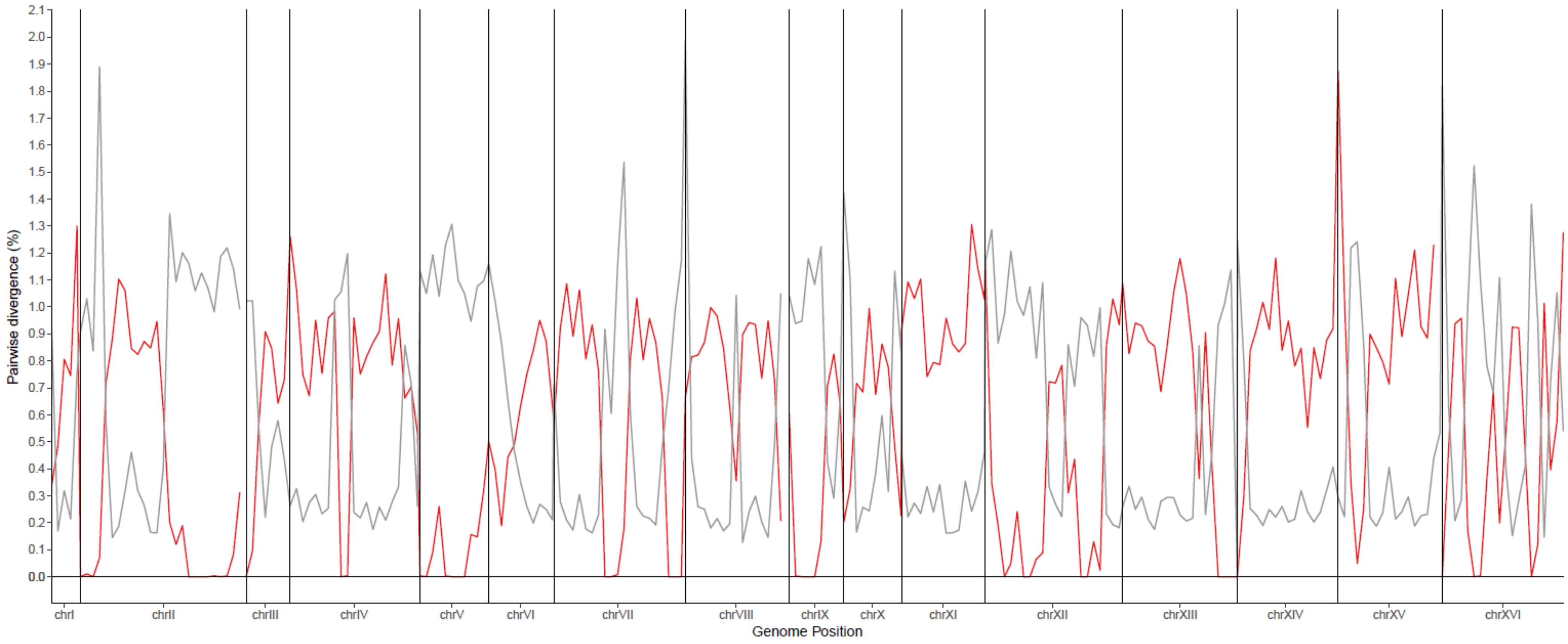

J

yHAB482

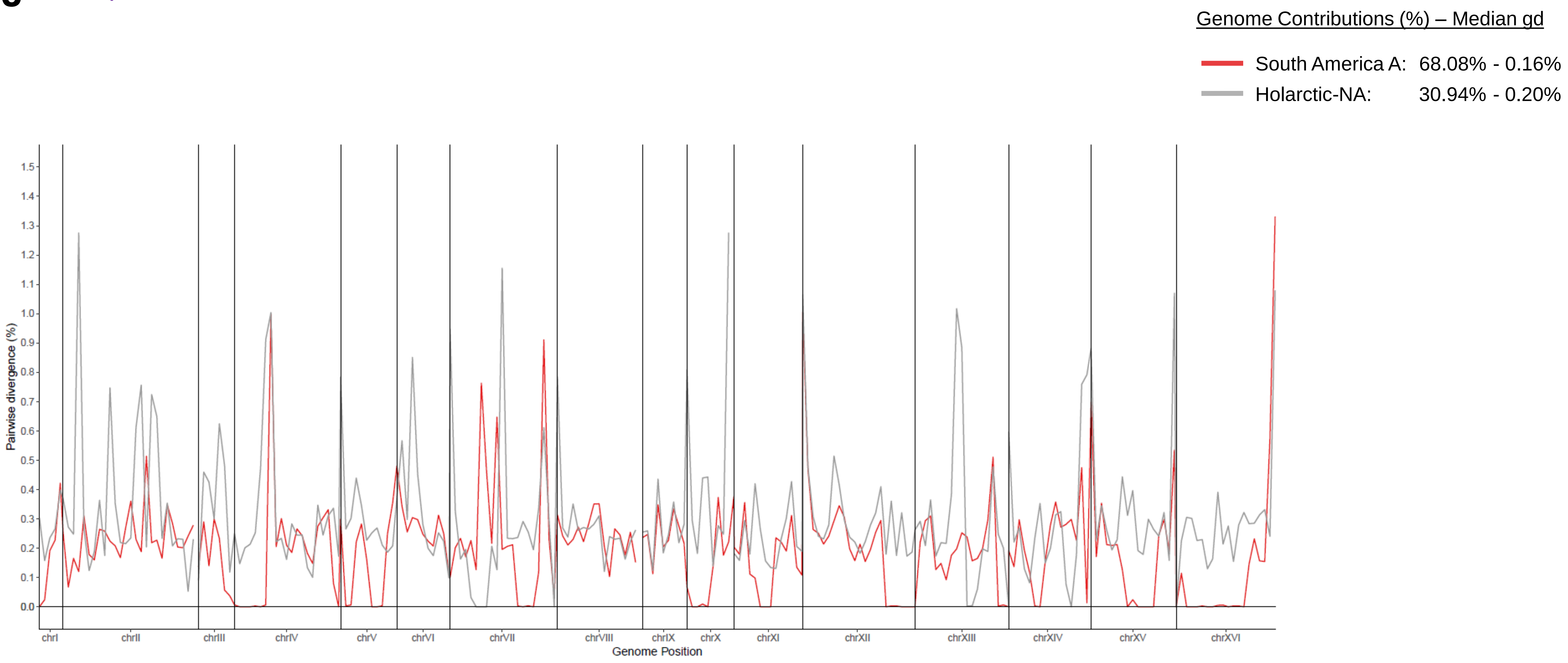
Genome Contributions (%) – Median gd
South America A: 68.08% - 0.16%
Holarctic-NA: 30.94% - 0.20%
Pairwise divergence (%)
0.0
0.1
0.2
0.3
0.4
0.5
0.6
0.7
0.8
0.9
1.0
1.1
1.2
1.3
1.4
1.5
chrI
chrII
chrIII
chrIV
chrV
chrVI
chrVII
chrVIII
chrIX
chrX
chrXI
chrXII
chrXIII
chrXIV
chrXV
chrXVI
Genome Position

K

FM1318

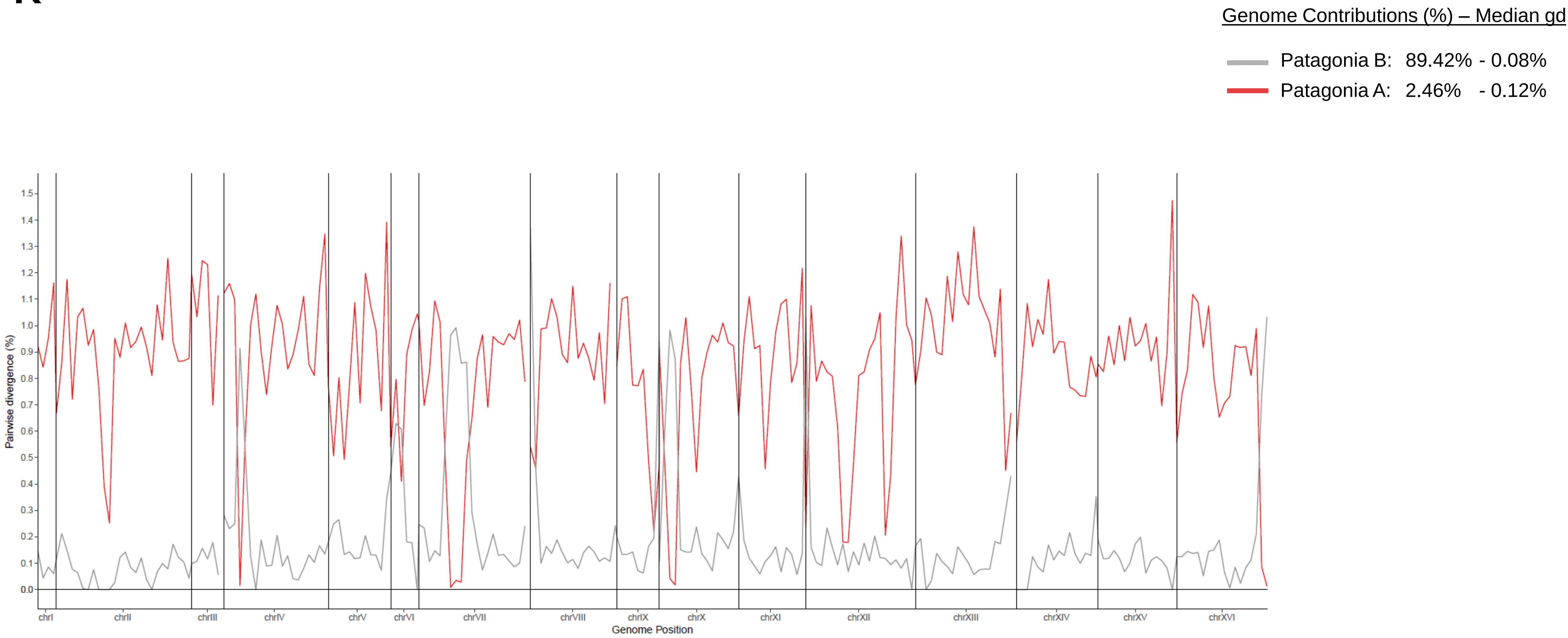
Genome Contributions (%) – Median gd
Patagonia B: 89.42% - 0.08%
Patagonia A: 2.46% - 0.12%
Pairwise divergence (%)
1.5
1.4
1.3
1.2
1.1
1.0
0.9
0.8
0.7
0.6
0.5
0.4
0.3
0.2
0.1
0.0
chrI
chrII
chrIII
chrIV
chrV
chrVI
chrVII
chrVIII
chrIX
chrX
chrXI
chrXII
chrXIII
chrXIV
chrXV
chrXVI
Genome Position

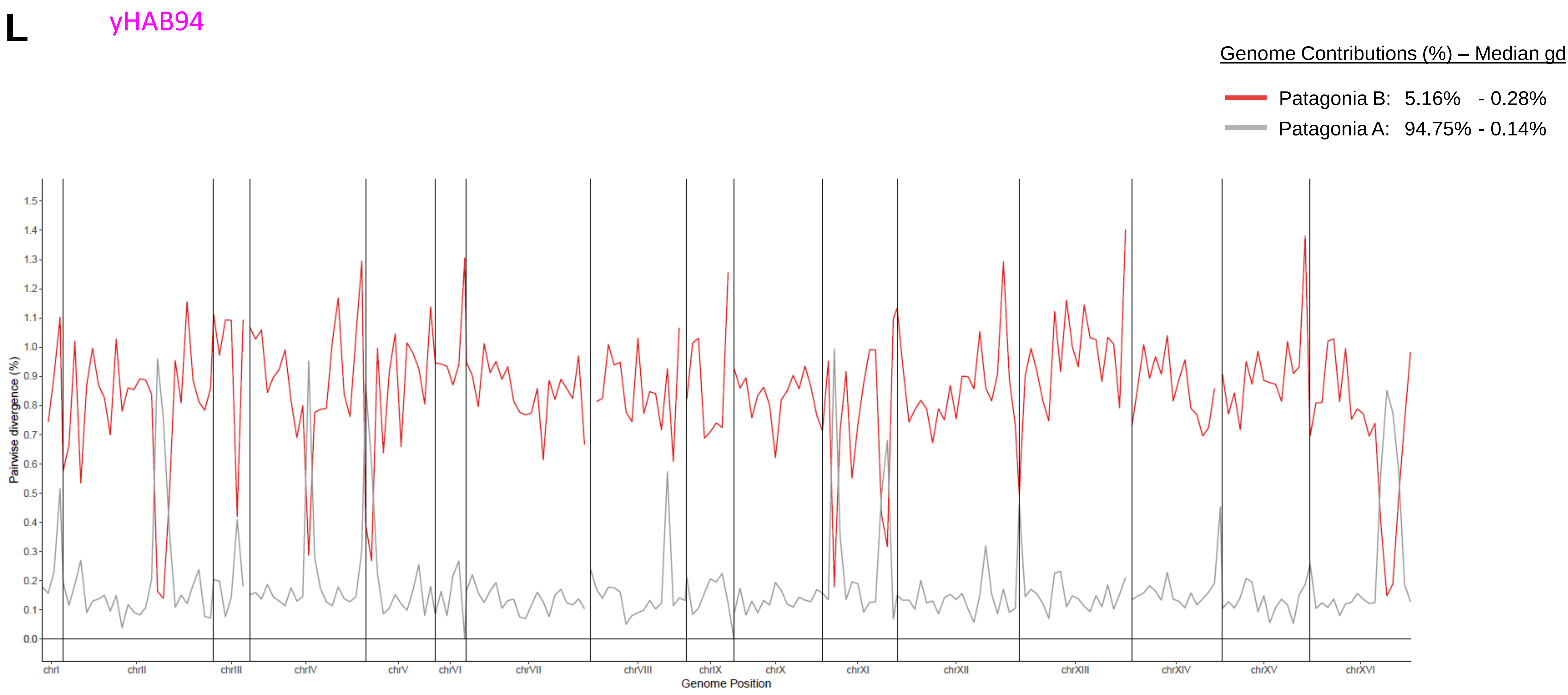
L
yHAB94
Genome Contributions (%) – Median gd
Patagonia B: 5.16% - 0.28%
Patagonia A: 94.75% - 0.14%
Pairwise divergence (%)
Genome Position
chrI
chrII
chrIII
chrIV
chrV
chrVI
chrVII
chrVIII
chrIX
chrX
chrXI
chrXII
chrXIII
chrXIV
chrXV
chrXVI

**M** yHCT101

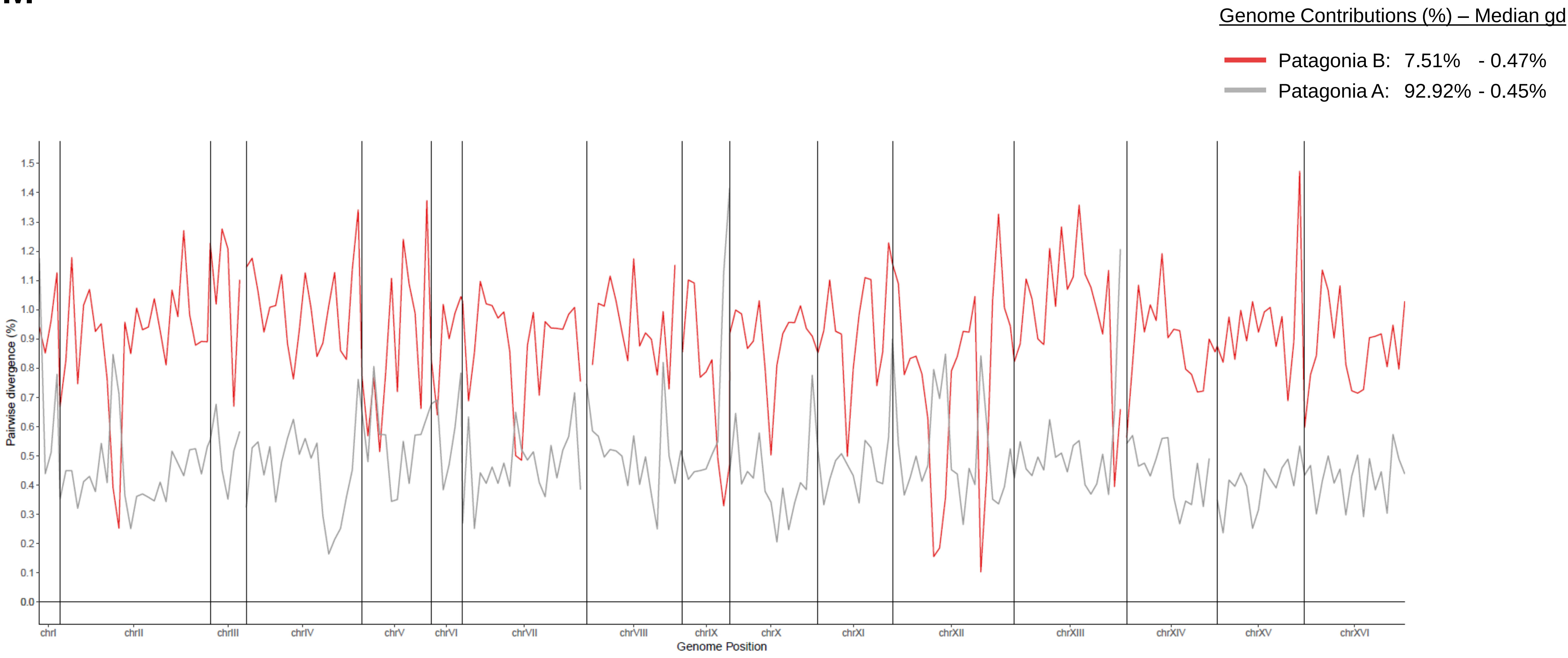

Genome Contributions (%) – Median gd
Patagonia B: 7.51% - 0.47%
Patagonia A: 92.92% - 0.45%
Pairwise divergence (%)
0.0
0.1
0.2
0.3
0.4
0.5
0.6
0.7
0.8
0.9
1.0
1.1
1.2
1.3
1.4
1.5
chrI
chrII
chrIII
chrIV
chrV
chrVI
chrVII
chrVIII
chrIX
chrX
chrXI
chrXII
chrXIII
chrXIV
chrXV
chrXVI
Genome Position

N

yHCT104

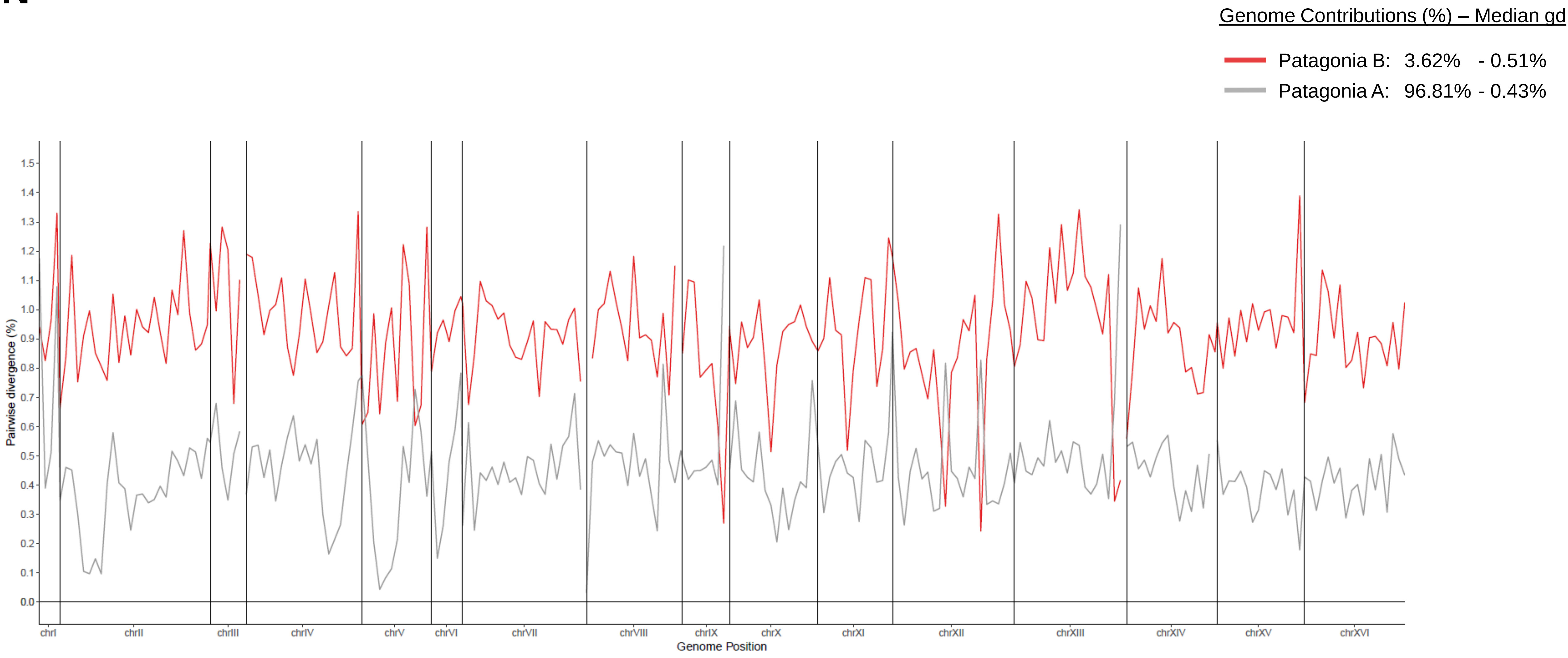
Genome Contributions (%) – Median gd
Patagonia B: 3.62% - 0.51%
Patagonia A: 96.81% - 0.43%
Pairwise divergence (%)
1.5
1.4
1.3
1.2
1.1
1.0
0.9
0.8
0.7
0.6
0.5
0.4
0.3
0.2
0.1
0.0
chrI
chrII
chrIII
chrIV
chrV
chrVI
chrVII
chrVIII
chrIX
chrX
chrXI
chrXII
chrXIII
chrXIV
chrXV
chrXVI
Genome Position

**O** yHCT96

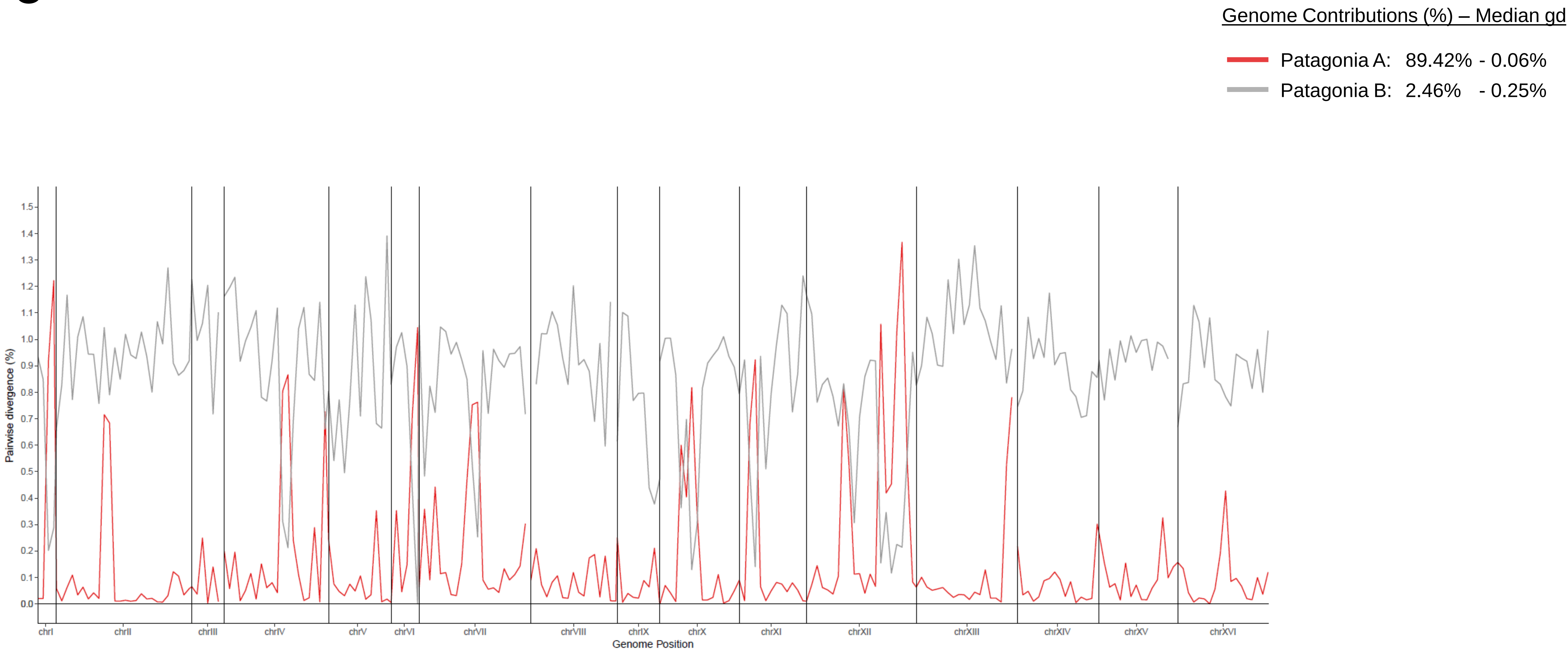
Genome Contributions (%) – Median gd
Patagonia A: 89.42% - 0.06%
Patagonia B: 2.46% - 0.25%
Pairwise divergence (%)
1.5
1.4
1.3
1.2
1.1
1.0
0.9
0.8
0.7
0.6
0.5
0.4
0.3
0.2
0.1
0.0
chrI
chrII
chrIII
chrIV
chrV
chrVI
chrVII
chrVIII
chrIX
chrX
chrXI
chrXII
chrXIII
chrXIV
chrXV
chrXVI
Genome Position
