## Supplementary material for "Macroevolutionary diversity of traits and genomes in the model yeast genus *Saccharomyces*": SuppFiguresTextANDOnlineMethods: Peris_etal_OnlineMethods_SSS2.pdf

### Methods

Yeast strains and maintenance

Strains submitted to the Portuguese Yeast Culture Collection (PYCC) are indicated in Table S1. Strains with codes FM[Number] (i.e. FM1198) or yHXX[Number] (i.e. yHAB33) are physically present and may be requested from the Hittinger Lab (Table S1). For the rest of the strains, references are provided in Table S1 to request them from the corresponding lab. Yeast strains are stored in cryotubes with YPD (1 % yeast extract, 2 % peptone, and 2 % glucose) and 15 % glycerol at -80 °C. Routine cultures were maintained in YPD plus 2 % agar plates at 24 °C. The taxonomic order of hosts was retrieved for each species based on the "Host" column of Table S1 using the `R` package `taxize v0.9.99` [1].

*COX2* and *COX3* PCR amplification, sequencing, and analyses

Partial gene sequences were obtained for *COX2* (471 bp) and *COX3* (491 bp) mitochondrial genes using primers and conditions described by Peris et al. [2]. *COX2* is highly polymorphic marker, which useful to trace ancient hybridization events [2], due to genetic footprints left by a free-standing homing endonuclease inserted into its coding sequence [3,4]. In contrast, *COX3* is less affected by homing activity than *COX2*, providing a better picture of mitochondrial inheritance [2]. Genomic DNA (gDNA) was isolated following the phenol:chloroform procedure [5]. Gene sequences were determined by PCR and Sanger sequencing. Sequences were edited and assembled with `STADEN Package` version 1.7 [6]. *COX2* and *COX3* sequences were deposited in GenBank under accession nos. MH813536-MH813939.

*COX2* and *COX3* sequences of *Saccharomyces* strains whose whole genomes were sequenced (Table S2) were retrieved from genome assemblies using a local BLAST v2.6 [7] or from raw Illumina reads by using HybPiper v1.2 [8]. New sequences were manually added to previously aligned gene sequences [4]. 1776 *Saccharomyces COX2* sequences, and 996 *COX3* sequences in FASTA format were classified by haplotype using DnaSP v5 [9] and converted to nexus format. The nexus file, with haplotype and biogeographic realm frequency information (Table S1), was used as input for PopART v1.7 (http://popart.otago.ac.nz) to reconstruct a phylogenetic network. The relationship among haplotypes was inferred by using the Templeton, Crandall, and Sing (TCS) method [10] (Figure 2A and Figure S3).

### Paired-end and Mate-pair Illumina library preparation

Representative strains of diverse *Saccharomyces* lineages were selected to prepare 2x300 bp or 2x250 bp paired-end and 2x100 bp mate-pair Illumina libraries (Table S2). To explore intrapopulation diversity across the genus (Figure 3A,B), additional Illumina paired-end libraries for 65 *Saccharomyces* strains from different species were made, and sequencing lengths were between 100 bp to 300 bp.

#### *Paired-end Illumina libraries*

Paired-end Illumina libraries were prepared as previously described [5]. Libraries were sequenced using Illumina HiSeq 2000, HiSeq 2500 Rapid, or MiSeq. The quality and quantity of the finished libraries were assessed using an Agilent DNA1000 series chip assay (Agilent Technologies) and Invitrogen Qubit HS Kit (Invitrogen, Carlsbad, CA), respectively, and the libraries were standardized to 2 nM. Images were analyzed using CASAVA version 1.8.2.

*Mate-pair Illumina libraries*

Genomic DNA was isolated following the phenol:chloroform procedure [5]. gDNA was quantified with a Qubit HS Kit (Invitrogen, Carlsbad, CA), and 4 µg were used to perform the Gel-plus protocol of the Nextera Mate Pair Library Prep Kit (Illumina, San Diego, CA). The target size selection was 8 kbp (Ty elements are around 6 kbp [11]), and fragments were isolated from the gel with a QIAEX II Gel Extraction Kit (QUIAGEN, Germantown, MD). Fragments were sonicated in a Covaris instrument using Covaris tubes (Covaris, Woburn, Massachusetts), and 300-400 bp fragments were targeted. gDNA cleanups were performed using Axygen Mag PCR Cleanup beads (Axygen, Union City, CA). The quality and quantity of the sonicated fragments and finished libraries were assessed using an Agilent DNA1000 series chip assay (Agilent Technologies, Santa Clara, CA) and Invitrogen Qubit HS Kit (Invitrogen, Carlsbad, CA), respectively. The libraries were standardized to 2 nM. Sequencing images were analyzed using CASAVA version 1.8.2.

Quality filtering, genome assembly, and annotations

Reads were demultiplexed, and Illumina adapters were removed using `Trimmomatic v0.33` [12] with parameters `2:30:10 TRAILING:3 MINLEN:`[`20` for reads shorter than 101 bp or `25` for larger reads] and `NextClip v1.3.1` [13], respectively. A quick phylogenetic assessment was performed with an `Alignment and Assembly Free (AAF) method v20150930` [14] to check that the correct strain was sequenced. Briefly, AAF reconstructed a phylogenetic tree using Illumina reads as an input and the correct phylogenetic position of the sequenced strains was assessed.

Trimmed reads were assembled using the meta-assembler pipeline `iWGS v1.1` [15]. Briefly, the wrapper performs quality-based read trimming, follow by *k*-mer length optimization, and uses multiple state-of-the-art assemblers to generate genome sequence assemblies. The quality of the assemblies was assessed using `QUAST v3.2` [16] as implemented in the wrapper, and the best assembly was chosen based on the number of contigs/scaffolds and N50 statistic (Table S2). For the collection of 22 representative *Saccharomyces* strains and the *Saccharomyces eubayanus* CDFM21L.1, additional steps were taken to decrease the number of scaffolds and generate a nearly complete genome assembly. Scaffolds longer than 10 kbp were retained. Ultrascaffolding was manually done in `Geneious vR6` [17] by using synteny information generated by `MUMmer v3.23` with parameters `--maxgap=500 --mincluster=150`. This process was used to order scaffolds by comparing them to previously assembled high-quality genomes and to the best newly assembled genomes. Concatenated scaffolds were separated by manual addition of 10,000 Ns. Error correction of the ultrascaffolded assembly was performed with `Pilon v1.22` [18]. These corrected assemblies were the final versions used for downstream analyses. `Qualimap v2.2.1` [19] generated quality statistics for Illumina reads by using the final assemblies as reference for where reads were mapped (Table S2).

Complete mitochondrial genomes were filtered out from our new assemblies by screening the assembly scaffolds. Mitochondrial genome scaffolds were of length between 40-90 kbp and GC content lower than 30 %. The extrachromosomal 2-µm plasmid scaffolds were filtered out by detecting those scaffolds matching 2-µm plasmid genes and having a length lower than 7 kbp. We calculated the nuclear (Figure S6B) and mitochondrial GC content and length (Figure S6A,C) using the `infoseq` with flags `--auto -only -name -length -pgc` from the `EMBOSS package v6.5.7` [20]. The previously sequenced and assembled *S. jurei* mitochondrial genome was corrected to remove an artifactual duplicated region produced by the original PacBio assembly pipeline [21]; this correction reduced the previously described length from ~111 kbp to 79 kbp. The completeness of the nuclear genomes was quickly assed by exploring the number of single-copy orthologous genes annotated by `BUSCO v2.0.1` [22] using the `saccharomycetales_odb9` database (Figure S30).

Genome annotations of our new nuclear assemblies were performed with `YGAP` [23]. To minimize downstream analysis errors, we also re-annotated all previously published genome assemblies using `YGAP`. `YGAP` output was in GenBank format, which was converted to gff3 format in `Geneious`. Paralogous genes are not fully resolved by `YGAP`, so synteny and manual inspection was used to resolve the genomic location of each paralog. Mitochondrial genomes were annotated with `MFannot` [24]. 2-µm plasmids were manually annotated in `Geneious`. Genome assemblies and annotations are available in

GitHub (http://bit.ly/2orfKyT) and ENA accession no. PRJEB48264. A genome browser and BLAST server can also be accessed through https://gxseq.glbrc.org/assemblies.

Phenotyping strains

We phenotyped the 87 strains whose genomes we sequenced, as well as 41 *Saccharomyces* strains whose genomes had been previously sequenced, in 26 media conditions (Table S2, S6). We tested carbon sources (2.5 % glycerol; 2 % glucose; 2 % maltose; 2 % maltotriose; 2 % and 0.1 % fructose; 2 % sucrose; 2 % and 0.1 % raffinose; 2 %, 0.4 %, 0.2 %, and 0.1 % galactose; 2 % and 0.1 % melibiose; 2 % mannose; 2 % xylose; and 2 % arabinose) and stresses, including osmotic (30 % must juice, 1.4 M NaCl), pH 7.5 (5 g/L histidine), oxidative (2mM $H_2O_2$), low temperatures (4 °C and 10 °C), and high temperatures (30 °C and 37 °C). All conditions, except low temperatures and high temperatures, were performed at 22 °C. The medium composition was minimal medium (6.7 g/L Yeast Nitrogen Base without amino acids, carbohydrate and with Ammonium Sulfate, pH 5.0) supplemented with 2 % glucose when no other carbon sources or different concentrations are specified. Must or grape juice was prepared according to Clowers [25], except 30 % was our final concentration. Glucose and fructose concentrations of the must were measured by high-performance liquid chromatography (HPLC) following the procedure described in Peris [2]. 14 % fructose and 12 % glucose were detected in the 30 % must.

The 128 *Saccharomyces* strains were pre-cultured in deep 96-well plates with 500 µl of minimal medium supplemented with 0.2 % glucose until saturation at room temperature. After pre-culturing, we used a pinner to inoculate 96-well plates (Nunc, Roskild, Denmark) containing 240 µl of all of the media being tested. Initial optical densities at 600 nm ($OD_{600}$) were below 0.2 (mean 0.037 ± 0.027). These 96 well-plates were designed with corner containers, which we filled with 3 mL of $dH_2O$ to maintain humidity during culturing. As a control, each plate contained 5 wells with YPD medium (1 % yeast extract, 2 % peptone and 2 % glucose) where the *S. cerevisiae* S288C strain was inoculated. To monitor the growth of strains in the different conditions, the inoculated 96-well plates were placed in a stacker of a BMG FLUOstar Omega plate reader (BMG Labtech, Ortenberg, Germany) located inside an incubator with an interior temperature set to 22 °C. Absorbance at 600 nm was monitored every 2 hours for 6-8 days with no shaking. Absorbance of low temperature and high temperature experiments were manually monitored (3-4 data points per day) in a BMG FLUOstar Omega, and the experiments were stopped when saturation of growing strains was detected. Strain location in the 96-well plates was randomized in each replicate. Two $\rho^-$ strains, *S. eubayanus* yHCT96 [26] and Bond Lab 1063 (this study), were removed from the phenotyping analysis. Background absorbance was subtracted from the average of five negative controls (uninoculated media). Kinetic parameters for each condition were calculated using `GCAT v6.3` [27]. Average, median, and standard deviations of kinetic parameters from the three independent biological replicates were calculated in `R v4.0.2` [28] (Table S6).

Flocculation can generate artificially low or high OD values, depending on which part of the well is measured by the spectrophotometer. To correct for flocculation, we took pictures of the first replicate of each 96-well plate and manually inspected the kinetic parameters to correct for false positive and negative values. In cases where the pictures displayed no growth but there were high OD values (e.g. due to condensation on the lid), we set the growth of that particular strain to 0. In cases where growth was not detected but exaggerated OD values are observed and the pictures showed evidence for flocculation, we removed the exaggerated values and kept the parameter values from other runs close to those observed on closely related strains.

Boxplots for kinetic parameters by species (Figure S26) and by lineages (Figure S29) were drawn using `ggplot2 v3.3.3` [29] and `gridExtra v2.3` packages in `R`. The stacked barplots showing the percentages of strains that grew above $OD_{600}$=0.5 normalized by species (Figure S18) were drawn with `ggplot2`. Dotplots of the median maximum $OD_{600}$ by species (Figure 5B top) were drawn with `ggplot2`. The variance of the median maximum OD by growth condition for each species (Figure S23A) was calculated in `R` and plotted with `ggplot2`. The correlation of mean Tamura-Nei corrected genetic distance (see *Species tree, genetic boundaries among species and concordance factors* section) within species and the average phenotypic variance for each species was tested using the Spearman correlation test implemented in `ggscatter` function of the `R` package `ggpubr v0.4` [30], and the scatterplot (Figure S23B) was drawn with `ggplot2`. The heatmap with median maximum $OD_{600}$ values normalized to the most extreme $OD_{600}$ value for each condition (Figure 5B) was drawn, together with the phylogenetic tree (see *Phylogenomics and quantification of reticulate evolution of phenotyped strains* section), using `iTOL v4.2.3` [31]. A principal component analysis (PCA) was performed by `prcomp` function in `R` using the median maximum $OD_{600}$ calculated from replicated growth curves. Selected PCs were plotted (Figure 5A, S24A,C) with `ggbiplot v0.55` [32] package in `R`.

The percentage of variance explained by each component (Figure S24B) was plotted with `factoextra v1.0.7` [33] package in `R`. The percentage of variance explained by each growth condition for each component was calculated in `R`, and the histogram plot (Figure S25) was drawn with `ggplot2`.

<u>Read mapping and variant calling</u>

Illumina reads from 163 *Saccharomyces* strains were generated in this study or downloaded (Table S2). We mapped Illumina reads to a reference genome belonging to one of the *Saccharomyces* species, following our previously developed pipeline [34]. The *Saccharomyces cerevisiae* reference genome was DBVPG6044 [35]; for *Saccharomyces paradoxus*, it was CBS432 [35]; for *Saccharomyces mikatae,* IFO1815 (this study); for *Saccharomyces kudriavzevii*, IFO1802 (this study); for *Saccharomyces arboricola*, CBS10644 [36]; for *S. uvarum,* CBS7001 (this study); and for *S. eubayanus*, FM1318 [37]. Reference genomes consisted of the assigned chromosomes, and extra scaffolds were discarded for mapping.

Illumina reads were first mapped with `bwa v0.7.12` using the `bwa-mem` algorithm [38]. The resulting SAM files were viewed and sorted using `samtools v1.4` [39], filtering for high quality reads `samtools view -q 30` (except for *S. kudriavzevii*, *S. arboricola*, and *S. eubayanus* where we set the quality to `20`). PCR duplicates were removed with `picard v1.98` (http://picard.sourceforge.net/) using `MarkDuplicates.jar REMOVE_DUPLICATES=true AS=true VALIDATION_STRINGENCY=SILENT`. Read groups were set using `picard v1.98 AddOrReplaceReadGroups.jar` with settings

“VALIDATION_STRINGENCY=SILENT SORT_ORDER=coordinate CREATE_INDEX=true”. Single nucleotide polymorphisms (SNPs) were called using the GATK v3.1 [40] haplotype caller using the setting --genotyping_mode DISCOVERY -mbq 20 -stand_emit_conf 31 -stand_call_conf 31. Genome coverage was measured using BEDTOOLS v2.27.0 genomeCoverageBed -d –ibam [41]. The VCF output of GATK was converted into FASTA format using a custom python script. A specific FASTA file for each strain, now called its Whole Genome Sequence (WGS), was obtained by using the reference genome as a template and replacing the called variant with the SNP reported by GATK. The presence of heterozygous sites in the sample were coded according to their IUPAC ambiguity codes. In downstream analyses, we considered only the homozygous SNPs, which represented the vast majority of the genome (>99.4 %) due to the low levels of heterozygosity (Figure S15). Insertions and deletions were masked by replacing the genomic sequence with a number of Ns corresponding to the length of the indel called. FASTA files for each sample were generated by masking regions with extremely high coverage (i.e. values greater than the 99.9th percentile of genome-wide coverage) and by masking regions with low coverage (i.e. either regions below 10X coverage or, for genomes with low coverage, below the 10th percentile of genome-wide coverage). Masked regions were replaced by Ns prior to downstream analyses.

Population genomics and quantification of reticulate evolution

WGSs were aligned by species. Frequent gaps were removed using trimal v1.4 [42] with parameters –gt 0.9. We calculated the average distance within species and populations with MEGA v7 [43] using complete gap removal and Tamura-Nei correction (Figure 3C). Polymorphism statistics from the WGS dataset were calculated in DnaSP v5 [9] and a stacked barplot (Figure 3A) with those results was drawn with ggplot2.

To delimit the number of populations in each *Saccharomyces* species, we used the program STRUCTURE v2.3.4 [44-46] (Figure S8i) and fineSTRUCTURE v2.0.7 [47]. VCF files were merged with GATK using the CombineVariants parameter and -genotypeMergeOptions UNIQUIFY. 10,000 random SNPs from the VCF data were picked for STRUCTURE analysis. We tested K clusters from 1 to 8 (except for *S. cerevisiae* where more clusters were tested), assuming an admixture model, with a 10,000-iteration burn-in and 100,000 iterations of sampling. Five independent runs were performed for each K cluster. The STRUCTURE output was used as input for STRUCTURE HARVESTER web v0.6.94 [48] to select the most likely number of populations. STRUCTURE HARVESTER output files were aligned in CLUMPP v1.1.2 [49] and visualized in STRUCTURE PLOT v2 [50]. In addition to delimiting the likely number of populations, fineSTRUCTURE gave a deeper picture of co-ancestry among *Saccharomyces* strains (Figure S9iv). We converted the FASTA dataset with the complete set of SNPs to a PHASED format, the input format of fineSTRUCTURE. To reconstruct the co-ancestry heatmap with the linkage model and to perform a principal component analysis (Figure S9v) of the SNP dataset, fineSTRUCTURE was run with default parameters, except ¨–ploidy 1” due to the low heterozygosity in the dataset; the genetic distance map was inferred by applying the specific genetic distance for each chromosome described on the SGD database.

We reconstructed the phylogenetic tree and network using the SNP dataset for each species (Figure S9ii and iii, respectively). ML phylogenetic tree reconstruction was done in `RAxML` as above, after correcting branch lengths for the presence of invariant sites. The SNP dataset was also the input of `SplitsTree`, which we used to detect incongruent data. For *S. arboricola, S. eubayanus,* and *S. kudriavzevii* species datasets, the outgroup was CBS7001. For *S. cerevisiae*, the outgroup was CBS432; for *S. paradoxus*, the outgroup was S288C; for *S. mikatae*, the outgroup was NCYC3947; and for *S. uvarum*, the outgroup was FM1318.

Supported admixture strains were further analyzed to quantify the genome contributions of parental strain relatives. The WGS alignment dataset was split in alignments consisting of 50,000 bp to calculate genetic contributions (Figure S10) using `PopGenome v2.2.4` package in `R` [51]. For detecting the ancestry of *GAL* genes in two Chinese *S. kudriavzevii* strains and the two closest relatives (Figure S10Hi), the WGS alignment dataset was split consisting of 5,000 bp to calculate genetic contributions using `PopGenome` and a $\log_2$ divergence ratio was plotted (Figure S10Hii). *GAL* gene coordinates were used to locate the coding sequences in the plot. Quantification of genome introgression between species was performed by analyzing `sppIDer v1` [52] plots (Table S3, Figure S16).

Phylogenomics of nuclear, mitochondrial, and 2-µm plasmid genomes of phenotyped strains

To build a phylogenetic tree of the nuclear genomes (Figure 5B) for the phenotyped 128 *Saccharomyces* strains, we searched for a common set of complete orthologous gene sequences. First, we annotated single-copy orthologous genes of our phenotyped collection of *Saccharomyces* strains with `BUSCO`. We detected 18 orthologs common to all genome assemblies of phenotyped strains, but we selected 14 well-resolved gene trees: *ASF1* (*YJL115W*), *MAF1* (*YDR005C*), *NIF3* (*YGL221C*), *NSL1* (*YPL233W*), *PET117* (*YER058W*), *PLP2* (*YOR281C*), *QCR7* (*YDR529C*), *RPA34* (*YJL148W*), *SGF73* (*YGL066W*), *SHB17* (*YKR043C*), *SMT3* (*YDR510W*), *SUB1* (*YMR039C*), *TUB2* (*YFL037W*), and *YAR1* (*YPL239W*). We blasted those genes to pull out the orthologs from the strains that were not phenotyped, as well as *Kluyveromyces lactis* as an outgroup (Table S2). Orthologous genes were concatenated with `FASconCAT v1.0` [53]. A maximum-likelihood (ML) phylogenetic tree for a concatenated alignment (~8.7 Kbp), with frequent gaps trimmed with `trimal`, was reconstructed in `RAxML v8.1` [54], performing 100 iterations to search for the best tree, using the model GTRGAMMA. Bootstrap branch support was assessed by performing 1,000 pseudoreplicates using the same model parameters as above. The ML phylogenetic tree can be accessed together with the phenotypic data of phenotyped strains at `iTOL` (http://bit.ly/2PYRuUc). The same concatenated alignment was used in `SplitsTree 4` [55] to reconstruct the phylonetwork using the NeighborNet (NN) method (Figure S14B). We followed a similar pipeline to reconstruct the mitochondrial and 2-µm plasmid phylogenetic networks, instead using mitochondrial and 2-µm plasmid genes (Figure 2B and Figure S17, respectively). For the mitochondrial genome, we were focused on the coding sequences (CDS) of 69 sequenced *Saccharomyces* strains representing the diverse *Saccharomyces* lineages

(Table S2): *ATP6*, *ATP8*, *ATP9*, *COB*, *COX1*, *COX2*, *COX3*, 15S rRNA, 21S rRNA, and *VAR1* (~9.7 kbp). Mitochondrial phylogenetic networks for individual gene alignments were also reconstructed (Figure S4). *COX2*, *COX3*, *ATP6, ATP8,* and *ATP*9 included the sequences of all non-$\rho^-$ phenotyped *Saccharomyces* strains. For the 2-µm plasmid, we analyzed the frequently observed genes *REP1* (*R0020C*) and *REP2* (*R0040C*) (extrachromosomal tag in Figure S17), which were used to classify the plasmid sequences by class [56,57] (Table S4). It is noteworthy that some plasmid genes were inserted in the nuclear genome (nuclear tag in Figure S17).

Species tree, genetic boundaries among species, and concordance factors

To infer the phylogenetic relationships of our species and lineages and the degree of genome-wide support, we performed several phylogenomic analyses. We first selected 23 strains to represent key *Saccharomyces* lineages, most of them high-quality genomes generated here, and an additional 15 previously assembled genomes (Table S2). Then, the coding sequences and amino acid sequences annotated with `YGAP` were extracted using Daniel Jeffares' `perl` script `process_gff_cds_proteins.pl` [58]. For each gene, two types of files were generated: one file containing all species'/lineages' CDS and another file containing all species'/lineages' protein sequences. Amino acid sequences were aligned using `MAFFT v7.21` [59], using the setting "`--preservecase --maxiterate 1000 --genafpair`". Amino acid alignments were back-translated to nucleotides using `pal2nal v14` [60]. Codon columns with gaps were removed from the alignments using `trimal 1.4.1 "-gt 1 –block 3"` [42]. Gene sequences present in all specimens that retained at least 50 % of positions and with equal or more than 300 nucleotides (100 amino acids) were selected for additional analyses. A total of 3850 genes passed our filters. ML phylogenetic trees from each CDS alignment were calculated in `IQTree v1.6.12` [61] following recommendations by Shen et al. [62]. The next settings were used in `IQTree` "`-bb 1000 -wbt -nt AUTO -seed 225494 -st DNA -m TEST`". The coalescent species tree was generated using the collection of ML phylogenetic trees in `ASTRAL v5.7.7` [63]. The gene concordance factor (gCF) (Figure 4A) for each branch in the coalescent species tree (reference tree) was assessed using all individual gene trees as an input for `IQTree v2.0.3`.

To assess reciprocal monophyly of each gene, we followed the bioinformatic pipeline developed by Peris *et al.* [64]. Briefly, ML phylogenetic trees were read in `R` using `treeio v1.12` [65] and converted to `ape v5.4` format [66]. Once species designations were associated with phylogenetic tip labels, the trees were rooted in the branch generating the *S. eubayanus* and *S. uvarum* clades. Monophyly tests were performed using `spider v1.5` [67], and when the test for one species was FALSE (Table S5), the tree was printed to a `TIFF` file for visual exploration. Trees were drawn using `R` package `ggtree v2.2.4` [68]. The monophyly test suggested that 0.77 % of genes were incorrectly annotated (e.g. due to cryptic paralogy) in at least one of the 38 genomes.

We reconstructed the concordance tree topology, which was congruent with the `ASTRAL` coalescent tree, and we inferred the branch support (CF, Figure 4A) using the `BUCKy v1.4.4` [69] pipeline as done previously [26]. Due to memory issues with large datasets in `BUCKy`, we reduced the number of *Saccharomyces* strains to 10. Strains were selected based on their Asian origin when possible. We included a *Kluyveromyces lactis* strain as an outgroup (Table S2). Before running `BUCKy`, the gene alignments were the input for `MrBayes v3.2.3` [70] for a Bayesian phylogenetic reconstruction. 49 genes failed to be parsed through the pipeline, so 3801 genes were analyzed. Settings in `MrBayes` were "`lset nst=6 rates=gamma; prset brlenspr=Unconstrained:Exp(50.0); mcmc nruns=2 temp=0.2 ngen=110000 burninfrac=0.0909; Nchains=4 samplefreq=10 swapfreq=10 printfreq=50000; mcmcdiagn=yes diagnfreq=50000`". Sample trees from each MCMC run were summarized for each gene with `mbsum` after a burn-in of 1000 trees. `BUCKy` was run with all collected mbsum files with settings "`-a 1 -k 3 -n 100000 -c 2 --calculate-pairs --create-joint-file --create-single-file`". `BUCKy` generated the posterior probability that pairs of loci share the same tree, which we represented in a histogram (Figure S13).

Boundaries among *Saccharomyces* species were calculated using the CDS alignments as an input. Distributions of genetic distance (Figure S11) and relative divergence (Fst) (Figure S12) were calculated in `R`. Tamura-Nei corrected genetic distance was calculated with `dist.dna` as implemented in the `R` package `ape v5.4`. Fst was calculated by the `R` package `PopGenome v2.7.5` [51].

<u>*GAL/MEL* pathway characterization</u>

To characterize the *GAL/MEL* pathway from the phenotyped strains, genes were retrieved from: i) the genome assemblies using the `YGAP` annotation files; ii) genome assemblies using a local `BLAST v2.6` [7]; iii) raw Illumina reads by using `HybPiper v1.2` [8]; or iv) PCR and Sanger sequencing of strains of interest (Table S7). ML phylogenetic trees for individual genes from the *GAL*/*MEL* pathway were reconstructed in `IQTree`, with similar settings as above. Phylogenetic trees were read and manipulated with `R` packages `treeio`, `ape`, and `phytools v0.7` [71], and drawn using `ggtree`. Conclusions about gene presence/absence and phylogenetics were displayed in heatmaps (Figure 6A, S27B) using the `R` package `ggplot2`.

To confirm the unexpected absence of the second copy of *GAL2* in some *S. eubayanus* and *S. uvarum* strains, we performed PCR and Sanger sequencing using primers and conditions described in Table S7. To optimize PCR conditions, we first performed gradient PCR, and the optimal annealing temperature was selected for amplifying the target region (Table S7). For amplifying long regions (expected length > 6 kbp, Table S7), LongAmp polymerase (New England Biolabs, Ipswich, MA, USA) was used, instead of Taq polymerase (New England Biolabs, Ipswich, MA, USA). Sanger-sequenced *GAL* sequences were deposited in GenBank under accession nos. OL660614-OL660618.

<u>Amino acid identity comparison between animals and *Saccharomyces*</u>

To compare the divergence among animals and among *Saccharomyces* species and lineages, we annotated a common set of single copy orthologous genes with `BUSCO v5.1.3` [72] using the `eukaryota_odb10` database. We first downloaded the genome assemblies for *Homo sapiens* GRCh38p13 (GCA_000001405.28), *Pan troglodytes*

ClintPTRv2 (GCA_002880755.3), *Macaca mulata* AG07107 (GCA_003339765.3), *Mus musculus* C57BL6J (GCA_000001635.9), *Takifugu rubripes* fTakRub1 (GCA_901000725.2), and *Gallus gallus domesticus* bGalGal1 (GCF_016699485.2). Then, we ran `BUSCO` on those genomes and high-quality genomes for *Saccharomyces* strains with no detected admixture or introgressions (Table S2). Each organism's amino acid sequences for each protein were pulled together. Amino acid alignments were performed using `MAFFT` with similar settings as above. Individual protein alignments were read in `R` with `seqinr v4.2` package [73]. Amino acid identity (AAI) values (Figure 3B) for each protein between lineages of the same *Saccharomyces* species, between *Saccharomyces* species, and between the chosen animals were calculated using `dist.alignment` function implemented in `seqinr` with the `"matrix=identity"` setting. Mean AAI values for each comparison was plotted with `ggplot2`.

28 R Development Core Team, "R: a Language and Environment for Statistical Computing,"in (Vienna, Austria: R Foundation for Statistical Computing, 2010).

29 H Wickham, *ggplot2: elegant graphics for data analysis* (Springer, NY, 2009).

30 Wickham, H., "Ggpubr: 'Ggplot2' Based Publication Reaqy Plots,"in 2021).

31 Letunic, I. and Bork, P., "Interactive tree of life (iTOL) v3: an online tool for the display and annotation of phylogenetic and other trees," *Nucl. Acids Res.* **44**, W242-W245 (2016).

32 Vu, V., "Ggbiplot,"in 2015).

33 Alboukadel Kassambara, *practical guide to cluster analysis in r: unsupervised machine learning*2017).

34 Peris, D., *et al.*, "Complex ancestries of lager-brewing hybrids were shaped by standing variation in wild yeast *Saccharomyces eubayanus*," **12**, e1006155 (2016).

35 Yue, J. X., *et al.*, "Contrasting evolutionary genome dynamics between domesticated and wild yeasts," *Nat Genet* **49**, 913-924 (2017).

36 Liti, G., *et al.*, "High quality *de novo* sequencing and assembly of the *Saccharomyces arboricolus* genome," *BMC Genomics* **14**, 69 (2013).

37 Baker, E., *et al.*, "The genome sequence of *Saccharomyces eubayanus* and the domestication of lager-brewing yeasts," *Mol. Biol. Evol.* **32**, 2818-2831 (2015).

38 Li, H. and Durbin, R., "Fast and accurate short read alignment with Burrows-Wheeler transform," **25**, 1754-1760 (2009).

39 Li, H., *et al.*, "The Sequence Alignment/Map format and SAMtools," **25**, 2078-2079 (2009).

40 McKenna, A., *et al.*, "The Genome Analysis Toolkit: a MapReduce framework for analyzing next-generation DNA sequencing data," *Genome Res.* **20**, 1297-1303 (2010).

41 Quinlan, A. R. and Hall, I. M., "BEDTools: a flexible suite of utilities for comparing genomic features," **26**, 841-842 (2010).

42 Capella-Gutiérrez, S., Silla-Martínez, J. M., and Gabaldón, T., "trimAl: a tool for automated alignment trimming in large-scale phylogenetic analyses," **25**, 1972-1973 (2009).

43 Kumar, S., Stecher, G., and Tamura, K., "MEGA7: Molecular Evolutionary Genetics Analysis version 7.0 for bigger datasets," *Mol. Biol. Evol.* **33**, 1870-1874 (2016).

44 Pritchard, J. K., Stephens, M., and Donnelly, P., "Inference of population structure using multilocus genotype data," *Genetics* **155**, 945-959 (2000).

45 Falush, D., Stephens, M., and Pritchard, J. K., "Inference of population structure using multilocus genotype data: linked loci and correlated allele frequencies," *Genetics* **164**, 1567-1587 (2003).

46 Hubisz, M., *et al.*, "Inferring weak population structure with the assistance of sample group information," *Mol Ecol Resour* **9**, 1322-1332 (2009).

47 Lawson, D. J., *et al.*, "Inference of population structure using dense haplotype data," **8**, e1002453 (2012).

48 Earl, D. and vonHoldt, B., "STRUCTURE HARVESTER: a website and program for visualizing STRUCTURE output and implementing the Evanno method," **4**, 359-361 (2012).

49 Jakobsson, M. and Rosenberg, N. A., "CLUMPP: a cluster matching and permutation program for dealing with label switching and multimodality in analysis of population structure," **23**, 1801-1806 (2007).

50 Ramasamy, R. K., *et al.*, "STRUCTURE PLOT: a program for drawing elegant STRUCTURE bar plots in user friendly interface," *SpringerPlus* **3**, 431 (2014).

51 Pfeifer, B., *et al.*, "PopGenome: an efficient Swiss Army Knife for population genomic analyses in R," *Mol. Biol. Evol.* **31**, 1929-1936 (2014).

52 Langdon, Q. K., *et al.*, "sppIDer: a species identification tool to investigate hybrid genomes with high-throughput sequencing," *Mol Biol Evol* **35**, 2835-2849 (2018).

53 Kück, P. and Meusemann, K., "FASconCAT, Version 1.0,"in (Zool. Forschungsmuseum A. Koenig, Germany, 2010).

54 Stamatakis, A., "RAxML version 8: a tool for phylogenetic analysis and post-analysis of large phylogenies," **30**, 1312-1313 (2014).

55 Huson, D. H., *et al.*, "Phylogenetic Super-Networks from partial trees," *IEEE/ACM Trans. Comput. Biol. Bioinformatics* **1**, 151-158 (2004).

56 Strope, P. K., *et al.*, "2µ plasmid in *Saccharomyces* species and in *Saccharomyces cerevisiae*," *FEMS Yeast Res.* **15**, fov090 (2015).

57 Peter, J., *et al.*, "Genome evolution across 1,011 *Saccharomyces cerevisiae* isolates," *Nature* **556**, 339-344 (2018).

58 Jeffares, D., "Perl Scripts,"in (figshare, figshare, 2016).

59 Katoh, K. and Standley, D. M., "MAFFT multiple sequence alignment software version 7: improvements in performance and usability," *Mol. Biol. Evol.* **30**, 772-780 (2013).

60 Suyama, M., Torrents, D., and Bork, P., "PAL2NAL: robust conversion of protein sequence alignments into the corresponding codon alignments," *Nucl. Acids Res.* **34**, W609-W612 (2006).

61 Nguyen, L. T., *et al.*, "IQ-TREE: A fast and effective stochastic algorithm for estimating maximum likelihood phylogenies," *Mol. Biol. Evol.* **32**, 268-274 (2014).

62 Shen, X. X., *et al.*, "An investigation of irreproducibility in maximum likelihood phylogenetic inference," *Nature Communications* **11**, 6096 (2020).

63 Zhang, C., *et al.*, "ASTRAL-Pro: quartet-based species-tree inference despite paralogy," *Mol. Biol. Evol.* **37**, 3292-3307 (2020).

64 Peris, D., *et al.*, "Large-scale fungal strain sequencing unravels the molecular diversity in mating loci maintained by long-term balancing selection," **In press** (2022).

65 Wang, L. G., *et al.*, "Treeio: an R package for phylogenetic tree input and output with richly annotated and associated data," *Mol. Biol. Evol.* **37**, 599-603 (2019).

66 Paradis, E. and Schliep, K., "ape 5.0: an environment for modern phylogenetics and evolutionary analyses in R," **35**, 526-528 (2018).

67 Brown, S. D. J., *et al.*, "Spider: An R package for the analysis of species identity and evolution, with particular reference to DNA barcoding," *Mol Ecol Resour* **12**, 562-565 (2012).

68 Yu, G., *et al.*, "ggtree: an R package for visualization and annotation of phylogenetic trees with their covariates and other associated data," *Methods Ecol Evol* , **8**, 28-36 (2016).

69 Larget, B. R., *et al.*, "BUCKy: Gene Tree / Species Tree reconciliation with bayesian concordance analysis," **26**, 2910-2911 (2010).

70 Ronquist, F., *et al.*, "MrBayes 3.2: efficient bayesian phylogenetic inference and model choice across a large model space," *Syst Biol* **61**, 539-542 (2012).

71 Revell, L. J., "phytools: an R package for phylogenetic comparative biology (and other things)," *Methods Ecol Evol* **3**, 217-223 (2012).

72 Waterhouse, R. M., *et al.*, "BUSCO applications from quality assessments to gene prediction and phylogenomics," *Mol. Biol. Evol.* **35**, 543-548 (2018).

73 Delphine Charif and Jean R. Lobry, "SeqinR 1.0-2: A Contributed Package to the R Project for Statistical Computing Devoted to Biological Sequences Retrieval and Analysis,"in 2007), p.207.
