## Supplementary material for "Macroevolutionary diversity of traits and genomes in the model yeast genus *Saccharomyces*": SuppFiguresTextANDOnlineMethods: Peris_etal_SuppFig-Notes_SSS2.pdf

### Supplementary Notes

#### Supplementary Note 1 – Nuclear and mitochondrial genome diversity

The lengths of chromosomes II, IV, VI, VIII, X, XIII, and XV differed among *Saccharomyces* strains. The differences in length were driven mainly by the presence of chromosomal translocations (Figure S5, Table S2). Most *S. cerevisiae*, most *S. paradoxus*, and all *S. kudriavzevii* strains were syntenic with *S. cerevisiae* S288C. The other *Saccharomyces* species had particular translocations relative to *S. cerevisiae* S288C. All *S. mikatae* and *S. jurei* strains share a common translocation between chromosomes VI and VII, as noted in a previous study [1]. *S. uvarum* and *S. eubayanus* share two translocations: between chromosomes II and IV and between VIII and XV. All *S. uvarum* populations fixed three translocations (IItIV, VItX and VIIItXV), except *S. uvarum* Australasia. *S. uvarum* Australasia maintains synteny with *S. eubayanus* genomes, where VItX is absent (Table S2, Figure S5). Both *S. arboricola* populations share a translocation between chromosomes IV and XIII. Particular strains from *S. paradoxus*, such as the America B strain UFRJ50816, America C strain yHDPN24, and the Hawaii strain UWOPS91-917.1, contain one or multiple new translocations. *S. mikatae* Asia A and *S. cerevisiae* Malaysia populations are also differentiated from other populations of the same species by unique translocations.

The *S. cerevisiae*, *S. paradoxus*, S. *jurei*, and *S. mikatae* genomes have lower GC-contents; *S. arboricola* and *S. kudriavzevii* genomes have intermediate GC-contents; and *S. eubayanus* and *S. uvarum* genomes have higher GC-contents (Figure S6B). *Saccharomyces eubayanus* and *S. uvarum* grew better at lower temperatures (Figure

S26C-E), whereas in bacteria, a positive correlation between optimal temperature and GC-content has been observed [2]. Some species that grew better at lower temperatures (Figure 5, 26C-E), such as *S. arboricola*, *S. eubayanus*, and *S. uvarum*, also have the smallest mitochondrial genomes (Figure S6C).

#### Supplementary Note 2 – *Saccharomyces* phenotypic diversity

*S. cerevisiae* was the most phenotypically diverse species (Figure 5A, Figure S23, S24), but some species also showed high phenotypic variance in particular conditions, often driven by lineage-specific phenotypic traits. For example, *S. kudriavzevii* had strains able to grow in 2 % galactose, 5 g/L histidine, and 4 °C, while other *S. kudriavzevii* strains could not. Some strains of *S. cerevisiae* grew in 2 mM $H_2O_2$, most of which were from domesticated lineages (Figure S29A). Wild populations of *S. cerevisiae* (Asia Islands, CHN IV, Malaysia, and North America) were more phenotypically similar than the domesticated lineages (Figure S24A). In general, conditions with more diversity within species were 2 % galactose, 2 % maltose, 2 % melibiose, 5 g/L histidine, and 4 °C (Figure S23A). Strains with negative values of PC1 (Figure 5A) were characterized by better growth in 2 % galactose and their ability to produce biomass from 2 % xylose (Figure 5B, S24A, S25). We note that all *Saccharomyces* strains produce little to no biomass from 2 % xylose, but the normalization accentuates the differences amongst the strains producing some biomass (Figure S18). PC2 classified strains by their ability to grow at 37 °C (positive values) or at 4 °C (negative values) (Figure S25). PC3 also split strains by their ability to grow at lower temperatures (4 °C and 10 °C) (Figure S24C, S25). The best performers at 4°C were *S. arboricola, S. eubayanus*, and *S. uvarum*, while the best performers at 37 °C were *S. cerevisiae* and some strains from *S. paradoxus* (Figure S26C-E). This suggests that low temperature growth is an ancestral trait (Figure 6B, S18).

*Saccharomyces* strains grew well in the presence of glucose, fructose, sucrose, raffinose, mannose, and in complex media, such as juice must, at temperatures of 22 °C (Figure S18). However, there were conditions where particular species and populations performed better than other strains. In general, sugar concentration affected kinetic parameters. For example, growth rate in both galactose and fructose generally increased as concentrations increased (Figure S21A-B). Despite this general trend, maximum growth rate at 2 % galactose was slightly lower than 0.4 % or had no further increase in *S. cerevisiae*, *S. paradoxus*, and *S. arboricola* (Figure S20A). In the case of galactose, in all species except *S. mikatae*, the increase of maximum growth rate in response to increased concentrations was faster in strains with a higher total number of putative Gal4-binding sites upstream of *PGM1* and *PGM2*, which likely allows enhanced flux from the *GAL* pathway into glycolysis (Figure S20E) [3]. Interestingly, lag time in both galactose and fructose did not have the same trend as growth rate; in fact, lag time in fructose was longer at a higher concentration for *S. arboricola*, *S. uvarum*, and *S. eubayanus* (Figure S20D). Surprisingly, at the level of individual strains, for both galactose and fructose, a majority of strains (59 % for galactose and 76 % for fructose) showed a shorter lag time in 0.1 % than in 2 % (dots on the upper left side of the gray dash line) (Figure S21A-B). Lag time variation on disaccharides or trisaccharides at the strain level may be partly explained by the growth variation on their constituent monosaccharides or disaccharides (Figure S22).

In the presence of maltose, non-domesticated *S. cerevisiae, S. arboricola, S. kudriavzevii* Asia A, *S. mikatae, S. eubayanus* Holarctic, and *S. uvarum* Australasia grew poorly (Figure S29). Some *S. mikatae* and *S. arboricola* Asia A strains were found to produce more biomass in the presence of melibiose than other *Saccharomyces* (Figure S18, S26C, S27C). Brewing and some *S. cerevisiae* wine/European strains were able to grow in the presence of maltotriose (Figure 5B, S29A), but surprisingly, some *S. mikatae* strains may also have been able to produce limited biomass from this carbon source (Figure S18, S26C, S29C). European *S. kudriavzevii,* except two strains from a Europe-Asia A admixed lineage (Figure S9D), grew on galactose as expected [4], while non-admixed strains from Asia A and Asia B did not (Figure 5B, 6B, S29D). As expected [5], *S. cerevisiae* West Africa did not grow well in the presence of galactose (Figure 5B, S29A). Some domesticated *S. cerevisiae*, *S. mikatae* Asia A*, S. kudriavzevii,* non-Holarctic *S. eubayanus,* and *S. uvarum* strains were able to grow at a higher pH (5 g/L histidine, pH 7.5) (Figure 5B, S18, S26, S29). *S. eubayanus* Patagonia B and few domesticated *S. cerevisiae* strains were more osmotolerant (1.4 NaCl) than other *Saccharomyces* strains (Figure S18, S26C, S29G). With the exception of many *S. eubayanus* and *S. uvarum* strains, few *Saccharomyces* strains grew on glycerol during the timeframe of the experiment (Figure 5B, S26C).

Some growth conditions promoted flocculation in specific *Saccharomyces* strains (Figure S19). In particular, mannose, 10 °C, 37 °C, histidine, and galactose were the most influential conditions, with more than 5 *Saccharomyces* strains flocculating in each.

#### Supplementary Figures

##### Figure S1. Geographic locations of *Saccharomyces* populations.

Panels A-G) show the geographic distribution of wild populations for each *Saccharomyces* species. Note that anthropic strains are not shown for clarity. The size of the symbols represent the number of strains from each locations. Symbol colors represent species designations according to the legend (Table S1). CHN: China; EU: Europe; FE: Far East; HOL: Holarctic; NA: North America; SA-A: South America A; SA-B: South America B. The map was generated using the `map_data` function implemented in `R` package `ggplot2` [6].

**Figure S2. Association biases for hosts and substrates of *Saccharomyces* strains.**

A) A stacked barplot for *Saccharomyces* strains isolated from different hosts (Table S1). The taxonomic rank of order was used to group *Saccharomyces* isolates by their host order. Human-related environments, such as vineyards, were grouped in the "Anthropic" category. B) A stacked barplot for wild *Saccharomyces* strains isolated from different substrates (Table S1). Bar plots are colored according to species.

#### **Figure S3**. ***COX3*** **phylogenetic network of** ***Saccharomyces*** **strains.**

A Templeton, Crandall, and Sing (TCS) phylogenetic network of 395 trimmed *COX3* sequences from wild *Saccharomyces* strains is shown. The *COX3* haplotype classification for the wild and anthropic *Saccharomyces* strains is shown in Table S1. Haplotypes are represented by circles, and haplotype names are colored based on species designations. When two different species shared the same haplotype, the haplotype name has both colors and is highlighted with an asterisk. Circle size is scaled according to the haplotype frequency. Pie charts show the frequency of haplotypes based on the biogeographic realm of origin. The number of mutations separating each haplotype is indicated by lines on the edges connecting different haplotype circles. Strain names for those yeasts phenotyped or studied in the main text are indicated. Population and strain names are colored according to their species designations. Afr: Africa; CHN: China; EU: European; HOL: Holarctic; Med: Mediterranean; NA-Jp: North America-Japan (=North America); SA-A: South America A; SA-B: South America B; W/EU: Wine/European.

**Figure S4. Phylogenetic networks of mitochondrial genes.**

Neighbor-Net phylogenetic networks for informative mitochondrial genes are shown in panels A-G). For the *COB* gene, the *S. mikatae* Asia B position is indicated with a discontinuous arrow due to an incomplete *COB* gene reconstruction. Strains found in an unexpected location are noted and colored according to their species designations. Populations are designated for all species, except for *S. cerevisiae* when all *S. cerevisiae* populations are clustered together. Scale bars represent nucleotide substitutions per site. *Candida castellii* (syn. '*Nakaseomyces*' *castellii*) is the outgroup.

**Figure S5. Genome dot plots of *Saccharomyces* strains compared to the *S. cerevisiae* S288C laboratory strain.**

Dot plot comparing the location of genes in the *Saccharomyces* strains with their location in *S. cerevisiae* (S288C). Blue dots indicate inversions. Strain names are colored according to their species designations. Chromosome and scaffolds boundaries are marked on the axes. Chromosomal translocations compared to S288C are highlighted in the coalescent phylogenetic tree (Figure 4A). Roman numerals represent chromosomes.

**Figure S6. Highly diverse genomic architectures among *Saccharomyces* species.**

Chromosome length, percent of GC in the nuclear genome, and mitochondrial genome (mtDNA) size for *Saccharomyces* strains are shown in panel A), B), and C), respectively. Dots or boxplots are colored according to the species designations. In panel B), the dots correspond to the GC-content for each chromosome per strain. The dashed blue line in panel C) corresponds to the median mitochondrial genome size of all mitochondrial genomes (Table S2).

**Figure S7. Mitochondrial genome dot plots of *Saccharomyces* strains compared to the *S. cerevisiae* S288C laboratory strain.**

Dot plot comparing the location of mitochondrial genes in the *Saccharomyces* strains with their location in *S. cerevisiae* (S288C). For consistency, all mitochondrial genomes were oriented to set the first nucleotide as the gene encoding tRNA-Serine, which is close to *VAR1*. Blue dots indicate inversions. Strain names are colored according to their species designations. Axes are represented in base pairs, and dashed lines are plotted every 10 kbp. Regions relocated compared to S288C are highlighted in the coalescent phylogenetic tree (Figure 4A).

**Figure S8. Mitochondrial genome dot plots of *Saccharomyces* populations compared to other populations.**

Dot plots comparing the location of mitochondrial genes in *Saccharomyces* populations with their location in other populations to highlight the diversity in the architectures among them. For consistency, all mitochondrial genomes were oriented to set the first nucleotide as the gene encoding tRNA-Serine, which is close to *VAR1*. Blue dots indicate inversions. Population names were colored according to their species designations. Axes are represented in base pairs, and dashed lines are plotted every 10 kbp.

**Figure S9. Population genomics of seven *Saccharomyces* species.**

Population genomic analyses for *S. cerevisiae*, *S. paradoxus*, *S. mikatae*, *S. kudriavzevii*, *S. arboricola*, *S. uvarum*, and *S. eubayanus* are shown in panels A), B), C), D), E), F), and G), respectively. i) Inference of the genetic clusters (*K*) and composition of individuals utilizing a random subsample of 10,000 SNPs in STRUCTURE. The most consistent number of genetic clusters/populations is highlighted in bold. Two summary plots for the most consistent number of genetic clusters/populations, from five independent runs are shown, except for *S. arboricola* because increasing K gave the same membership plot. Each color in the bar plots represents the cluster membership coefficients. The presence of several colors in the same strain suggests admixture. ii) Maximum likelihood phylogenetic tree reconstructed using all SNPs and corrected for invariant sites (see Online Material and Methods). The scale bars show the number of substitutions per site. Bootstrap values above 75 are reported at their corresponding nodes. Colored boxes enclose strains by their lineage designation. iii) Neighbor-Net phylogenetic network reconstructed with the SNP dataset. Incongruent data are represented by nodes subtended by multiple edges. iv) Coancestry heatmap where darker colors indicate higher coancestry between strains. Hypothesized donor strains are on the y-axis, while hypothesized recipients are on the x-axis. Colored bars indicate populations. In *S. cerevisiae*, for simplification, populations are described on the top tree branches, and larger groups were described in the bars. The *S. cerevisiae* groups are described according to the PCA in panel v. v) Principal Component

Analysis (PCA) plots of PC1 versus PC2. The percent of the variance accounted for by each component is indicated. For *S. cerevisiae*, due to the low genetic diversity among domesticated strains and their close relatives, we reanalyzed the dataset by removing CHN I, CHN II, CHN IX, and the Taiwanese strains.

**Figure S10. Genome-wide pairwise nucleotide sequence divergence plots for admixture *Saccharomyces* strains.**

Pairwise nucleotide divergence comparisons for admixture strains of five different *Saccharomyces* species compared to representative strains from the potential donor population are shown in panels A-Hi,I-N. The percentage of pairwise divergence for 50-kbp windows is shown on the y-axis. Roman numerals represent chromosomes. Strain names are colored according to their species designations. Panel H ii) represents the $\log_2$ divergence ratio of the pairwise nucleotide divergence comparisons calculated in panel H i), but for 5-kbp windows. Ancestry is indicated by arrows at the right of the plot. In panel H ii), we include the location of genes involved in galactose utilization (Figure 6A), whose phylogenetic tree reconstruction is displayed in Figure S28I-K. Holarctic-NA: Holarctic-North America. gd: average genome-wide divergence of the portion of the genome contributed by that population.

#### Figure S11. Genetic distance distributions.

Tamura-Nei corrected genetic distance distribution for comparisons between species using all annotated `YGAP` orthologous genes (3858 genes). Each panel, from A) to H), represents the comparison of a species against all other species with lines colored according to the species compared. Panel I) represents the genetic distance within a species with each line colored according to the legend. TN93: Tamura-Nei 1993 model correction.

**Figure S12. Fst distributions.**

Relative divergence (Fst) distribution for comparisons between species using all annotated `YGAP` orthologous genes (3858 genes). Each panel represents the comparison of a species against the other species with lines were colored according to the species compared.

**Figure S13. BUCKy concordance primary tree and alternative topologies.**

Distribution of the most probable Bayesian tree topologies for each `YGAP` orthologous gene (from a set of 3802) for a selection of 10 strains, mostly from Asia, chosen to represent each species or the most divergent population within a species. Tip circles are colored according to the species designation.

#### Figure S14. Phylogenomic network of *Saccharomyces* single-copy orthologous genes.

Neighbor-Net phylogenetic network reconstructed with a concatenated alignment of 3859 (~5.5 Mbp) YGAP genes for 38 strains (panel A), as well as for 14 (11.5 Kbp) single-copy orthologous (BUSCO) genes for 160 strains (panel B). Strain names in A), and linages in B), are colored according to their species designations. The scale bar represents the number of substitutions per site. *Kluyveromyces lactis* is the outgroup. A ML phylogenetic tree of 14 BUSCO genes is shown in Figure 5B, and the schematic representation of the ML phylogenetic tree is shown in Figure 4Bi-ii.

**Figure S15. Levels of heterozygosity among *Saccharomyces* strains.**

Boxplots show the percentage of heterozygous sites across the genomes for each *Saccharomyces* strain, grouped by species. Color dots correspond to the species designations. Admixture, anthropic, and wild strains are indicated by a circle, triangle, or a square shape, respectively.

**Figure S16. Introgressions between *S. cerevisiae* and *S. paradoxus*.**

`sppIDer` plots of three *S. cerevisiae* strains with evidence of *S. paradoxus* introgressions are shown in panels A-C). The y-axis is the average coverage depth. As reference genomes, we used a representative high-quality genome for each *Saccharomyces* lineage. Strain names are colored according to their species designations.

**Figure S17.** ***Saccharomyces* 2-µm plasmid inheritance.**

Neighbor-Net phylogenetic networks for *REP1* (panel A) and *REP2* (panel B) 2-µm plasmid genes. *Saccharomyces* population, species 2-µm plasmid class, and strain names of interest are colored according to their species designations. Class D was colored in black due to its uncertain species designation. Asterisks highlight sequences that were too short or incomplete and were individually explored to confirm their relationships. The scale is given in nucleotide substitutions per site.

**Figure S18. Percentage of *Saccharomyces* that grew above $OD_{600}$=0.5 in various growth conditions.**

A stacked barplot for *Saccharomyces* strains growing above $OD_{600}$=0.5 is shown. Values were normalized by species such that each species represents 1/7 of the total, and each species is colored according to its species designation. Bars are colored according to the species designations.

**Figure S19. Growth conditions promoting flocculation among *Saccharomyces* strains.**

A stacked barplot shows the number of *Saccharomyces* strains flocculating in various growth conditions. Bars are colored according to the species designations.

**Figure S20. Growth variation in simple sugars across concentrations and the impact of Gal4-binding sites.**

A) The maximum growth rate distribution of each species across galactose concentrations from 0.1 % to 2 %. B) The lag time distribution of each species across galactose concentrations from 0.1 % to 2 %. Panels C) and D) show dot plots for the maximum growth rate and lag time, respectively, for each species at 0.1 % fructose and 2 % fructose. E) Slope of maximum growth rate against galactose concentrations plotted against the number of putative Gal4-binding sites ($CGGN_{11}CCG$) upstream (maximum 1 Kbp distance) of *PGM1/2*; *S. mikatae* strains are shown in the subpanel on the right because they did not show the correlation. ANOVA followed by Tukey's test was performed in A, B, and E: significant differences (*p*-value < 0.05) are indicated by different letters, n.s.: not significant. Unpaired two-tailed t-test was performed in C and D: *: *p*-value < 0.05, **: *p*-value < 0.01, ***: *p*-value < 0.001, ****: *p*-value < 0.001. Violin plots (panels A and B) were built with `vioplot 0.3.2 R` package [7], and dotplots (Panels C to E) were drawn with `GraphPad Prism 8` (RRID:SCR_002798).

**Figure S21. Lag time and maximum growth rate correlations between low and high sugar concentrations.**

Correlation for lag time and maximum growth rates of strains cultured at low (0.1 %) and high (2 %) concentrations of galactose (A) and fructose (B). Linear regression lines are plotted as black solid lines. The dashed gray line is y=x. Data points are colored according to their species designations. $R^2$ values from linear regression and *p*-values calculated from F-tests for regression are indicated in the top right corner in each plot. Plots were drawn with Igor Pro 9 (RRID:SCR_000325).

**Figure S22. Lag time correlations between monosaccharides and their disaccharides or trisaccharides.**

A) Lag time on sucrose (y-axis) is plotted by variation in its constituent monosaccharide units, glucose (x-axis) and fructose (color-coded by lag time). B) Lag time on melibiose (y-axis) is plotted by variation in its constituent monosaccharide units, glucose (color-coded by lag time) and galactose (x-axis). C) Lag time on raffinose (y-axis) is plotted by variation in the constituent disaccharides sucrose (x-axis) and melibiose (color-coded by lag time). Data point sizes are proportional to the lag time on fructose, which is generally the first constituent monosaccharide unit consumed, leaving behind melibiose. Each data point represents an average of three biological replicates from one strain. Data for a total of 96 strains are shown (*S. arboricola*=5, *S. cerevisiae*=15, *S. eubayanus*=22, *S. kudriavzevii*=7, *S. mikatae*=18, *S. paradoxus*=17, *Suva*=12). Plots were drawn with `Igor Pro 7` (RRID:SCR_000325).

**Figure S23. Phenotypic variance across *Saccharomyces* species.**

A) Phenotypic variance in maximum $OD_{600}$ across the genus *Saccharomyces* is shown for several growth conditions. Median variance is represented by a horizontal line inside the box, and the upper and lower whiskers represent the highest and lowest values of the 1.5*IQR (inter-quartile range), respectively. Growth conditions with unusually high variance for a species are noted close to their values. Gal: Galactose; His: Histidine; Mal: Maltose; Maltr: Maltotriose; Mel: Melibiose. B) Spearman correlation test of the average Tamura-Nei corrected genetic distance within species and the average of phenotypic variance for the species.

**Figure S24. Principal component analysis of maximum $OD_{600}$.**

A higher image resolution PCA (Figure 5A) of PC1 and PC2 of the maximum $OD_{600}$ as calculated from growth curves ($n$ = 3) (Table S6) is shown in panel A). PC1 and PC2 accounted for 37.4 % of the total variation. Growth condition weights (Figure S25) are represented by black arrows. Strains are colored according to their species designations, and different shapes represent their lineage/group designations. The percentage of variance explained by each component is shown in panel B). A PCA of PC1 and PC3 accounting for 33.6 % of the total variation is shown in panel C). The groups in panel A) and C) are defined as follows:

i) *S. cerevisiae*: Group1 (Domesticated strains: Bioethanol, Beer 1 & 2, Wine/European, Sake), Group 3 (West African), Group 4 (CHN IV), Group 5 (Asian Islands, Malaysian, North American).
ii) *S. paradoxus*: Group 1 (European), Group 2 (Far East), Group 3 (America B), Group 4 (America C).
iii) *S. mikatae*: Group 1 (Asia A), Group 2 (Asia B).
iv) *S. kudriavzevii*: Group 1 (EU), Group 2 (Asia A), Group 3 (Asia B).
v) *S. arboricola*: Group 1 (Asia A), Group 2 (Oceania).
vi) *S. uvarum*: Group 1 (Holarctic), Group 2 (South America A), Group 3 (South America B), Group 4 (Australasia).
vii) *S. eubayanus*: Group 1 (Holarctic), Group 2 (Patagonia B), Group 3 (Patagonia A).

**Figure S25. Variance contributed to each component by growth condition.**

The variances contributed to PC1, PC2, and PC3 by each growth condition is shown in bar plots. Ara: Arabinose; Fru: Fructose; Gal: Galactose; Gly: Glycerol; His: Histidine; Mal: Maltose; Maltr: Maltrotriose; Man: Mannose; Mel: Melibiose; Raf: Raffinose; Suc: Sucrose; Xyl: Xylose.

**Figure S26. Kinetic parameters of *Saccharomyces* strains in different growth conditions.**

*Saccharomyces* species violin boxplots of adaptation time (lag time, hours), maximum specific growth rate (µ, defined as $(\ln(OD_2)-\ln(OD_1))/(T_2-T_1)$), and biomass production (maximum $OD_{600}$) are represented in panels A), B), and C), respectively. Boxplots are colored according to their species designations. Panels D) and E) show the maximum biomass production ($OD_{600}$) at 4°C and 37°C temperatures in minimal media with 2 % glucose growth conditions, respectively. Names of the top-performing strains of each species at 4 °C and 37 °C are shown. Data points are split based on the species designations, and dashed lines connect the median values for each species. F) Median maximum growth rates for *S. eubayanus* and *S. uvarum* strains with and without *GAL2B* genes. Shapes highlight different *Saccharomyces* populations/groups. Dotted lines connect the median values of all *Saccharomyces* species. 2_4C_Glu: 2 % glucose at 4°C; 2_10C_Glu: 2 % glucose at 10°C; 2_22C_Glu: 2 % glucose at 22°C; 2_30C_Glu: 2 % glucose at 30°C; 2_37C_Glu: 2 % glucose at 37°C; 2_Mal: 2 % maltose; 2_Maltr: 2 % maltotriose; 2_Fru: 2 % fructose; .1_Fru: 0.1 % fructose; 2_Suc: 2 % sucrose; 2_Raf: 2 % raffinose; .1_Raf: 0.1 % raffinose; 2_Gal: 2 % galactose; .4_Gal: 0.4 % galactose; .2_Gal: 0.2 % galactose; .1_Gal: 0.1 % galactose; 2_Mel: 2 % melibiose; .1_Mel: 0.1 % melibiose; 2_Man: 2 % mannose; 2_Xyl: 2 % xylose; 2_Ara: 2 % arabinose; 30_Must: 30 % must; 5_His: 5 g/L histidine; 1.4_NaCl: 1.4 mM NaCl; 2_H2O2: 2mM $H_2O_2$; 2.5_Gly: 2.5 % glycerol. The groups are defined as follows:

i) *S. cerevisiae*: Group1 (Domesticated strains: Bioethanol, Beer 1 & 2, Wine/European, Sake), Group 3 (West African), Group 4 (CHN IV), Group 5 (Asian Islands, Malaysian, North American).

ii) *S. paradoxus*: Group 1 (European), Group 2 (Far East), Group 3 (America B), Group 4 (America C).

iii) *S*. *mikatae*: Group 1 (Asia A), Group 2 (Asia B).

iv) *S*. *kudriavzevii*: Group 1 (EU), Group 2 (Asia A), Group 3 (Asia B).

v) *S*. *arboricola*: Group 1 (Asia A), Group 2 (Oceania).

vi) *S*. *uvarum*: Group 1 (Holarctic), Group 2 (South America A), Group 3 (South America B), Group 4 (Australasia).

vii) *S*. *eubayanus*: Group 1 (Holarctic), Group 2 (Patagonia B), Group 3 (Patagonia A).

**Figure S27. Melibiose phenotypic diversity generated through complex genomic ancestries.**

The *GAL*/*MEL* pathway is represented in panel A). Proteins or activities specific to *S. eubayanus* and *S. uvarum* are colored according to the species' color [3,8]. *Saccharomyces* strains with complex ancestries for the *GAL/MEL* pathway genes are shown in panel B). Names of strains with genome-wide admixture (Table S3) are boxed. Incomplete gene sequences due to low coverage are labelled as black. Complete genes with a phylogenetic position (Figure S28) as expected based on population genomic analysis (Figure S9) are labeled as white. Genes acquired from another lineage by gene flow (within species) or introgression (between species) are labelled orange. Genes with premature stop codons or in a more advanced state of pseudogenization are labelled gray. Genes with a complex ancestries are labelled cyan. Genes not detected by any of the methods employed in this study (see Online Material and Methods) were considered to have been evolutionarily lost and are labelled red. Maximum biomass production ($OD_{600}$) in 2 % melibiose is shown in panel C). Each point is a strain colored by its species designation. Data were split based on the absence, gene flow/introgression, presence, or pseudogene state of *MEL1*. The groups in panel C) are defined as follows:

i) *S. cerevisiae*: Group1 (Domesticated strains: Bioethanol, Beer 1 & 2, Wine/European, Sake), Group 3 (West African), Group 4 (CHN IV), Group 5 (Asian Islands, Malaysian, North American).

ii) *S. paradoxus*: Group 1 (European), Group 2 (Far East), Group 3 (America B), Group 4 (America C).

iii) *S*. *mikatae*: Group 1 (Asia A), Group 2 (Asia B).

iv) *S*. *kudriavzevii*: Group 1 (EU), Group 2 (Asia A), Group 3 (Asia B).

v) *S*. *arboricola*: Group 1 (Asia A), Group 2 (Oceania).

vi) *S*. *uvarum*: Group 1 (Holarctic), Group 2 (South America A), Group 3 (South America B), Group 4 (Australasia).

vii) *S*. *eubayanus*: Group 1 (Holarctic), Group 2 (Patagonia B), Group 3 (Patagonia A).

**Figure S28. Individual phylogenetics trees of the *GAL/MEL* pathway.**

Maximum Likelihood (ML) phylogenetic trees of individual genes from the *GAL/MEL* pathway are represented in panels A-J. Strain names are colored according to their species designations. Branch support was assessed by using the UltraFast (UF) bootstrap method implemented in `IQTree`. Nodes with UF bootstrap higher than 50 % are reported and colored according to the legend. Panel K) displays the Neighbor-Joining (NJ) phylogenetic tree for the *S. kudriavzevii* genes, pseudogenes, and intergenic sequence (covering from one neighboring gene to the other, in the case of *GAL7/GAL10/GAL1*) alignments. Branch support was assessed by using 1,000 non-parametric bootstrap resampling and the Maximum Composite Likelihood model in `MEGA v5`. Panel B) and G) display the ML phylogenetic tree for the Gal2 and Gal80 protein sequences of *Saccharomyces*, respectively. The scale bars for all panels represented the number of nucleotide substitutions per site, except panel B) and G) where scale bars represented the number of amino acid substitutions per site. In panel B) Gal2 and Gal2B proteins are indicated by A or B, respectively. In panel G) Gal80 and Gal80B proteins are indicated by A and B, respectively. Phylogenetic location of T73Bio, DBVPG6044, and Y55 were manually added due to the high degree of gene degeneration, but our findings are supported by recent analyses with additional *S. cerevisiae* and *S. paradoxus* strains [9].

**Figure S29. Maximum $OD_{600}$ violin boxplots of *Saccharomyces* populations/groups.**

Populations/groups of *Saccharomyces* violin boxplots for biomass production (maximum $OD_{600}$) are shown in different panels and colored according to their species designations. Shapes highlight different *Saccharomyces* populations/groups. Dotted lines connect the median values of each *Saccharomyces* population/group. 2_4C_Glu: 2 % glucose at 4°C; 2_10C_Glu: 2 % glucose at 10°C; 2_22C_Glu: 2 % glucose at 22°C; 2_30C_Glu: 2 % glucose at 30°C; 2_37C_Glu: 2 % glucose at 37°C; 2_Mal: 2 % maltose; 2_Maltr: 2 % maltotriose; 2_Fru: 2 % fructose; .1_Fru: 0.1 % fructose; 2_Suc: 2 % sucrose; 2_Raf: 2 % raffinose; .1_Raf: 0.1 % raffinose; 2_Gal: 2 % galactose; .4_Gal: 0.4 % galactose; .2_Gal: 0.2 % galactose; .1_Gal: 0.1 % galactose; 2_Mel: 2 % melibiose; .1_Mel: 0.1 % melibiose; 2_Man: 2 % mannose; 2_Xyl: 2 % xylose; 2_Ara: 2 % arabinose; 30_Must: 30 % must; 5_His: 5 g/L histidine; 1.4_NaCl: 1.4 mM NaCl; 2_H2O2: 2mM $H_2O_2$; 2.5_Gly: 2.5 % glycerol.

**Figure S30.** **Summary statistics of *Saccharomyces* genome assemblies.**

Stacked bar plots for the four categories (single-copy, duplicated, fragmented, and missing) of genes annotated by `BUSCO` [10] for each *Saccharomyces* strain assembled. Strain names were colored according to their species designations. Genomes were classified as high, mid, and low quality according to the percentage of single-copy `BUSCO` genes (> 90 % High, between 70 % and 90 % Mid, and < 70 % Low).
