## Supplementary figures and images for "Macroevolutionary diversity of traits and genomes in the model yeast genus *Saccharomyces*"

### FigureS1.pdf

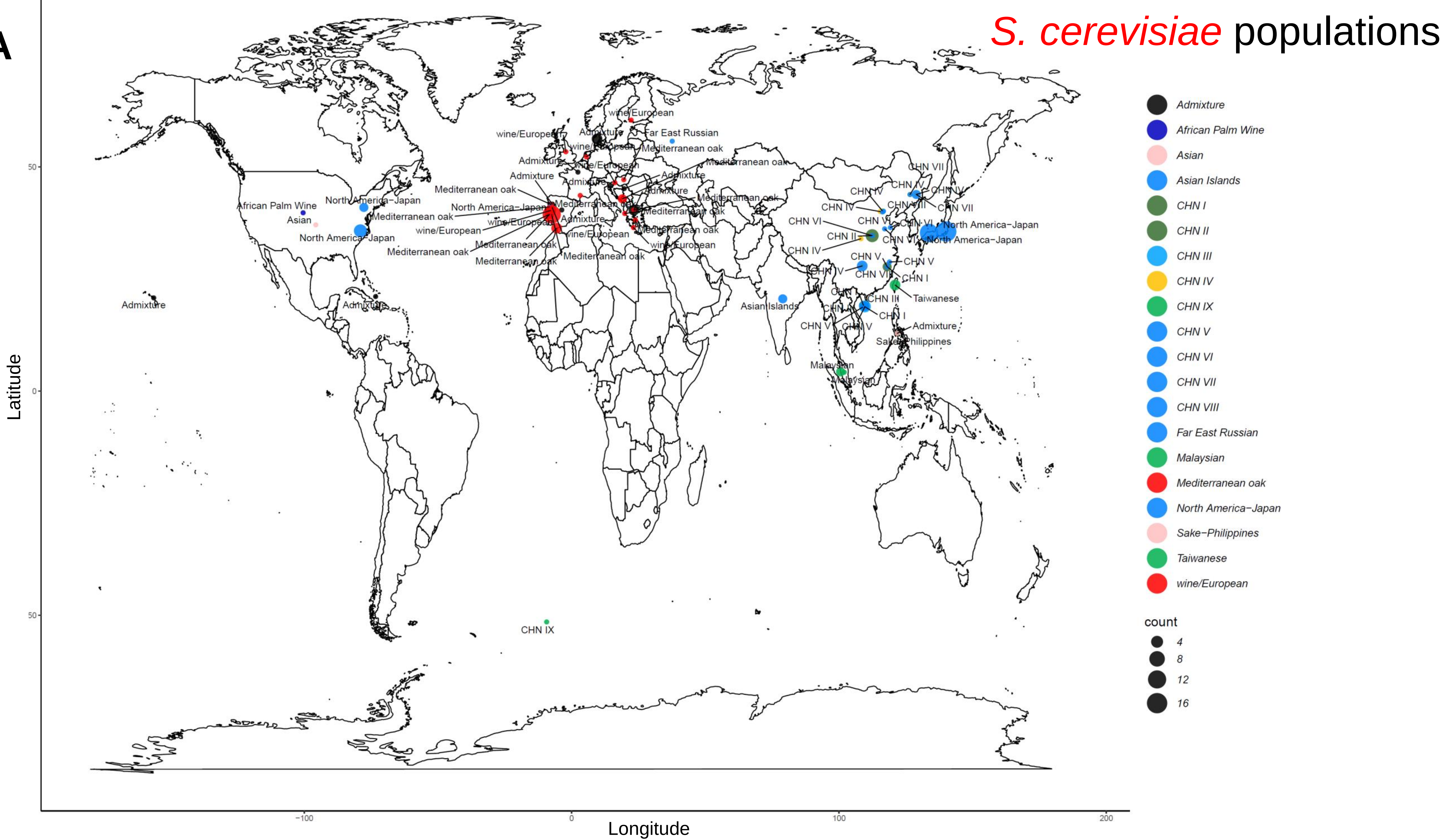

**B**

#### *S. paradoxus* populations

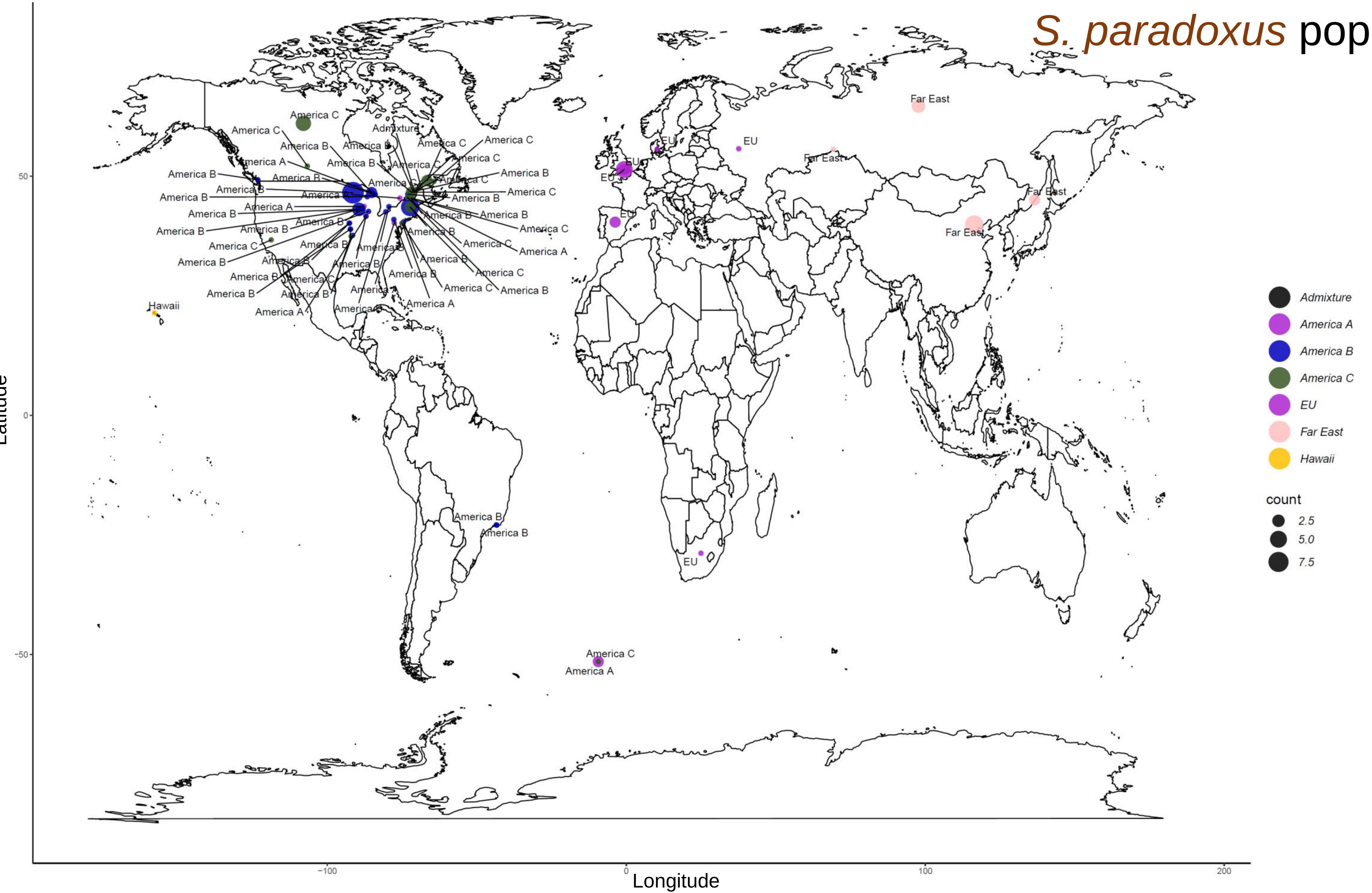

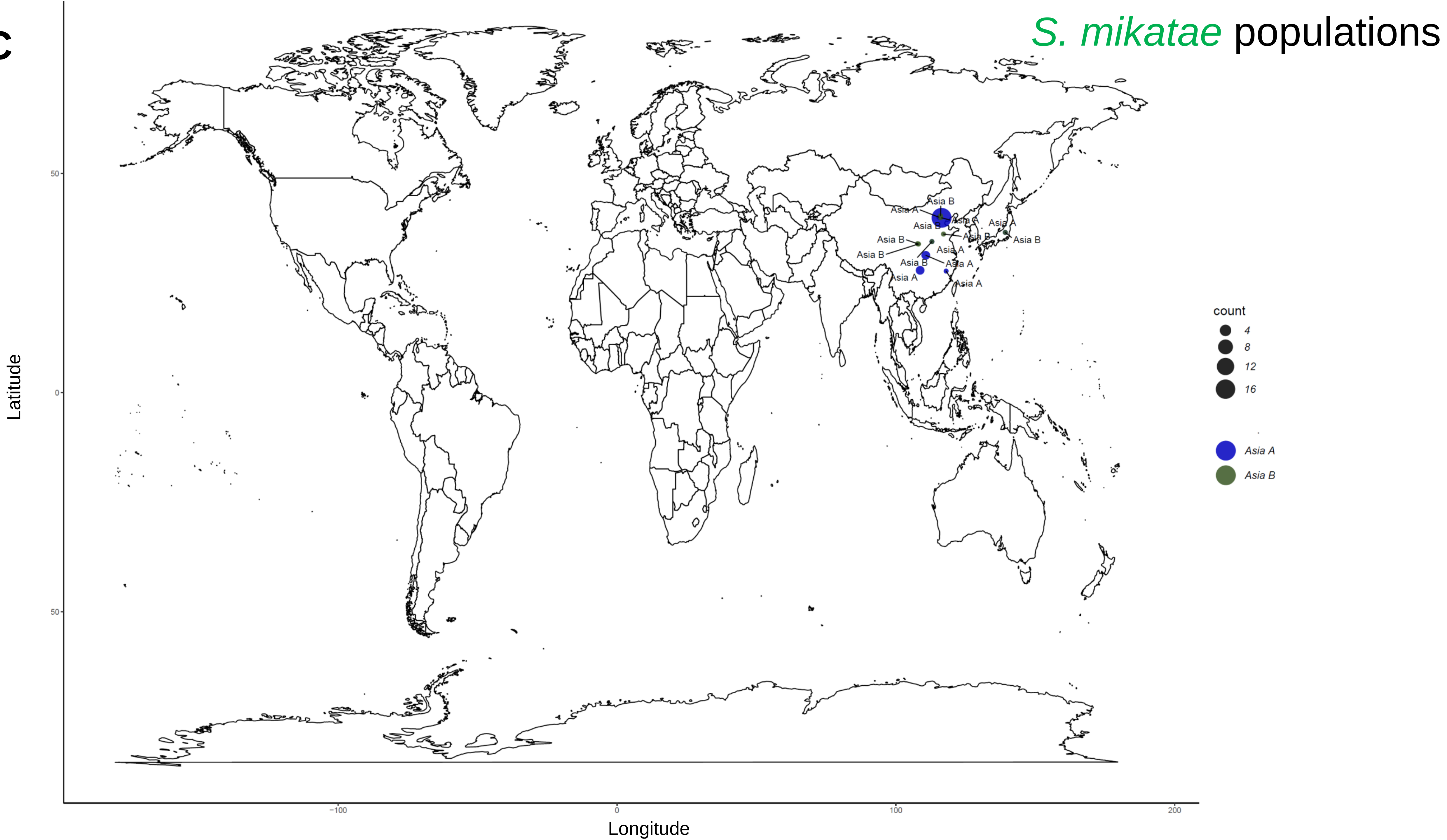

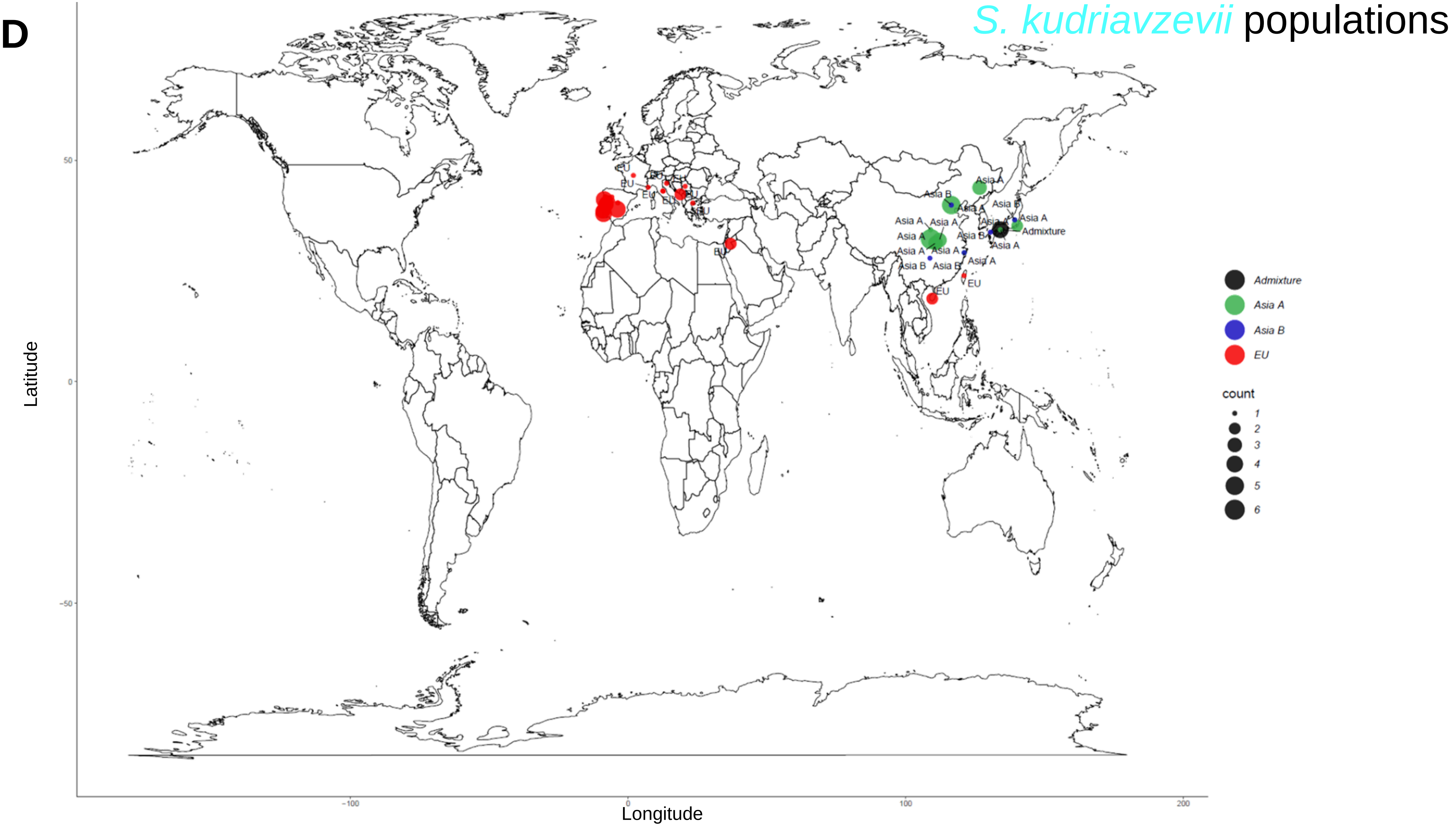

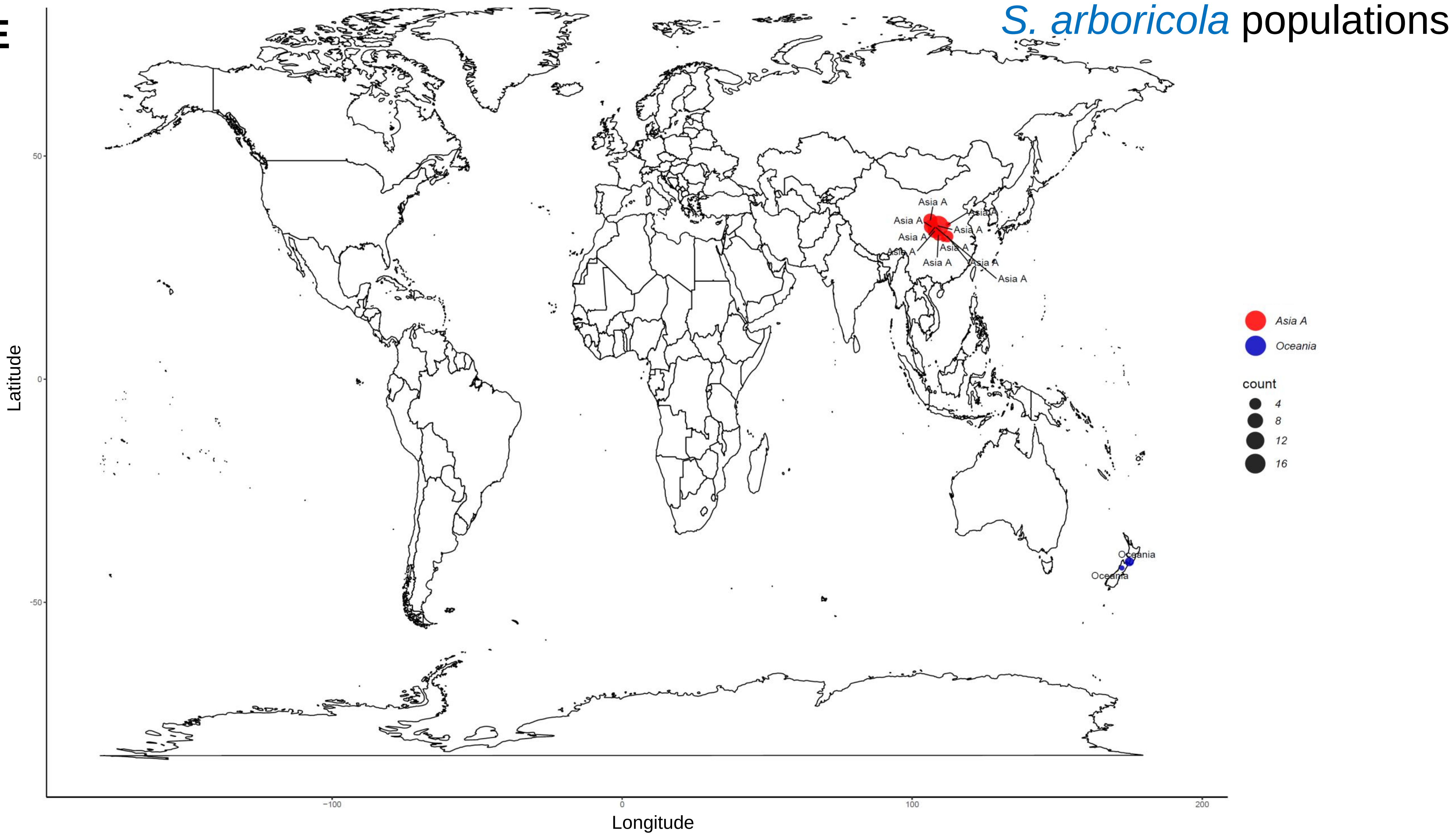

**F**

##### *S. uvarum* populations

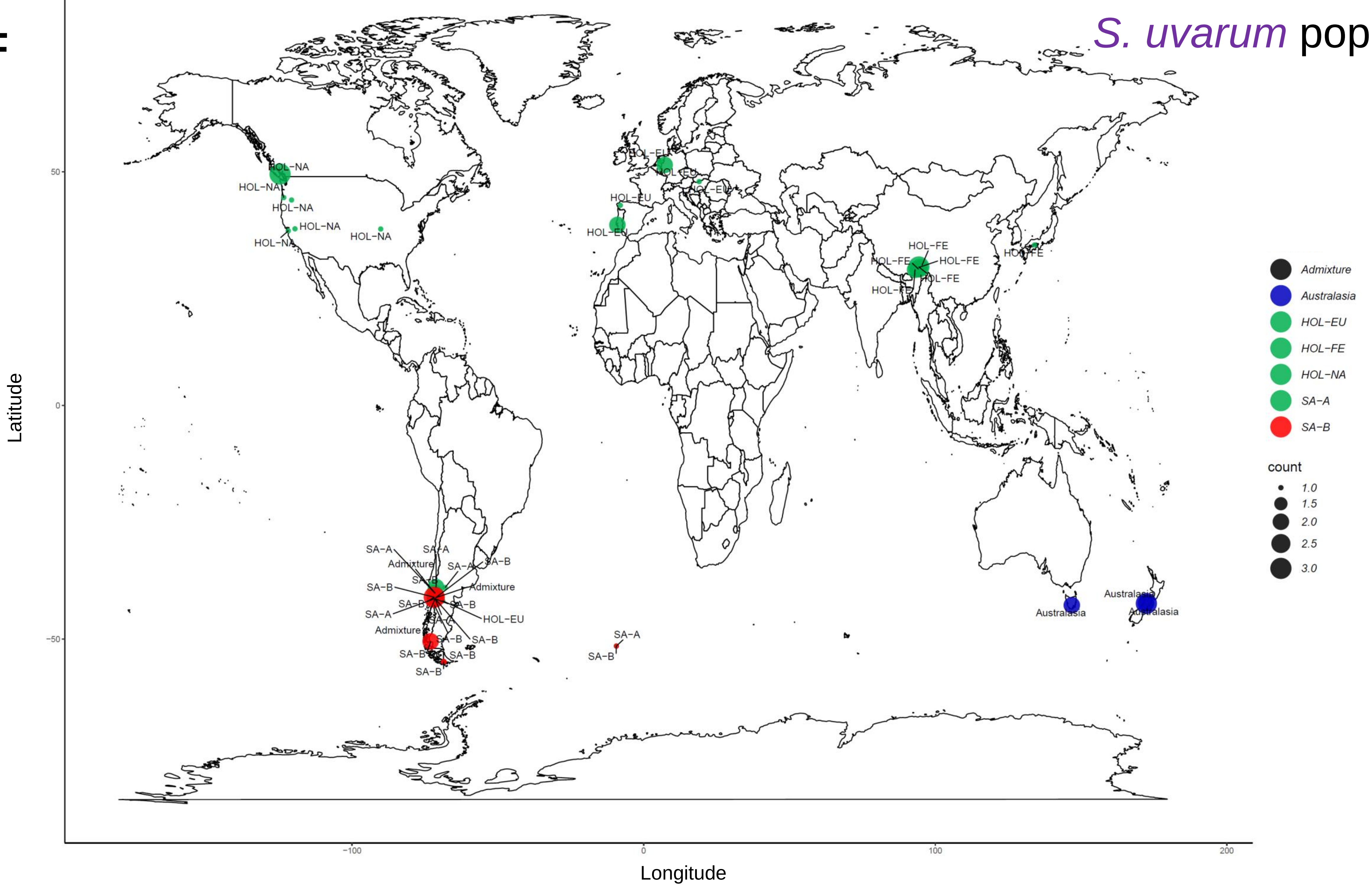

G

*S. eubayanus* populations

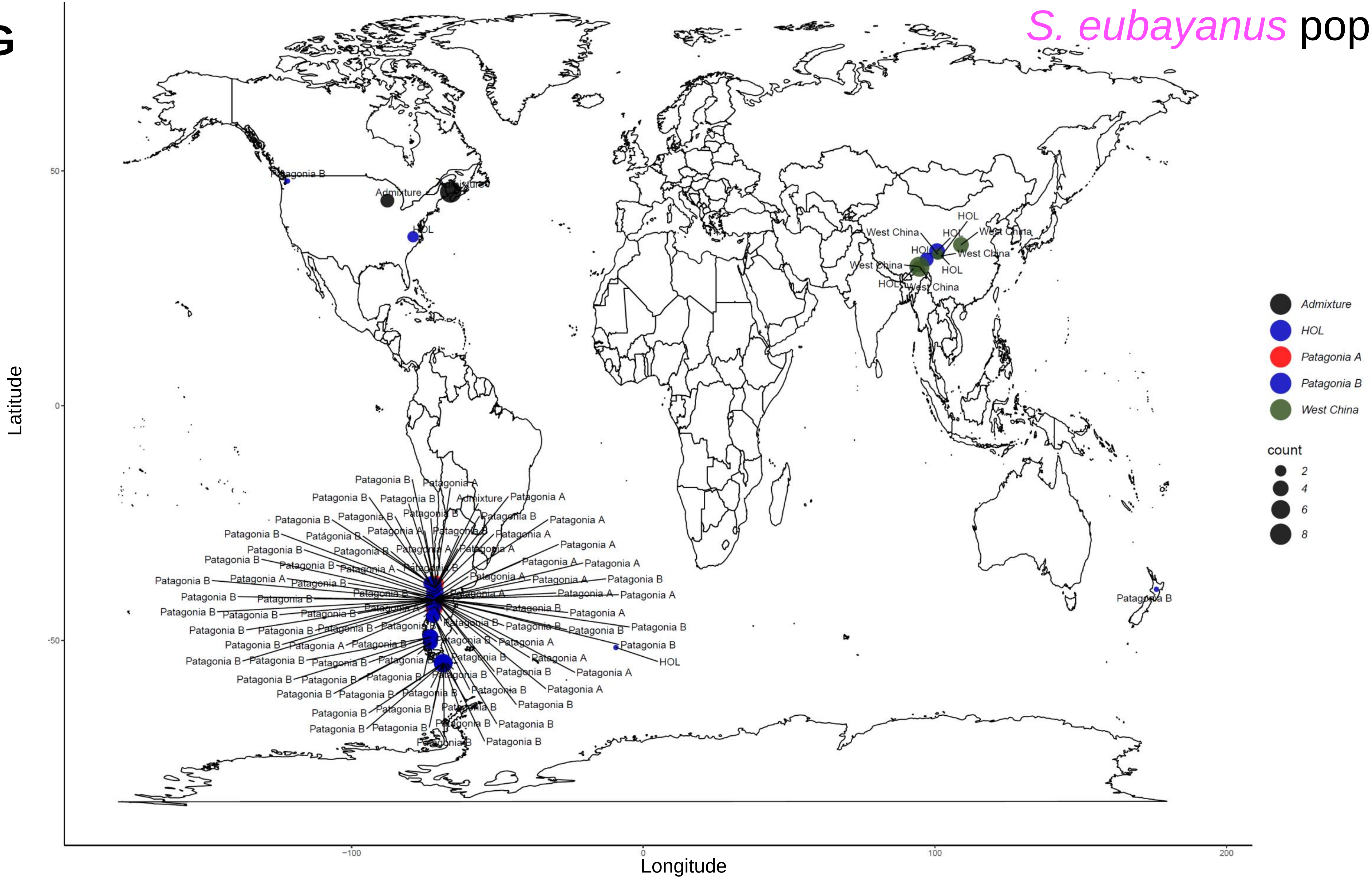

### FigureS2.pdf

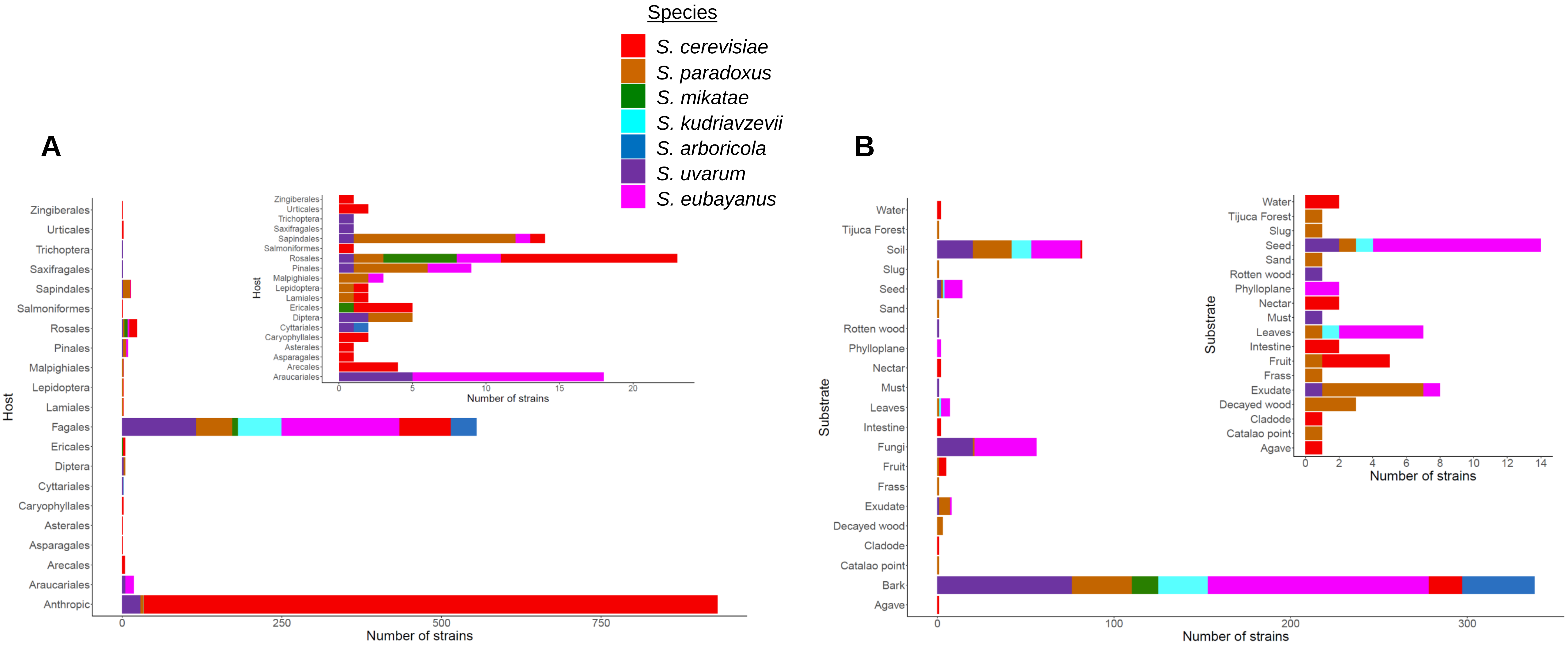

### FigureS3.pdf

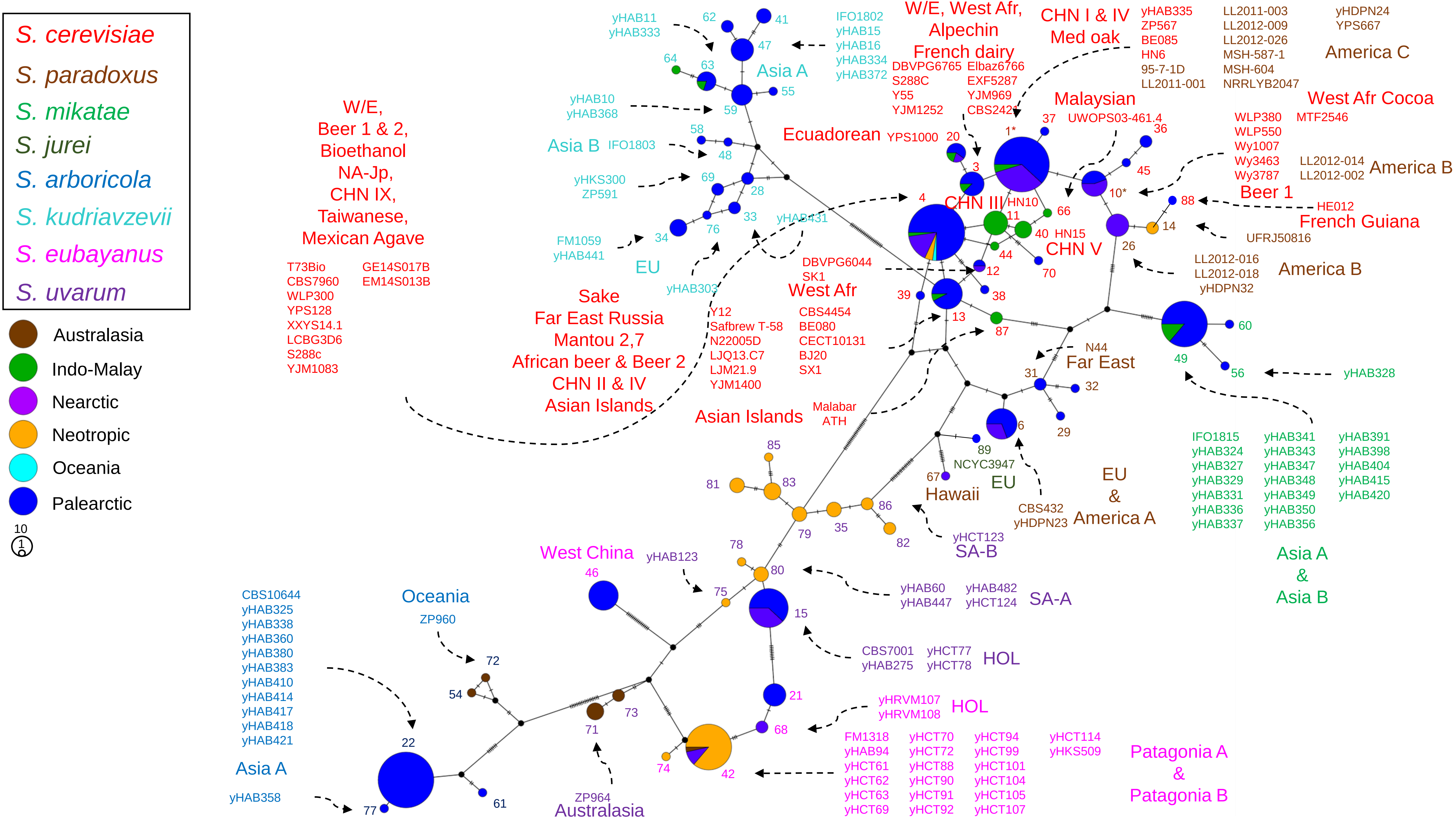

### FigureS4.pdf

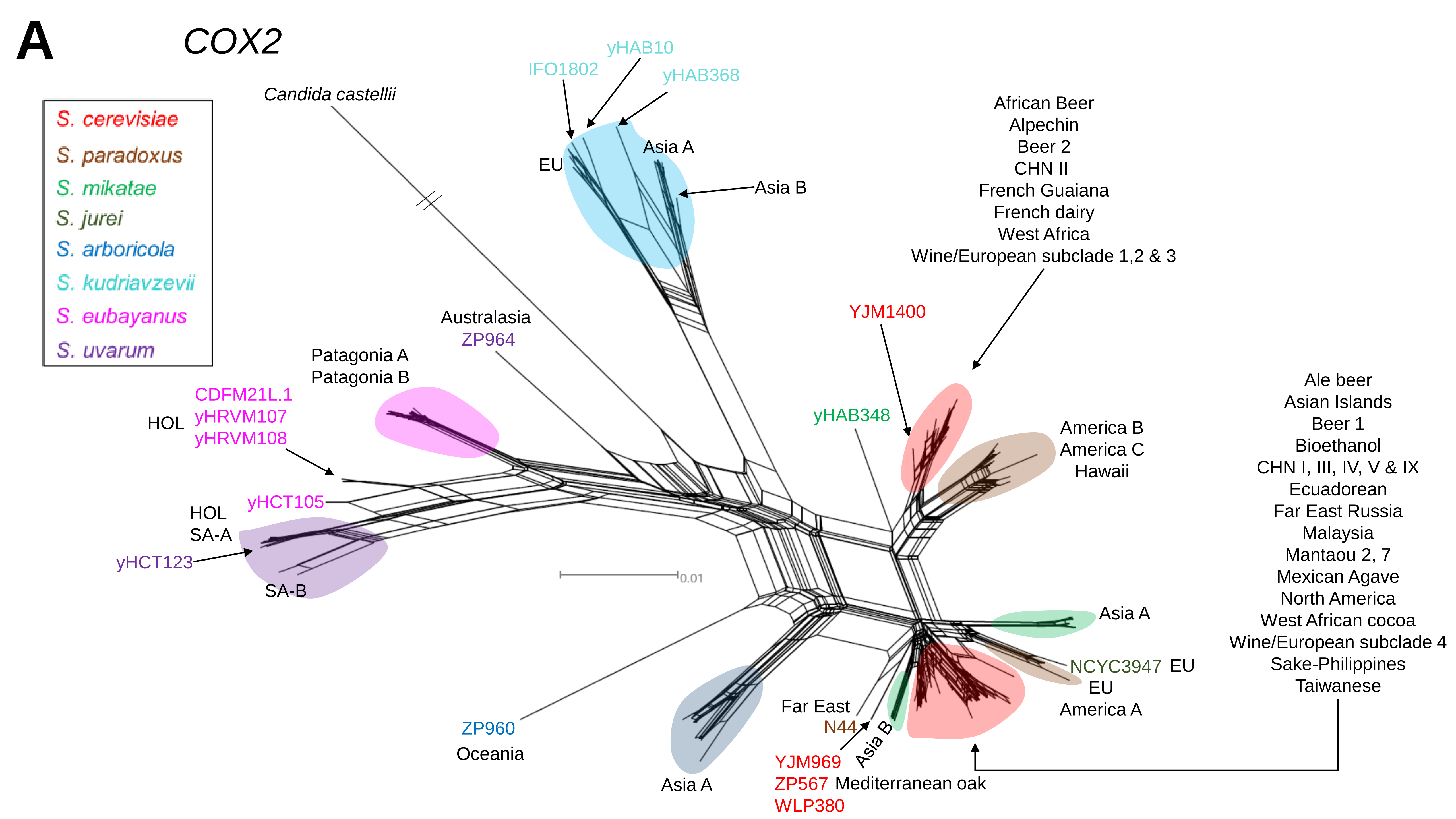

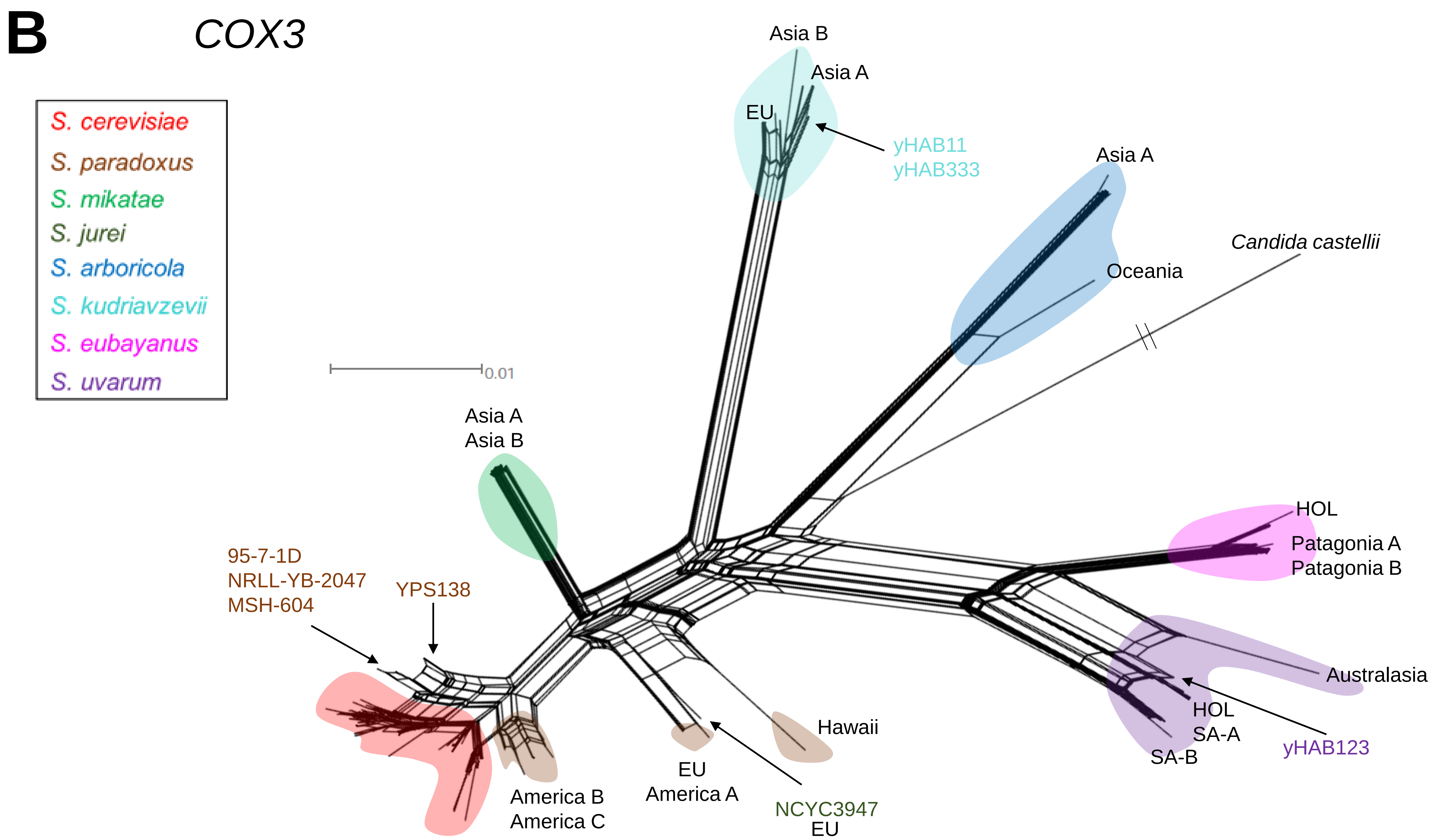

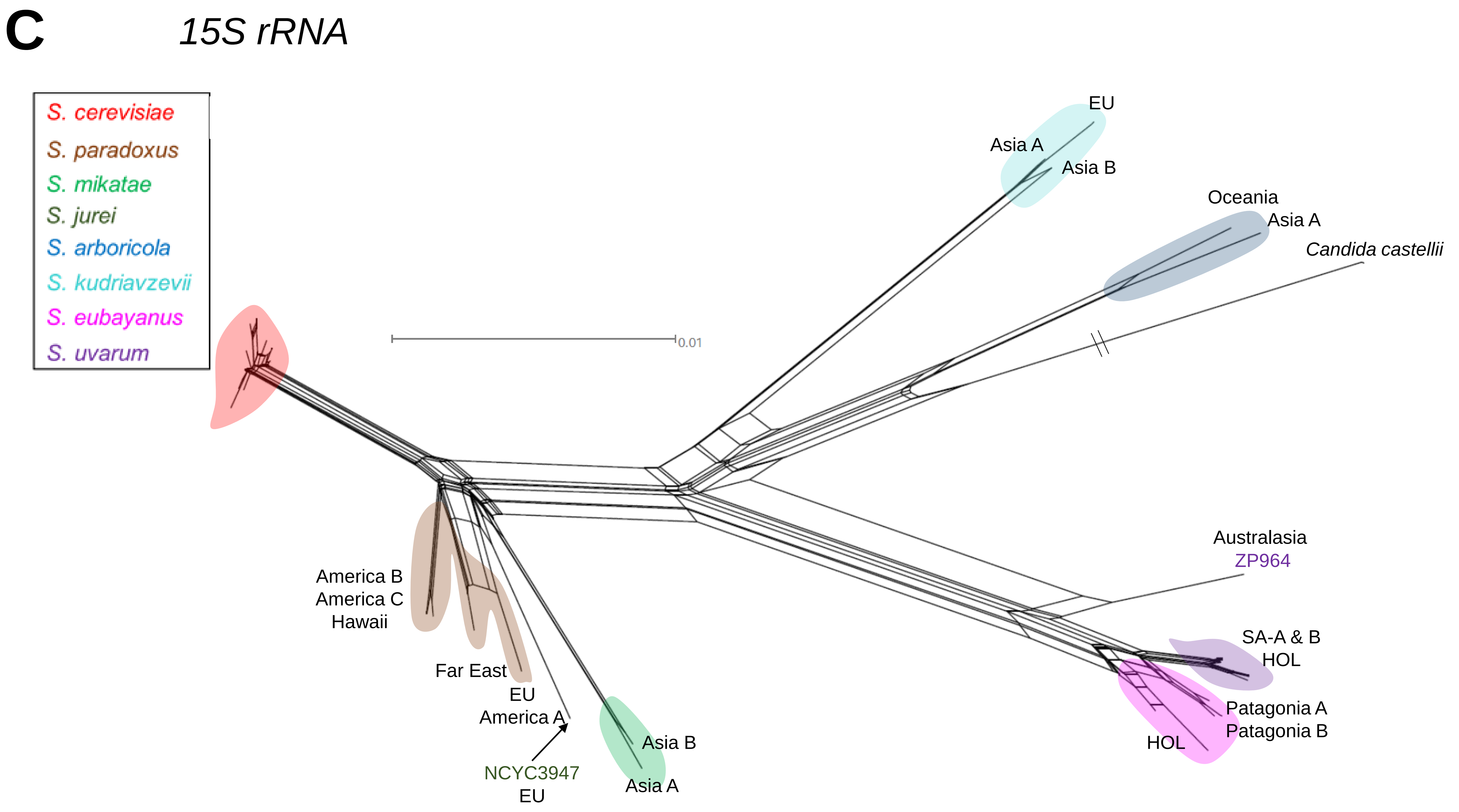

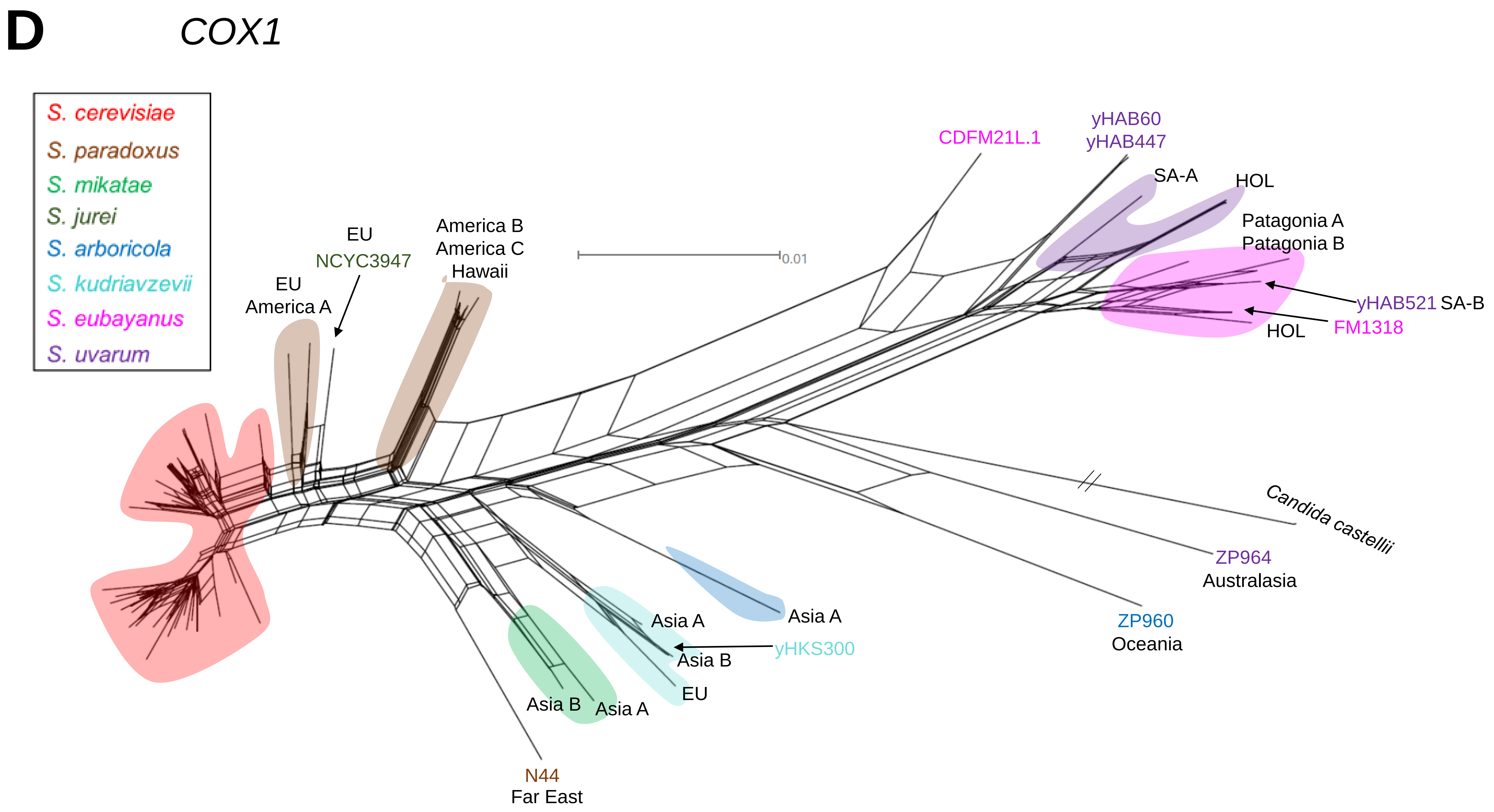

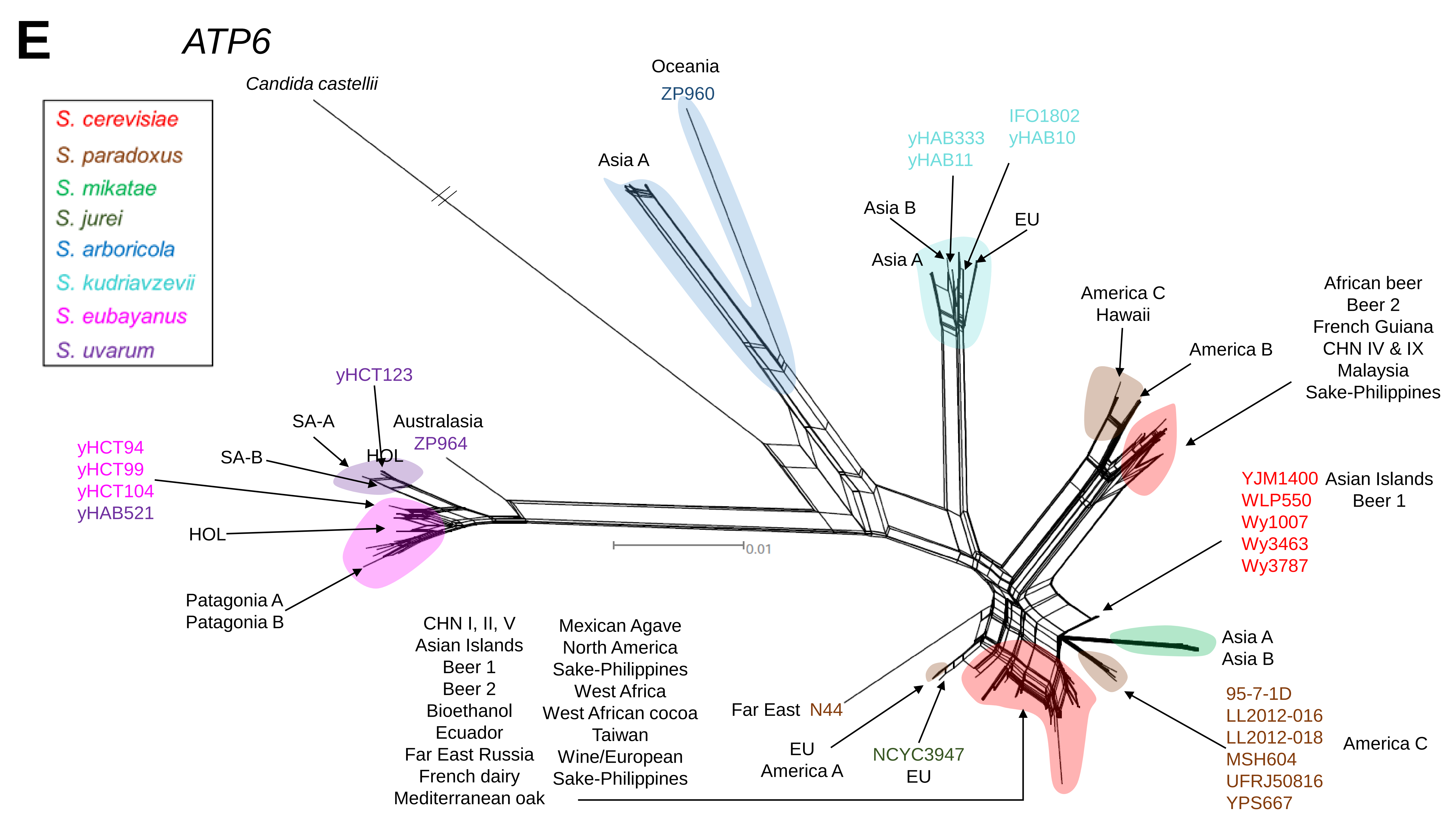

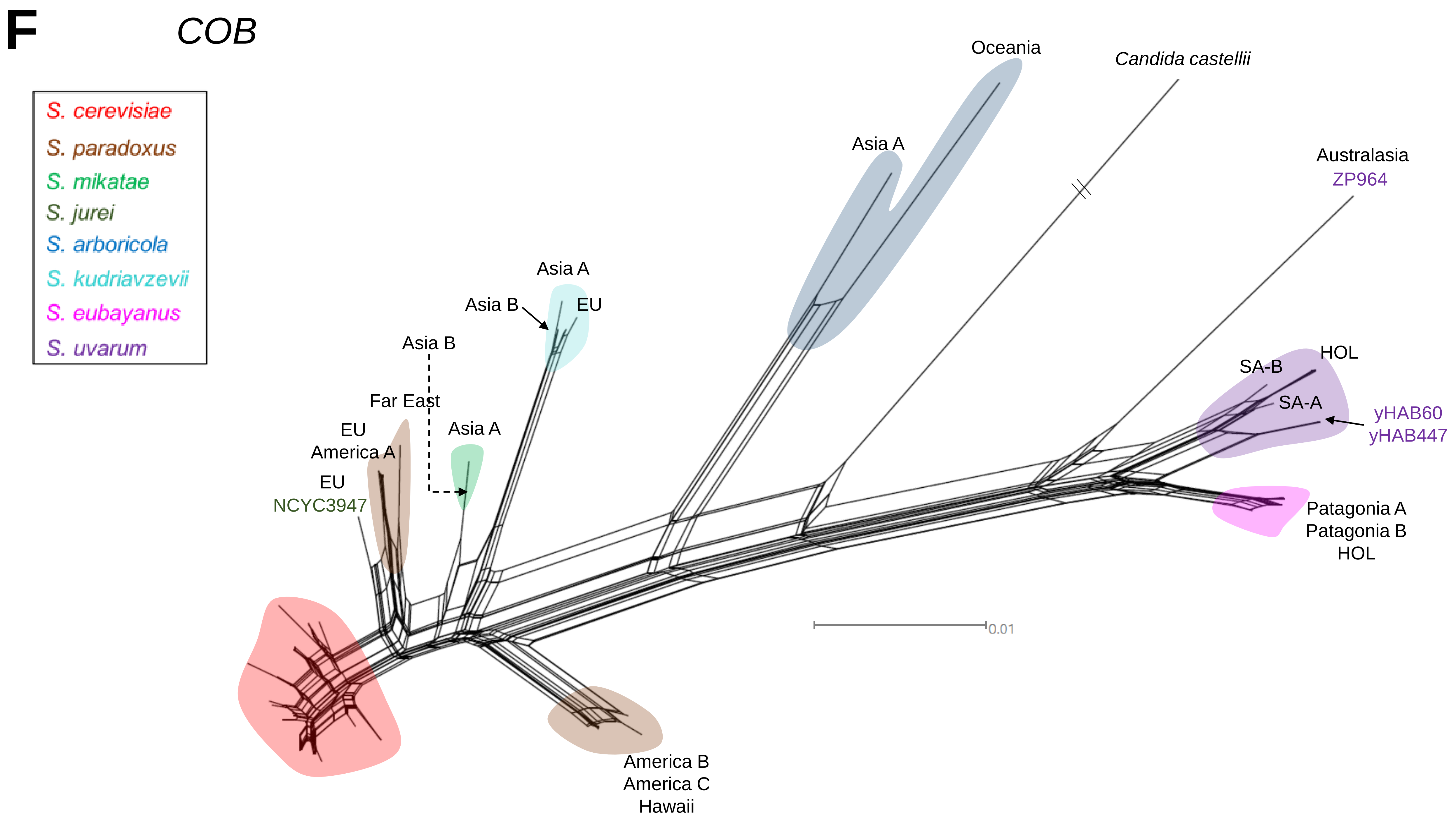

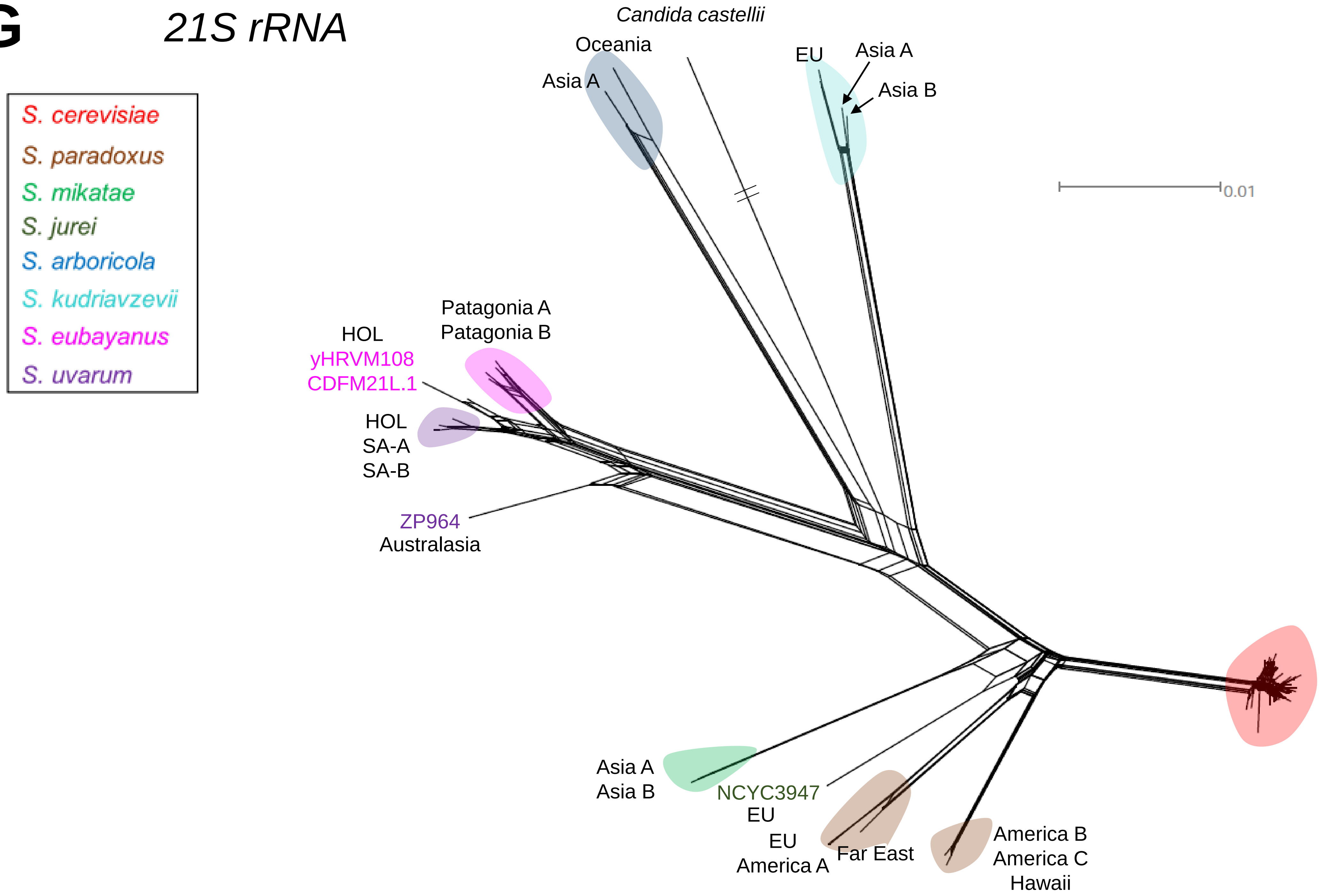

### FigureS5.pdf

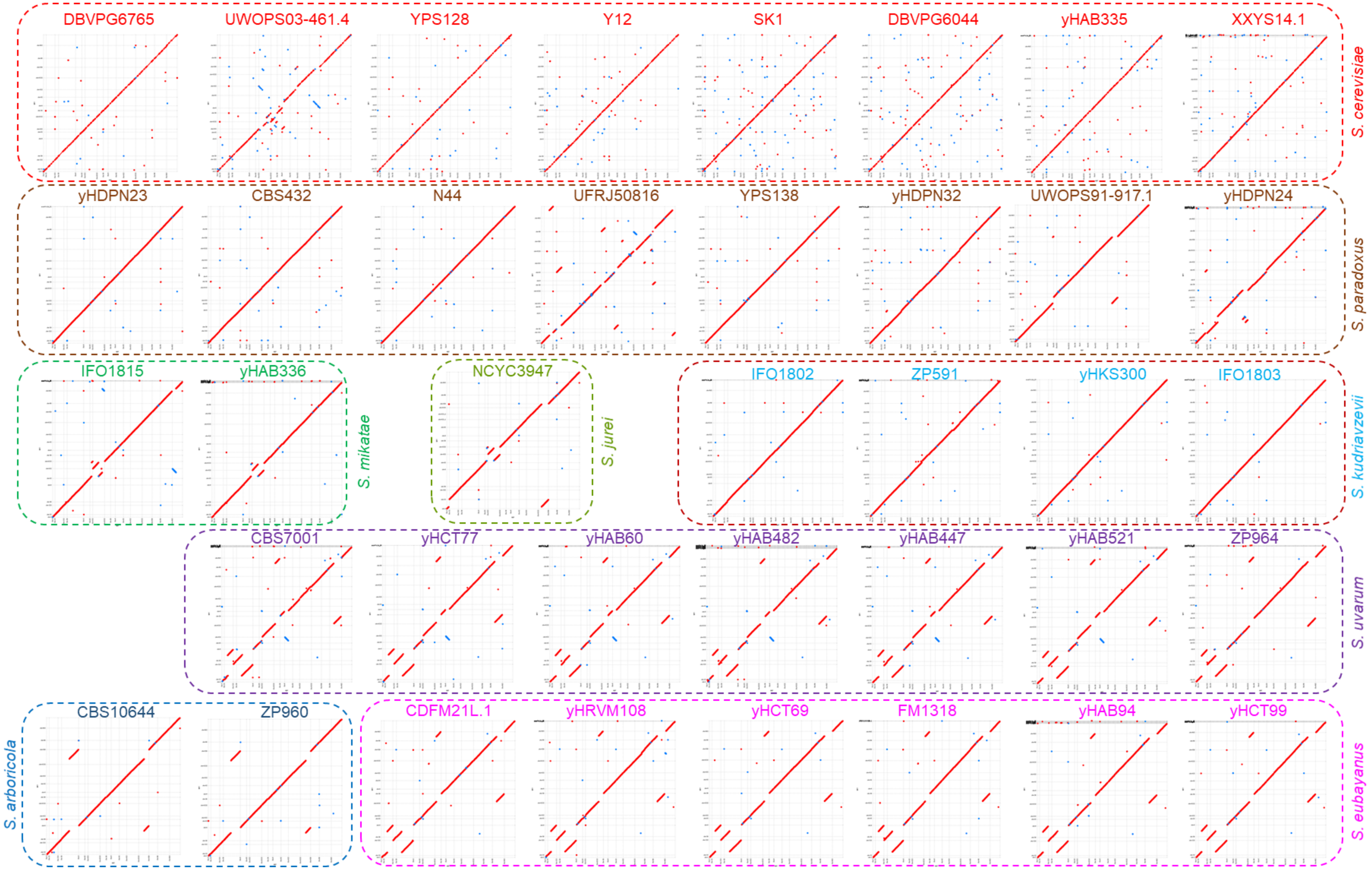

### FigureS6.pdf

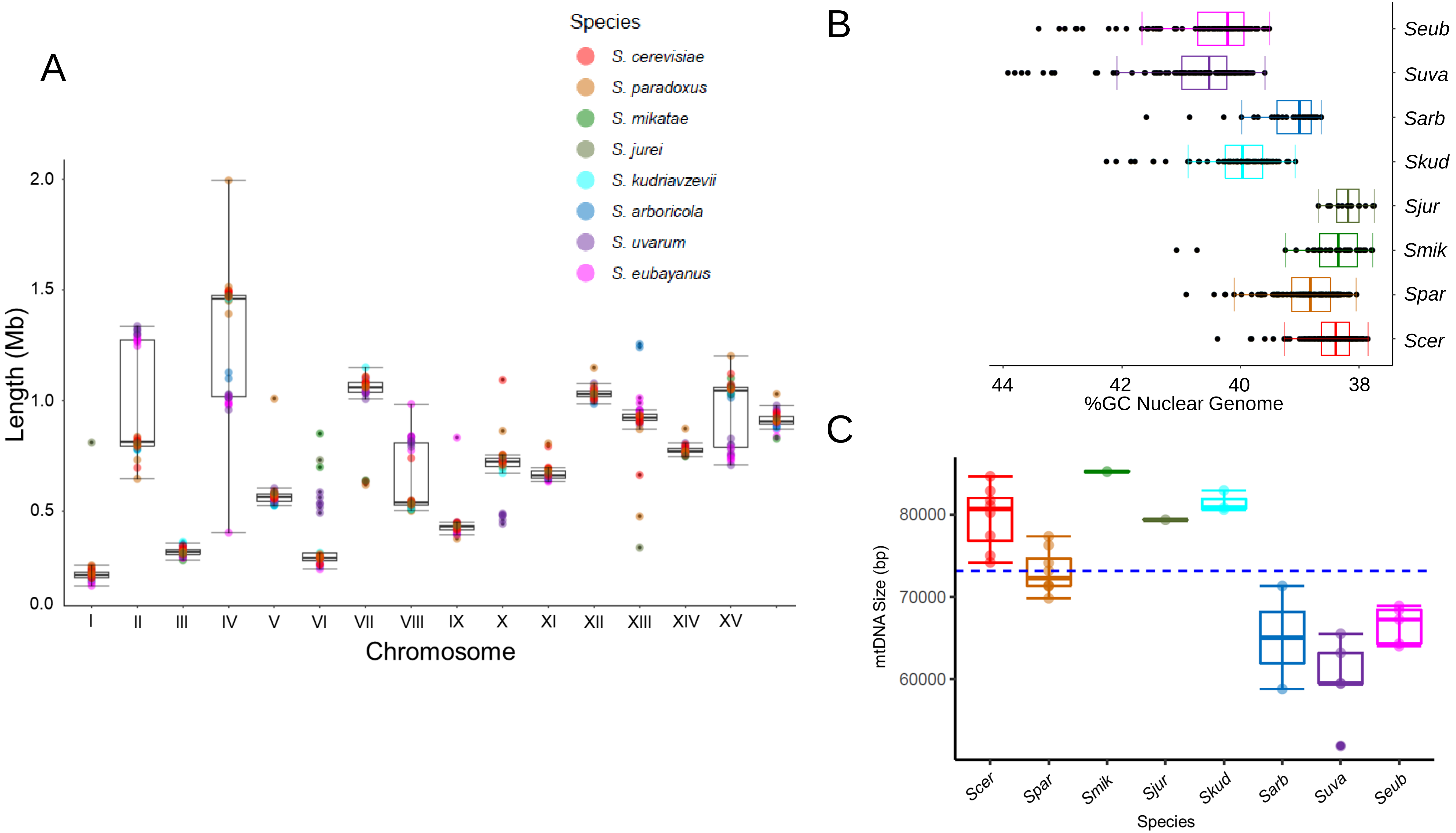

### FigureS7.pdf

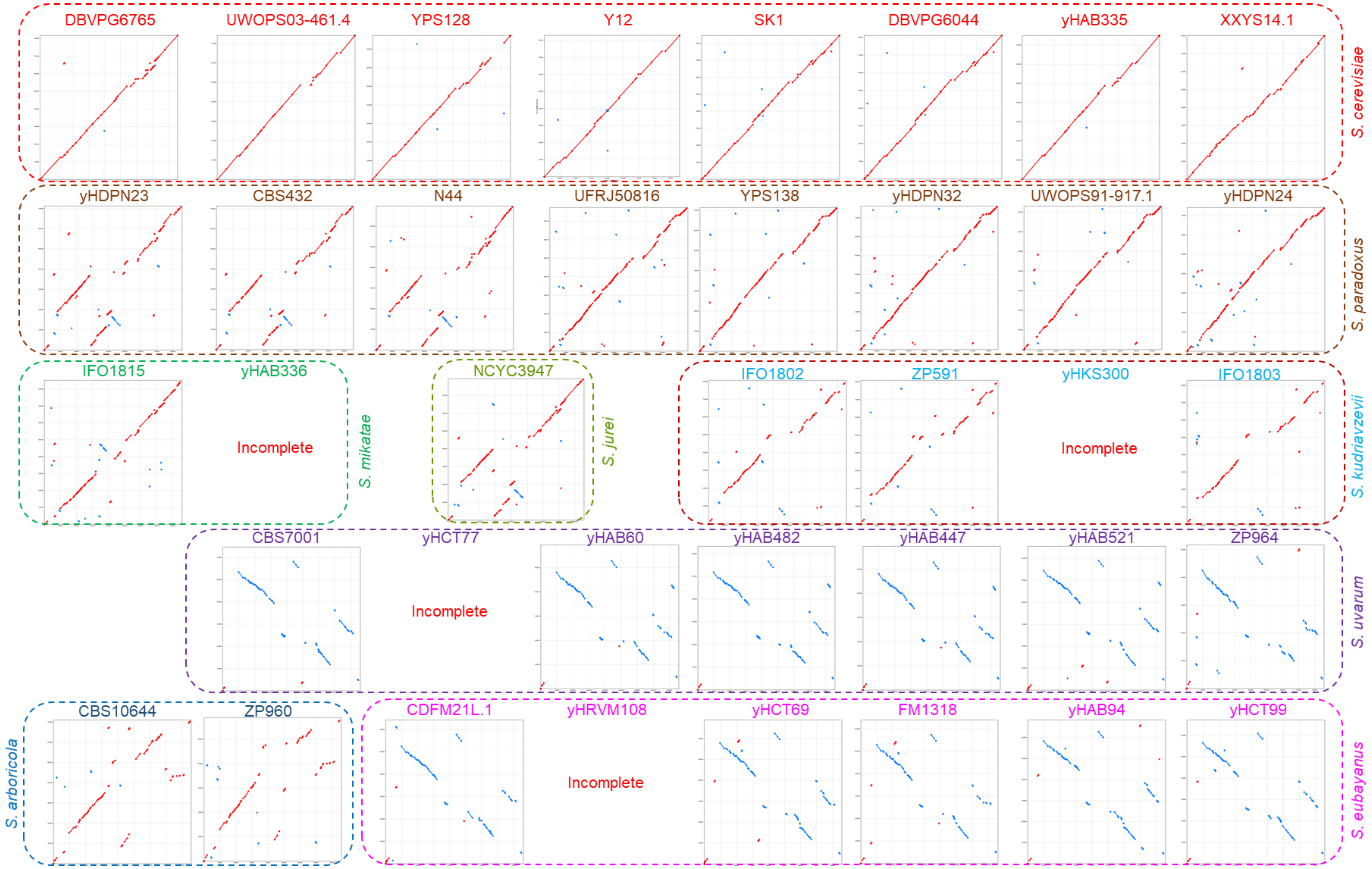

### FigureS8.pdf

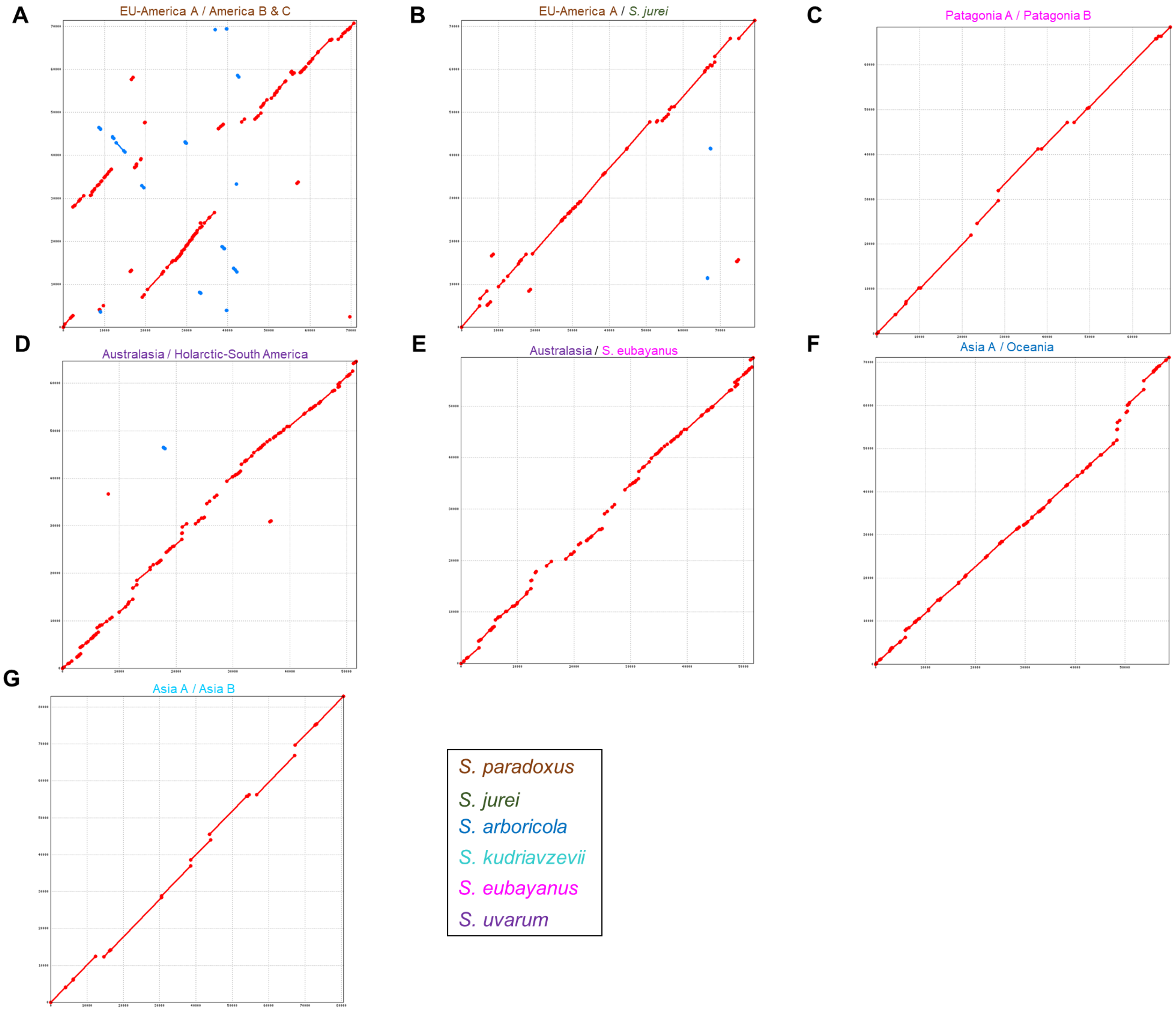

### FigureS9.pdf

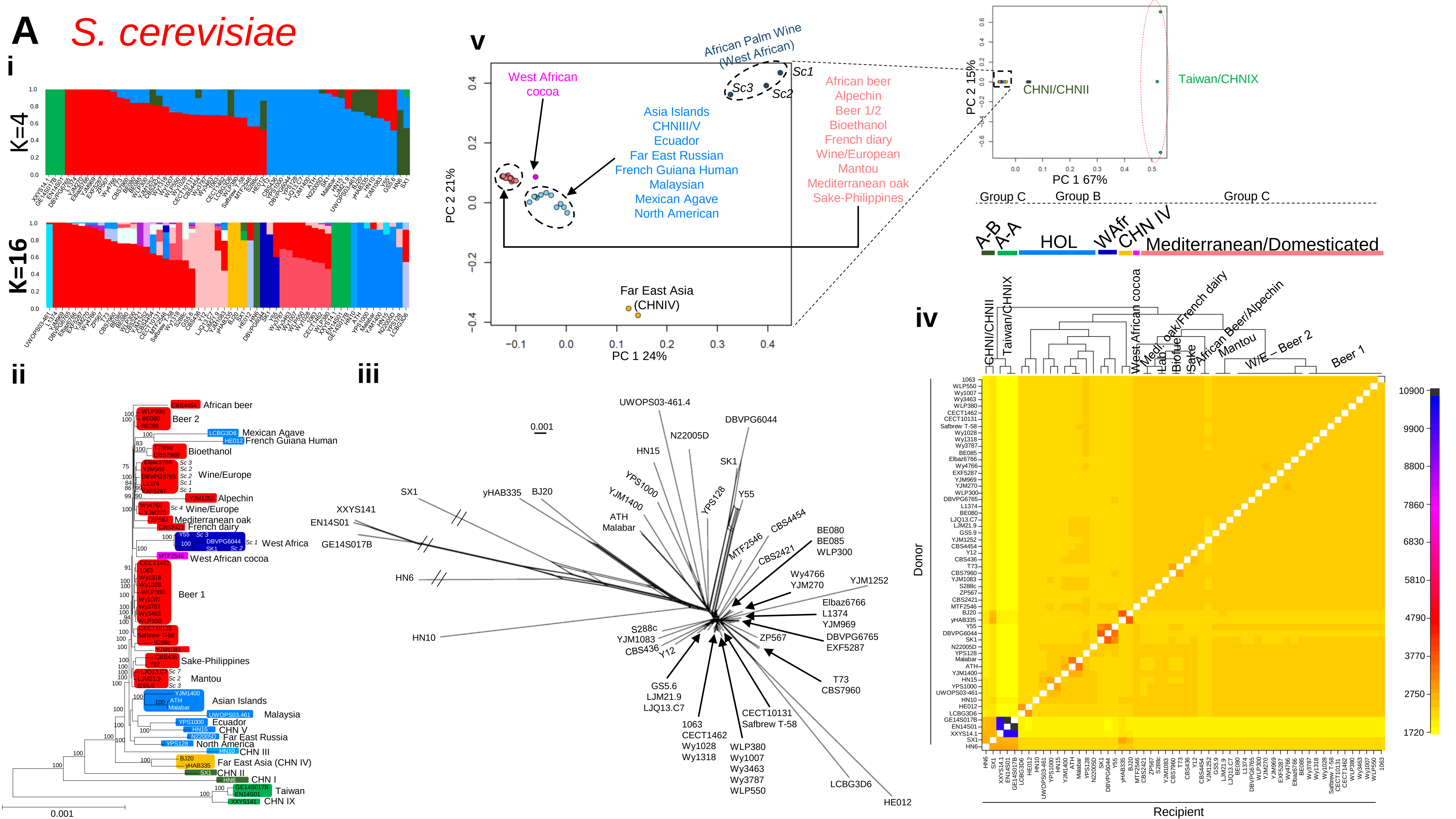

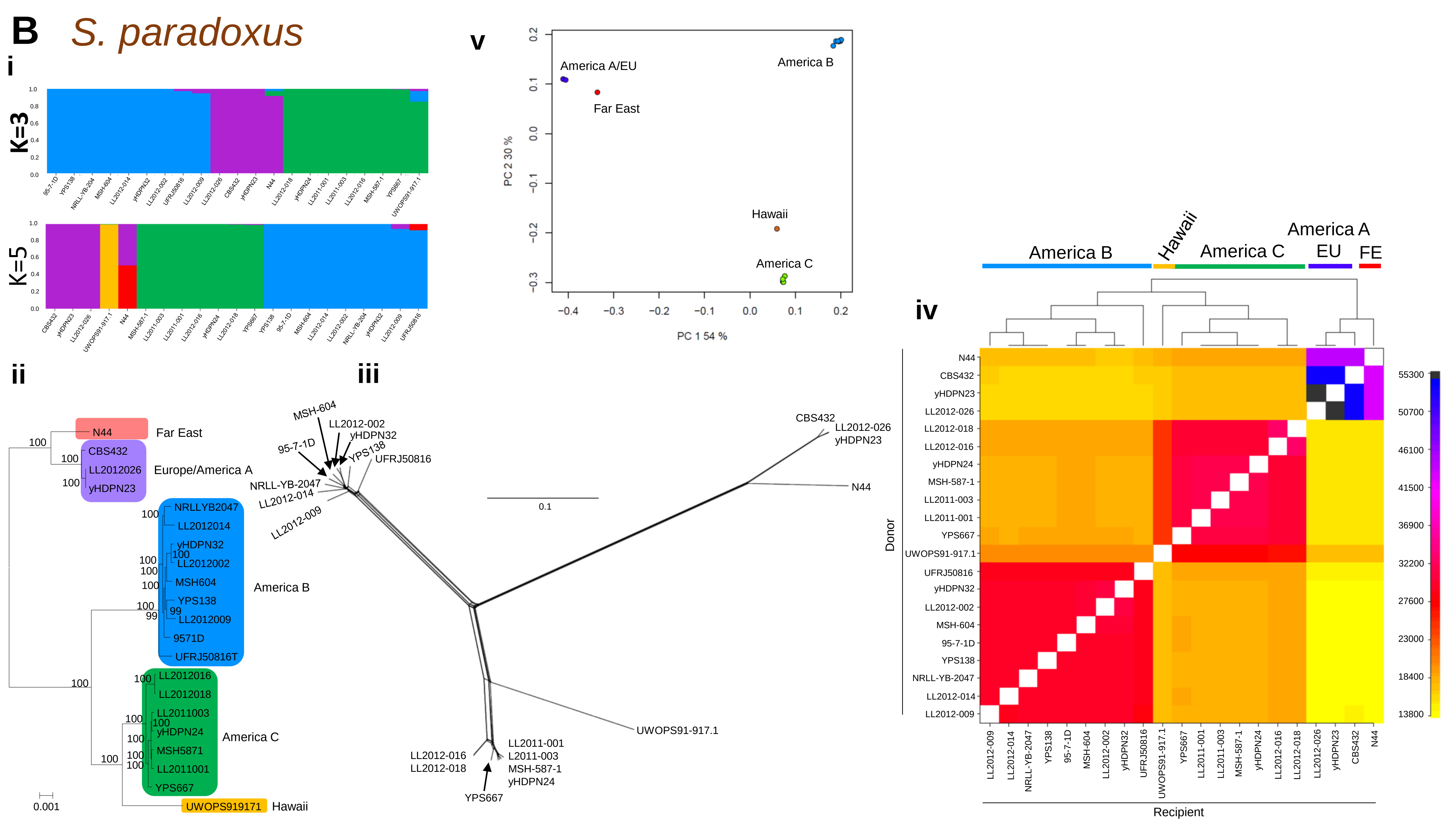

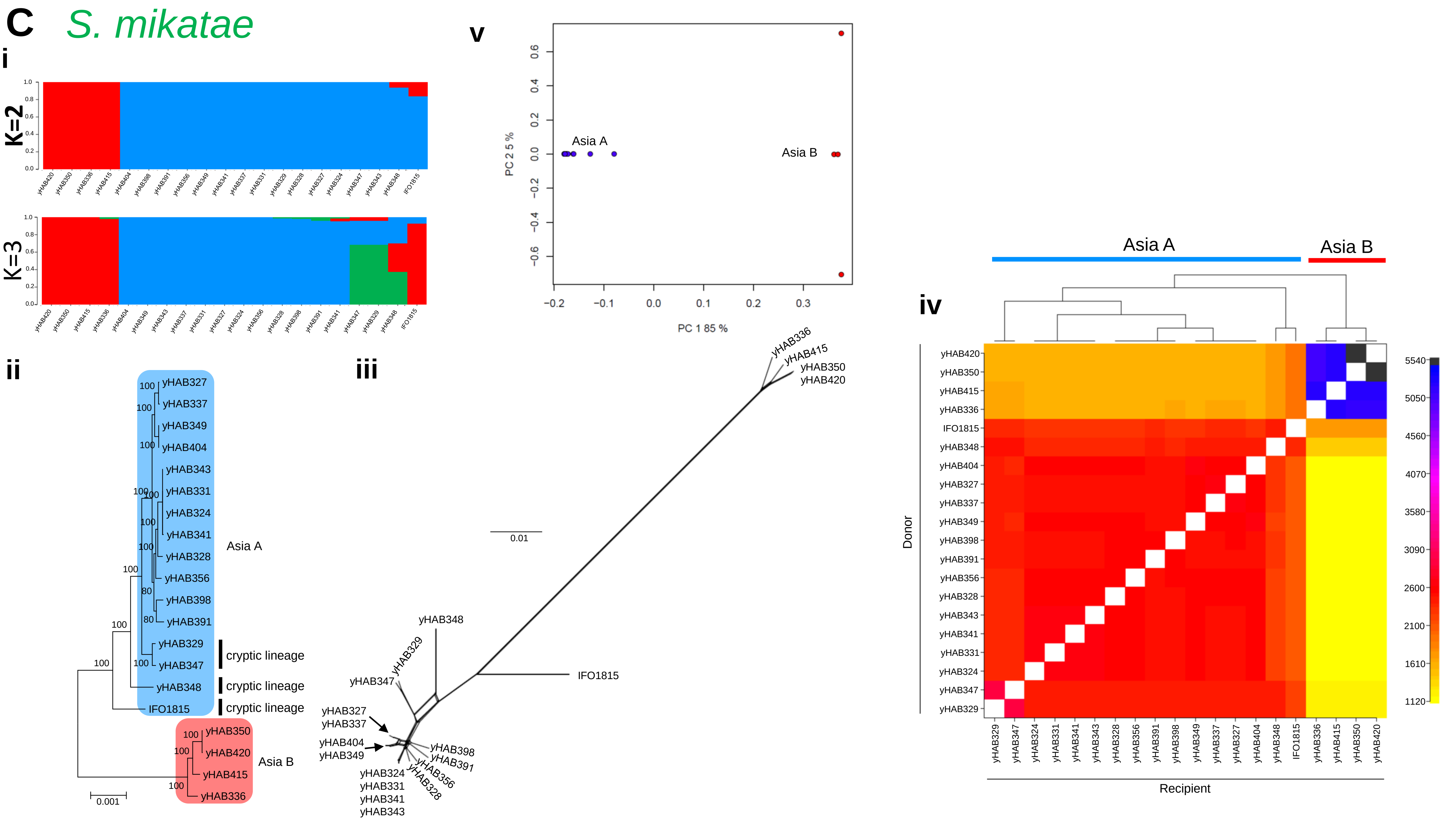

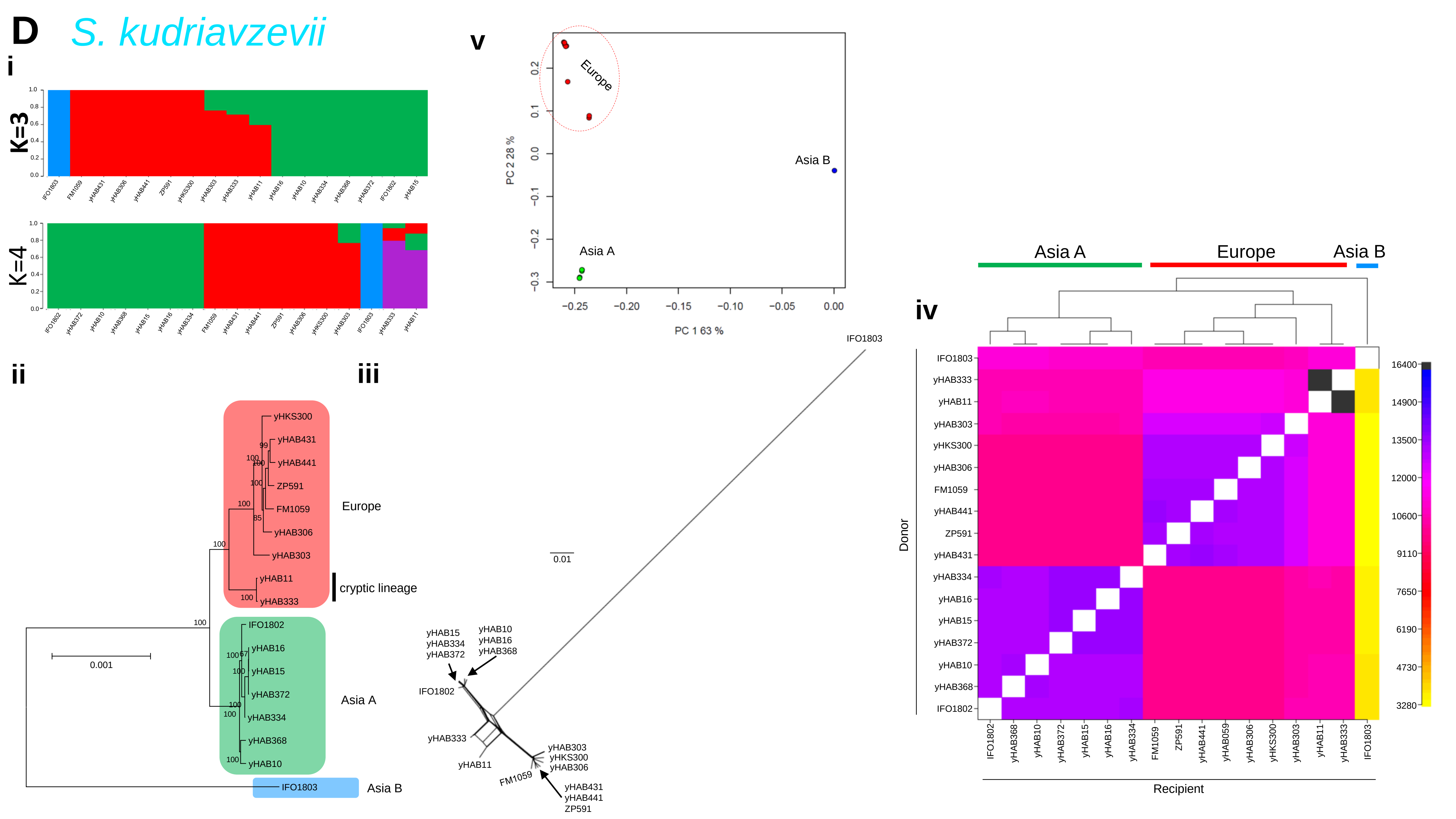

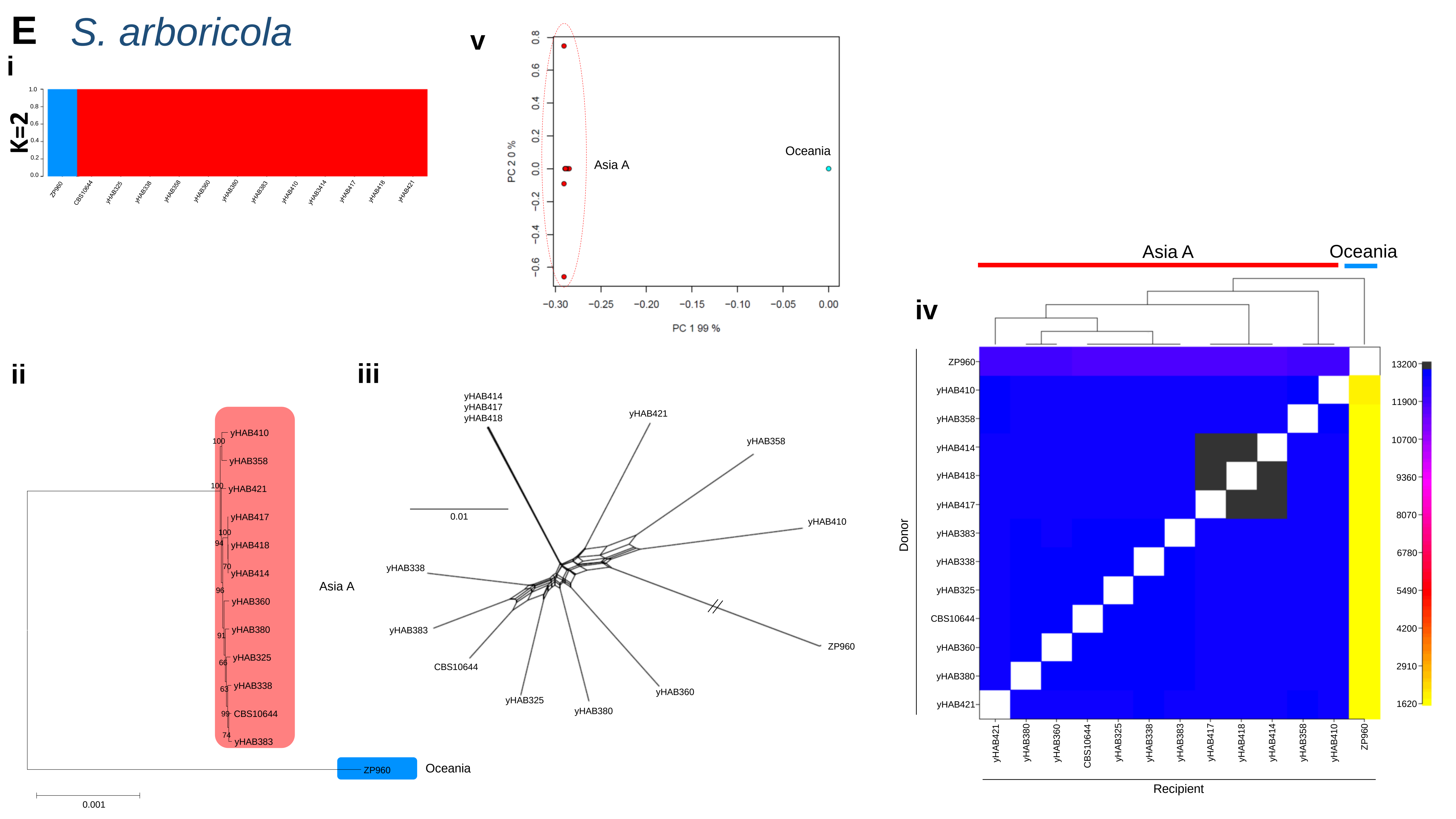

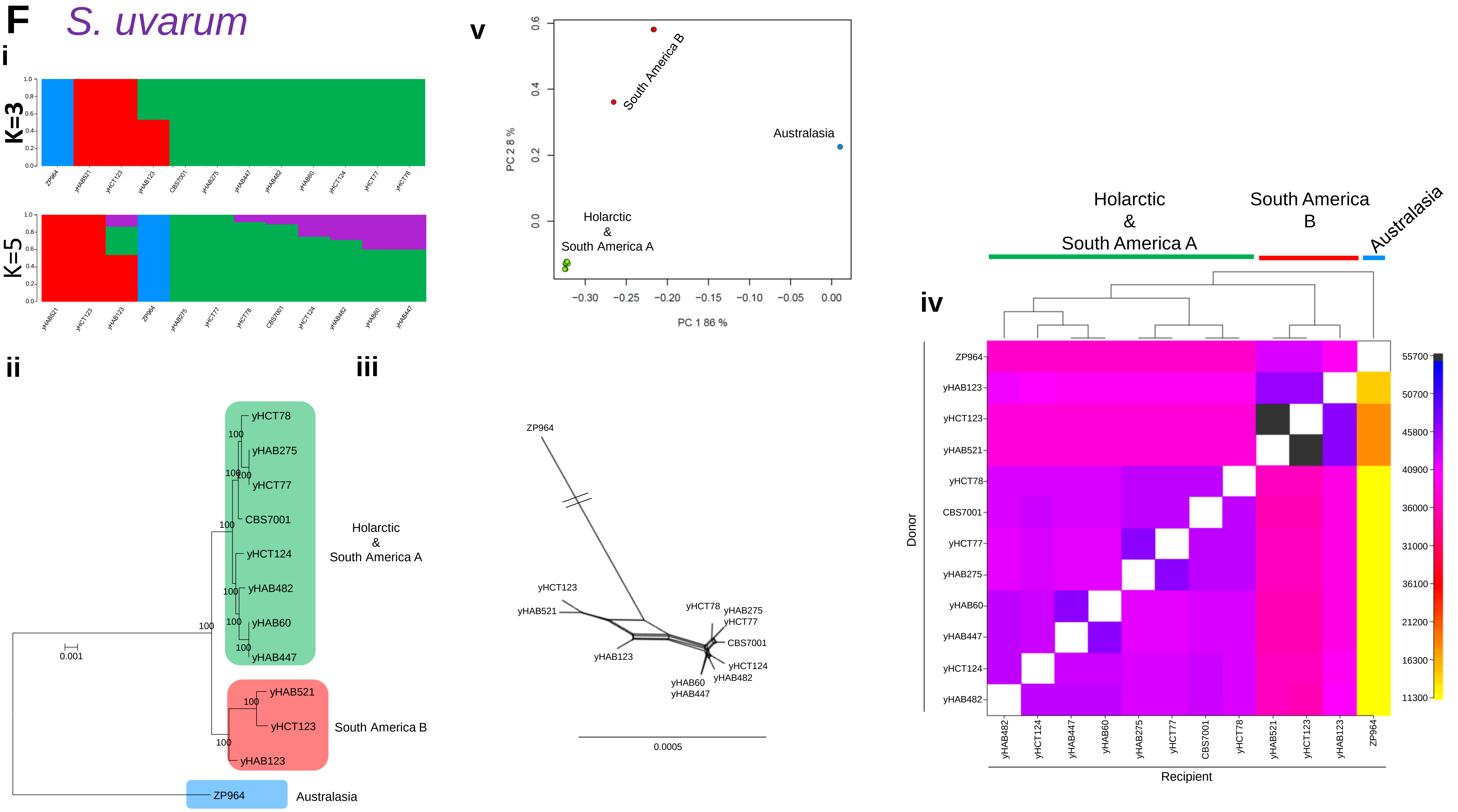

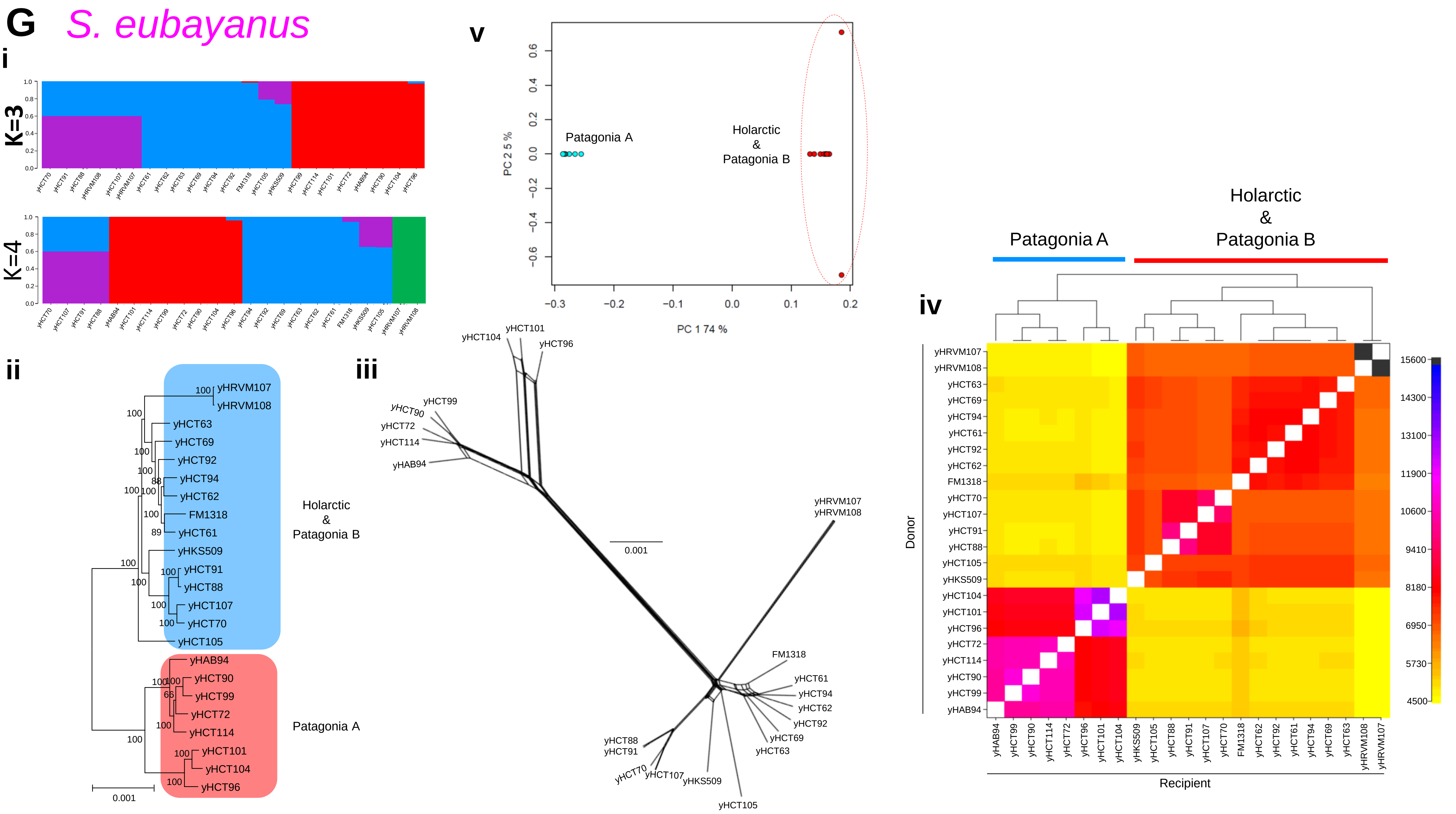

### FigureS11.pdf

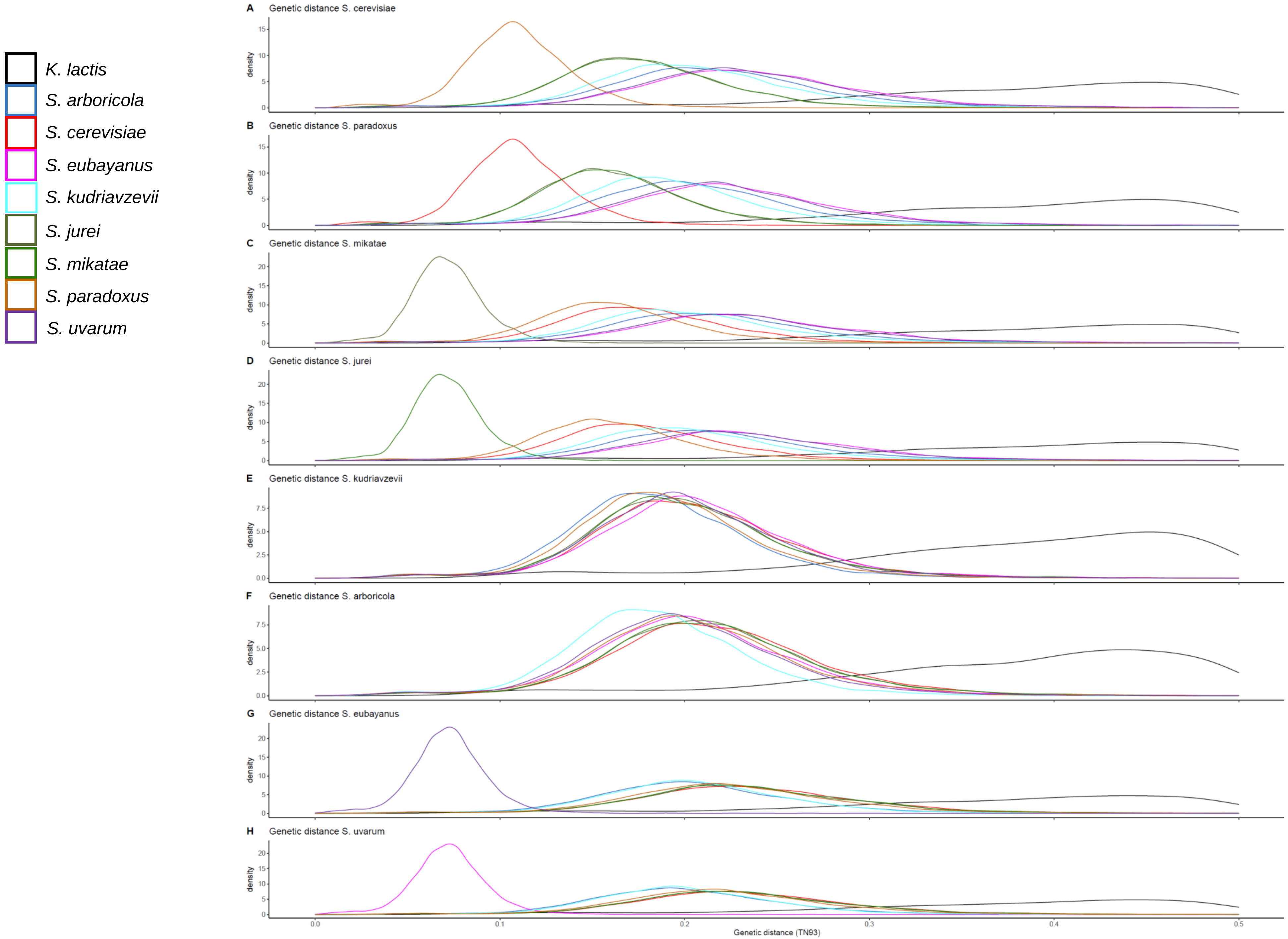

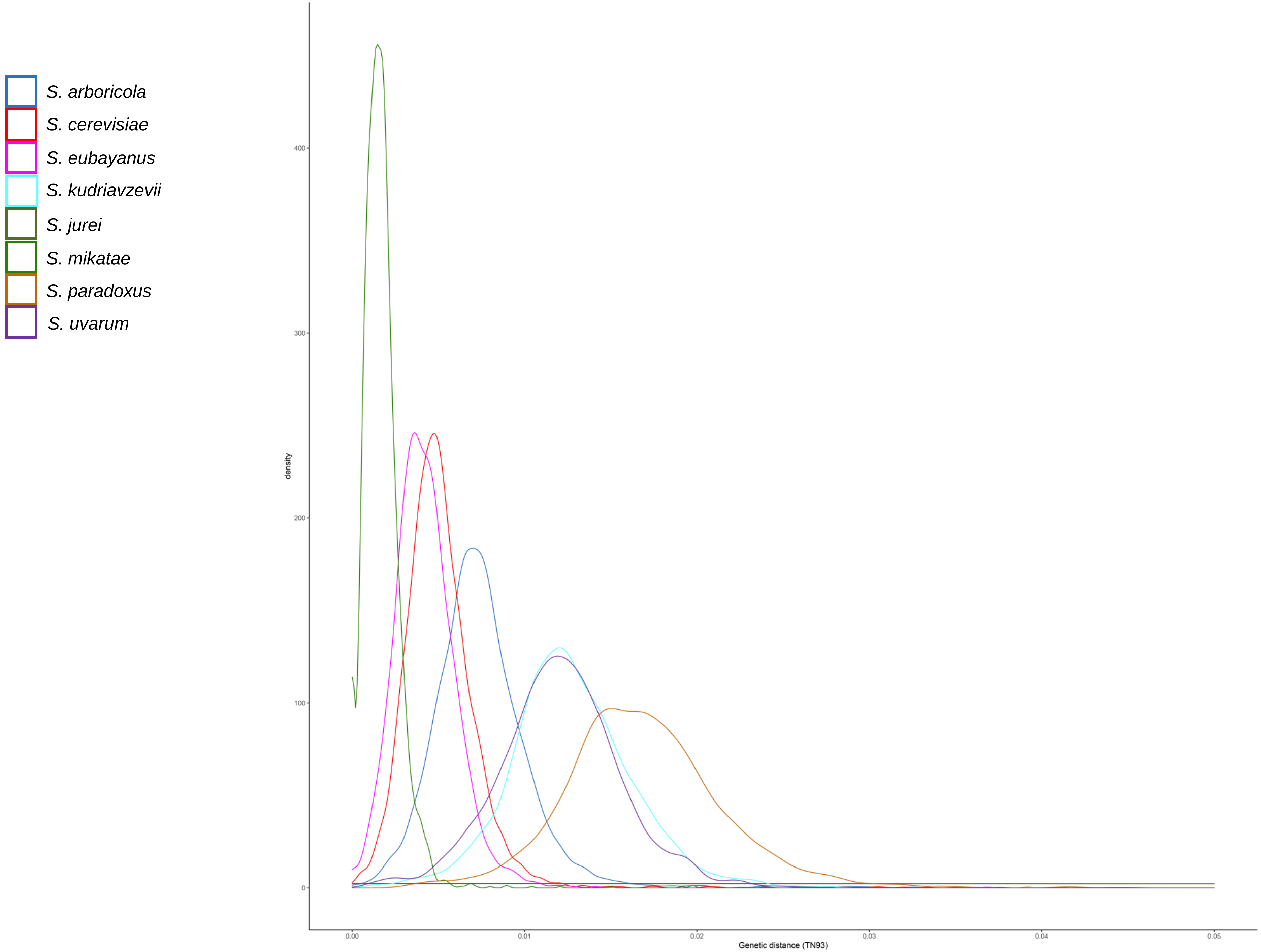

### FigureS12.pdf

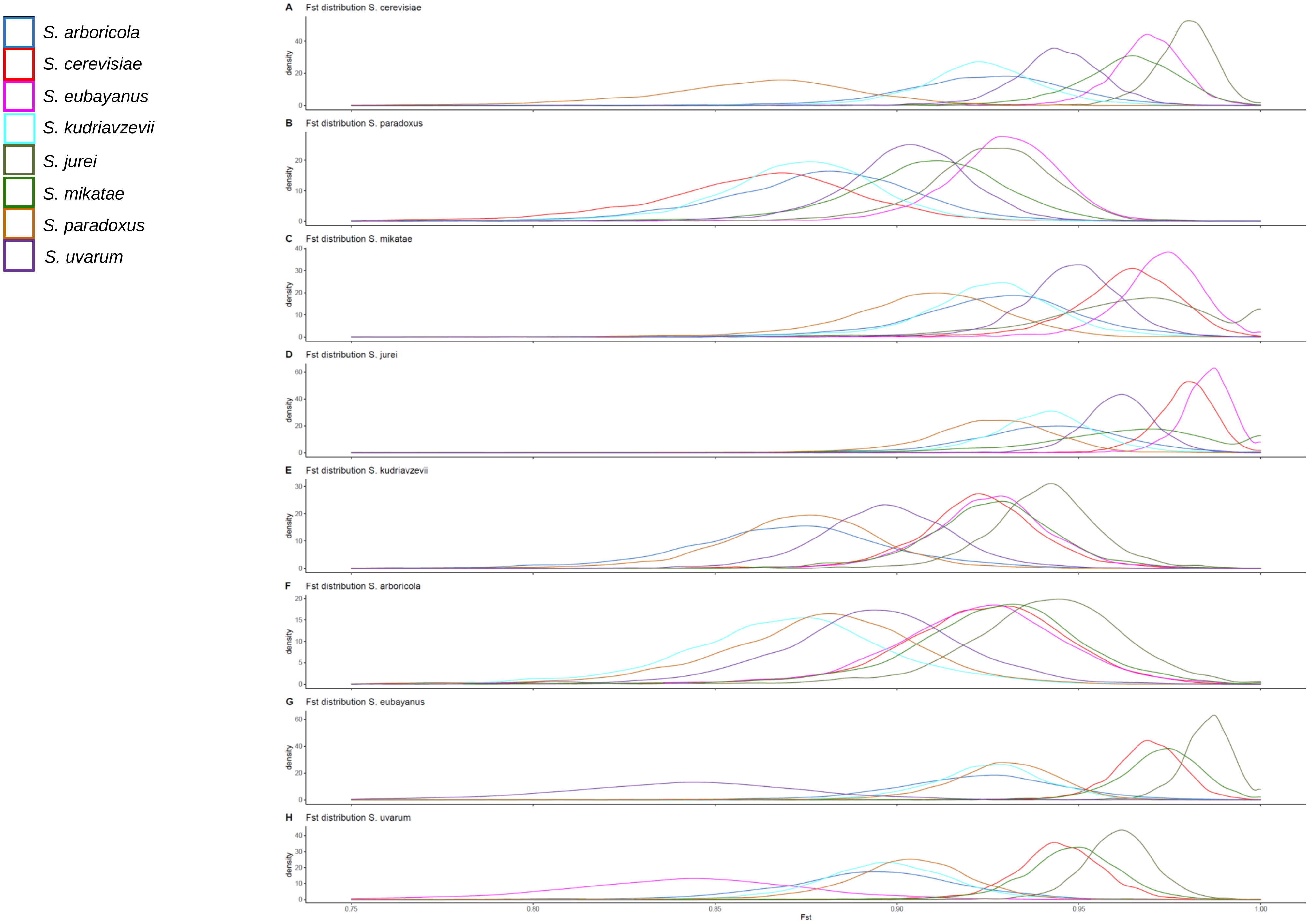

### FigureS13.pdf

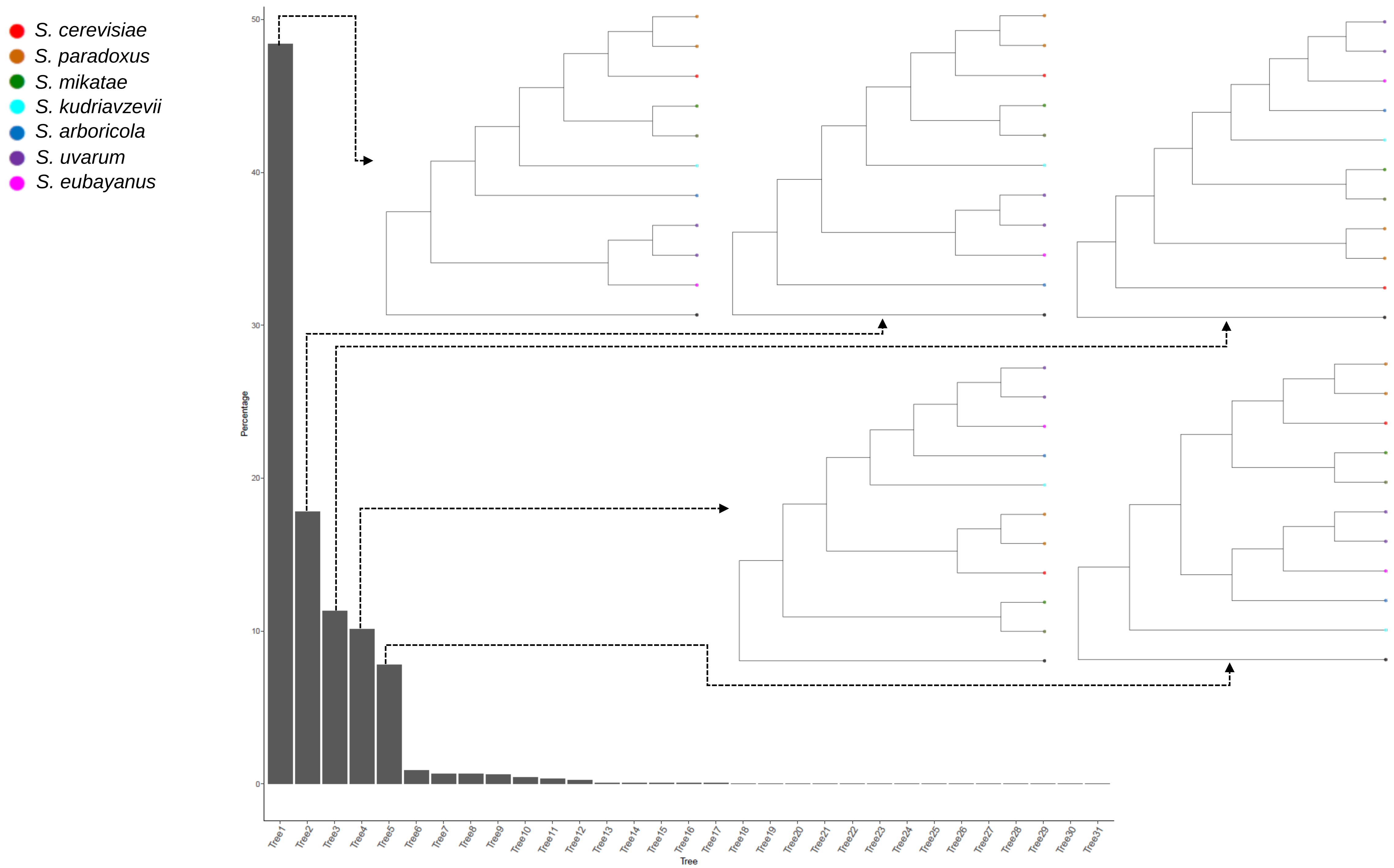

### FigureS14.pdf

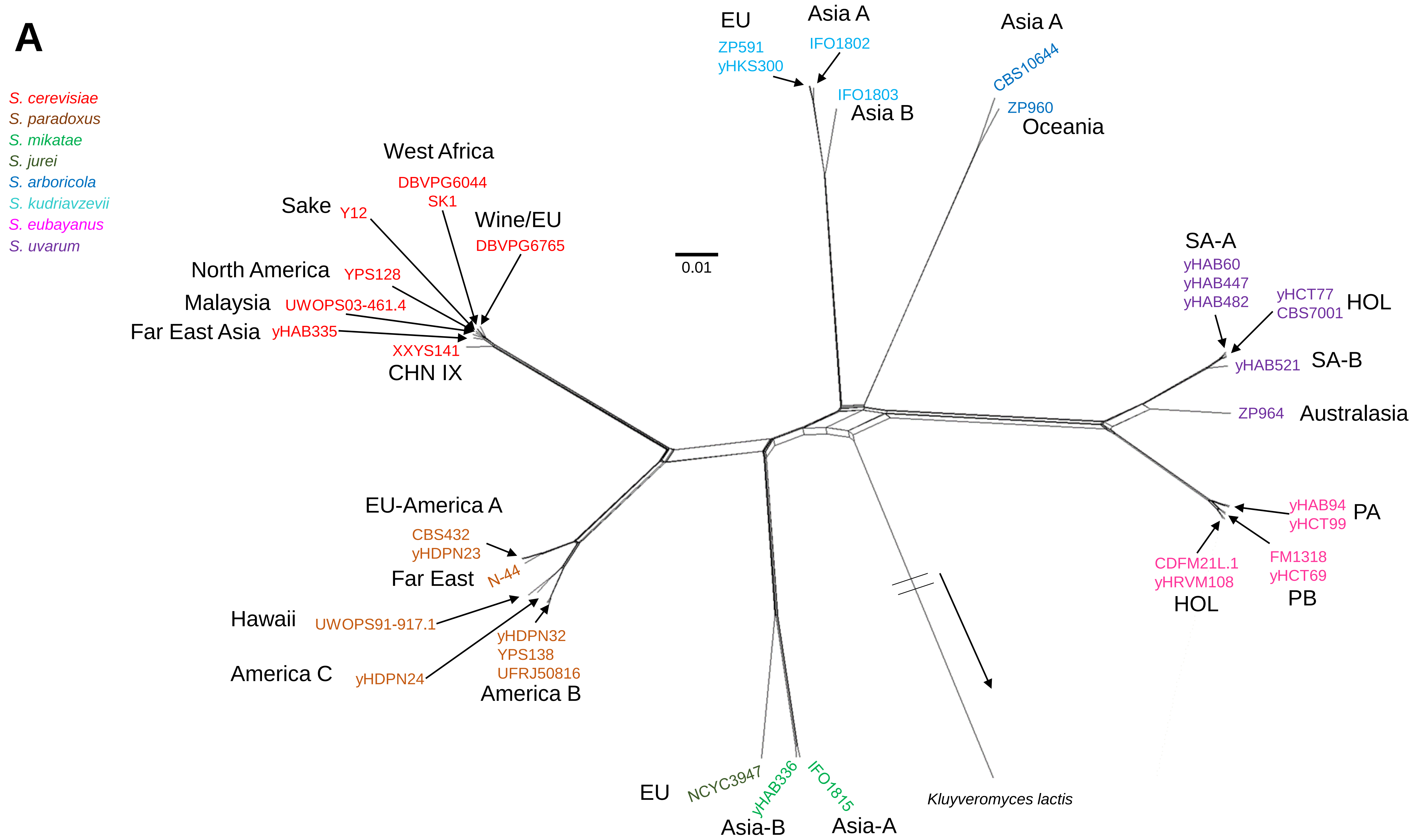

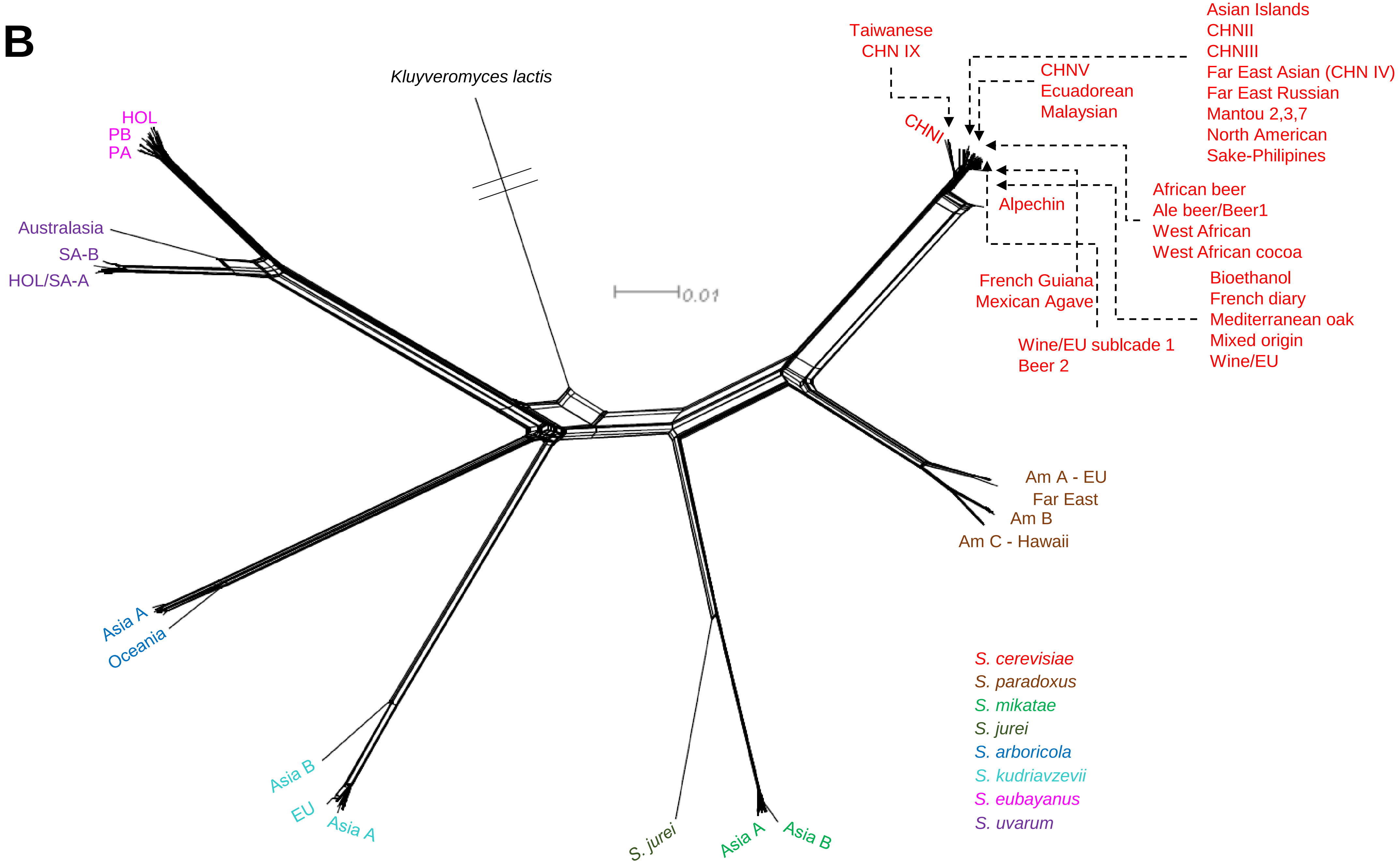

### FigureS15.pdf

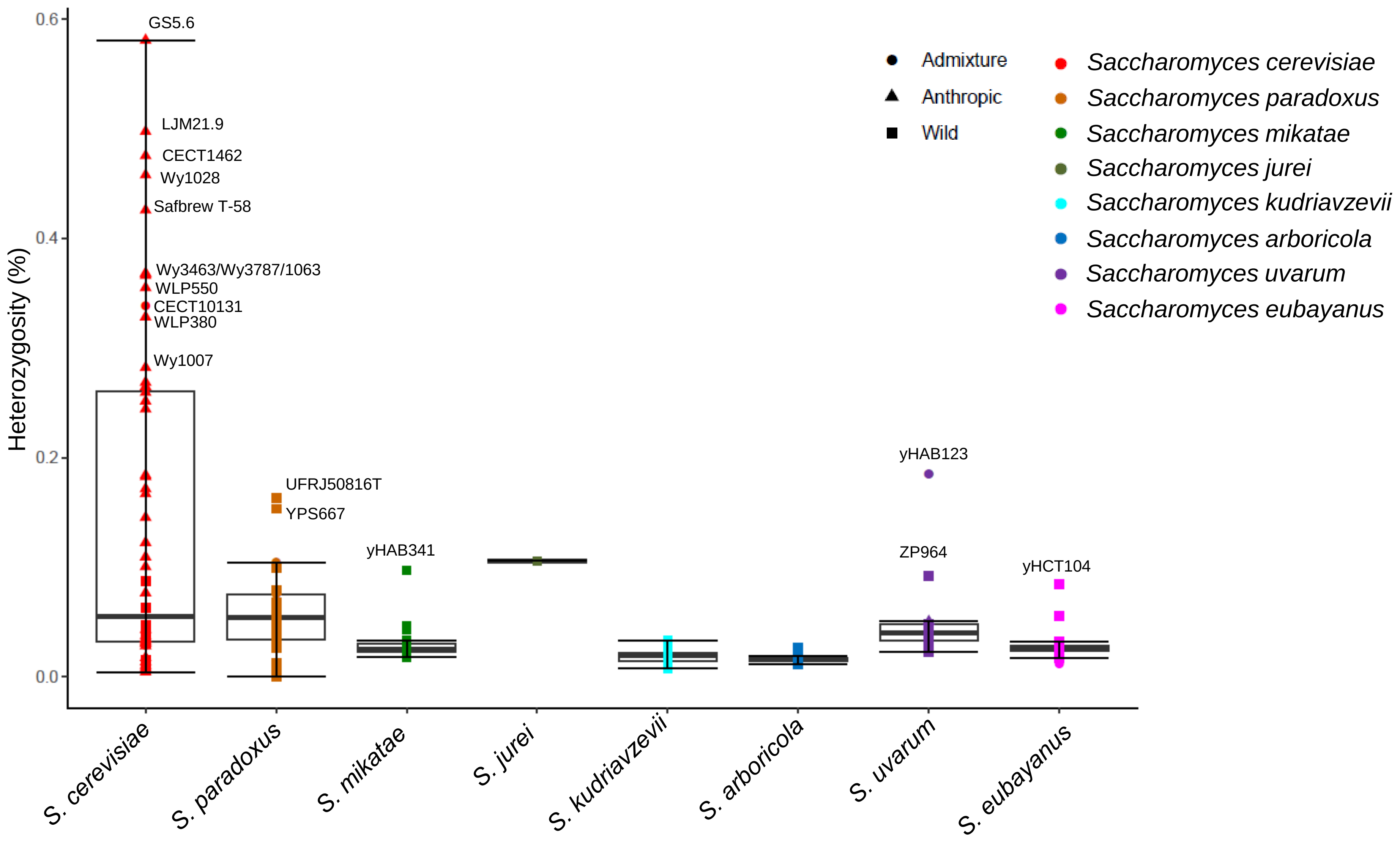

### FigureS16.pdf

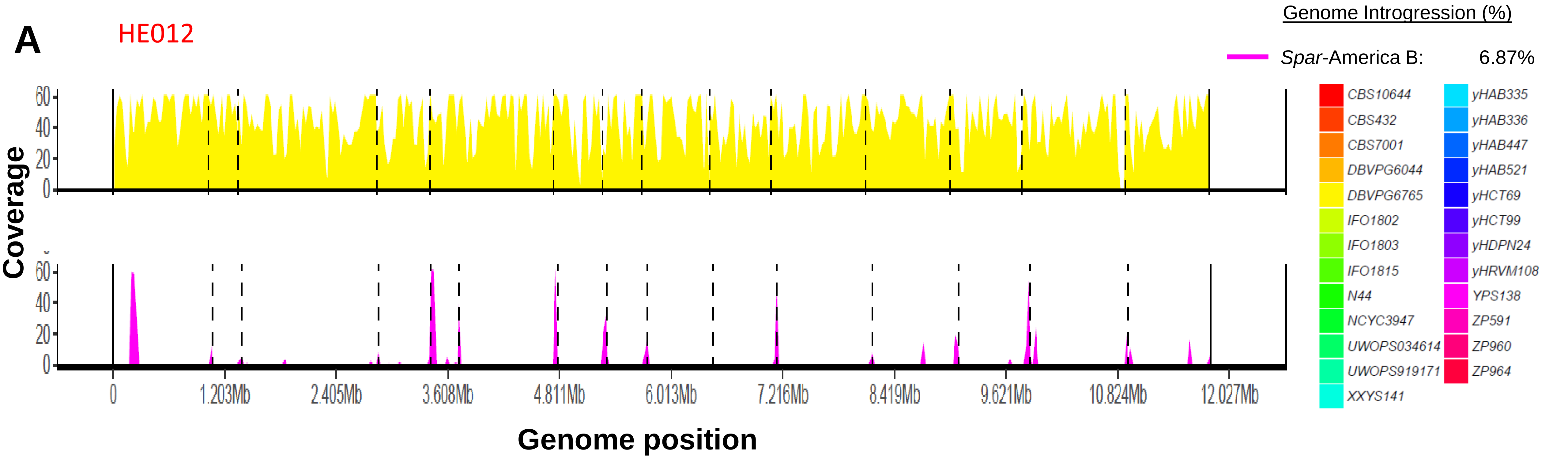

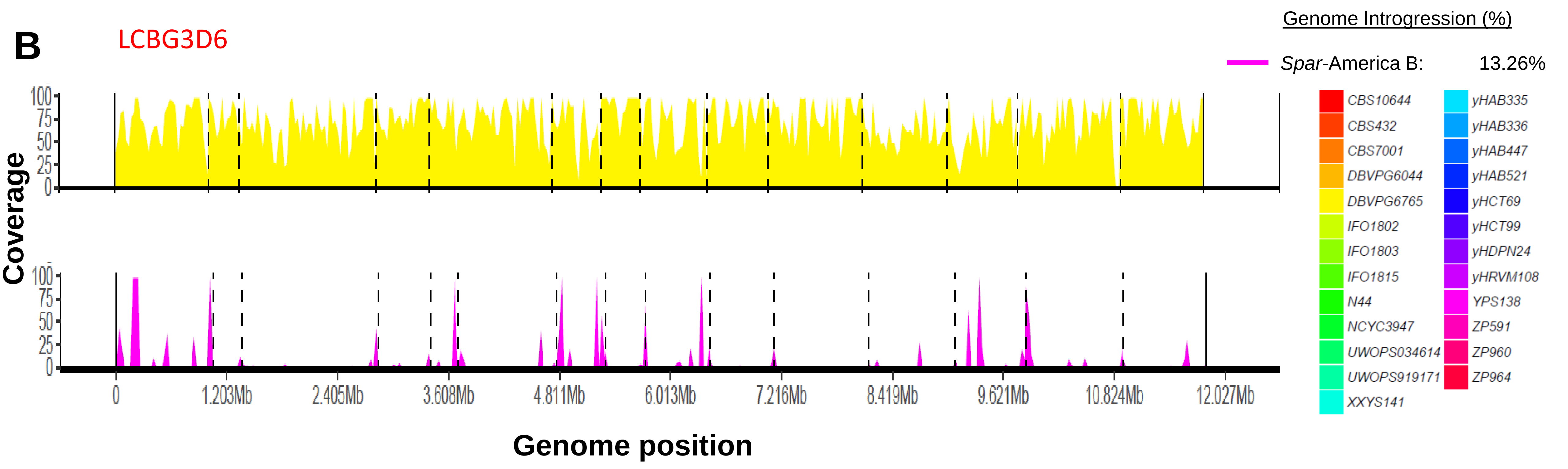

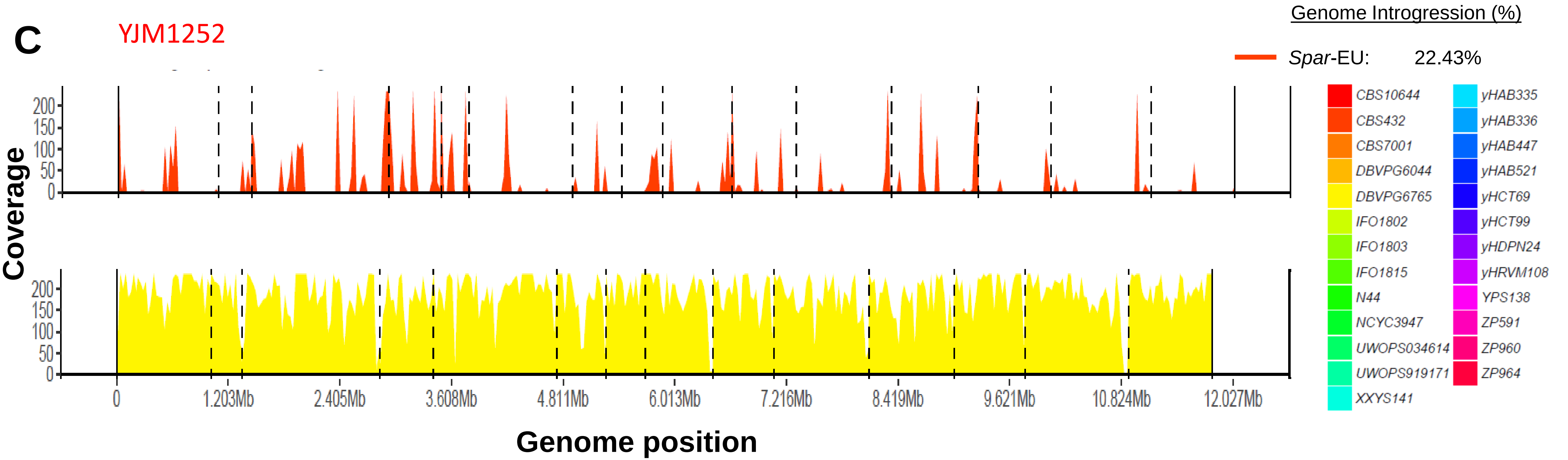

### FigureS17.pdf

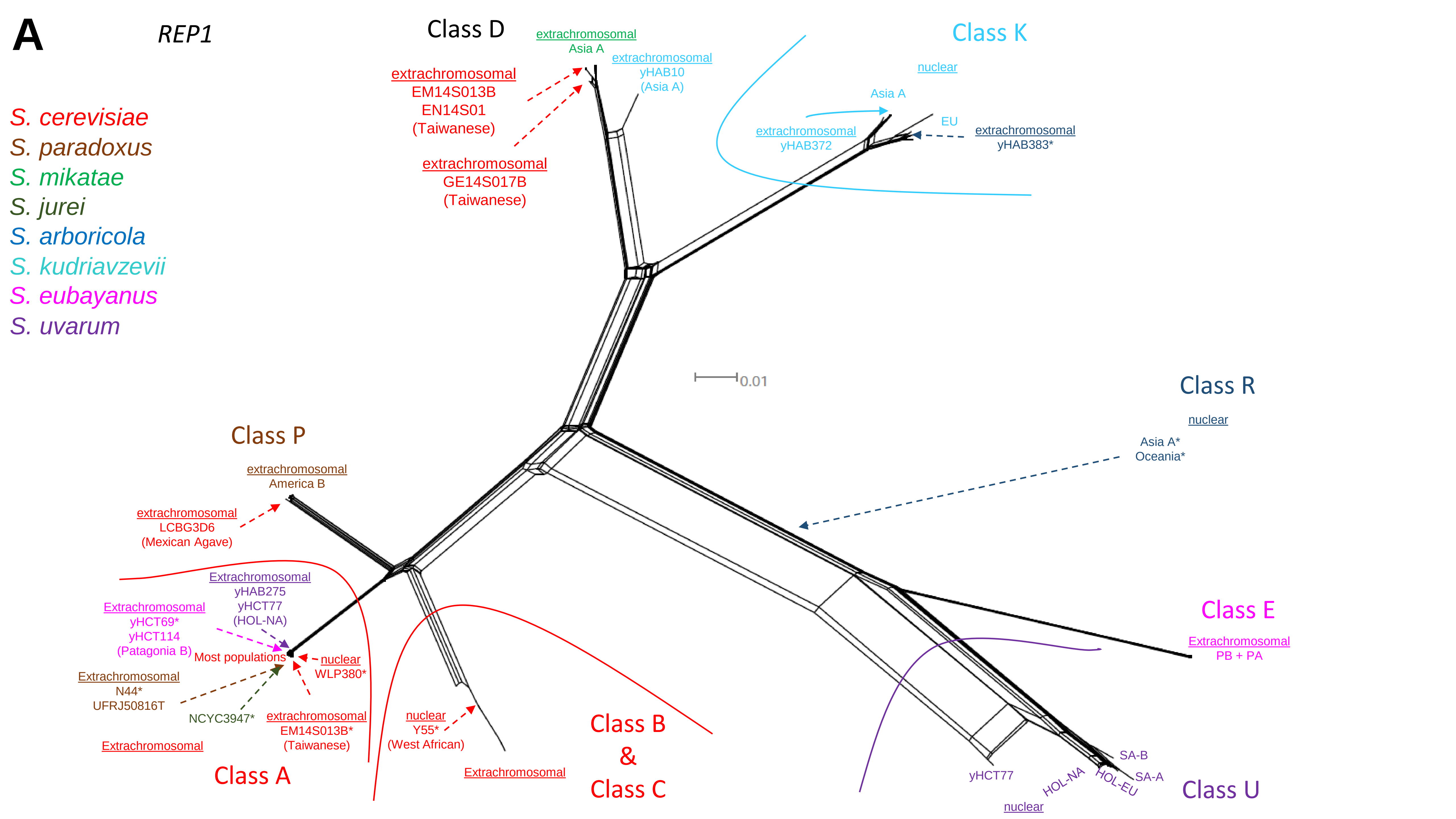

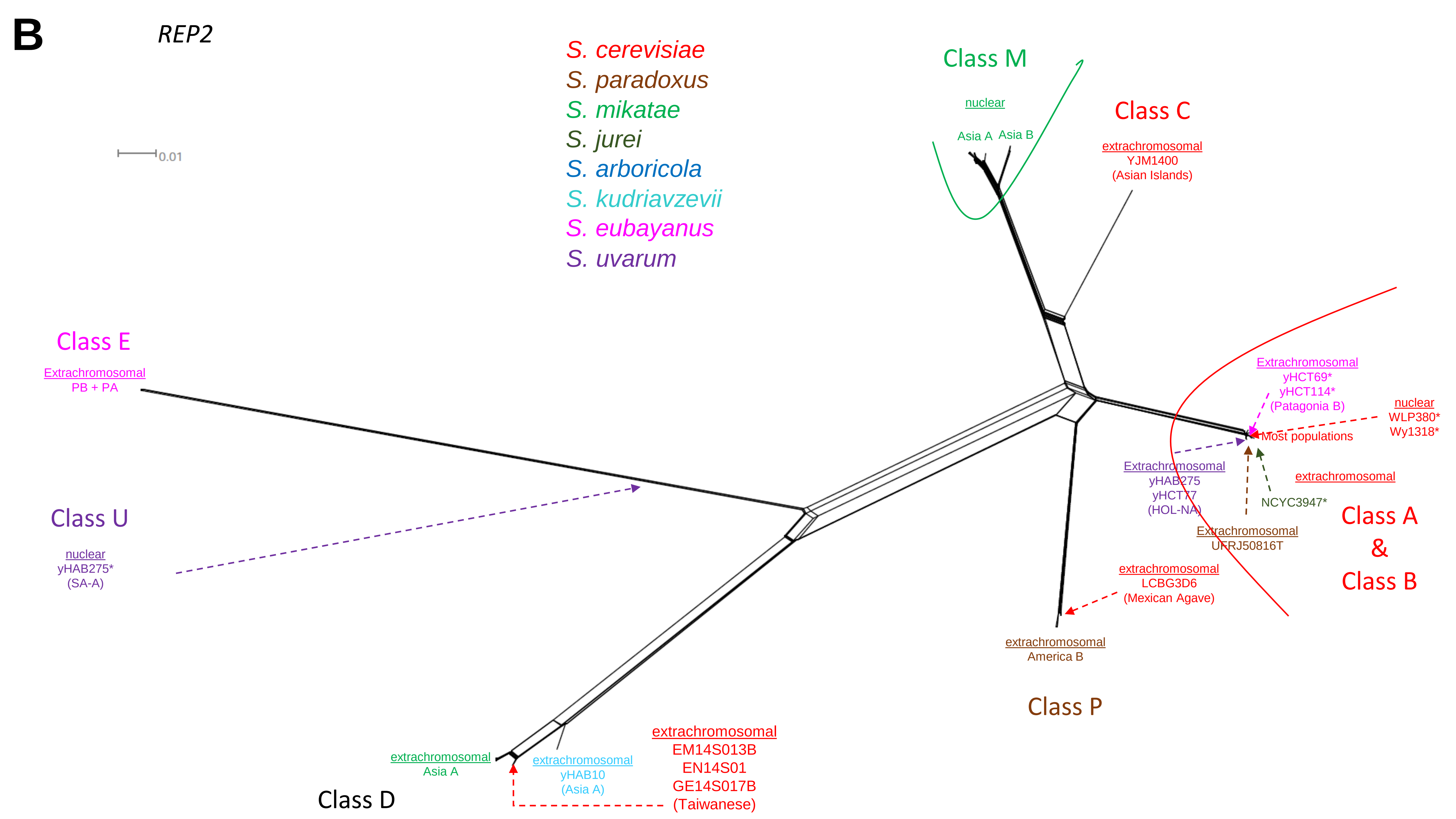

### FigureS18.pdf

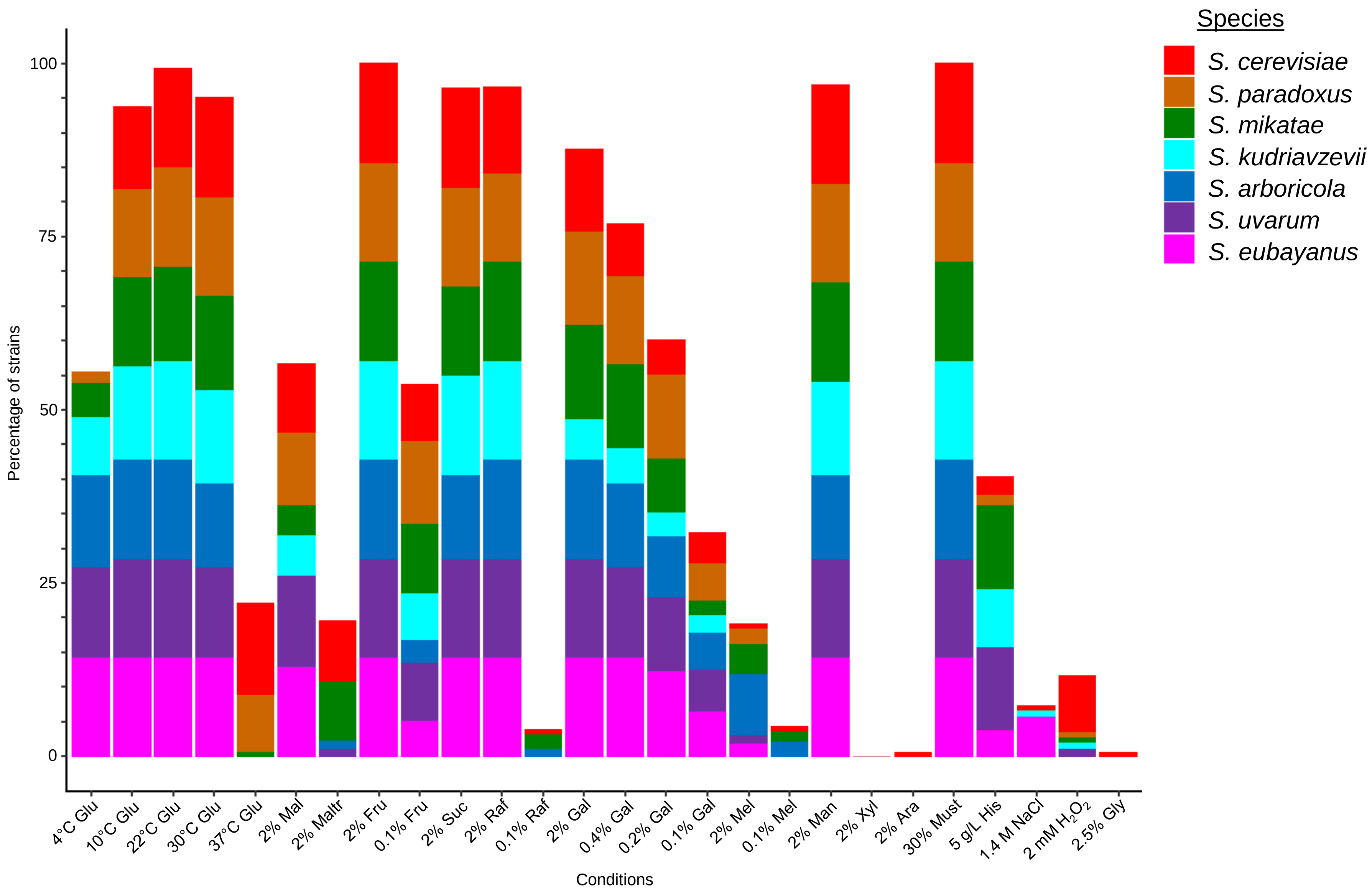

### FigureS19.pdf

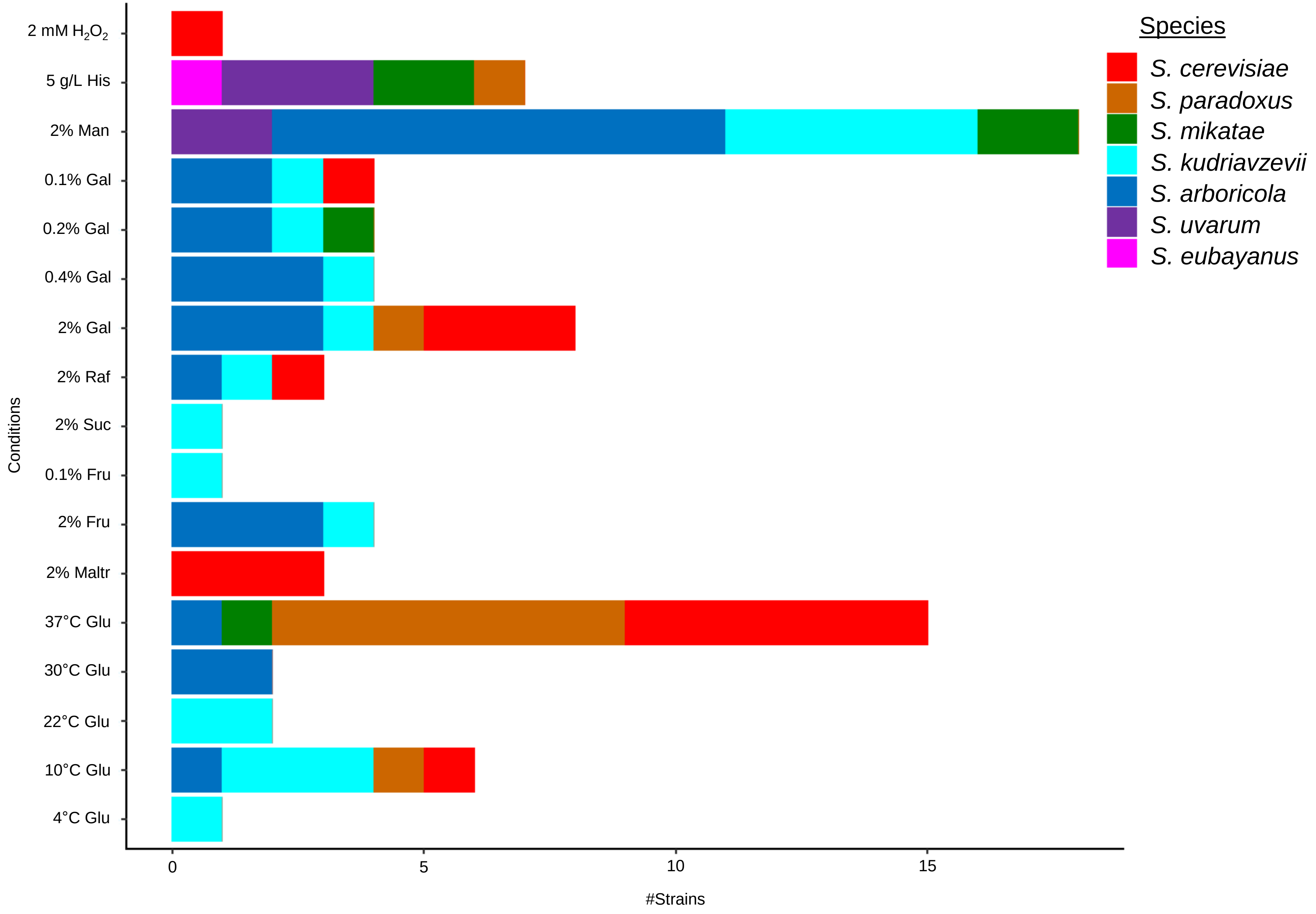

### FigureS20.pdf

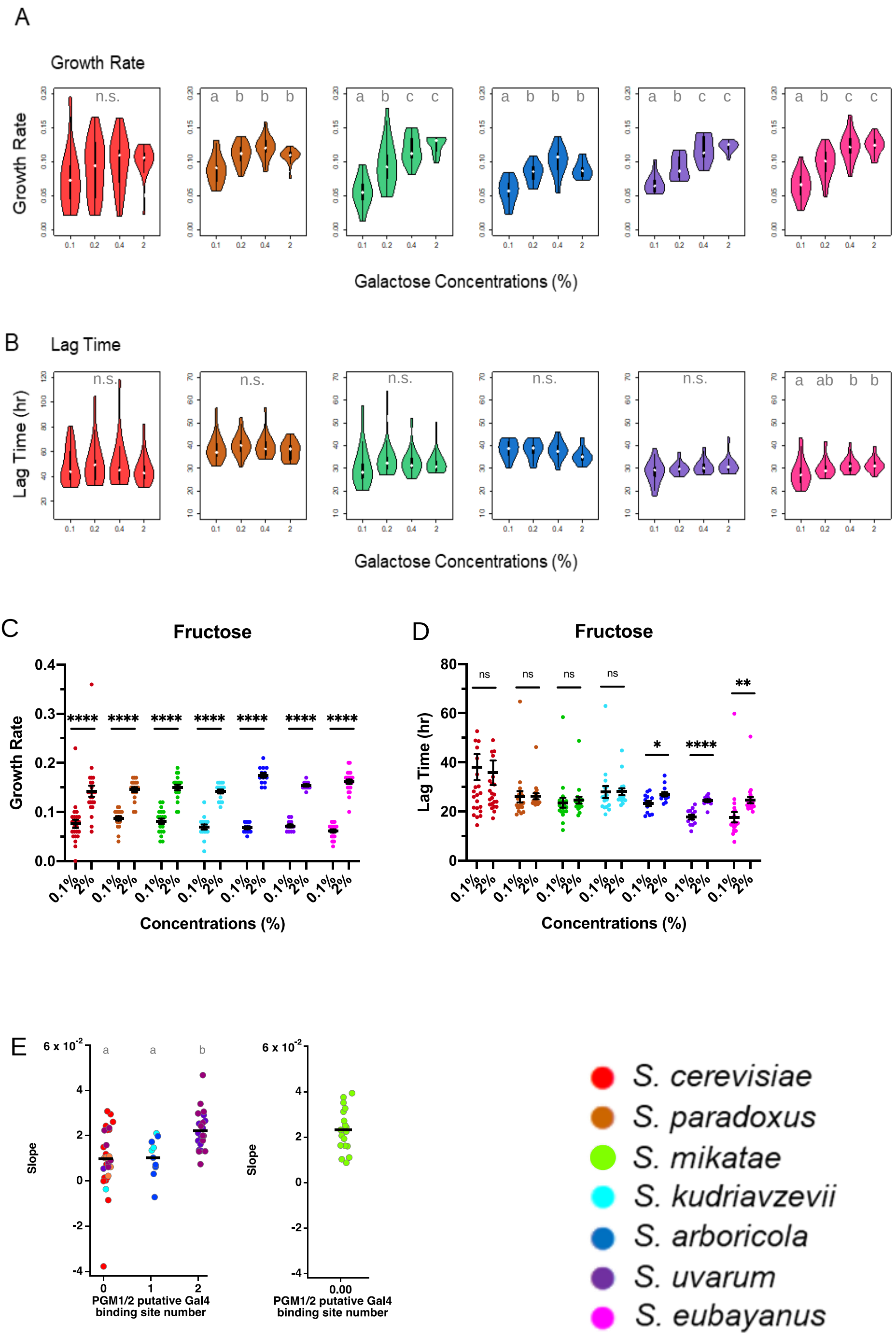

### FigureS21.pdf

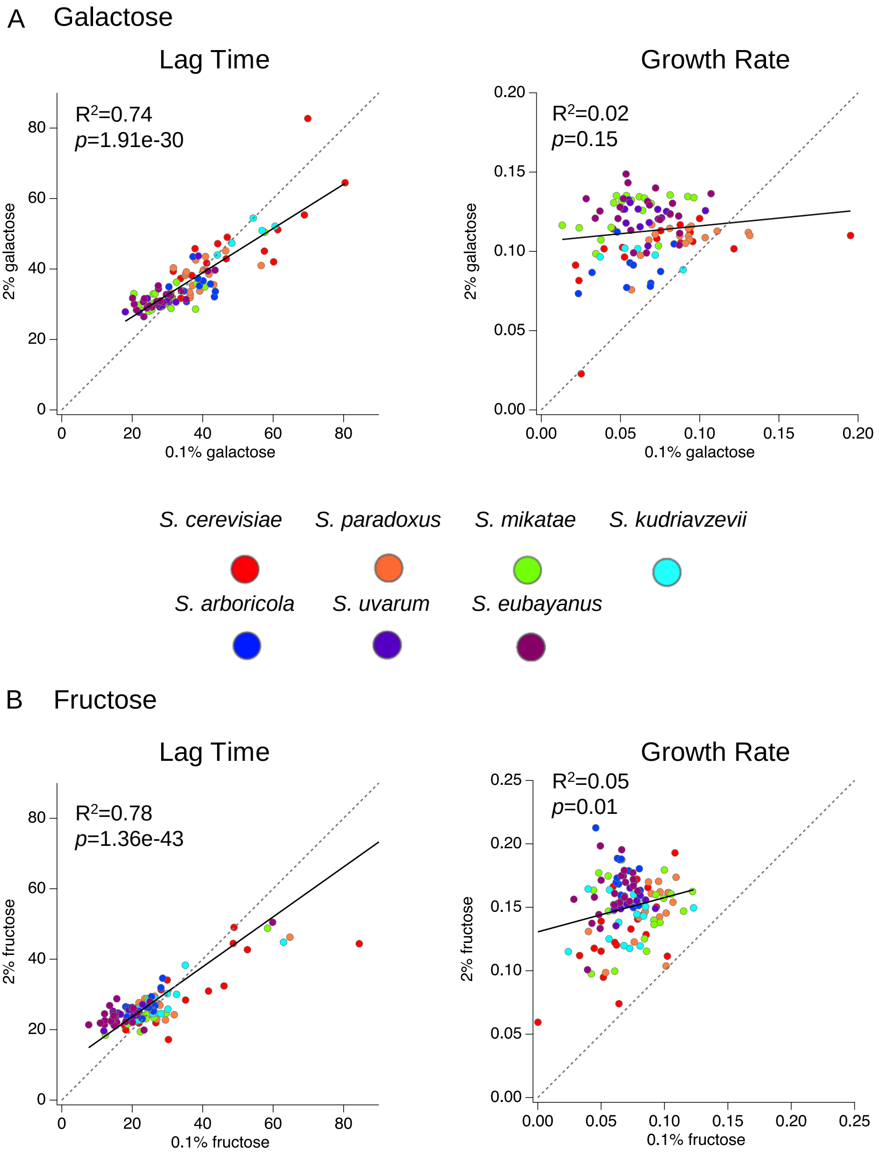

### FigureS22.pdf

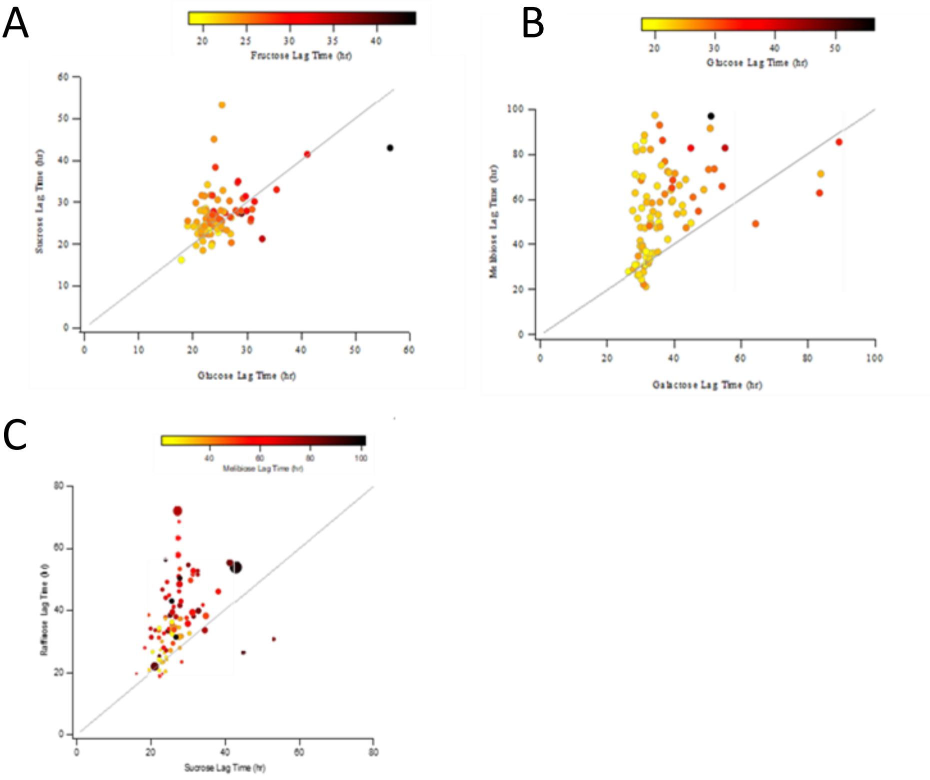

### FigureS23.pdf

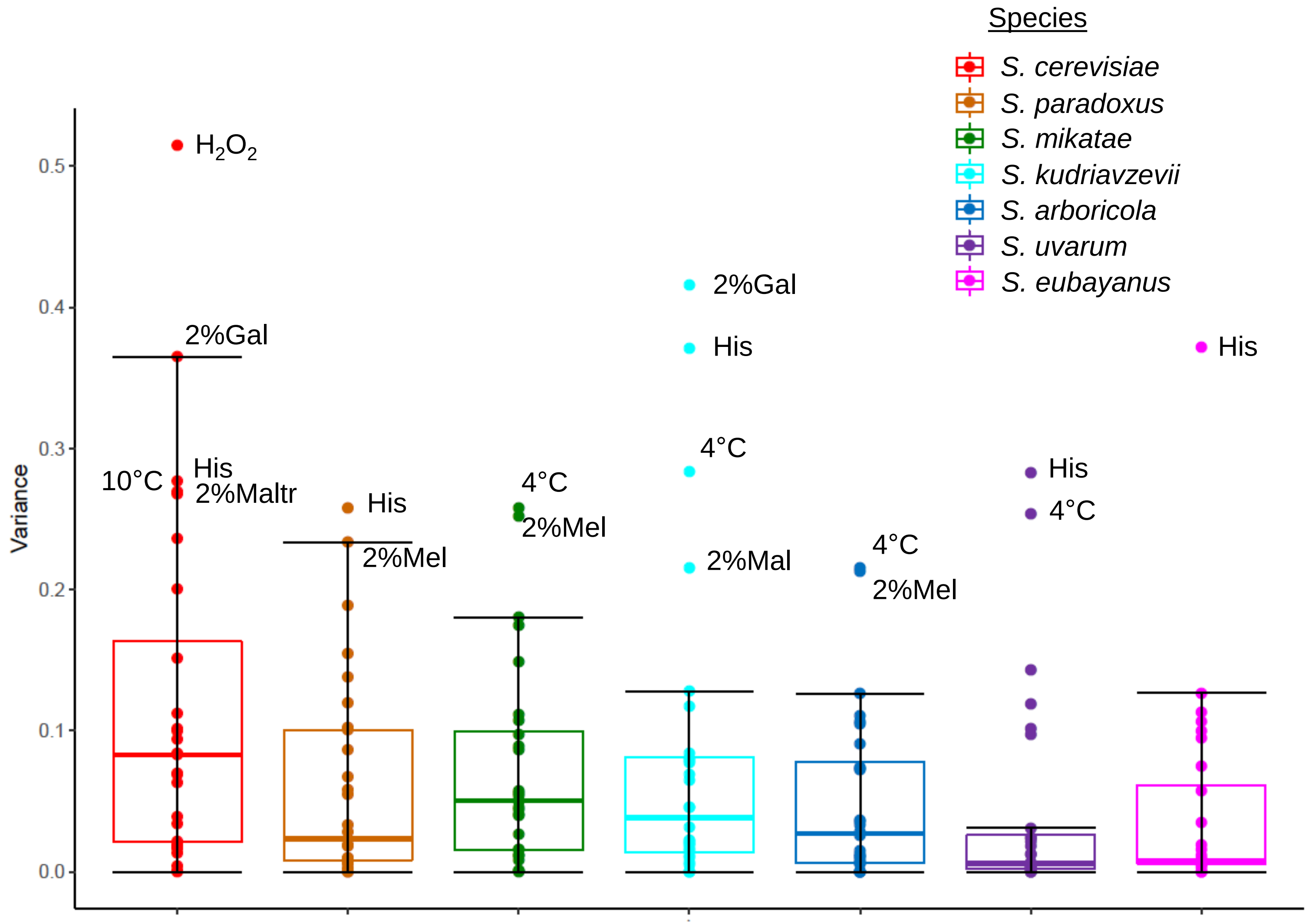

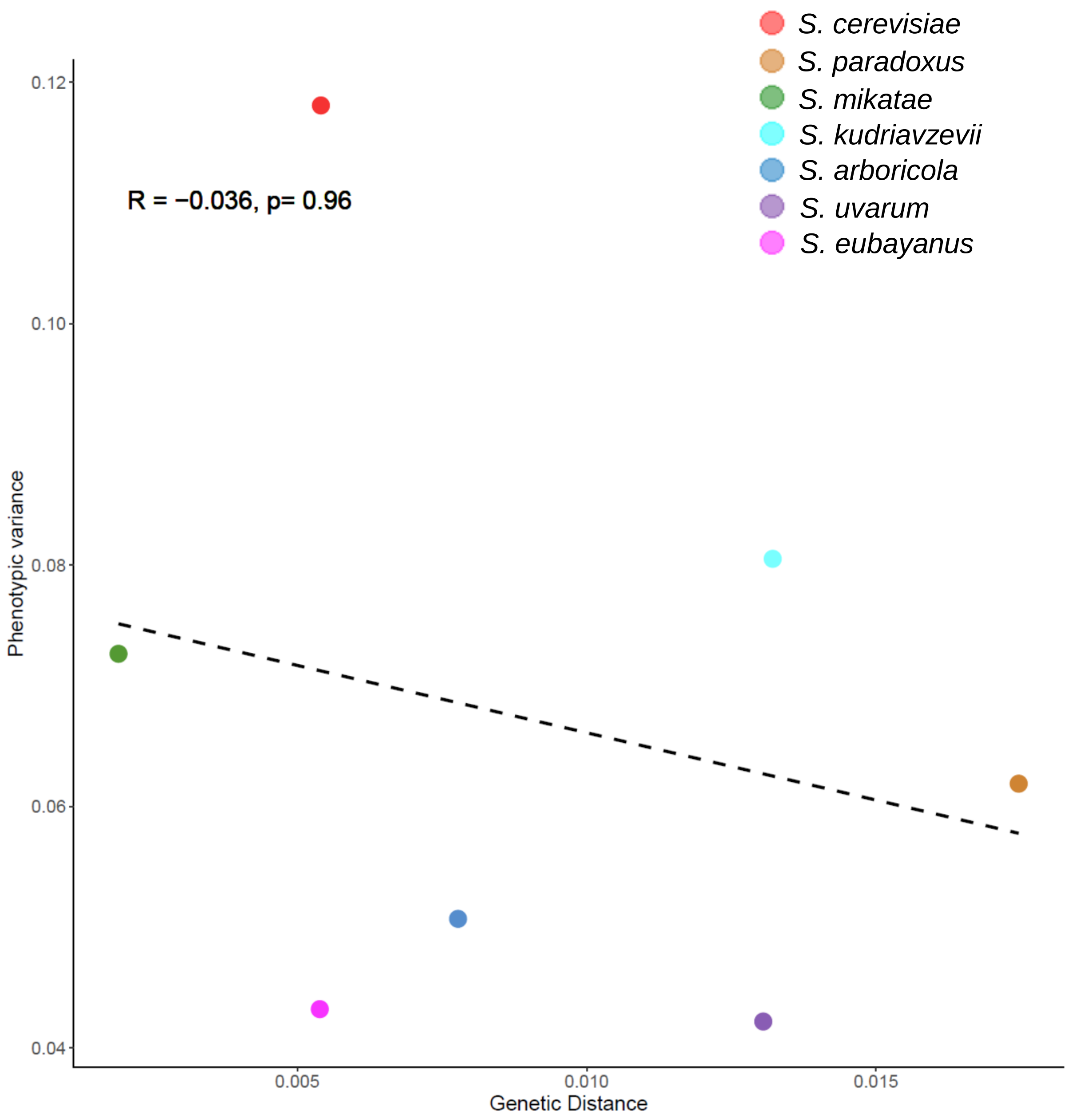

### FigureS24.pdf

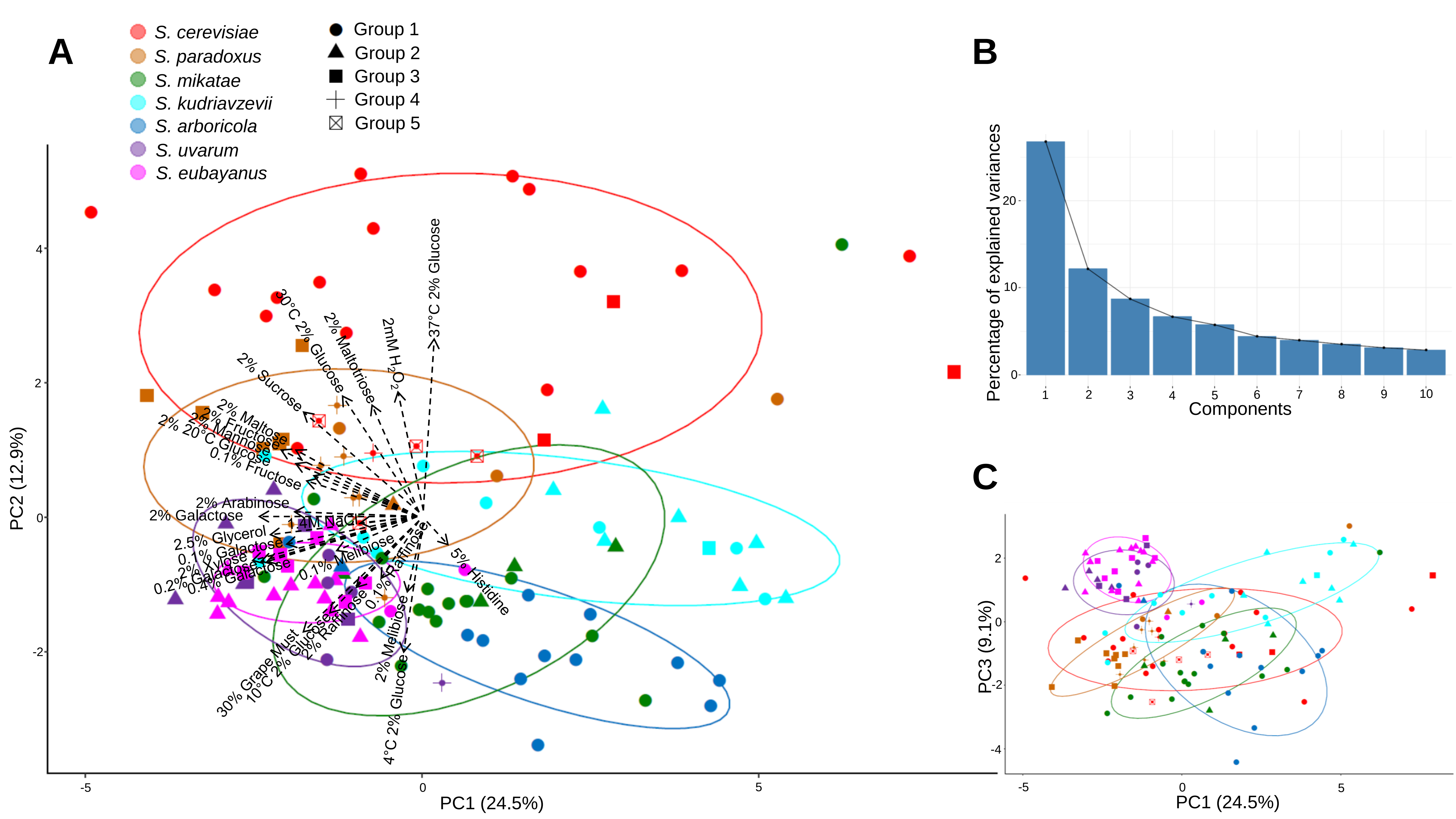

### FigureS25.pdf

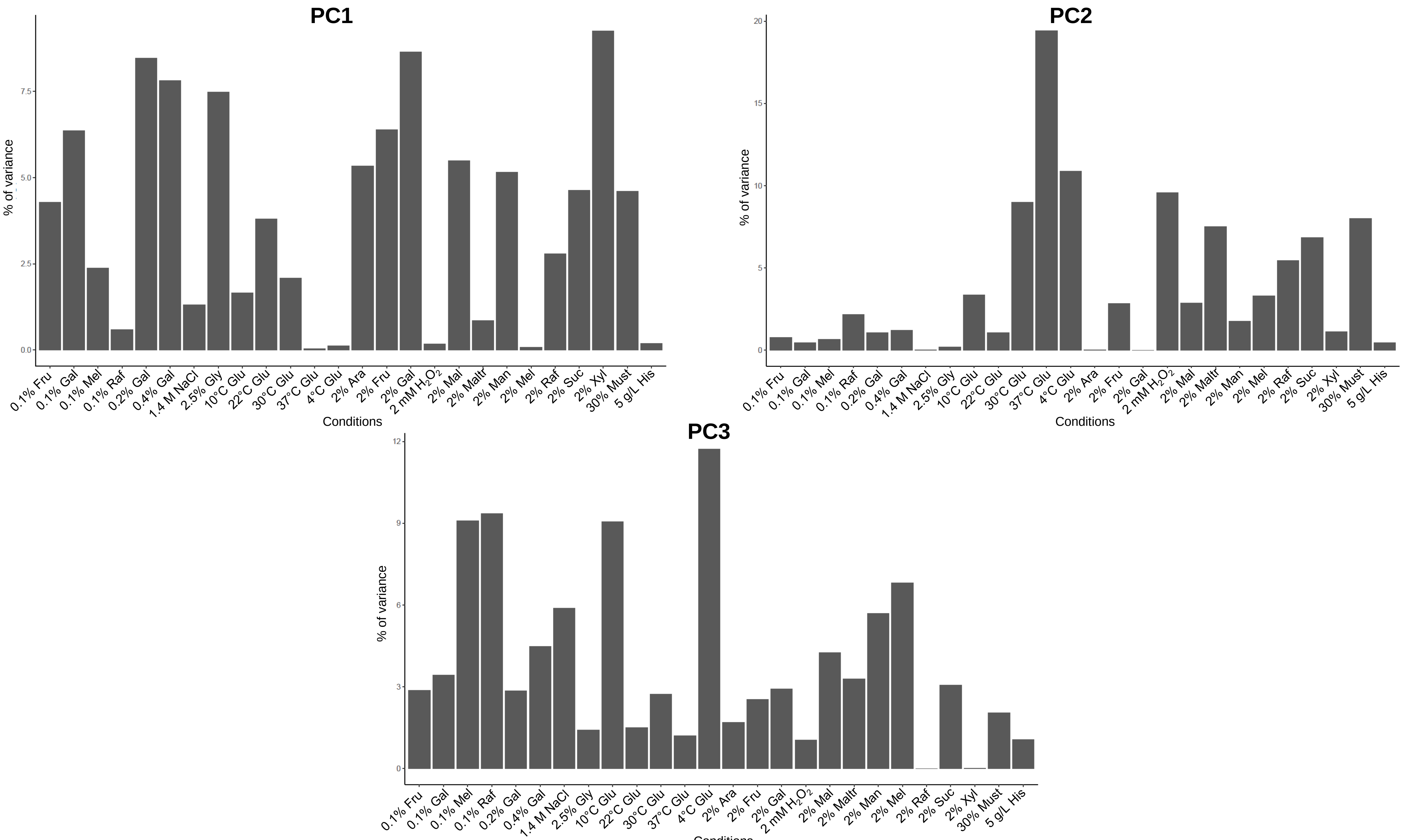

### FigureS26.pdf

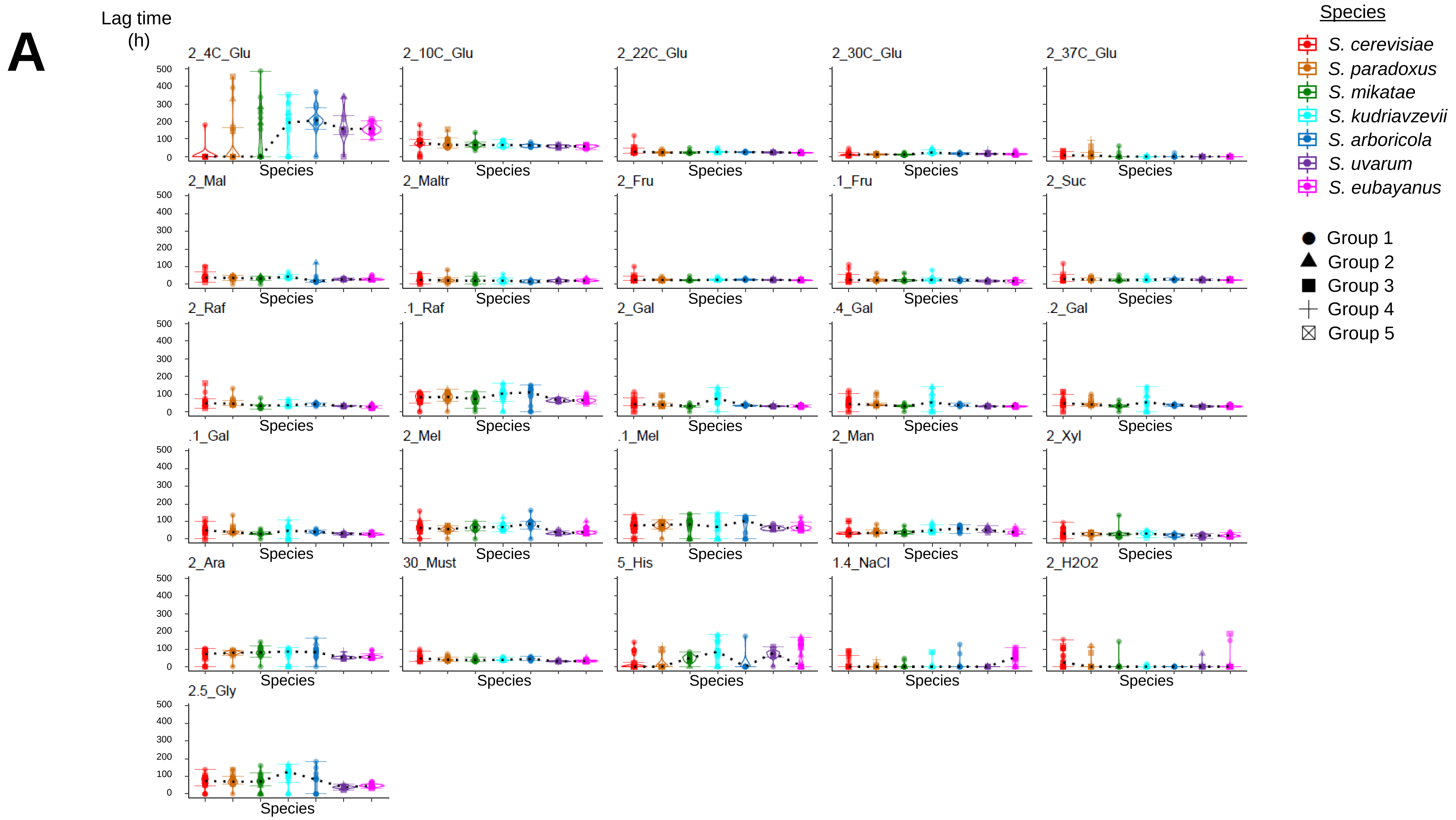

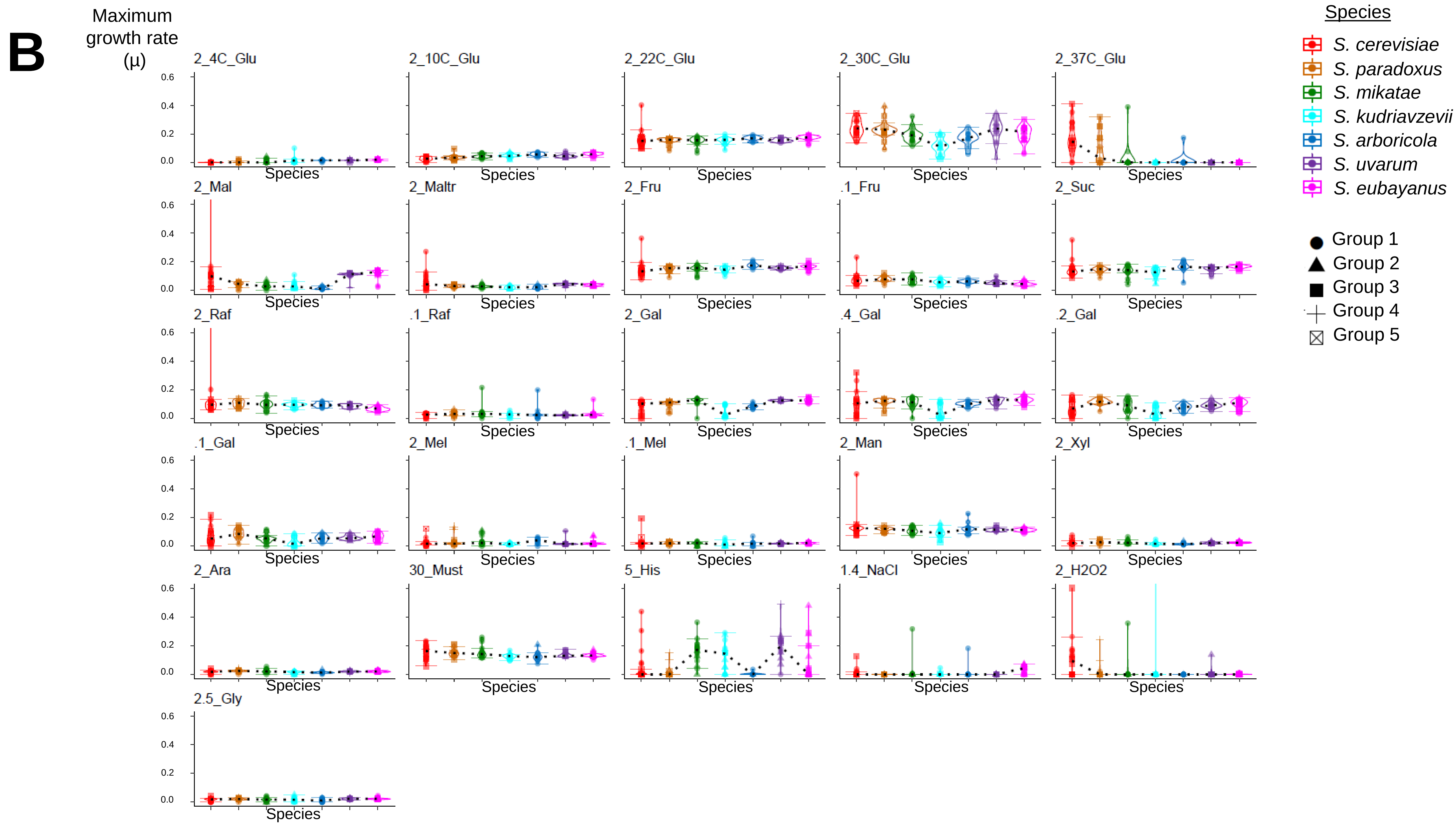

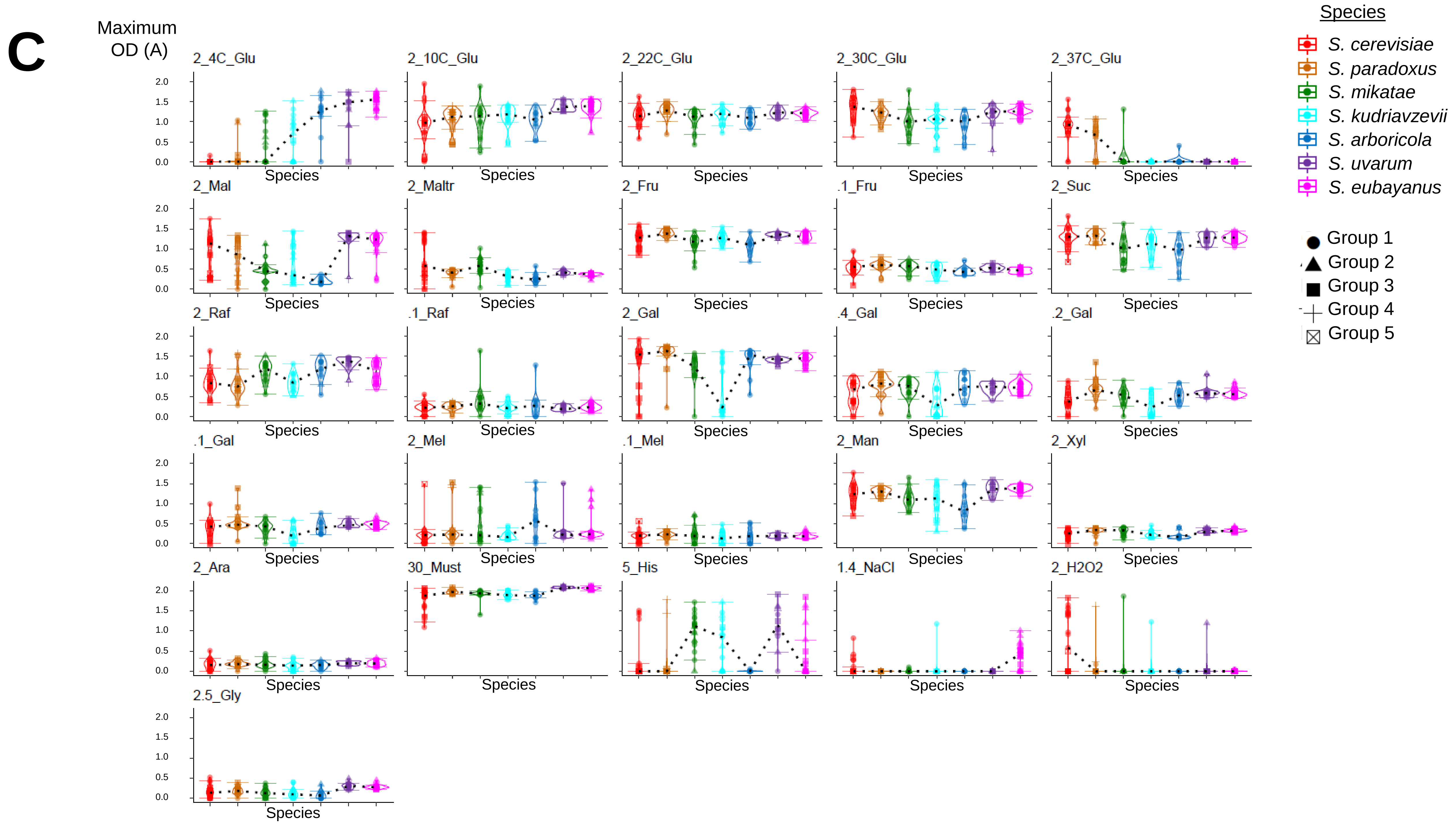

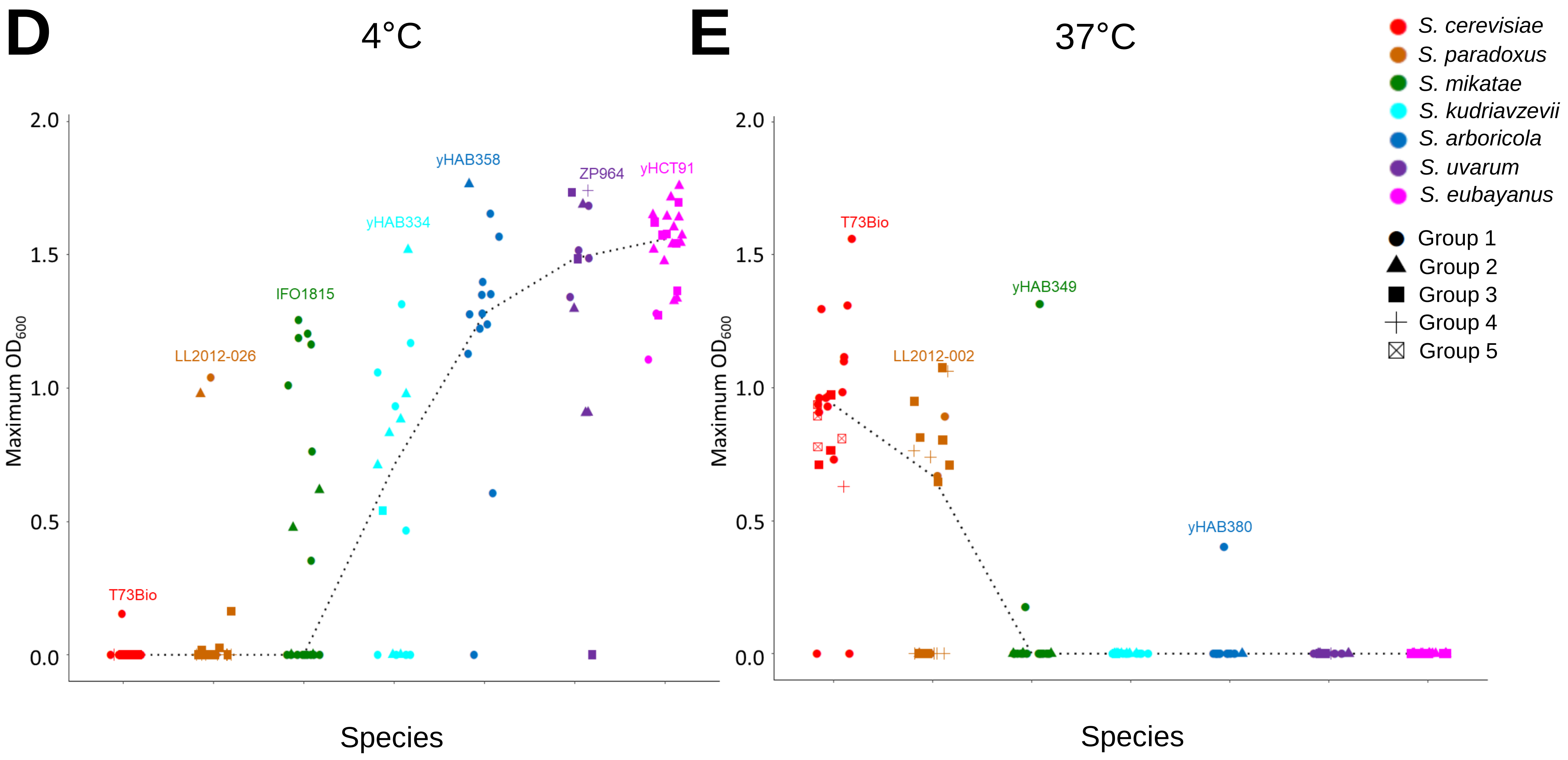

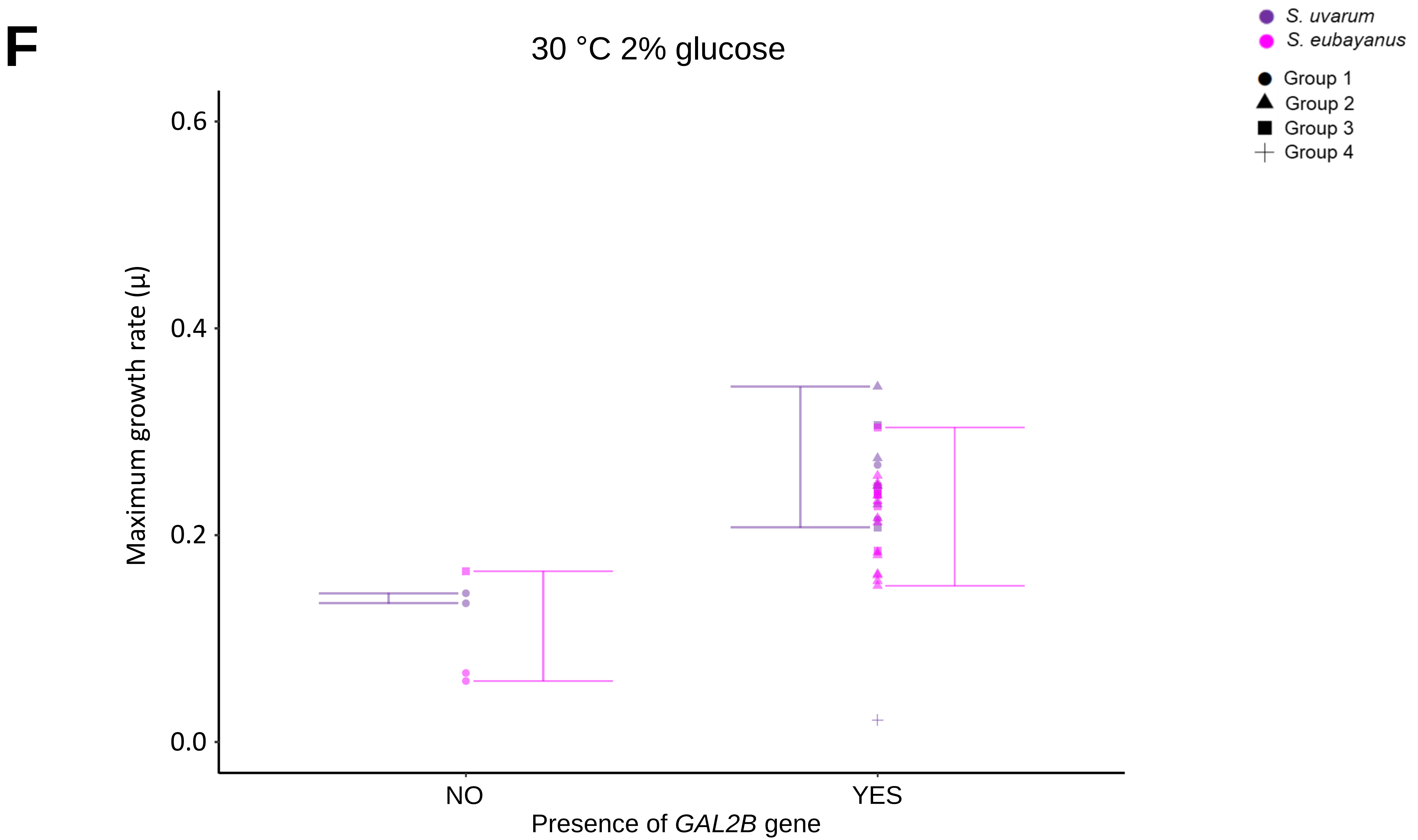

### FigureS27.pdf

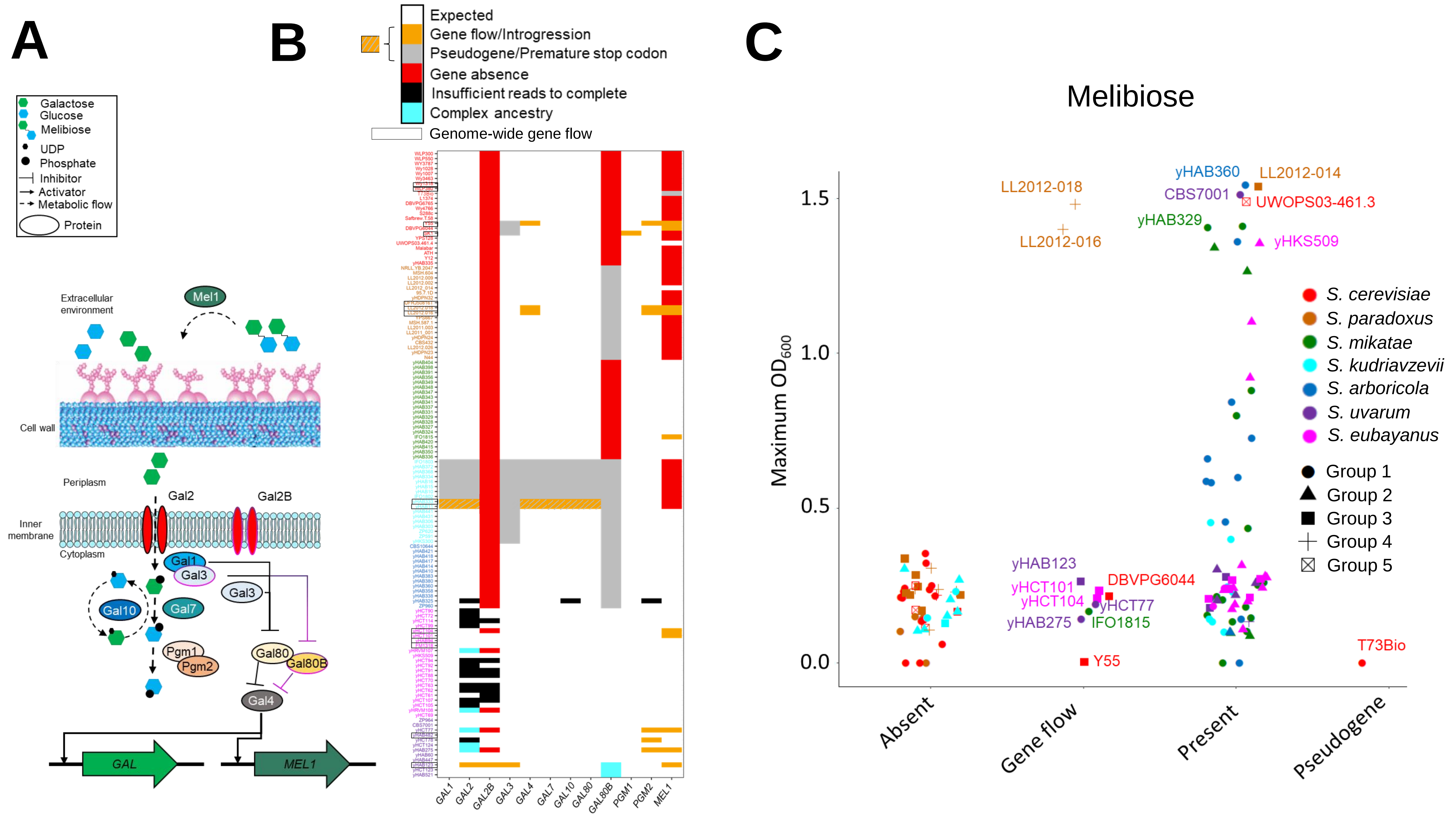

### FigureS28.pdf

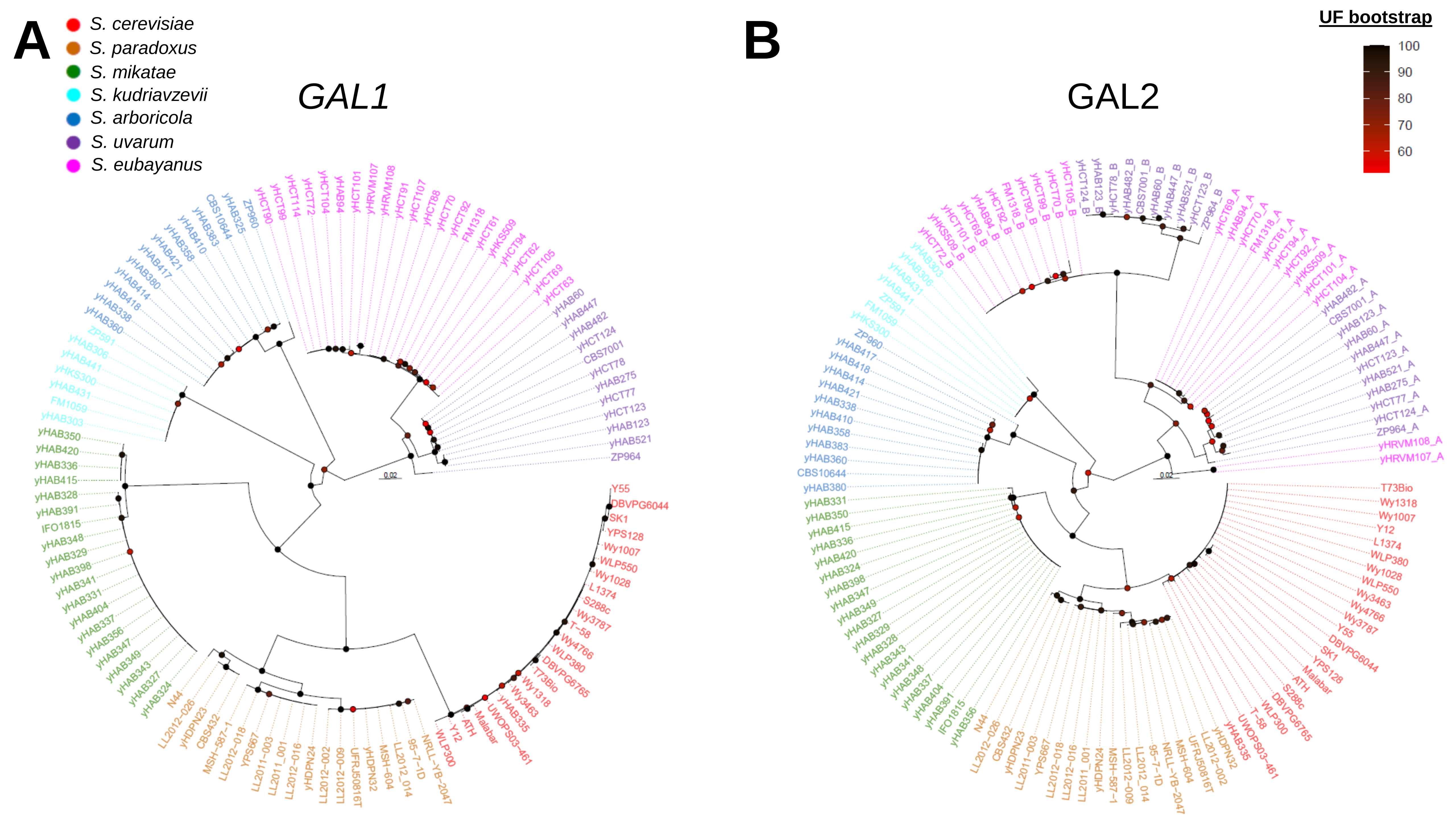

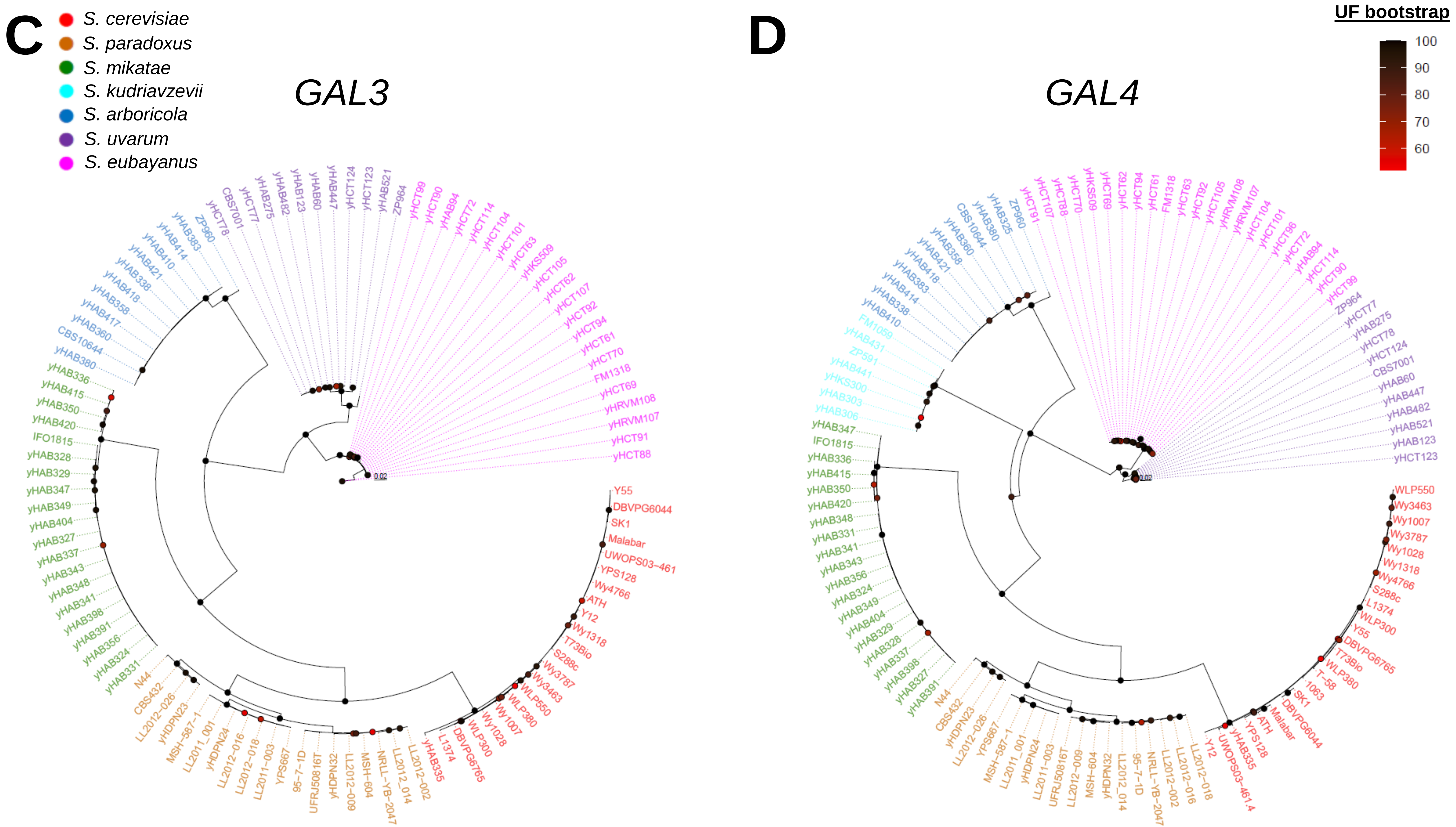

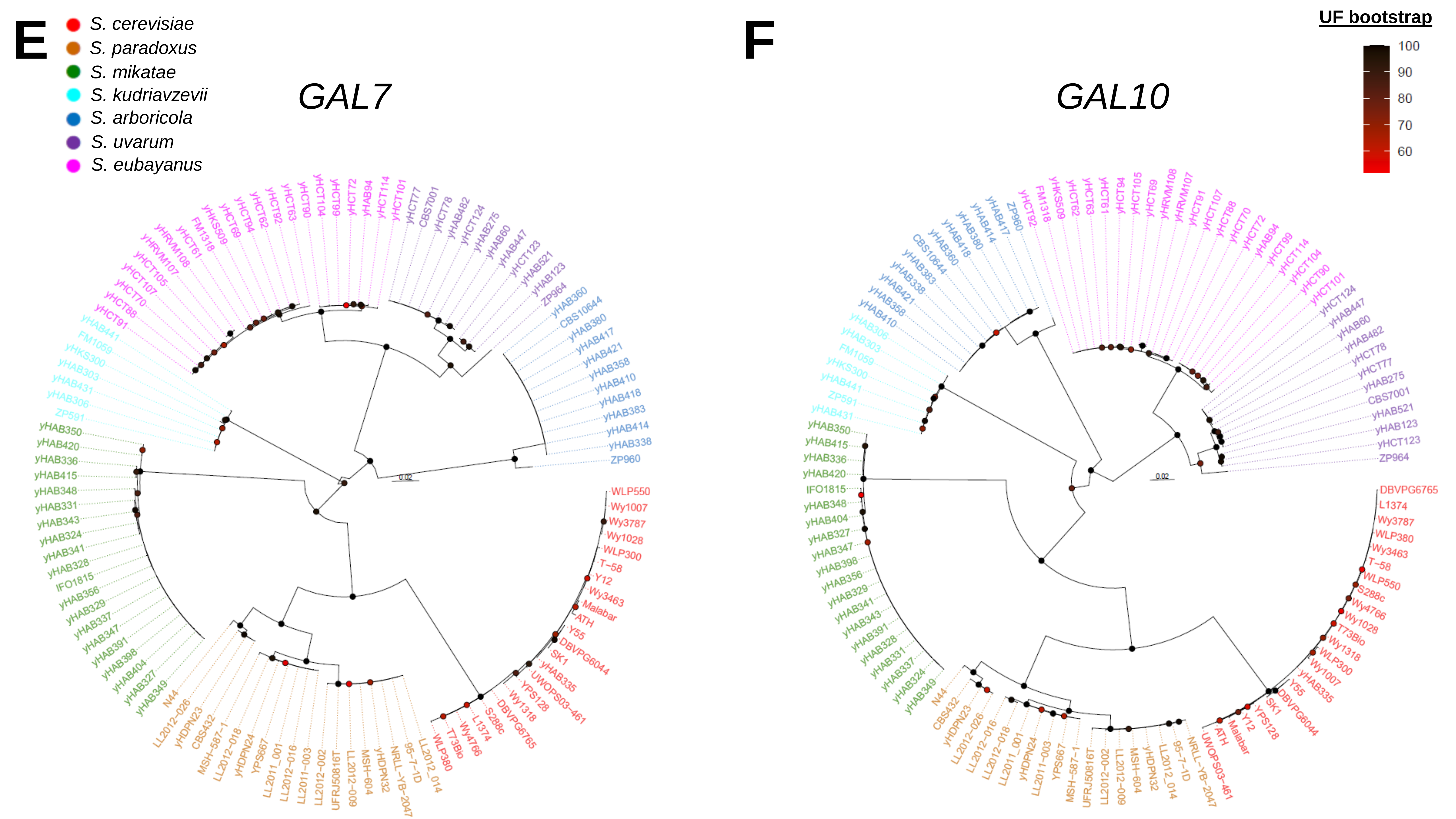

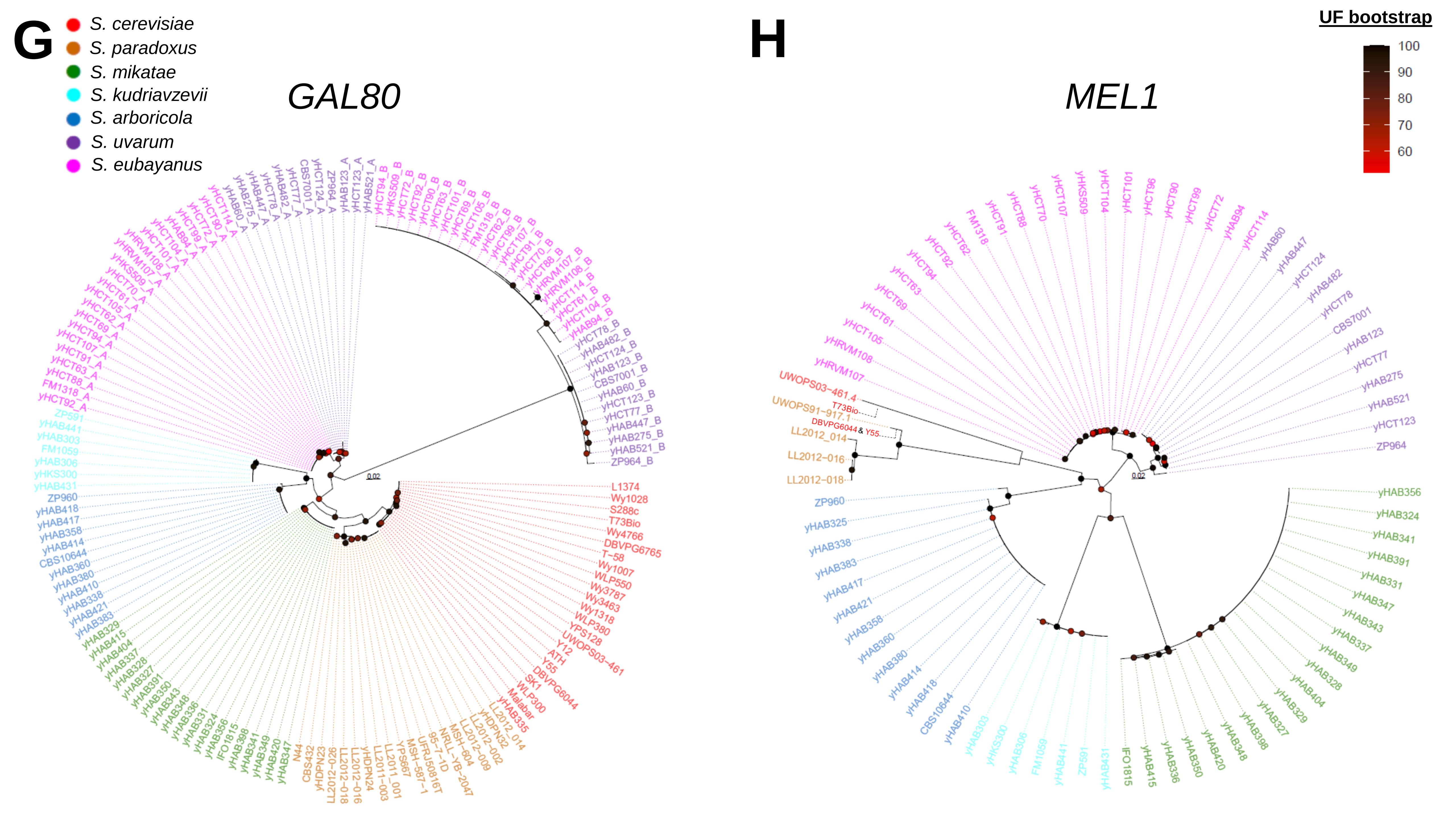

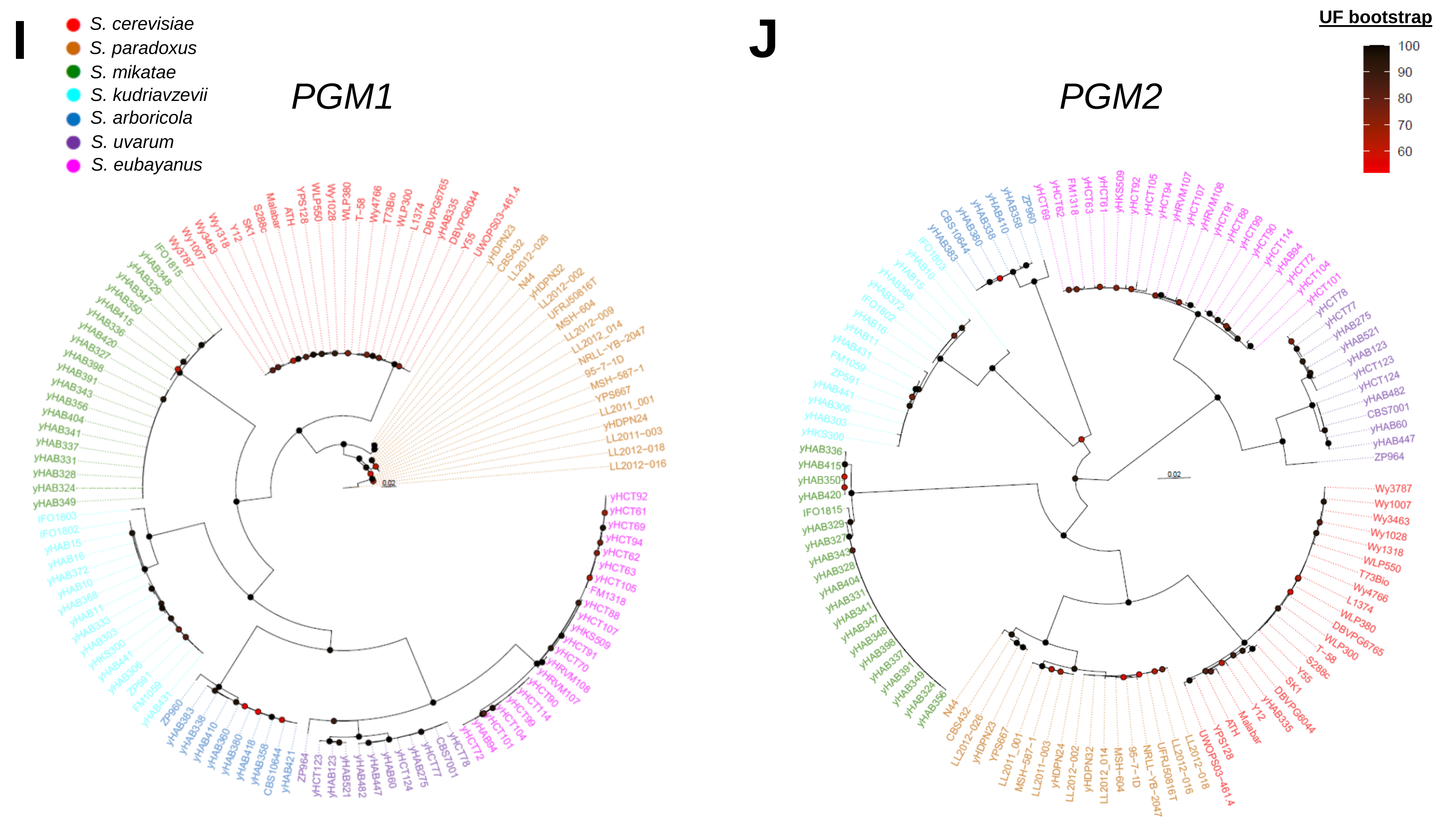

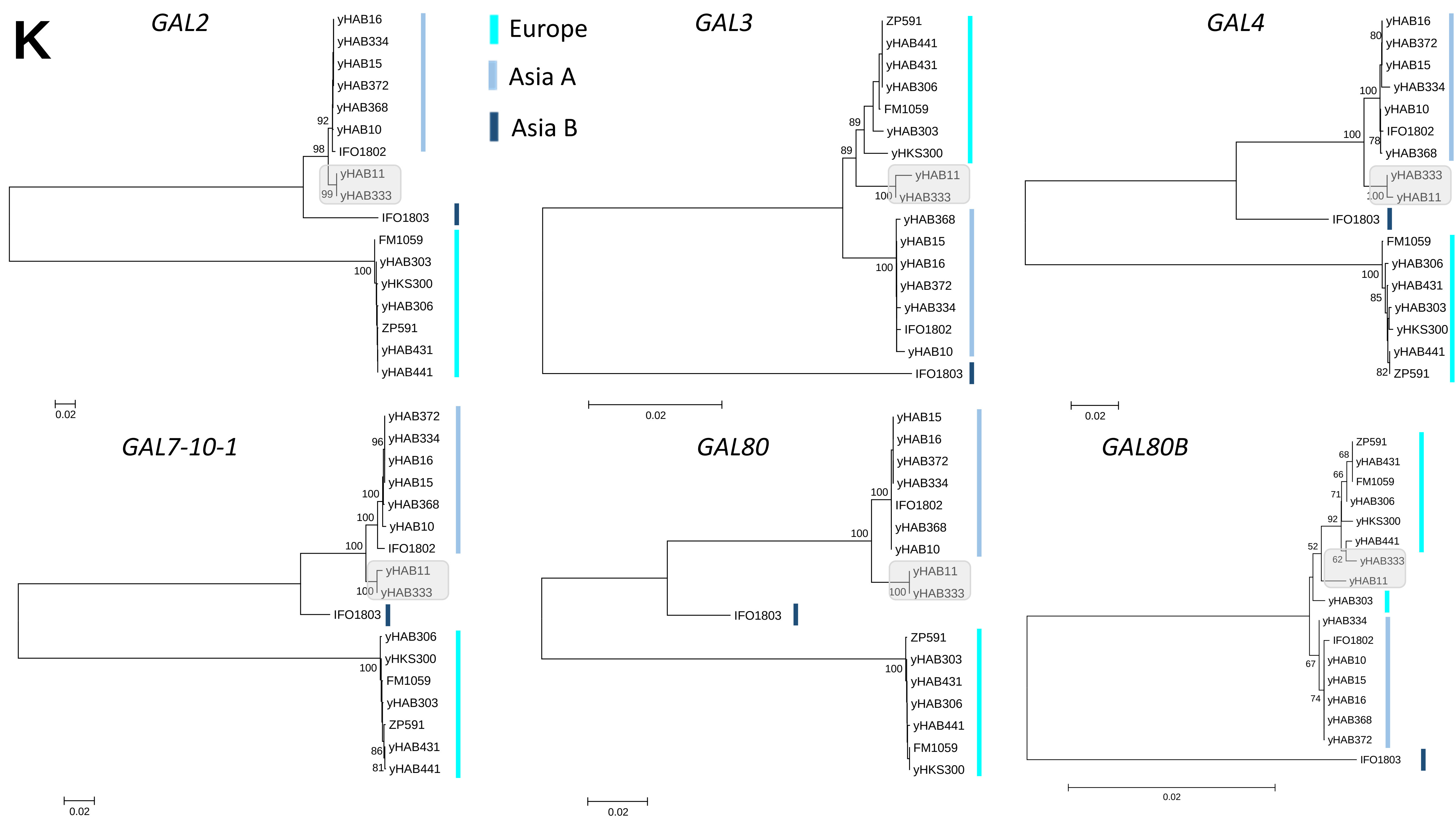

### FigureS29.pdf

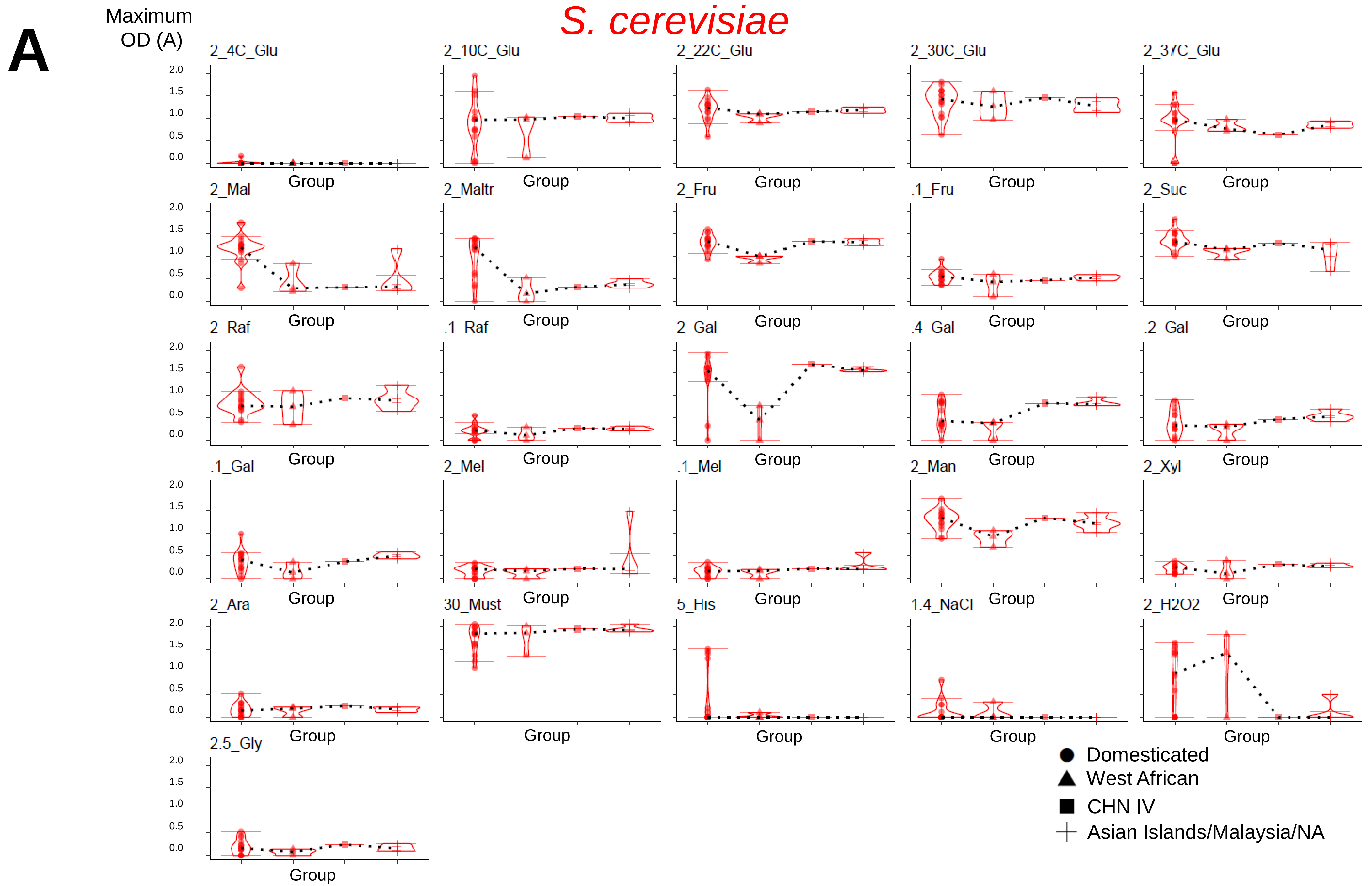

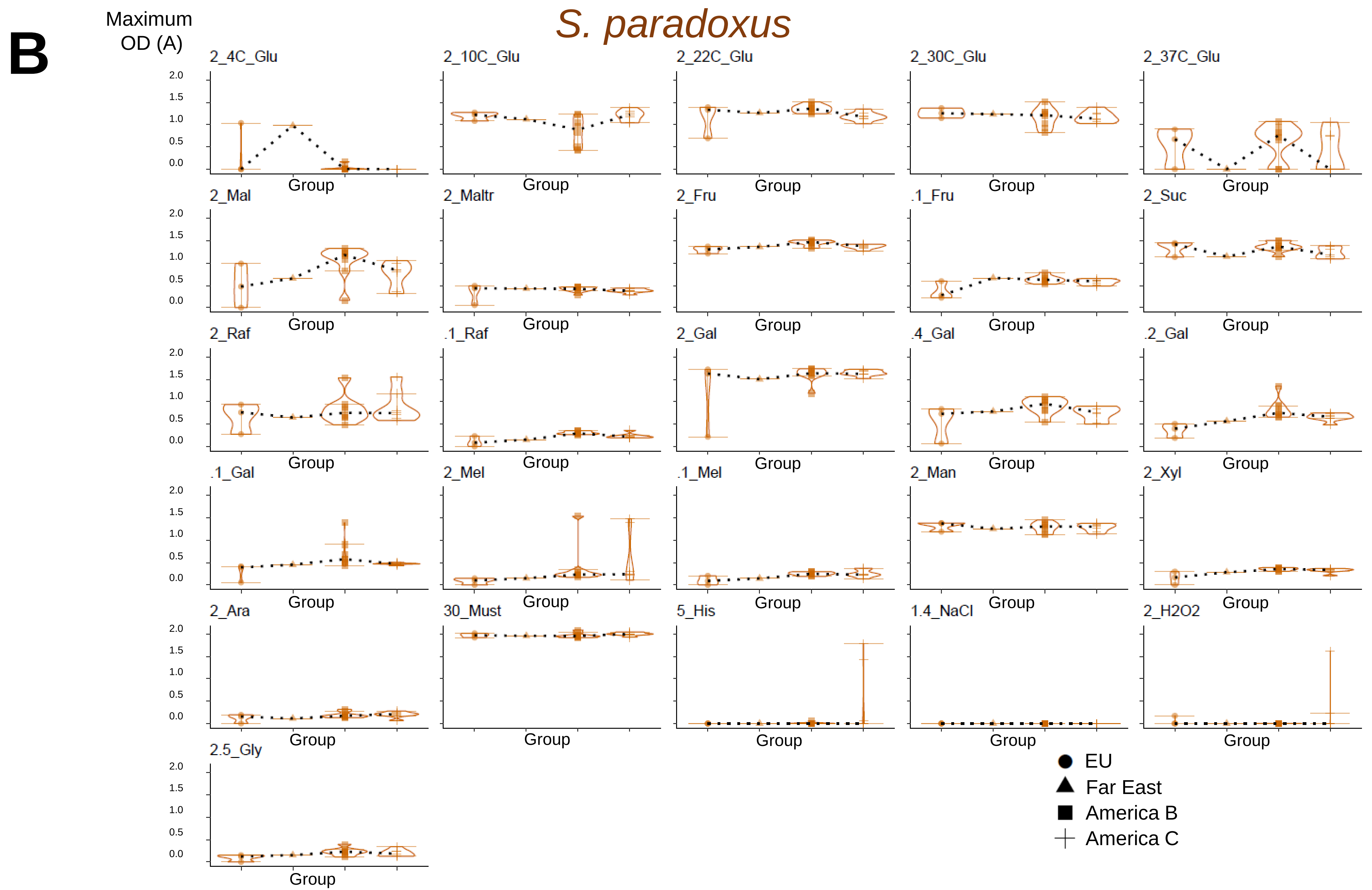

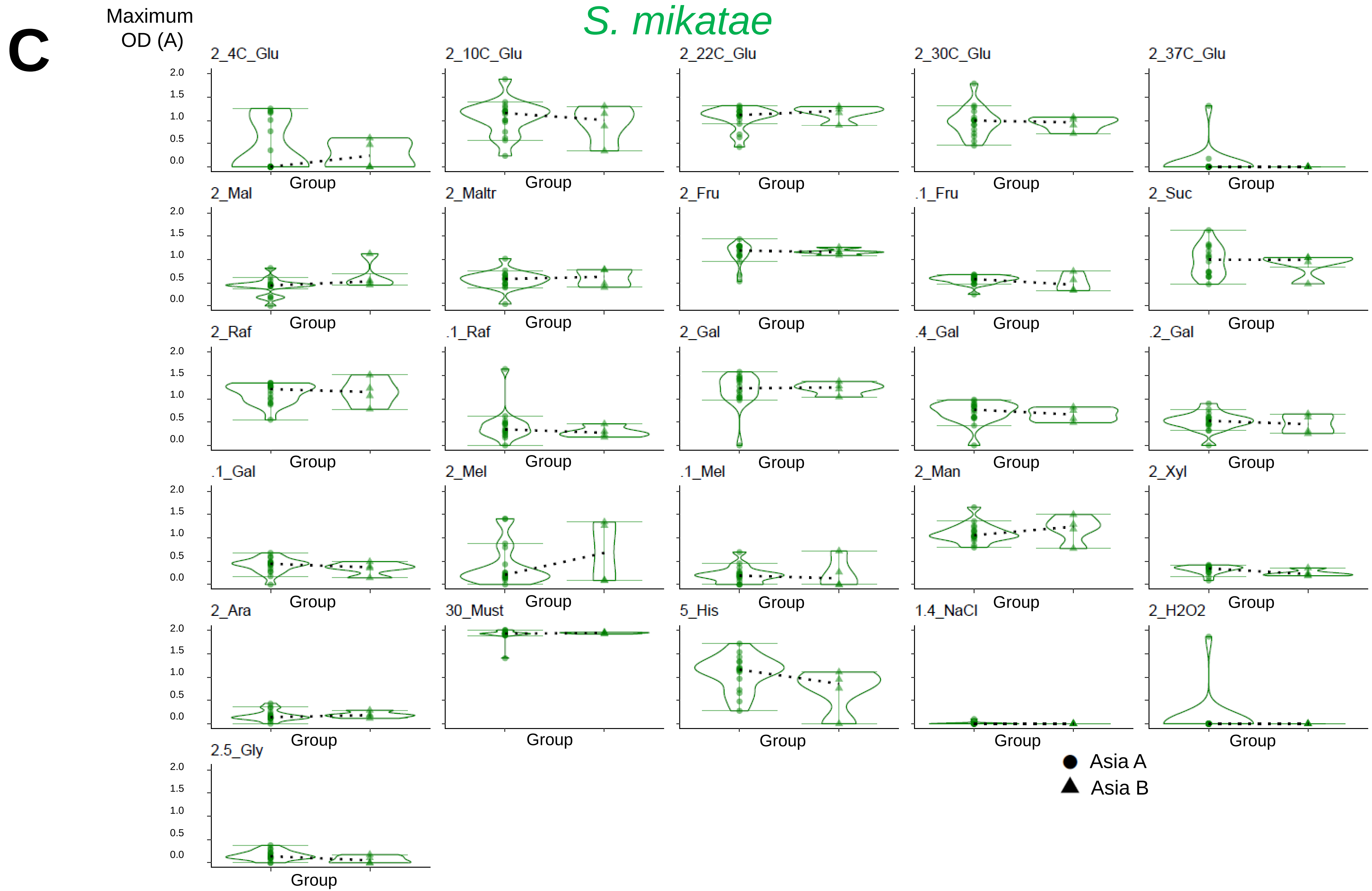

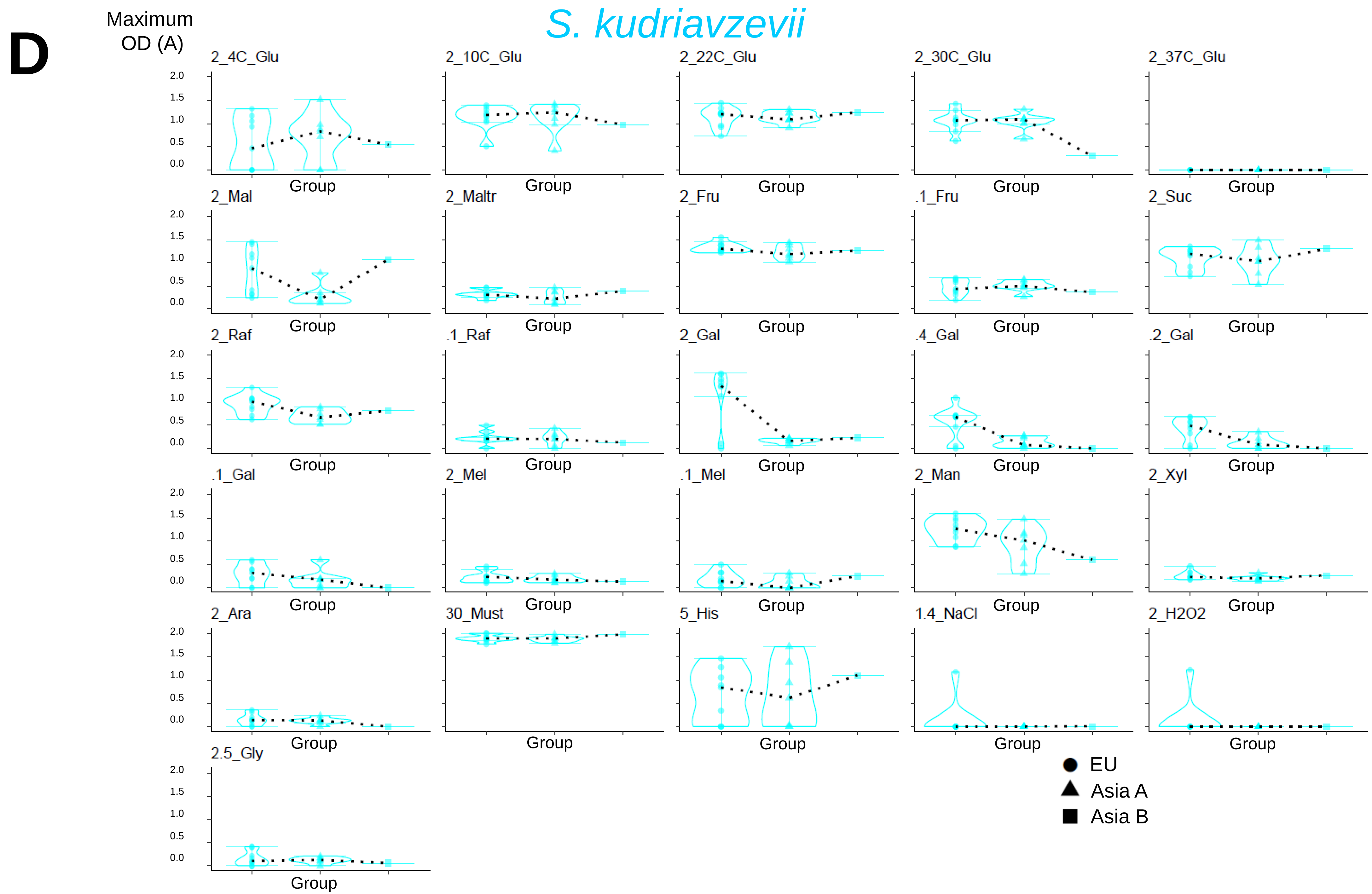

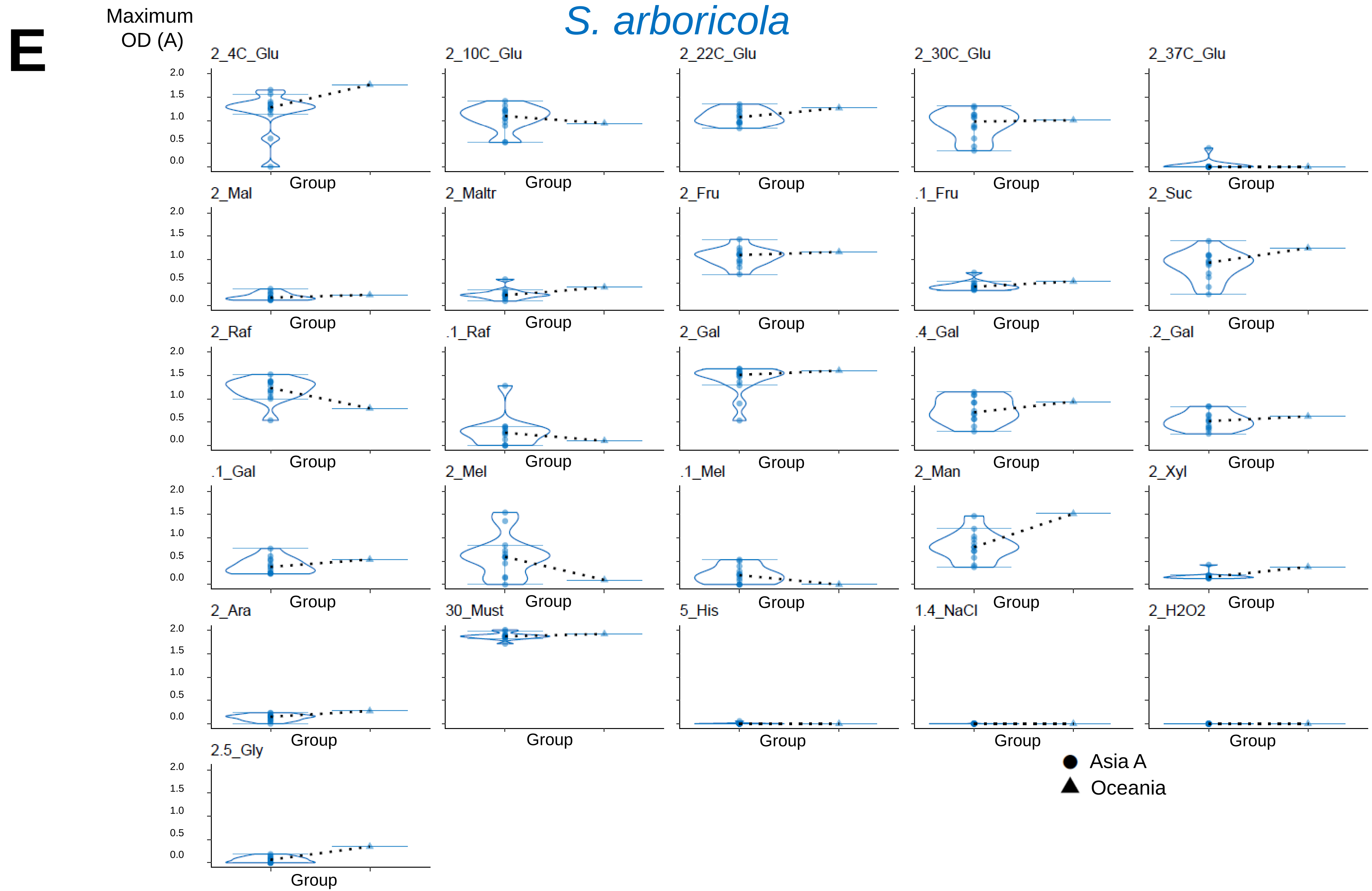

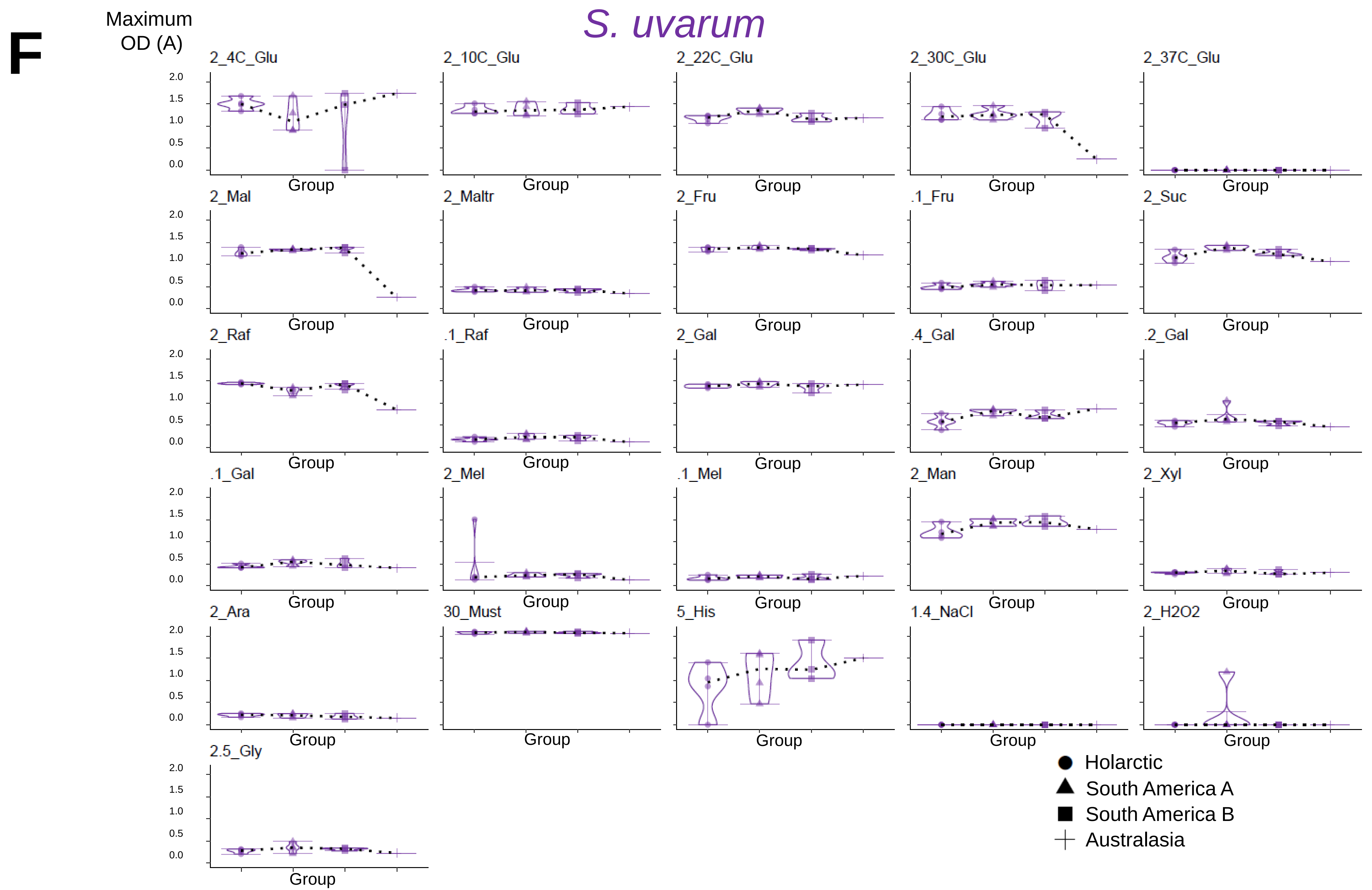

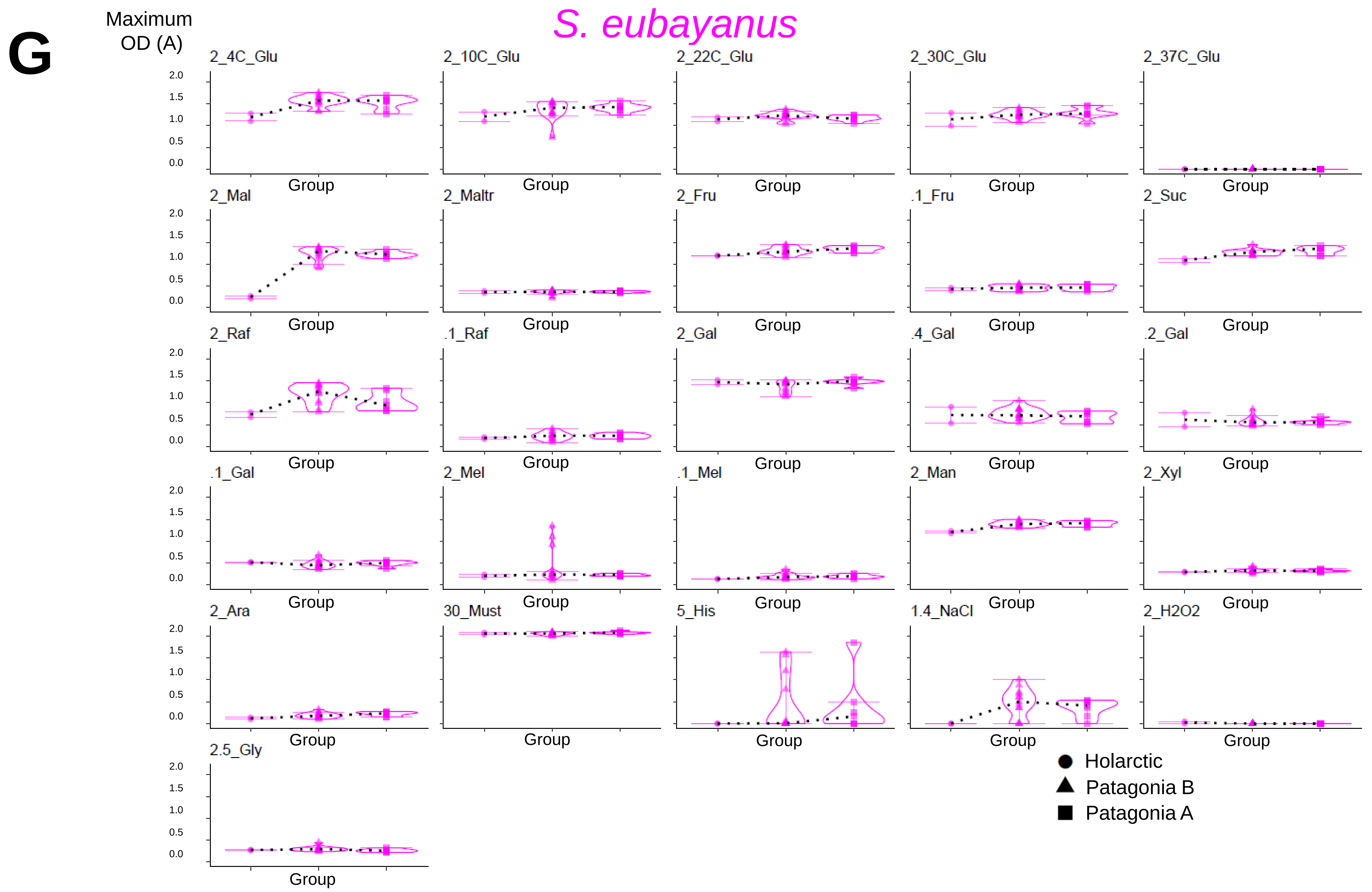

### FigureS30.pdf

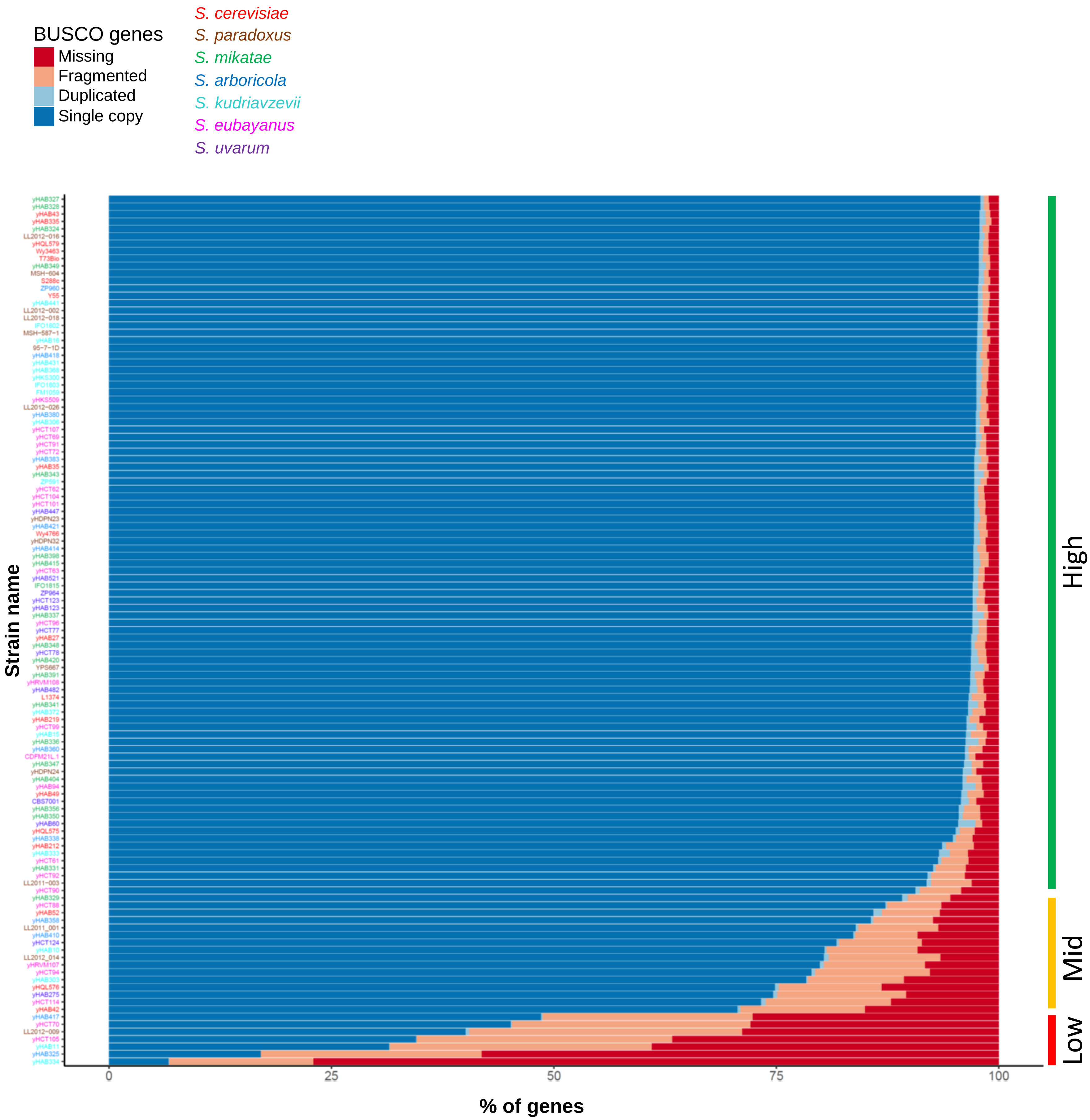
